# Membrane bound geranylated RNAs establish a primitive peptide synthesis system

**DOI:** 10.1101/2024.08.02.606298

**Authors:** Chun-Yin Chan, Johannes Singer, Thomas Carell

## Abstract

The origin of an RNA-based translation system necessitates a specific interaction of certain RNAs with defined amino acids. This must have happened in a protocellular environment where these molecules were concentrated so that a connection between the encoding RNA and the amino acids could be established that allowed the formation of peptides. A model of how such a system could have evolved does not exist. Here we show that geranylated non-canonical nucleotides that are potential fossils in an early RNA world, allow RNA to anchor to lipid membranes. This creates RNA-geranylating lipids on which a primitive peptide synthesis can then operate with rudimentary chemoselectivity. The system creates a protocellular model of how RNAs and amino acids could have been mutually selected based on their physicochemical properties.

## Introduction

A prerequisite for encoded translation is the transfer of encoded information into a defined peptide sequence. This requires as a first step the selective charging of a unique RNA (transfer RNA, tRNA) with a specific amino acid. Today, this is performed by specialized amino acid-tRNA synthetases (aaRS). These amino acid-charged RNAs need then to selectively base pair with the complementary encoding RNA sequences (messenger RNA, mRNA) to allow the sequential assembly of amino acids to generate a peptide with a sequence of amino acid that corresponds to the mRNA sequence.

Today, this highly sophisticated process happens at the ribosome, which is arguable one of the most complex machines present in cells. It unites the mRNA with the amino acid-loaded tRNAs and it catalyzes the peptide bond formation step. This whole process translates genetic information into specific proteins, which is a hallmark of all life on Earth. However, the ribosome itself is unable to recognise tRNAs with a mischarged amino acid^1,2^ apart from their chirality.^3^ Hence, it is the aaRS-catalyzed charging of the tRNA adapters with specific amino acids which creates the link between a specific RNA code and a defined amino acid.

One of the greatest mysteries of the origin of life is how this complex machinery evolved in a primitive, primordial RNA world about 4.2 billion years ago. A plausible concept attempting to explain the origin of translation suggests that functionalized RNAs learnt at some point to colonize liposomes, which may have established an environment that could concentrate RNA and amino acids to enable the co-evolution of specific peptides and encoding nucleic acids.^4,5^ This scenario, however, requires the highly negatively charged RNA to phase separate from the aqueous environment onto liposome surfaces. In this context, it was found that RNA can be lipidated under prebiotic conditions in the presence of cations,^6,7^ fatty acid modifications^8^ or in association with specific peptides.^9,10^ This all provides an unspecific lipidation of RNA for membrane anchoring. Furthermore, all scenarios in which fatty acids are coupled to terminal phosphates present on the RNA strand,^8^ limits the system to one fatty acid modification per RNA end, which greatly limits the membrane-anchoring capabilities, particularly for longer oligonucleotides.

Others^11–14^ and us^15–17^ introduced recently the concept that non-canonical nucleosides present in contemporary RNA could be relics (living fossils) of an early RNA world that have vastly expanded RNA’s capabilities. Analysing the structures of non-canonical nucleotides, particularly of those found within the anticodon stem loop, which is arguably the oldest part of the tRNA-based decoding system,^18^ reveals the presence of *S*-geranyl-2-thiouridine (ges^2^U) and its derivatives. These non-canonical nucleosides are found in the anticodon loops of Lys, Glu and Gln tRNAs.^19^ With the knowledge that the membranes of archaebacteria are composed of structurally similar isoprenoid-based lipids,^20–22^ we thought that ges^2^U bases could have been the missing link for early RNA systems to transition into protocells.

## Results

### Geranylated RNAs phase separate onto liposomal membranes

To explore whether ges^2^U can anchor RNA onto lipid surfaces, we first explored methods to generate this non-canonical nucleotide in RNA. We thought that for a prebiotically more plausible scenario it would be desirable to achieve this in a post-synthetic manner by first incorporating the prebiotically plausible base, s^2^U (ref.^23^), into RNA, followed by its geranylation (see Ext. Data Table 1 for information of all RNAs). For initial studies, we prepared first the homo-U RNA strand 5’-U_4_(s^2^U)_2_-3’ (ON2-2S), containing two s^2^U nucleosides and post-synthetically geranylated it.^24^ We used as a model for a geranyl-electrophile, geranyl bromide (geBr) as depicted in Fig. 1a.^24^ Indeed, this procedure allowed the efficient formation of the geranylated RNA strands 5’-U_4_(ges^2^U)_2_-3’ (ON2-2geS) in about 50 % yield (see Methods and Fig. S1). Comparing it with the result of direct incorporation of ges^2^U phosphoramidite to form 5’-U_5_(ges^2^U)-3’ (ON1-geS), this method avoided the potential conversion to *iso*-(m^2^)C and as such appears to be the best method for multiple ges^2^U incorporations (Fig. S2).^24^ We therefore synthesized all ges^2^U-RNAs in this study using this post-modification method (Fig. S3-S14).

**Figure 1.**
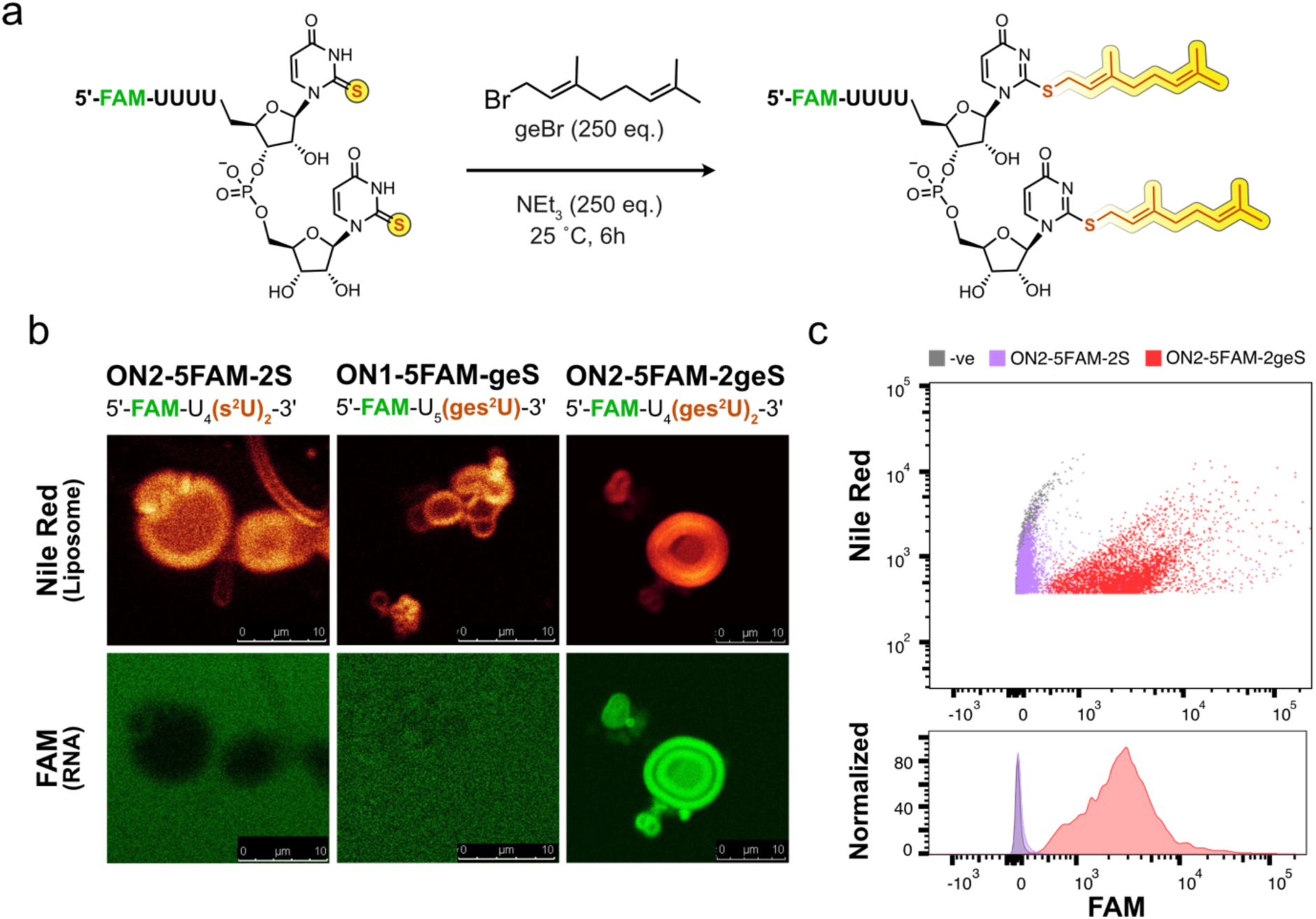
Lipidation of homo-U RNAs by ges^2^U modification. (a) General procedure for post-synthetic geranylation of s^2^U-containing oligos -purified s^2^U oligos are incubated with 250 eq. of geBr and NEt_3_ in 50% EtOH for 6 h at r.t. The crude is then quenched and purified (see Methods). (b) Confocal images display localisation of FAM-labelled, s^2^U or ges^2^U modified RNA oligos onto Nile Red stained Egg PC liposome surface, supporting the lipidating capacity of ges^2^U but not s^2^U. (c) Flow cytometry data on resuspension experiments of liposome binding ges^2^U-modified oligos supporting potential enrichment by sedimentation. The negative control contains only Nile Red stained liposomes.

In order to explore the membrane-anchoring properties of ges^2^U-containing RNA, we synthesized two 5’-FAM-labelled versions of these RNA strands (ON2-5FAM-2S and ON2-5FAM-2geS). The distribution of these strands was then analyzed by confocal microscopy in the presence of zwitterionic Egg PC liposomes, prepared by a reported lipid film rehydration method (see Methods).^25^ It is noteworthy that phosphatidylcholine-based zwitterionic lipids were recently suggested to be prebiotically plausible.^26^ The synthesized FAM-labelled ges^2^U oligos were added to the Nile Red stained liposomes and the distribution of the RNAs was analyzed by confocal imaging. As expected, the image of the non-geranylated reference strand ON2-5FAM-2S did not show any association with the liposome (Fig. 1b). The presence of the two geranyl chains in ON2-5FAM-2geS, however, leads to strict localization of the RNA to the liposome (Fig. 1b). We then performed the same experiment with the same RNA strand carrying just one geranyl unit and also in this case, an association of ON1-5FAM-geS with the liposome surface is clearly detected. This is surprising given that the oligonucleotides have in total 6 negatively charged phosphordiester groups (Fig. 1b). The result that one geranyl chain already allows membrane-anchoring of RNA confirms the initial idea that geranylation of RNA with the help of the non-canonical nucleoside s^2^U (ges^2^U) opens the possibility for small RNAs that certainly dominated on the early Earth to escape from the aqueous phase into a lipophilic environment.

In order to explore if the membrane association is affected by the position of the ges^2^U units within RNA, we next varied the position of the geranyl side chains on the RNA. Even when we separated the two ges^2^U nucleotides in 5’-FAM-U_2_(ges^2^U)U_2_(ges^2^U)-3’ (ON3-5FAM-2geS), only a slightly weakened lipid-binding effect was observed in the confocal image with liposome (Fig. S33), which confirms that ges^2^U allows stable anchoring of RNA to membrane surfaces.

A functional primitive translation system requires the stable presence of different RNA species in one single compartment. Today, the various RNAs are tightly associated to create a translating ribosome. In order to explore if modified RNAs can concentrate on liposomes, we investigated the possibility to enrich liposome-bound RNAs via sedimentation. To this end, we spun down the RNA-attached liposomes and then resuspended them in fresh buffer (see Methods). Even after three re-suspension cycles, the samples showed a strong FAM emission on the liposome with ON2-5FAM-2geS in the flow cytometry analysis (Fig. 1c & S31), contrary to ON2-5FAM-2S (Fig. 1c), ON1-5FAM-geS (Fig. S32), or ON3-5FAM-geS (Fig. S33), proving that the liposome association does allow the enrichment of ges^2^U containing oligos that may be present in solution.

In order to establish a peptide synthesis on the surface of a liposome, longer and more sequence diverse RNAs need to be attached to the liposome (Fig. 2a & S34-S36). To explore this possibility, we performed a confocal analysis of the 6, 8 and 11mer RNA strands 5’-FAM-GCGA(ges^2^U)_2_-3’ (ON4-5FAM-2geS), 5’-FAM-CAGCGA(ges^2^U)_2_-3’ (ON5-5FAM-2geS), and 5’-FAM-GUACAGCGA(ges^2^U)_2_-3’ (ON6-5FAM-2geS). In all of these cases, we observed that the two terminal geranyl-units allow the stable association of the corresponding RNA with the liposomes. However, when we further increased the length of the RNA to reach a nucleotide ratio of 12:2 (canonical:ges^2^U), as in 5’-FAM-AUCGUACAGCGA(ges^2^U)_2_-3’ (ON7-5FAM-2geS), the localization effect slowly vanished (Fig. 2a and Fig. S37). This suggests that a certain ratio of canonical:ges^2^U nucleotides must be met for the lipid-anchoring effect to happen.

**Figure 2.**
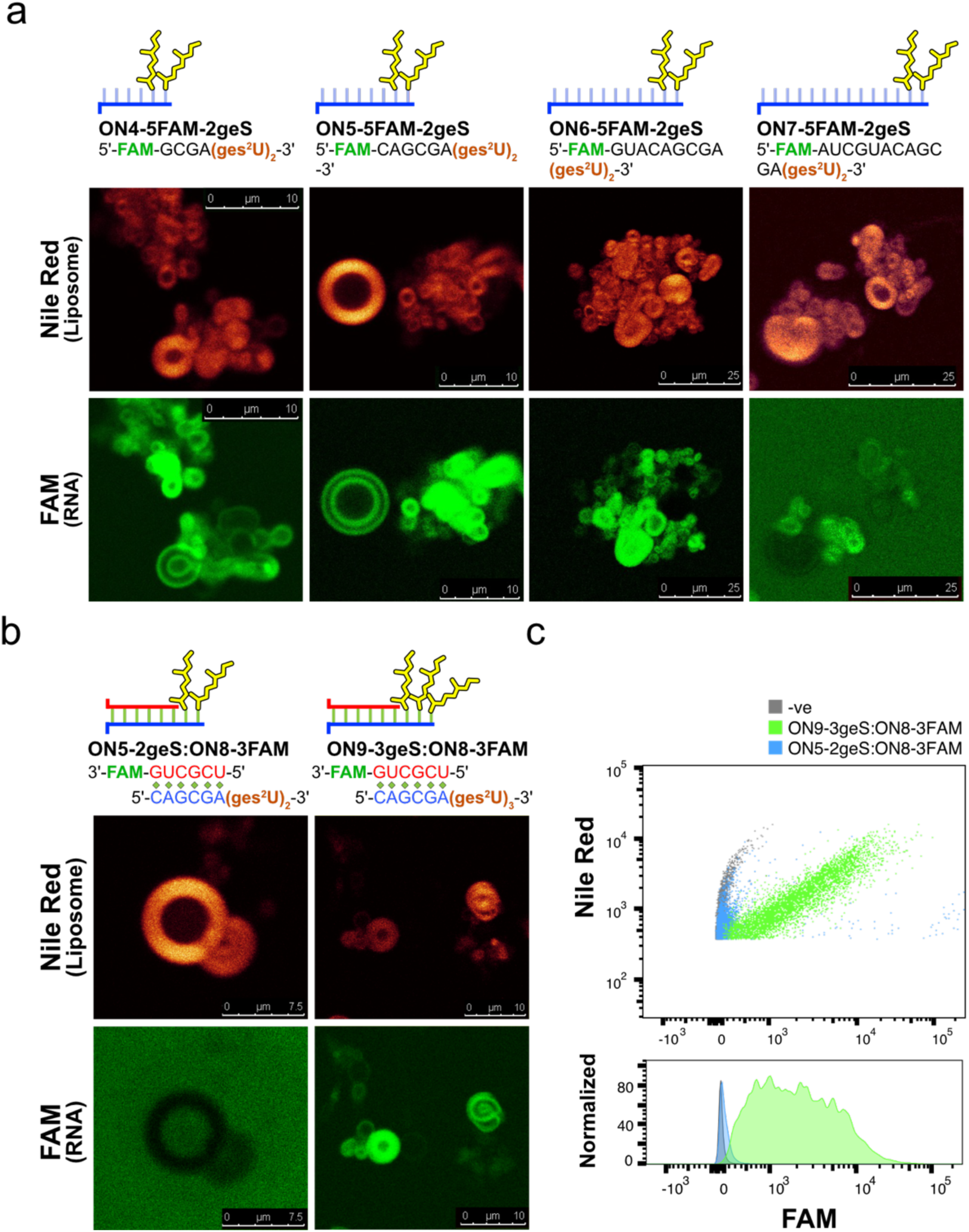
Length and duplex effects on liposome localisation. (a) Confocal images of oligos consist of a (ges^2^U)_2_ lipid-anchor with increasing number of canonical nucleotides suggest a length limitation in ges^2^U lipidating capacity. (b) Confocal images of two six-basepair RNA duplexes suggest a FAM-labelled canonical oligo can be recruited by its couterstrand containing three but not two ges^2^Us onto the liposome surface. (c) Flow cytometry data of the mentioned RNA duplexes in the resuspension experiment, showing possibility of co-enrichment. The negative control contains only Nile Red stained liposome and ON8-3FAM.

Next, we asked if a membrane-bound geranylated RNA can capture a complementary non-geranylated RNA strand from solution to form a stable membrane bound RNA duplex, which is the prerequisite for sequence-specific reactions, like template-directed peptide synthesis (Fig. 2b & c). We therefore investigated the distribution of a 3’FAM-labelled, canonical RNA strand 5’-UCGCUG-FAM-3’ (ON8-3FAM) in the presence of liposomes loaded with either a bis- or tris-geranylated (unlabelled) 5’-CAGCGA(ges^2^U)_2_-3’ (ON5-2geS) or 5’-CAGCGA(ges^2^U)_3_-3’ (ON9-3geS) RNA strand respectively, using confocal and flow cytometry analyses. In the experiment of the duplex ON5-2geS:ON8-3FAM on liposomes, we initially observed an exclusion of the FAM-labelled oligo from the Nile Red stained liposome, similar to what we observed for the negative control with just ON8-3FAM (Fig. S38). However, in the experiment performed with ON9-3geS:ON8-3FAM, a clear accumulation of the FAM emission on the liposome surface could be observed (Fig. 2b & c). To confirm that the accumulation is based on proper base pairing, the experiment was repeated by replacing ON8-3FAM with a mutated version of the labelled strand, 5’-ACGGU-FAM-3’ (ON10-3FAM). As expected, no FAM signal on the liposome surface was observed (Fig. S39), showing that correct base pairing is essential for stable lipid anchoring of the ungeranylated strand. This experiment shows that tris-geranylated RNAs can capture RNAs with the correct sequence to sequester them from the aqueous phase onto the lipid surface.

### Liposome-catalyzed specific RNA geranylations

We next investigated if liposome can facilitate the geranylation of RNAs to anchor initially non-geranylated RNA to its membrane. This could have been the basis to establish a selection system that enriches ges^2^U RNAs (Fig. 3). Using again geBr as a model compound of activated geraniols, we studied the geranylation of a single stranded (ss) RNA 5’-(s^2^U)_2_UCGCUG-3’ (ON11-2S) containing two s^2^U sites (Fig. 3a and 3b-I). This oligonucleotide was placed in a solution containing 20 mM geBr, 100 mM borate buffer (pH10) and 100 mM NaCl. Geranylation was compared in the absence and presence of liposomes (-Lip and +Lip, respectively) by HPLC and MALDI-TOF after 2 h (see Methods). In the experiment we indeed detected both single and double geranylation of s^2^U in ON11-2S. Importantly, the geranylation was significantly enhanced by the presence of the liposomes. For instance, the formation of the double-geranylated RNA (5’-(ges^2^U)_2_UCGCUG-3’; ON11-2geS) tripled from 3.1±0.4% to 10.5±1.8% by the addition of liposomes (Fig. 3a & S40). Since geBr is hardly miscible with water, we believe that the accumulation and concentration of the lipophilic geBr on the liposome membrane is responsible for the effect.

**Figure 3.**
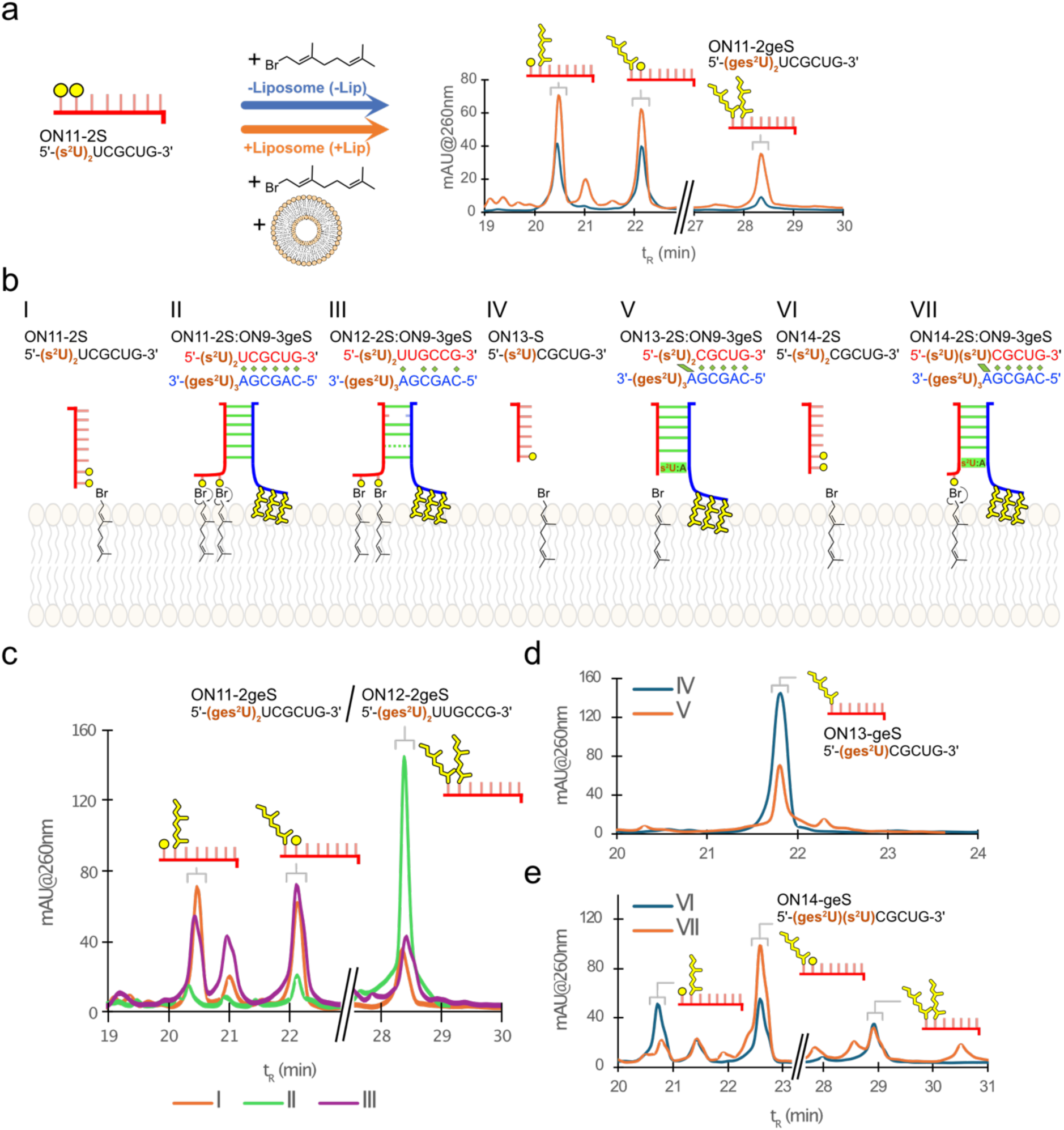
Selective geranylation of s^2^U on liposomal membrane. (a) Geranylation of ON112-S catalysed by liposomes. The HPLC chromatograms showed that both single and double geranylations are improved, with the yield of double geranylation increased from 3.1±0.4% to 10.5±1.8%. (b) Seven experiment setups in the presence of liposome to test the effect of sequence-specific geranylation (I-III) and site-specific geranylation (IV-VII) on the surface of catalytic liposomes (see Methods). (c) Overlaying HPLC chromatograms showing the catalytic effect of complementary, ges^2^U oligos on s^2^U geranylation of its counterstrand in II and the loss of such catalytic activity when the dsRNA is weakened by mutation in III. (d) & (e) HPLC chromatograms showing the effect of protective base pairing of s^2^U results in reduced geranylation (IV & V), and such protection can catalyse site-specific single geranylation and selectively form isomer with a terminal ges^2^U.

We next asked if the experiment shown in Fig. 3b-I can be further developed to achieve base pairing of an s^2^U-containing counterstrand with a membrane-bound complementary ges^2^U-modified oligonucleotide for subsequent geranylation (Fig. 3b-II). For this selection-based geranylation experiment, we added in addition to the experiment in Fig. 3b-I the ges^2^U-containing counterstrand, ON9-3geS. Analysis of the reaction products showed that the peaks of the single geranylated product of ON11-2S almost completely disappeared, while the yield of double geranylated oligonucleotide, 5’-(ges^2^U)_2_UCGCUG-3’ (ON11-2geS), showing an increase of almost 5-fold from 10.5±1.8% to 48.2±2.7% (Fig. 3b-II & c). To confirm that base pairing is responsible for this effect, we repeated the experiment with an s^2^U oligonucleotide mutated by a C/U swap (Fig. 3b-III). This gives the oligonucleotide 5’-(s^2^U)_2_UUGCCG-3’ (ON12-2S), which can only form a G:U wobble base pair and a C:A mismatch with ON9-3geS. As expected, now the yield of fully geranylated ON12-2geS dropped to only 14.2±0.6%, while those of the single geranylated products recovered (Fig. 3c & S41). Combined, these data show, that geranylated RNAs on the surface of liposomes are able to capture and select their non-geranylated counterpart from solution to achieve their geranylation on the liposome.

We then investigated how the position and the base pairing situation of the (to be geranylated) s^2^U in RNA affects the liposome-catalysed geranylation (Fig. 3b-IV-VII). For the experiments, we prepared the RNA strand 5’-(s^2^U)CGCUG-3’ (ON13-S) with a single s^2^U unit and reacted it with geBr in the presence of liposomes, first as the single strand (ssRNA) (Fig. 3b-IV) and later as a duplex (dsRNA) with ON9-3geS (Fig.3b-V). In the dsRNA situation, the s^2^U is base pairing with an A in the counterstrand (Fig. 3b-V). The obtained data are depicted in Fig. 3d. We observed that the liposome-catalysed geranylation now drops from 30.0±1.2% in the ssRNA to 16.0±1.0 in the dsRNA situation (Fig. 3d & S42). This can be explained by the known fact that base paired nucleobases have reduced reactivity.^27^

Interestingly, in situations (Fig. 3b-VI & VII) where we inserted a second s^2^U to form an overhang in 5’-(s^2^U)_2_CGCUG-3’ (ON14-2S), we observed an increase of the yield for one of the single geranylated isomers, from 19.6±1.6% to 36.4±2.9%, and a decrease of the other, while no significant change for the double geranylated species was detected, when the dsRNA with ON9-3geS (Fig. 3b-VII) was compared to the ssRNA situation (Fig. 3b-VI). The data are presented in Fig. 3e & S43). The experiment shows that the liposome-assisted geranylation is operating selectively on unpaired s^2^U of the ungeranylated RNA strand.

Finally, we studied the stability of the geranyl group on RNA, since it is reported that ges^2^U hydrolyses under basic conditions.^24^ We first incubated ON12-2geS at pH7 and pH10 respectively, in the presence or absence of 20 mM Egg PC liposomes and monitored the integrity of the RNA by HPLC over 4 d (see Methods). While we detected only minor difference at pH7, we observed a mitigation of ges^2^U hydrolysis at pH10 in the presence of liposomes. In the absence of liposomes (-Lip), we detected a survival of the geranylation of 33.7±2.3% of ON12-geS, while this increased to 47.7±3.9% after 4 d in the presence of liposomes (+Lip) (see Ext. Data Fig. 2 and Fig. S44-S45). This result shows that not only can liposomes function as geranylating catalysts, but they also protect the membrane-anchored RNA from de-geranylation and its potential removal from the membrane.

### Geranylation influences the peptide coupling reaction

In order to study how an RNA-liposomal association, mediated by s^2^U and its geranylation, would influence a primitive amino acid selectivity, we used the previously introduced RNA-peptide world concept, in which RNAs gain the ability to decorate themself with peptides in the presence of the non-canonical nucleotides *aa*^6^A and nm^5^U.^17^ For the experiment, we synthesized oligonucleotides that contain in total three non-canonical fossil-nucleosides ges^2^U, *aa*^6^A and nm^5^U. From a synthetic point, it was rewarding that the unprotected carboxylic acid of *aa*^6^A did not interfere with the post-synthetic geranylation step.^17,28^ The amino group of the nm^5^U was synthetically deprotected after the geranylation step (see Methods, Fig. S13-S14).

First, the melting temperature of a 5-basepair dsRNA formed between 5’-(g^6^A)AGCGA(s^2^U)_3_-3 (ON15-g^6^A-3S) and ON16-nm^5^U was determined to be 34°C (Fig. S46), confirming duplex formation at room temperature. We then determined the yields of the RNA-peptide hairpin formation based on HPLC calibration curves (Fig. S47), obtained from canonical oligonucleotides that substitute the modifications with their closest canonical analogs (CON-1-4, Ext. Data Table 1, also see Methods).^17^ To obtain an overview of these similar reactions performed in the presence (+Lip) and absence (-Lip) of liposomes, relative yields were calculated by dividing the peak area of the formed RNA-peptide products -Lip by that of it in +Lip (Fig. 4b, also see Methods and Ext. Data Table 2 for all coupling yields). To ensure that the data are not influenced by the background inhibition from the Egg PC lipids, peptide-coupling reactions between the non-geranylated oligonucleotides, 5’-UCGCU(nm^5^U)-3‘(ON16-nm^5^U) and 5’-(g^6^A)AGCGA-3‘ (ON17-g^6^A), were compared in the presence (+Lip) and absence (-Lip) of liposomes (Fig. 4a-I). The experiment showed very similar yields, with relative yield determined to be 112.6±18.0 % (Fig. S48), showing that liposomes do not inhibit the reaction. When we, however, performed the peptide coupling step in the membrane-bound situation with RNA strands containing ges^2^U units, we observed, first of all, a general reduction of the coupling yields (Fig. 4a-II-IV). In the coupling experiments of ON16-nm^5^U:ON15-g^6^A-3geS (Fig. 4a-II) and ON18-nm^5^U-geS:ON17-g^6^A (Fig. 4a-III), where the lipid-anchoring (ges^2^U)_3_ tail was located on either the *aa*^6^A strand or the nm^5^U strand, the relative yields of coupling decreased to 61.7±9.6 % and 65.4±3.4 %, respectively (Fig. 4c). When both strands of the dsRNA were geranylated with the (ges^2^U)_3_ tails facing each other (Fig. 4a-IV), the coupling yield further decreased in the case of dsRNA ON15-g^6^A-geS:ON18-nm^5^U-geS to finally only 50.3±1.7 % %. The inhibition is likely due to the repulsion of the lipid membrane against the hydrophilic DMTMM•Cl activator, which makes it hard to access the carboxy group of *aa*^6^A at the lipid-water interphase.

**Figure 4.**
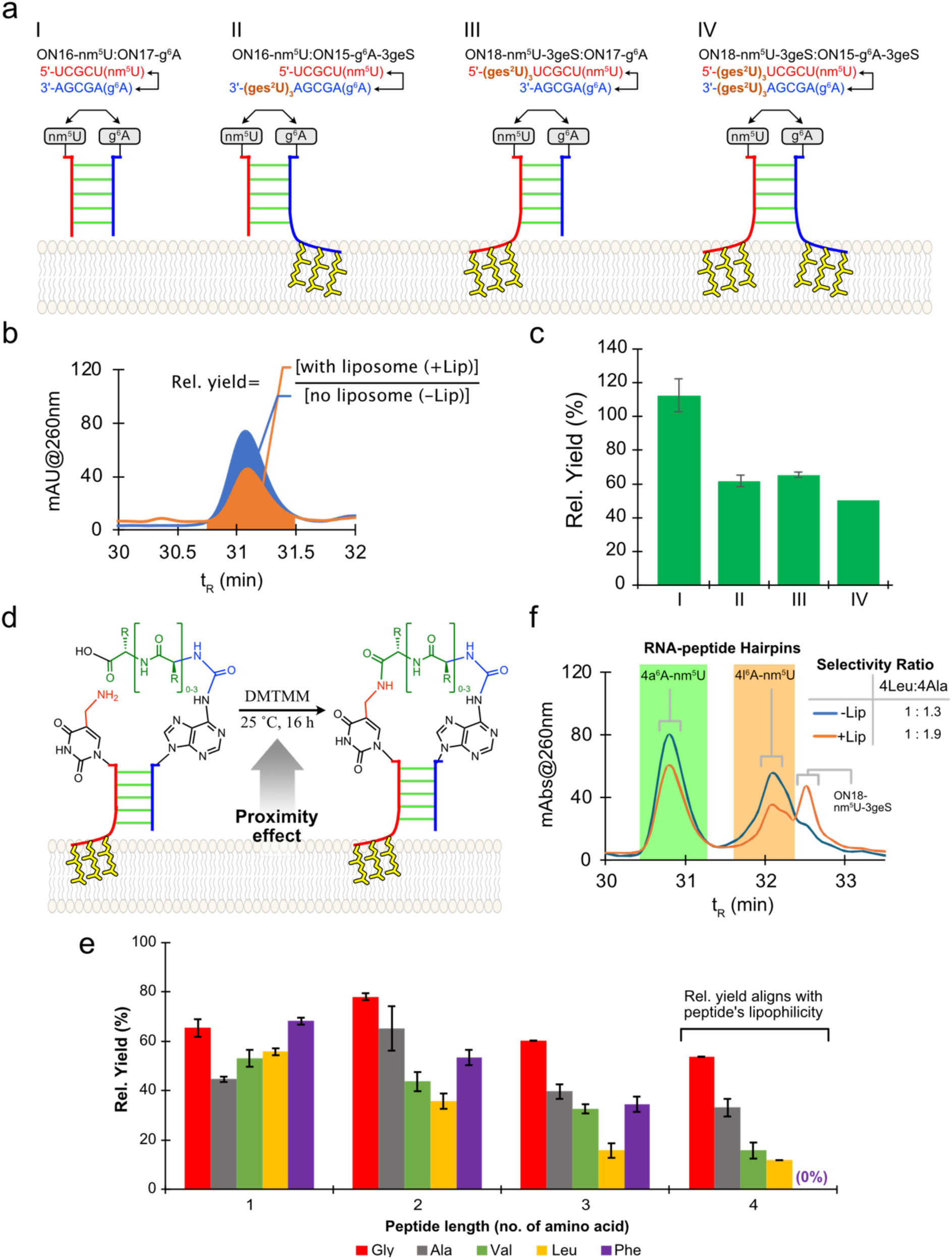
Template-directed peptide couplings on liposome surface. (a) Depiction of 5 experiment setups to investigate the lipid-anchoring effect on peptide coupling reactions. (b) Relative yields of each coupling reaction are calculated by dividing the peak area of formed RNA-peptide hairpin in +Lip by that in -Lip. (c) Relative yields calculated for coupling reactions described in (a). (d) Schematic of lipid surface template directed coupling by incubating 10 µM dsRNA with the corresponding modifications in 100 mM NaCl, 100 mM MES pH=6.0, and 50 µM DMTMM•Cl in the absence (-Lip) or presence (+Lip) of liposomes. (d) Result of coupling reaction relative yields with *aa*^6^A oligos charged with amino acids or their repeating oligopeptide and reacted with ON18-nm^5^U-geS. The coupling reactions of tetrapeptide-RNA features decreasing relative yields with increasing lipophilicity of the reacting peptides. (e) Competition experiment between ON17-4a^6^A and -4l^6^A with ON18-nm^5^U-geS. The ratios of formed RNA-peptide hairpins of 4l^6^A to 4a^6^A are 1:1.3 and 1:1.9 in -Lip and +Lip setups, respectively.

Inspired by these results, we designed a system in which an nm^5^U and (ges^2^U)_3_ modified universal amino acid acceptor strand, 5’-(ges^2^U)_3_UCGCU(nm^5^U)-3’ (ON18-nm^5^U-3geS), was reacted with non-geranylated donor strands with different amino acids or peptides coupled to the *aa*^6^A nucleotide, 5’-(*aa*^6^A)AGCGA-3’ (ON15-*aa*^6^A, with *aa*^6^A = g^6^A, a^6^A, v^6^A, l^6^A or f^6^A). This ON15-*aa*^6^A contained different amino acids or short *homo*-peptide of these amino acids (ON15-<u>n</u>*aa*^6^A, with <u>n</u> = 2-4), which we all prepared by our reported method.^17^ We then performed coupling reactions between ON18-nm^5^U-3geS and all these 16 ON15-*aa*^6^A oligonucleotides in separate experiments, as depicted in Fig. 4d. The data obtained (Fig. 4e) with the RNAs charged with a single amino acid (ON17-*aa*^6^A) show already surprisingly different relative yields. A clear order of reactivity Phe ≈ Gly > Leu ≈ Val > Ala was detected, ranging from Phe 68±1% to Ala 45±1%. As the peptide length increases, the relative yields for the different oligopeptides started to deviate. Beginning at the dipeptidal RNAs (ON17-2*aa*^6^A), we started to observe relative yields in line with the lipophilicity of peptides, with 2Gly ≈ 2Phe > 2Ala > 2Val ≈ 2Leu. Most interesting are the coupling reactions of tetrapeptidal RNAs (ON17-4*aa*^6^A), where we observed a perfect alignment of relative yields and peptide lipophilicity, giving the order of 4Gly > 4Ala > 4Val ≈ 4Leu. For 4Phe, the formation of RNA-peptide hairpin was no longer detectable in the presence liposome (+Lip). We obtained data ranging from Gly 54±3% to Phe 0% (undetectable). We also noted the emergence of solubility problems when we tried to prepare ON17-5l^6^A and ON17-5f^6^A (data not shown) and therefore, we stopped the study at the RNA-tetrapeptide level.

To demonstrate selectivity in a direct competition experiment, we set up coupling experiments with a mixture of 1:1:1 ON17-4a^6^A:ON17-4l^6^A:ON18-nm^5^U-geS (Fig. 4f & S71). Here, we observed that the ratio of the formed RNA-peptide hairpins from 4l^6^A to 4a^6^A was initially 1:1.3 in -Lip setup, which was then increased to 1:1.9 in the +Lip case, showing that the liposomes can amplify subtle differences. The result of this study shows that lipid-anchored RNAs can be involved in peptide formation reactions, and certain amino acids react more preferably when the RNA is bound to a liposome surface. This suggests a potential scenario where short peptides and RNAs could be co-selected by a lipid phase. One can imagine that liposomes formed by different lipid compositions could attract different peptides and RNAs with different sequences or modifications, which slowly evolved to form the basis of the modern genetic code.

### Conclusion

Life is characterized by compartimentalization. All living processes are shielded from the outside world inside a membrane surrounded space (cell) and within this space, defined processes happen in defined places, often again associated with membranes such as the Golgi apparatus. It is currently assumed that in the phase of chemical evolution, RNA must have gained the ability to first associated with membranes early on, and to be trapped inside membrane surrounded vesicles to create protocells later.^4,5^ However, given the highly negatively charged nature of RNA, it is unclear how a membrane association could have occurred. Previous studies revealed that RNA can get lipidated in the presence of cations,^6,7^ fatty acid modifications^8^ or in association with specific peptides,^9,10^ but this generates uncontrolled and unspecific lipidations. Ideas were also formulated of how RNA could have become lipidated at their ends, but this at best allows to charge RNA with two lipid units, which is insufficient for stable membrane anchoring.^8^

In order to solve this problem we analyzed the chemical structure of non-canonical nucleosides, which are thought to be living fossils of an early RNA World and discovered geranylated nucleosides such as ges^2^U in the anticodon loop of certain RNAs. With the knowledge that the membranes of archaea are derived from geranyl units, we explored the possibility to anchor RNA to membranes with ges^2^U units, and discovered that two or three of these building blocks within single and double stranded RNA allows the efficient sequestering of these RNAs from an aqueous solution to the lipid membranes. Importantly, RNA containing the ges^2^U precursor nucleoside 2-thiouridine (s^2^U) can be selectively recognized by membrane anchored RNA counterstrand, which, in the presence of activated geranyl molecules, allows specific membrane associated geranylation. This establishes a selection system, in which only those RNA strands are geranylated and hence, membrane-anchored that are fully complementary.

We finally show that these membrane-anchored RNAs can perform primitive peptide synthesis directly at the surface of the membrane and this all together then provides a scenario in which amino acid-carrying donor RNAs are attracted from the aqueous environment to the lipid surface by membrane-bound acceptor RNAs to feed a peptide synthesis on the lipid surface. Although we show here only the first step of this membrane-bound peptide growth cycle, our previous reports show that the reported chemistry can be extended to allow the growth also of longer peptides.^17,29^

The surprising result of the peptide growth study is that certain amino acids are more preferably reacting when the RNA is bound to a liposome surface. The differences became particularly apparent, when the peptides grew a bit longer. This suggests that selectivity can be achieved between certain RNA with specific sequences or modifications and specific amino acids depending on how well the specific membranes allow the growths of specific peptides. This in turn would offer the possibility to decipher the origin of the genetic code. Although we are far from proposing why a certain amino acid is encoded by a certain base triplet, our system suggests a route along which amino acids and RNA sequences could co-evolve along a mutually beneficial selection path. One can imagine that liposomes formed by different lipid compositions could attract different peptides, and RNAs with different sequences or modification patterns, to slowly evolve the modern genetic code.

## Methods

### Post-synthetic modification of s^2^U to ges^2^U

Purified and desalted s^2^U containing ON was dissolved in 66% EtOH to achieve a final concentration of 2 mM. Geranyl bromide (250 eq.) was added to the solution and the mixture was vortexed thoroughly to fully dissolve the geranyl bromide. After that, Et_3_N (250 eq.) was added and the solution was incubated at r.t. with 1400 rpm in a thermomixer for 6 h. The reaction was then quenched by adding 5X volume of 20% EtOH in HPLC buffer A and dried with a SpeedVac concentrator. The crude was then resuspended in water, filtered, purified by HPLC and desalted with standard protocol (see Scheme 1 & SI).

**Scheme 1.**
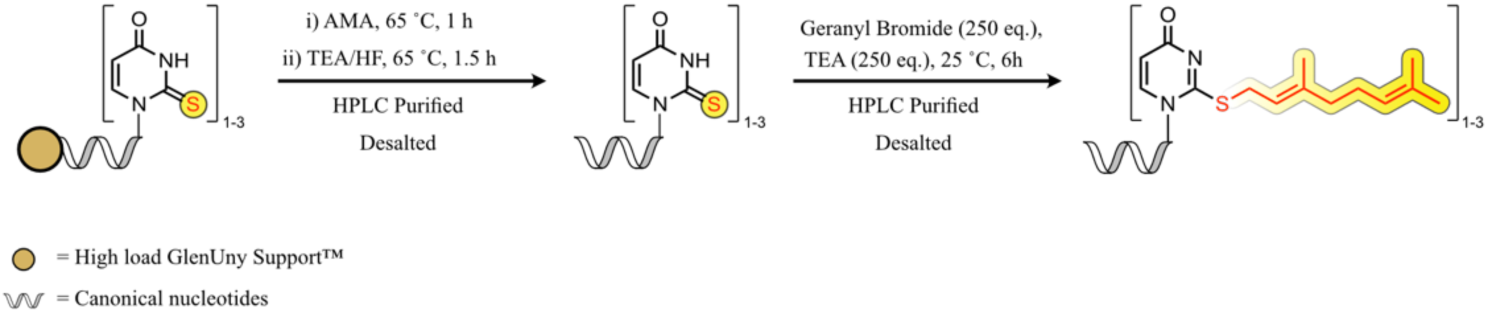
RNAs containing s^2^U were first synthesized and purified according to standard protocol (see SI). Afterwards, it was reacted with geBr and Et_3_N to achieve multiple incorporation of ges^2^U.^24^

### Preparation of liposome

Required volume of Egg PC chloroform solution was dried in a 10 mL round bottom flask to form a lipid film, which was then rehydrated with a 2-fold reaction buffer. The liposome suspension was then pressed through a mini-extruder (Avanti Polar Lipids) equipped with a ø=100 nm Nucleopore™ Track-Etched membrane (Whatman) for 15 times and a small sample of the suspension was analysed by dynamic light scattering. For confocal microscopy, the extrusion and DLS steps were skipped. For confocal microscopy and FACS analysis, 0.1% and 1.0% mol of Nile Red was added to stain the liposome respectively.

### Selective geranylation of s^2^U in the presence of liposome

Vesicle solution was prepared by thin film rehydration with an aqueous solution containing NaCl (100 mM) and borate buffer (pH = 10.0, 100 mM) giving a final Egg PC concentration of 40 mM. The ONs for the reaction (2 nmol each) were put into an 1.5 mL eppendorf tube and the ONs were dried in vacuo. Vesicle solution (100 µl) was added to the dried ONs and geBr (2.0 µmol, 0.397 µL) was added to the mixtures followed by vortexing. The reaction mixtures were incubated at 1000 rpm in r.t. over a period of 2 h. They were then injected to HPLC, and peaks were collected and analysed by MALDI-TOF mass spectrometry.

### Stabilisation of ges^2^U in the presence or absence of liposome

Liposome solution was prepared as mentioned above containing NaCl (200 mM) and glycine buffer (pH = 10, 100 mM) giving a final Egg PC concentration of 20mM. The ONs for the reaction (5.5 nmol each) were put into a 1.5mL eppendorf tube and the ONs were dried in vacuo. Vesicle solution (275 µl) was added to the dried ONs followed by addition of rehydration buffer (275 µl), to achieve a final concentration of 100 mM NaCl, 50 mM buffer and 10 mM Egg PC. For the -Lip setup, Egg PC was excluded. The samples were vortexed and shaken with a Thermomixer (1000 rpm) at r.t. for 4 days. After 0, 24, 48, 72 and 96 h an aliquot of 100 µl were removed from the samples and analysed by HPLC. The peaks were colleced and identified by MALDI-TOF mass spectrometry.

### Synthesis of ONs containing ges^2^U and *aa*^6^A

Solid support of s^2^U and *aa*^6^A containing ON was suspended in 10% DBU in THF in r.t. for 3 h. It was then washed with DMC 3 times and dried, followed by the standard procedure of deprotection and purification of ONs (see SI). The purified and desalted ON was then geranylated according to the method above (Scheme 2).

**Scheme 2.**
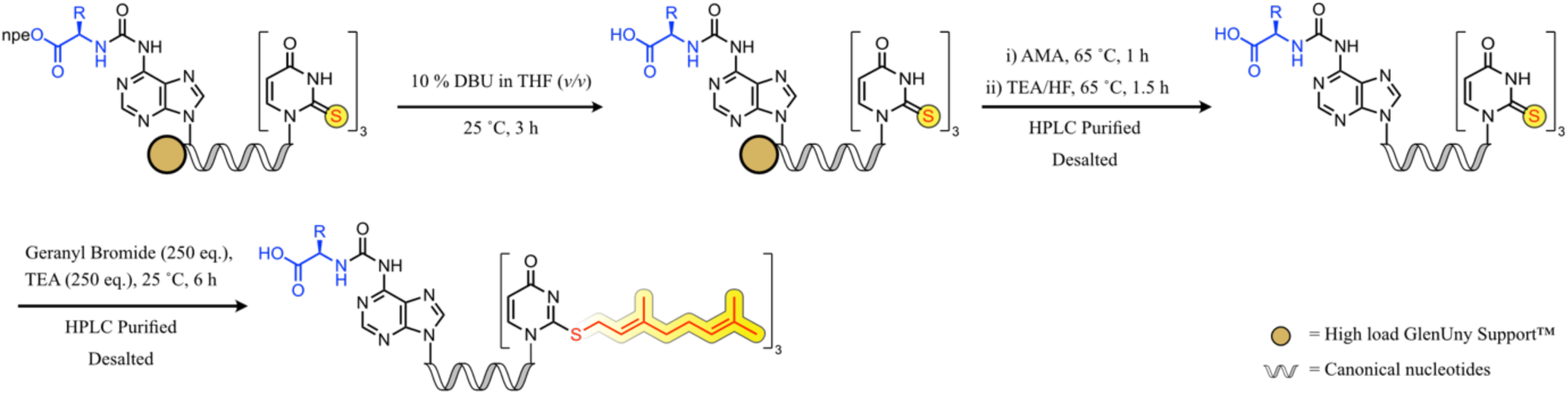
The solid support of *aa*^6^A and s^2^U containing ON was incubated in 10% DBU in THF for 3 h. The oligo was then deprotected and purified according to standard protocol (see SI). It was then geranylated according to the method above.

### Synthesis of ONs containing ges^2^U and nm^5^U

The HPLC purified and desalted ON containing teoc-nm^5^U and s^2^U was post-synthetically geranylated as described above. The purified and desalted ON was cooled on ice and cold 1M TBAF in THF solution was added to obtain a 0.5 - 1mM solution of the oligo. The solution was then incubated at 10 °C for 1.5 h. After that, it was cooled in an ice bath for 5 min and diluted with 5 mL cold HPLC buffer A. The solution was then dried and purified by HPLC (Scheme 3).

**Scheme 3.**
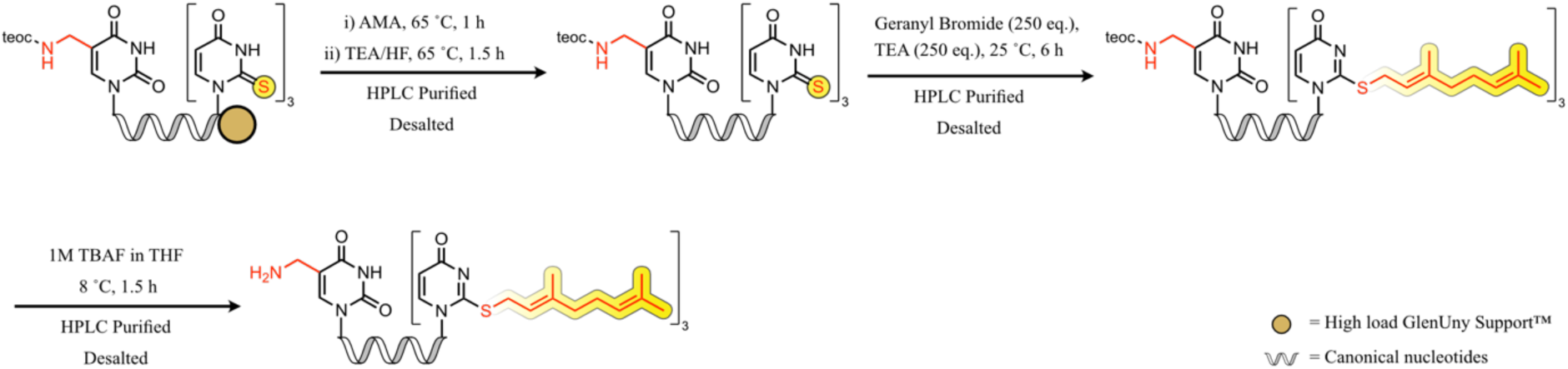
After the incorporation of s^2^U and teocnm^5^U on the solid phase, the ON was deprotected and purified according to standard protocol (see SI), followed by geranylation as described in Scheme 1. The teoc group was afterwards removed by incubating the oligo in 1M TBAF in THF at 10 °C for 1.5 h, then HPLC purified and desalted.

### Peptide coupling of *aa*^6^A and nm^5^U containing ONs

A solution of 10 µM *aa*^6^A and nm^5^U containing ONs, 100 mM NaCl, 100 mM MES pH=6.0, and 50 µM DMTMM•Cl was incubated at r.t. for 1 d in the presence (+Lip) or absence (-Lip) of 10 mM Egg PC.

The crudes of the coupling reactions were analyzed by RP-HPLC using an EC 250/4 Nucleodur 100-3 C18ec column from Macherey-Nagel. Buffers: A) 0.1 M AcOH/Et_3_N in H_2_O at pH 7 and B) 0.1 M AcOH/Et_3_N in 80% (v/v) MeCN in H_2_O. Gradient: unless otherwise stated, 0-80% of B in 45 min. Flow rate = 1 mL·min^-1^. Injection: 100 μL (1 nmol).

The yields of the reactions were calculated by integration of the chromatographic peaks of the products and the use of the calibration curves of the corresponding canonical ONs (CON1-4, Ext. Data Table 1). To simplify the calculations, we assumed that the formed products and the canonical oligonucleotides used for calibration featured identical extinction coefficients, which were calculated for single stranded RNAs, as previously described. It was expected that double strands and/or secondary structures were disrupted under the HPLC conditions used.

Relative yields of coupling reactions on liposome surface were calculated as follow:

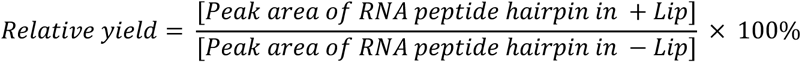

The isolated products were desalted with a Sep-Pak tC18 1 cc Vac Cartridge (Waters) and analysed by MALDI-TOF mass spectrometry.

## Data availability

The data that support the findings of this study are available within the paper and its Supplementary Information.

## Acknowledgements

We acknowledge the Core Facility Flow Cytometry at the Biomedical Center, Ludwig-Maximilians-Universität München, for providing the equipment and assistance with data analysis. This project has received funding from the European Research Council (ERC) under the European Union’s Horizon 2020 research and innovation programme under grant agreement no. 741912 (EPiR) and under the Marie Skłodowska-Curie grant agreement no. 861381 (Nature-ETN).

## Author Contributions

C.-Y.C. and J.S. synthesised the nucleoside phosphoramidites and performed all experiments described in this study. T.C. conceived the project and directed the research. All authors contributed to the analysis of the results and writing of the manuscript.

## Extended Data

**Ext. Data Table 1.**
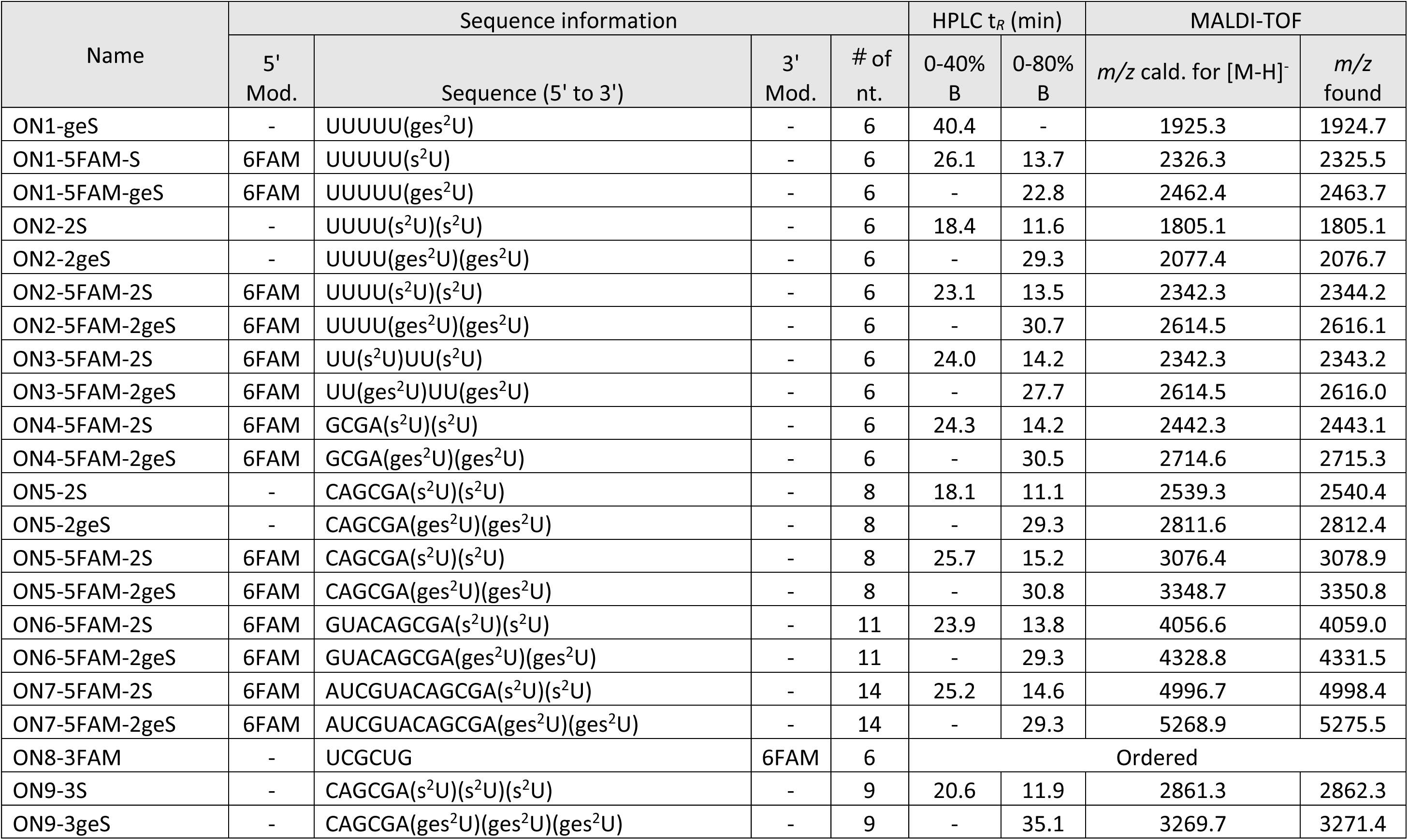
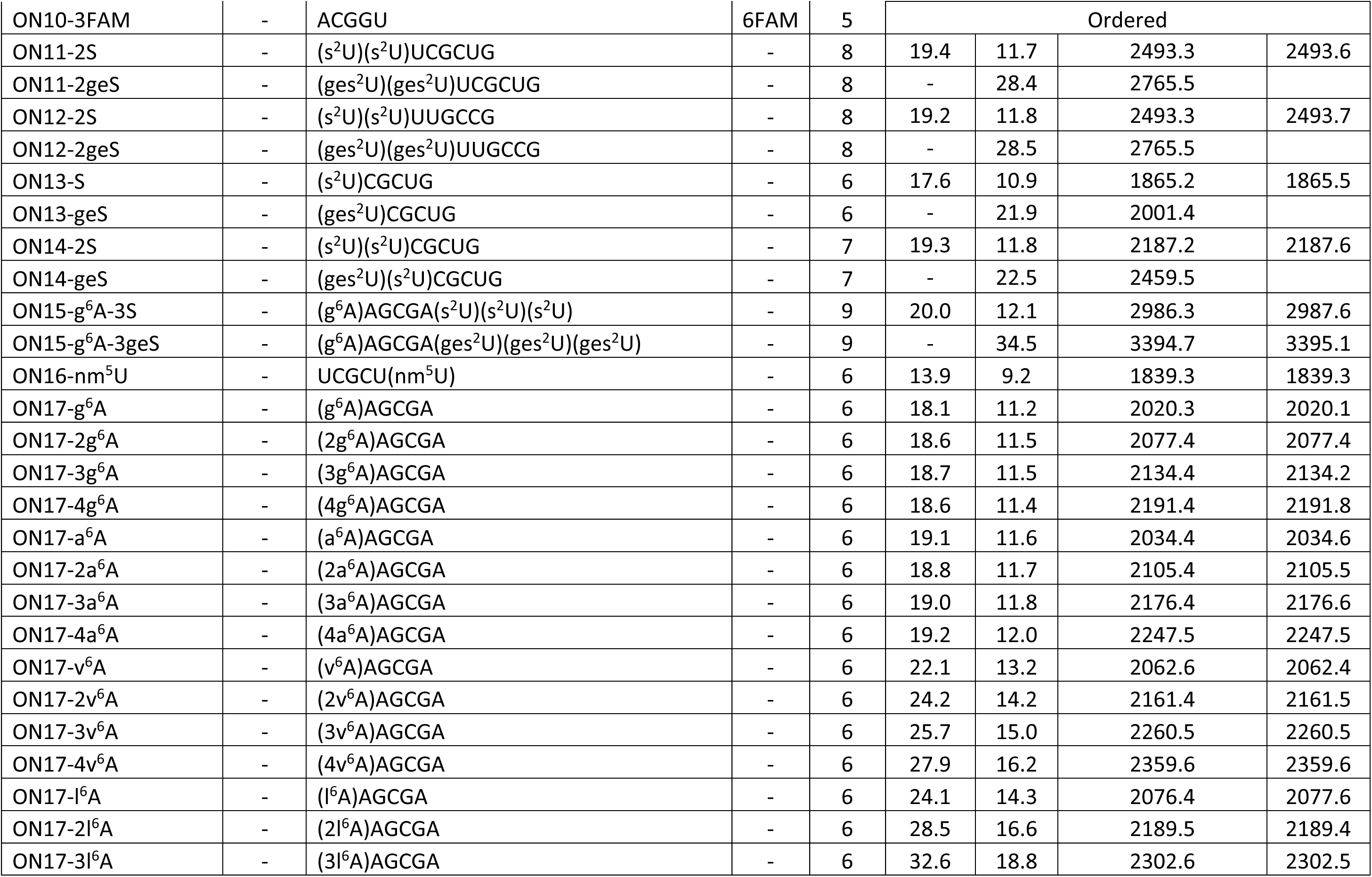
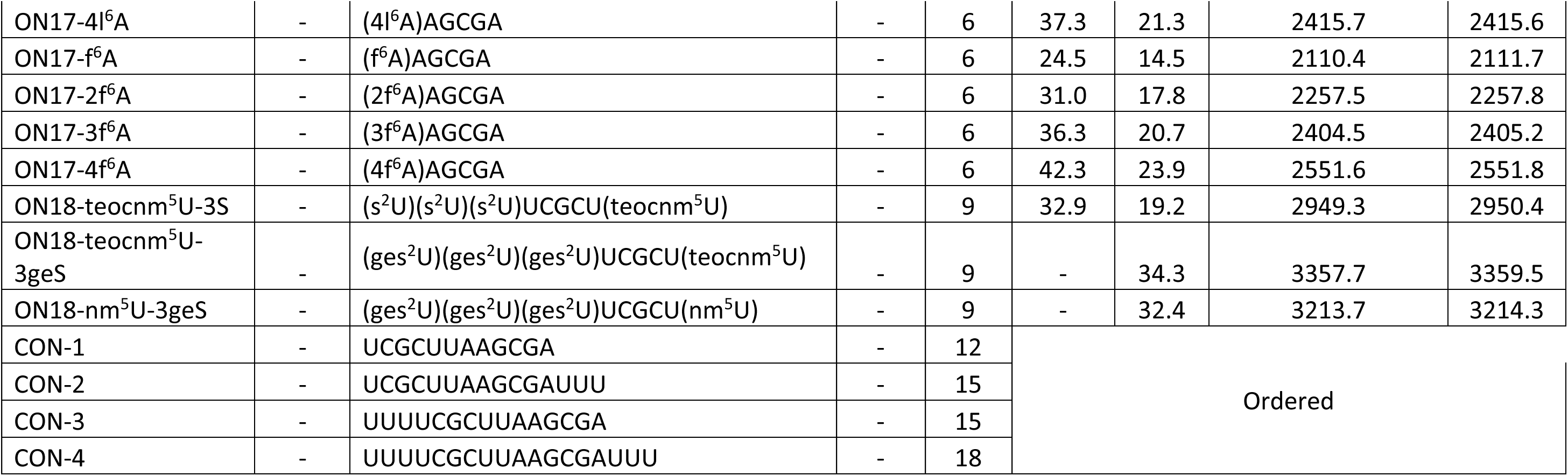
Information on ONs involved in this study. Modifications, sequences, HPLC retention time (t*_R_*) and gradient (0 → 40% or 0 → 80% buffer B), MALDI-TOF mass spectra of all ONs mentioned in this study.

**Ext. Data. Table 2.** Information on RNA-Peptide coupling reactions. Left to right: Parent ONs, HPLC retention times, MALDI-TOF and yields of performed *aa*^6^A ONs and nm^5^U ONs coupling reactions. *: Two product peaks of the same mass were observed in this reaction.

| Synthesised from | | HPLC $t_R$ (min) | MALDI-TOF | | Peptide formation yield (%) | | |
| --- | --- | --- | --- | --- | --- | --- | --- |
| *aa*<sup>6</sup>A Strand | nm<sup>5</sup>U Strand | 0-80% B | $m/z$ calcd. for [M-H]<sup>-</sup> | $m/z$ found | -Lip | +Lip | Rel. Yield |
| ON17-g<sup>6</sup>A | ON16-nm<sup>5</sup>U | 12.4 | 3841.6 | 3842.7 | 28.1 ± 2.0 | 30.9 ± 2.6 | 112.6 ± 18.0 |
| ON15-g<sup>6</sup>A-3geS | ON16-nm<sup>5</sup>U | 33.8 | 5217.0 | 5224.4 | 30.5 ± 3.1 | 19.5 ± 2.6 | 61.7 ± 9.6 |
| ON17-g<sup>6</sup>A | ON18-nm<sup>5</sup>U-3geS | 38.9 | 5217.0 | 5222.4 | 24.2 ± 0.3 | 16.9 ± 0.7 | 65.4 ± 3.4 |
| ON15-g<sup>6</sup>A-3geS | ON18-nm<sup>5</sup>U-3geS | 31.1 | 6591.4 | 6601.3 | 23.8 ± 1.9 | 13.4 ± 1.0 | 50.3 ± 1.7 |
| ON17-2g<sup>6</sup>A | ON18-nm<sup>5</sup>U-3geS | 31.3 | 5274.0 | 5273.2 | 31.7 ± 1.5 | 25.4 ± 1.1 | 78.0 ± 4.5 |
| ON17-3g<sup>6</sup>A | ON18-nm<sup>5</sup>U-3geS | 31.3 | 5331.0 | 5333.6 | 33.6 ± 1.4 | 21.5 ± 0.6 | 60.4 ± 2.8 |
| ON17-4g<sup>6</sup>A | ON18-nm<sup>5</sup>U-3geS | 31.3 | 5388.1 | 5379.6 | 28.3 ± 0.7 | 16.7 ± 0.5 | 53.8 ± 2.6 |
| ON17-a<sup>6</sup>A | ON18-nm<sup>5</sup>U-3geS | 31.4 | 5231.0 | 5229.3 | 25.8 ± 0.8 | 13.3 ± 0.2 | 44.7 ± 1.1 |
| ON17-2a<sup>6</sup>A | ON18-nm<sup>5</sup>U-3geS | 31.3 | 5302.0 | 5307.2 | 36.2 ± 1.6 | 24.5 ± 1.9 | 65.2 ± 9.1 |
| ON17-3a<sup>6</sup>A | ON18-nm<sup>5</sup>U-3geS | 31.3 | 5373.1 | 5378.2 | 37.9 ± 3.1 | 16.9 ± 1.0 | 39.7 ± 2.9 |
| ON17-4a<sup>6</sup>A | ON18-nm<sup>5</sup>U-3geS | 31.3 | 5444.1 | 5449.6 | 34.2 ± 3.6 | 13.3 ± 1.1 | 33.2 ± 3.6 |
| ON17-v<sup>6</sup>A | ON18-nm<sup>5</sup>U-3geS | 30.8 | 5259.0 | 5268.4 | 24.1 ± 0.5 | 14.2 ± 0.5 | 53.1 ± 3.5 |
| ON17-2v<sup>6</sup>A | ON18-nm<sup>5</sup>U-3geS | 30.9 | 5358.1 | 5364.1 | 42.2 ± 1.6 | 20.3 ± 2.1 | 43.8 ± 3.9 |
| ON17-3v<sup>6</sup>A | ON18-nm<sup>5</sup>U-3geS | 31.2 | 5457.2 | 5464.7 | 44.1 ± 1.1 | 15.7 ± 0.4 | 32.6 ± 1.9 |
| ON17-4v<sup>6</sup>A | ON18-nm<sup>5</sup>U-3geS | 31.2 | 5556.2 | 5565.8 | 19.7 ± 2.3 | 5.6 ± 0.1 | 15.8 ± 3.3 |
| ON17-l<sup>6</sup>A | ON18-nm<sup>5</sup>U-3geS | 30.8 | 5273.1 | 5278.7 | 22.4 ± 0.3 | 13.9 ± 0.2 | 55.9 ± 0.5 |
| ON17-2l<sup>6</sup>A | ON18-nm<sup>5</sup>U-3geS | 31.3 | 5386.1 | 5393.9 | 40.2 ± 1.3 | 16.3 ± 1.6 | 35.8 ± 5.8 |
| ON17-3l<sup>6</sup>A | ON18-nm<sup>5</sup>U-3geS | 31.7 | 5499.2 | 5508.6 | 26.1 ± 0.5 | 6.8 ± 0.5 | 15.8 ± 2.0 |
| ON17-4l<sup>6</sup>A | ON18-nm<sup>5</sup>U-3geS | 32.2 | 5612.3 | 5616.4 | 21.3 ± 0.3 | 5.3 ± 0.4 | 11.9 ± 1.9 |
| ON17-f<sup>6</sup>A | ON18-nm<sup>5</sup>U-3geS | 30.9 | 5307.0 | 5313.1 | 29.0 ± 0.4 | 20.8 ± 0.4 | 68.3 ± 1.4 |
| ON17-2f<sup>6</sup>A | ON18-nm<sup>5</sup>U-3geS | 31.2 | 5454.1 | 5464.6 | 30.5 ± 1.5 | 17.7 ± 0.2 | 53.5 ± 3.1 |
| ON17-3f<sup>6</sup>A | ON18-nm<sup>5</sup>U-3geS | 31.7 & 31.9* | 5601.2 | 5588.5 | 30.6 ± 1.5 | 12.6 ± 0.6 | 34.5 ± 3.0 |
| ON17-4f<sup>6</sup>A | ON18-nm<sup>5</sup>U-3geS | 32.2 | 5748.2 | 5739.7 | 28.9 ± 2.5 | n.d. | n.d. |

**Ext. Data Fig. 1.**
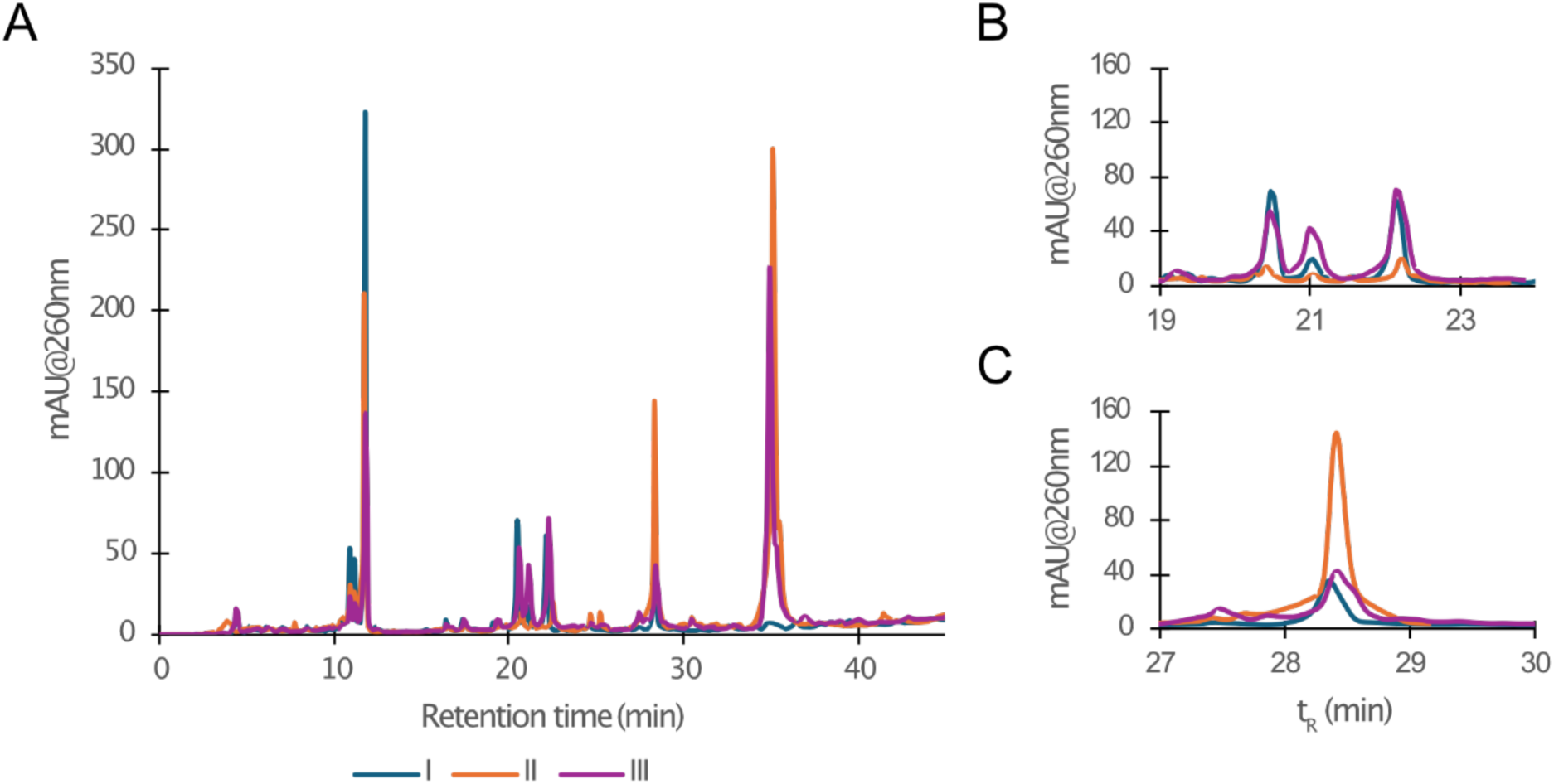
Liposome-catalysed geranylation of s^2^U ONs. (A) Overlaying HPLC chromatograms of situations (I) (blue), (II) (orange) and (III) (purple) in Fig. 3, with the starting material, ON11-2S, at 11.7 min; single geranylated ON11 species at 20-23 min, fully geranylated ON11-2geS at 28.4 min and ON9-geS for (II) and (III) at 35.1 min. The chromatograms were aligned with the ON11-geS peak. (B) Zoomed in region of 19-23.5 min, showing the decrease of single geranylated species in (II). (C) Zoomed in region of 17-30 min, showing the significant increase in formation of ON11-2geS in (II).

**Ext. Data Fig. 2.**
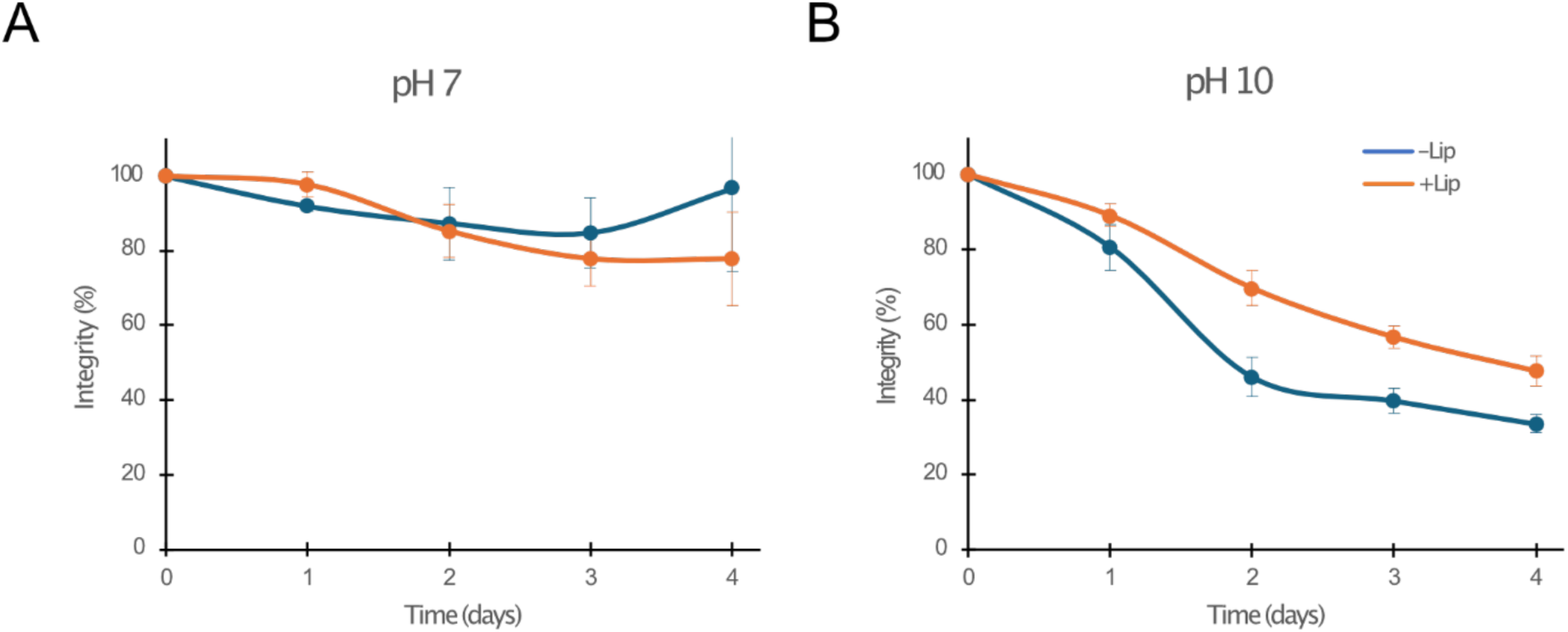
Stability study of ges^2^U against hydrolysis. Degradation of ON12-2geS over time in (A) phosphate buffer (pH=7.0) and (B) glycine buffer (pH=10.0) over 4 d in -Lip (blue) and +Lip (orange). Representative chromatograms of each time point are available in Fig. S44-S45.

