## Supplementary Information for "Membrane bound geranylated RNAs establish a primitive peptide synthesis system" (from another version of this preprint: version 1, 2024-08-02)

**Table of contents**

1. General information for materials and instrumentations____S2
2. Synthesis and characterisation data____S2
    2.1. Synthesis of protected amino acids and peptides____S2
    2.2. Synthesis of 2-thiouridine phosphoramidite____S13
    2.3. Synthesis of 5-aminomethyluridine phosphoramidite____S17
    2.4. Synthesis of $N^6$-carbamoyladenosine phosphoramidites____S21
3. General information and instrumentation for oligonucleotides____S32
    3.1. Synthesis and purification of oligonucleotides____S32
    3.2. Modification of oligonucleotides through post-synthetic reactions____S33
4. Characterisation data of synthesised oligonucleotides____S35
    4.1. Comparison of ges$^2$U incorporation with 2 different methods____S35
    4.2. HPLCs and MALDI-TOFs of post-synthetic geranylations____S37
    4.3. HPLCs and MALDI-TOFs of coupling of amino acids and peptides____S41
5. General information of liposome preparation____S46
    5.1. Dynamic Light Scattering (DLS) measurements____S47
6. General information for confocal microscopy and flow cytometry____S48
    6.1. General information for confocal microscopy____S48
    6.2. General information for flow cytometry____S48
    6.3. Controls for flow cytometry data____S49
    6.4. Confocal images and cytometry data____S50
7. General information for liposome-catalysed geranylation____S54
    7.1. Liposome-catalysed geranylation of s$^2$U ONs____S54
    7.2. HPLCs for liposome catalysed geranylation____S54
8. General information for ges$^2$U hydrolysis____S56
    8.1. HPLCs of ges$^2$U hydrolysis____S56
9. General information for UV-Vis melting____S57
    9.1. UV-Vis melting for ON17-g$^6$A-S and ON16-nm$^5$U____S57
10. General information of coupling reactions between *aa*$^6$A and nm$^5$U ONs____S57
    10.1. HPLC calibrations using canonical oligonucleotides____S59
    10.2. HPLCs and MALDI-TOFs of RNA peptide coupling reactions____S60

Reference____S72

NMR spectra____S73

#### 1. General information for materials and instrumentation

Reagents were purchased from commercial suppliers and used without further purification unless otherwise stated. All anhydrous solvents stored under inert atmosphere were also purchased. All reactions involving air/moisture sensitive reagents/intermediates were performed under inert atmosphere using oven-dried glassware. Routine $^{1}$H NMR, $^{13}$C NMR and $^{31}$P NMR were recorded on a Bruker AVANCE NEO™ 500 MHz spectrometer. Deuterated solvents used are indicated in the characterization and chemical shifts ($\delta$) are reported in ppm. All NMR *J* values are given in Hz. NMR spectra were analysed using MestReNova software version 14.3.0. High Resolution Mass Spectra (HRMS) were measured on a Thermo Finnigan™ LTQ-FT MS™ with ESI as ionization mode. IR spectra were recorded on a Perkin-Elmer Spectrum BX II FT-IR instrument equipped with an ATR accessory. Column chromatography was performed with silica gel technical grade (Macherey-Nagel), 40-63 µm particle size. Reaction progress was monitored by Thin Layer Chromatography (TLC) analysis on silica gel 60 F254 and stained with *para*-anisaldehyde solution.

#### 2. Synthesis and characterisation data

##### 2.1 Synthesis of protected amino acids and peptides

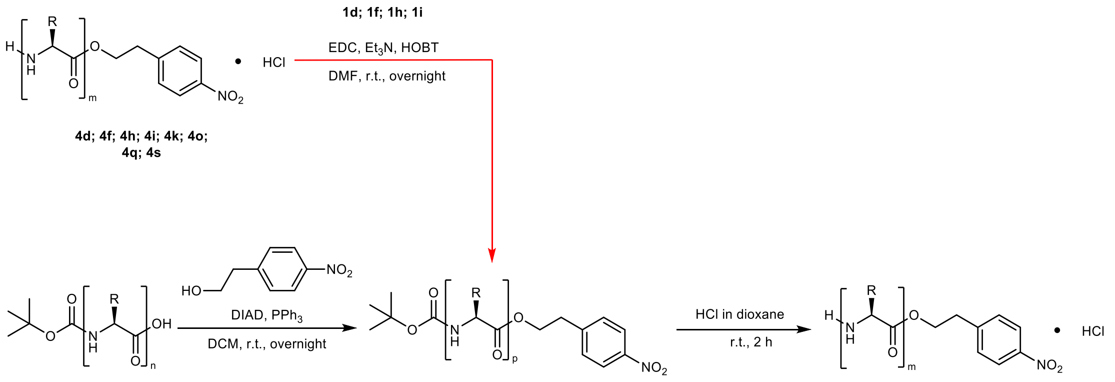


**1a:** R = H, n = 1
**1b[a]:** R = H, n = 2
**1c[a]:** R = H, n = 3
**1d:** R = $CH_2CH(CH_3)_2$, n = 1
**1f:** R = $CH_2C_6H_6$, n = 1
**1h:** R = $CH_3$, n = 1
**1i:** R = $CH(CH_3)_2$, n = 1

**2a:** R = H, p = 1
**2b:** R = H, p = 2
**2c:** R = H, p = 3
**2d:** R = $CH_2CH(CH_3)_2$, p = 1
**2f[b]:** R = $CH_2C_6H_6$, p = 1
**2h[b]:** R = $CH_3$, p = 1
**2i[b]:** R = $CH(CH_3)_2$, p = 1
**p = m + n**
**2k:** R = $CH_2CH(CH_3)_2$, p = 2
**2l:** R = $CH_2CH(CH_3)_2$, p = 3
**2o:** R= $CH_2C_6H_6$, p = 2
**2p:** R= $CH_2C_6H_6$, p = 3
**2q:** R= $CH_3$, p = 2
**2r:** R= $CH_3$, p = 3
**2s:** R= $CH(CH_3)_2$, p = 2
**2t:** R= $CH(CH_3)_2$, p = 3

**4a:** R = H, m = 1
**4b:** R = H, m = 2
**4c:** R = H, m = 3
**4d:** R = $CH_2CH(CH_3)_2$, m = 1
**4f:** R = $CH_2C_6H_6$, m = 1
**4h:** R= $CH_3$, m = 1
**4i:** R= $CH(CH_3)_2$, m = 1
**4k:** R= $CH_2CH(CH_3)_2$, m = 2
**4l:** R= $CH_2CH(CH_3)_2$, m = 3
**4o:** R= $CH_2C_6H_6$, m = 2
**4p:** R= $CH_2C_6H_6$, m = 3
**4q:** R= $CH_3$, m = 2
**4r:** R= $CH_3$, m = 3
**4s:** R= $CH(CH_3)_2$, m = 2
**4t:** R= $CH(CH_3)_2$, m = 3

**Scheme S1.** Synthesis of npe-protected amino acids and peptides **4a**-**4t**. [a] The boc-protected peptides **1b** and **1c** were synthesized from the unprotected peptides by using $Boc_2O$ and TEA in methanol. [b] The compounds were not purified and the crude compounds were used for further reactions.

##### General procedure for the syntheses of 2a-i

The reactions were performed similar to a procedure published before.[1]

The *N*-Boc-protected amino acids or peptides **1a** – **i** (1.0 equiv.) were dissolved in dry DCM under inert atmosphere and cooled to 0°C. Subsequently, npeOH (1.3 equiv.) and $PPh_3$ were

added to the cold solution. DIAD (1.3 equiv.) was added dropwise and the reaction mixture was stirred overnight at r.t.. The reaction mixtures were diluted with DCM and washed with water twice, aqueous saturated solution of $NaHCO_3$ twice and brine once. The solvents were removed *in vacuo* and the crude products were purified by flash column chromatography to afford the target compounds **2a-i**.
(Some of the compounds did not separate well from side products by flash chromatography. However, the following deprotections of the boc-groups were compatible with the crude compounds and washing with diethyl ether removed side products efficiently.)

**Compound 2a**

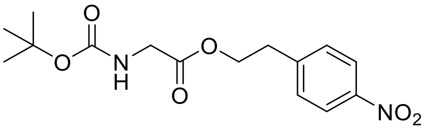


Eluent: DCM
**Yield:** 86%; ***R*f:** 0.50 (95:5 DCM/MeOH); **$^1$H NMR (500 MHz, $CDCl_3$) δ:** 8.19 – 8.13 (m, 2H), 7.41 – 7.34 (m, 2H), 4.99 (s, 1H), 4.39 (t, *J* = 6.7 Hz, 2H), 3.87 (d, *J* = 5.8 Hz, 2H), 3.06 (t, *J* = 6.7 Hz, 2H), 1.43 (s, 9H); **HRMS (ESI):** Calcd. for $C_{15}H_{20}N_2O_6Na^+$: *m/z* = 347.1219 [M+Na]$^+$, found: *m/z*: 347.1215 [M+Na]$^+$.
Literature: ref.[1,2]

**Compound 2b**

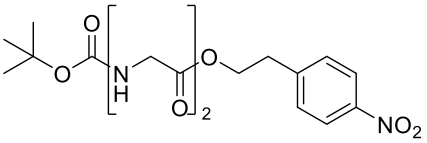


Eluent: DCM/MeOH (99/1, v/v).
**Yield:** 60%; ***R*f:** 0.32 (95:5 DCM/MeOH); **IR:** ṽ (cm$^{-1}$) = 3295 (w), 2979 (w), 1750 (m), 1710 (m), 1677 (m), 1600 (w), 1516 (s), 1437 (w), 1367 (w), 1344 (s), 1249 (w), 1165 (vs), 1118 (s), 1029 (w), 946 (w), 856 (m), 749 (m), 720 (s), 694 (s), 537 (s); **$^1$H NMR (500 MHz, $CDCl_3$) δ:** 8.20 – 8.12 (m, 2H), 7.41 – 7.33 (m, 2H), 6.73 (d, *J* = 5.6 Hz, 1H), 5.22 (s, 1H), 4.39 (t, *J* = 6.7 Hz, 2H), 4.01 (d, *J* = 5.4 Hz, 2H), 3.83 (d, *J* = 5.9 Hz, 2H), 3.06 (t, *J* = 6.7 Hz, 2H), 1.44 (s, 9H); **$^{13}$C NMR (126 MHz, $CDCl_3$) δ:** 170.0, 169.6, 156.2, 147.1, 145.3, 129.9, 124.0, 80.6, 64.9, 44.4, 41.2, 34.9, 28.4; **HRMS (ESI):** Calcd. for $C_{17}H_{24}N_3O_7Na^+$: *m/z* = 404.1434 [M+Na]$^+$, found: *m/z*: 404.1431 [M+Na]$^+$.

**Combound 2c**

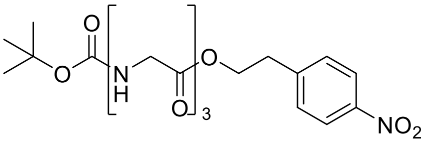


Eluent: DCM/MeOH (96/4, v/v).
**Yield:** 45%; ***R*f:** 0.27 (95:5 DCM/MeOH); **IR:** ṽ (cm$^{-1}$) = 3305 (s), 2982 (w), 2936 (w), 1753 (m), 1706 (w), 1676 (w), 1645 (s), 1604 (w), 1513 (vs), 1439 (w), 1394 (w), 1368 (w), 1344 (s), 1250 (w), 1184 (s), 1164 (m), 1109 (w), 1052 (m), 1029 (m), 987 (m), 945 (w), 855 (s), 783 (w), 769 (w), 747 (w), 697 (m), 558 (w), 527 (w), 499 (w), 463 (w); **$^1$H NMR (500 MHz, $CDCl_3$) δ:** 8.20 – 8.13 (m, 2H), 7.42 – 7.35 (m, 2H), 7.06 (m, 2H), 5.37 (t, *J* = 5.8 Hz, 1H), 4.38 (t, *J* = 6.7 Hz, 2H), 4.00 (d, *J* = 5.7 Hz, 2H), 3.97 (d, *J* = 5.6 Hz, 2H), 3.81 (d, *J* = 5.8 Hz, 2H), 3.06 (t, *J* = 6.7 Hz, 2H),

1.42 (s, 9H); **$^{13}$C NMR (126 MHz, $CDCl_3$) δ:** 170.4, 169.7, 169.4, 156.6, 147.1, 145.3, 129.9, 124.0, 80.8, 65.0, 44.6, 43.0, 41.3, 34.9, 28.4; **HRMS (ESI):** Calcd. for $C_{19}H_{26}N_4O_8Na^+$: *m/z* = 461.1648 [M+Na]$^+$, found: *m/z*: 461.1638 [M+Na]$^+$.

**Compound 2d**

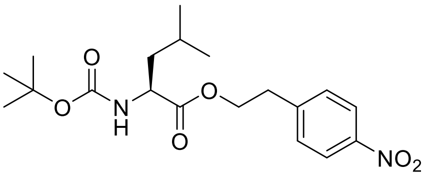

Eluent: Hex/EtOAc (85/15, v/v).
**Yield:** 91%; ***R*f:** 0.47 (3:1 Hex/EtOAc); **IR:** $\tilde{\nu}$ (cm$^{-1}$) = 3380 (w), 2960 (m), 2937 (w), 2871 (w), 2363 (w), 1742 (m), 1707 (s), 1600 (w), 1519 (s), 1472 (w), 1457 (w), 1390 (w), 1367 (m), 1344 (s), 1272 (w), 1249 (m), 1224 (w), 1158 (s), 1109 (m), 1048 (m), 1016 (m), 957 (w), 856 (m), 773 (w), 749 (m), 698 (m) 617 (w), 536 (w); **$^1$H NMR (500 MHz, $CDCl_3$) δ:** 8.19 – 8.13 (m, 2H), 7.41 – 7.35 (m, 2H), 4.82 (d, *J* = 8.7 Hz, 1H), 4.37 (td, *J* = 6.7, 3.6 Hz, 2H), 4.24 (td, *J* = 8.9, 5.6 Hz, 1H), 3.06 (t, *J* = 6.7 Hz, 2H), 1.59 (tdd, *J* = 16.7, 11.1, 4.5 Hz, 1H), 1.51 – 1.34 (m, 11H), 0.87 (dd, *J* = 9.3, 6.6 Hz, 6H); **$^{13}$C NMR (126 MHz, $CDCl_3$) δ:** 173.5, 155.5, 147.0, 145.5, 129.9, 123.9, 80.1, 64.6, 52.1, 41.8, 35.0, 28.4, 24.9, 22.9, 22.1, 22.0; **HRMS (ESI):** Calcd. for $C_{19}H_{28}N_2O_6Na^+$: *m/z* = 403.1845 [M+Na]$^+$, found: *m/z*: 403.1839 [M+Na]$^+$.

**Compound 2f**

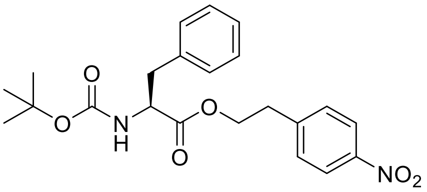

The crude product was used in the following step without further purification.

**Compound 2h**

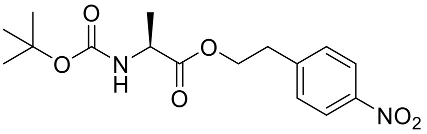

The crude product was used in the following step without further purification.

**Compound 2i**

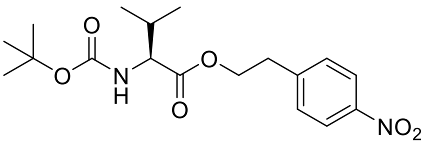

The crude product was used in the following step without further purification.

**General procedure for synthesizing boc- and npe-protected polypeptides 2k-t**
The reaction was performed similar to a procedure published before.[3]
The npe-protected amino acids/peptides **4d**, **4f**, **4h**, **4i**, **4k, 4o, 4q** or **4s** (1.0 equiv.) and the corresponding boc-protected amino acids **1d**, **1f**, **1h** or **1i** (1.2 equiv.) were dissolved in DMF and HOBT·$H_2O$ (1.2 equiv.), EDC-Cl (1.2 equiv.) and TEA (2.3 equiv.) were added to the stirred solution. The resulting mixture was stirred overnight at r.t., and the solvent was removed *in*

*vacuo*. The residue was dissolved in EtOAc and washed with water, aqueous solution of citric acid (10%) (three times), aqueous saturated solution of $NaHCO_3$ (two times) and with brine. Drying the organic solvents over $Na_2SO_4$ and removing of the solvents *in vacuo* yielded the target compounds **2k**-**t** as yellowish foams. Compounds which showed impurities on TLC or NMR were further purified by flash column chromatography.

**Compound 2k**

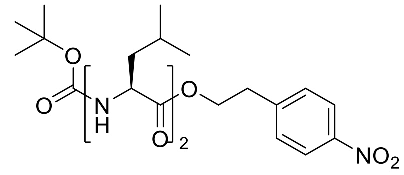

Eluent: Hex/EtOAc (80/20, v/v).

**Yield:** 85%; ***R*f:** 0.49 (6:4 Hex/EtOAc); **IR:** $\tilde{\nu}$ ($cm^{-1}$) = 3315 (m), 2957 (m), 2935 (w), 2871 (w), 1746 (m), 1686 (w), 1651 (s), 1602 (w), 1519 (s), 1469 (w), 1391 (w), 1366 (w), 1345 (s), 1321 (w), 1295 (w), 1272 (w), 1247 (m), 1164 (vs), 1048 (m), 1026 (w), 877 (w), 855 (m), 747 (w), 697 (w), 632 (w), 589 (w), 535 (w), 500 (w); **$^1$H NMR (500 MHz, $CDCl_3$) δ:** 8.20 – 8.13 (m, 2H), 7.42 – 7.35 (m, 2H), 6.47 (d, *J* = 8.0 Hz, 1H), 4.84 (d, *J* = 8.3 Hz, 1H), 4.53 (ddd, *J* = 9.1, 8.2, 5.2 Hz, 1H), 4.37 (qt, *J* = 11.0, 6.7 Hz, 2H), 4.08 (q, *J* = 7.9 Hz, 1H), 3.06 (t, *J* = 6.5 Hz, 2H), 1.72 – 1.43 (m, 6H), 1.43 (s, 9H), 0.92 (dd, *J* = 8.1, 6.2 Hz, 6H), 0.86 (dd, *J* = 6.3, 5.4 Hz, 6H); **$^{13}$C NMR (126 MHz, $CDCl_3$) δ:** 172.7, 172.4, 155.9, 147.1, 145.4, 129.9, 123.9, 80.3, 64.7, 53.0, 50.7, 41.5, 40.7, 34.9, 28.4, 24.8, 24.8, 23.0, 22.9, 22.2, 21.9; **HRMS (ESI):** Calcd. for $C_{25}H_{39}N_3O_7Na^+$: *m/z* = 516.2685 $[M+Na]^+$, found: *m/z*: 516.2675 $[M+Na]^+$.

**Compound 2l**

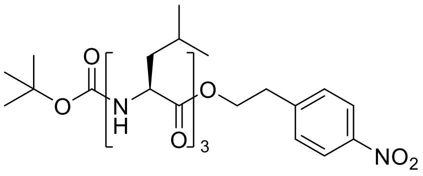

Eluent: Hex/EtOAc (70/30, v/v).

**Yield:** 95%; ***R*f:** 0.26 (7:3 Hex/EtOAc); **IR:** $\tilde{\nu}$ ($cm^{-1}$) = 3287 (m), 2957 (m), 2934 (w), 2871 (w), 1743 (m), 1690 (w), 1643 (s), 1607 (w), 1519 (s), 1470 (w), 1455 (w), 1388 (w), 1367 (m), 1345 (s), 1321 (w), 1246 (m), 1161 (s), 1046 (m), 1017 (m), 856 (m), 747 (w), 697 (w), 536 (w), 468 (w); **$^1$H NMR (500 MHz, $CDCl_3$) δ:**8.21 – 8.14 (m, 2H), 7.43 – 7.36 (m, 2H), 6.59 (d, *J* = 8.0 Hz, 1H), 6.47 (d, *J* = 8.0 Hz, 1H), 4.85 (d, *J* = 7.5 Hz, 1H), 4.49 (td, *J* = 8.3, 5.1 Hz, 1H), 4.46 – 4.31 (m, 3H), 4.09 – 4.02 (m, 1H), 3.07 (t, *J* = 6.7 Hz, 2H), 1.72 – 1.59 (m, 4H), 1.57 – 1.44 (m, 5H), 1.43 (s, 9H), 0.97 – 0.82 (m, 18H); **$^{13}$C NMR (126 MHz, $CDCl_3$) δ:** 172.7, 172.4, 171.6, 155.9, 146.9, 145.4, 129.9, 123.8, 80.5, 64.6, 53.2, 51.5, 50.8, 41.2, 40.7, 40.2, 34.8, 28.3, 24.8, 24.7, 24.6, 23.0, 22.9, 22.7, 22.0, 21.8; **HRMS (ESI):** Calcd. for $C_{31}H_{50}N_4O_8Na^+$: *m/z* = 629.3526 $[M+Na]^+$, found: *m/z*: 629.3521 $[M+H]^+$.

**Compound 2o**

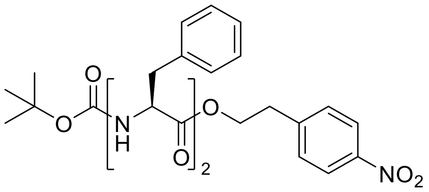

The compound was used after extraction with column purification.

**Yield:** 95%; ***R*f:** 0.84 (95:5 DCM/MeOH); **IR:** $\tilde{\nu}$ (cm$^{-1}$) = 3335 (w), 3029 (w), 2972 (w), 2934 (w), 1740 (m), 1681 (m), 1666 (s), 1605 (w), 1512 (vs), 1495 (m), 1454 (m), 1447 (m), 1391 (m), 1368 (m), 1351 (s), 1319 (m), 1292 (m), 1246 (m), 1223 (m), 1165 (s), 1111 (m), 1081 (m), 1042 (m), 1030 (m), 1019 (m), 1007 (m), 1002 (m), 862 (m), 851 (m), 747 (m), 697 (s) 625 (m), 594 (m), 566 (m), 522 (m), 507 (m); **$^1$H NMR (500 MHz, $CD_2Cl_2$) δ:** 8.18 – 8.12 (m, 2H), 7.38 – 7.32 (m, 2H), 7.31 – 7.25 (m, 2H), 7.25 – 7.20 (m, 4H), 7.16 (m, 2H), 6.98 – 6.91 (m, 2H), 6.38 (s, 1H), 4.95 – 4.86 (m, 1H), 4.72 (dt, *J* = 7.7, 6.1 Hz, 1H), 4.38 – 4.23 (m, 3H), 3.11 – 2.90 (m, 6H), 1.37 (s, 9H); **$^{13}$C NMR (126 MHz, $CD_2Cl_2$) δ:** 171.3, 171.2, 155.7, 147.4, 146.0, 137.1, 136.3, 132.4, 130.3, 129.7, 129.6, 129.0, 128.9, 127.4, 127.3, 124.1, 80.5, 65.3, 56.0, 53.7, 38.3, 35.0, 28.3; **HRMS (ESI):** Calcd. for $C_{31}H_{35}N_3O_7Na^+$: *m/z* = 584.2373 [M+Na]$^+$, found: *m/z*: 584.2374 [M+Na]$^+$.

**Compound 2p**

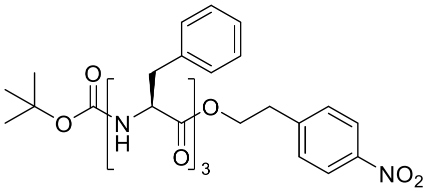

The compound was used after extraction with column purification.

**Yield:** 93%; ***R*f:** 0.58 (95:5 DCM/MeOH); **IR:** $\tilde{\nu}$ (cm$^{-1}$) = 3284 (w), 3064 (w), 3031 (w), 2975 (w), 2930 (w), 1742 (m), 1716 (m), 1684 (m), 1670 (m), 1647 (s), 1604 (m), 1559 (m), 1541 (m), 1516 (vs), 1497 (s), 1456 (m), 1393 (m), 1364 (m), 1344 (vs), 1319 (m), 1249 (m), 1213 (m), 1165 (s), 1109 (m), 1081 (m), 1016 (m), 856 (m), 744 (s), 697 (vs), 632 (m), 618 (m), 594 (m), 532 (m), 493 (m), 474 (m), 461 (m), 448 (m), 434 (m), 415 (m); **$^1$H NMR (500 MHz, $CD_2Cl_2$) δ:** 8.19 – 8.09 (m, 2H), 7.38 – 7.31 (m, 2H), 7.33 – 7.17 (m, 9H), 7.18 – 7.12 (m, 2H), 7.08 – 7.03 (m, 2H), 7.01 – 6.95 (m, 2H), 6.48 (s, 1H), 6.45 (d, *J* = 7.8 Hz, 1H), 4.89 – 4.78 (m, 1H), 4.65 (dt, *J* = 7.7, 6.6 Hz, 1H), 4.56 (dt, *J* = 7.9, 6.6 Hz, 1H), 4.29 (ddt, *J* = 30.7, 10.8, 6.7 Hz, 3H), 3.12 – 2.85 (m, 8H), 1.34 (s, 9H); **$^{13}$C NMR (126 MHz, $CD_2Cl_2$) δ:** 171.5, 171.2, 170.4, 155.9, 147.3, 146.1, 137.0, 136.8, 136.4, 132.4, 132.3, 130.3, 129.7, 129.6, 129.5, 129.1, 129.0, 128.9, 127.4, 127.4, 127.4, 124.0, 80.7, 65.2, 56.3, 38.2, 37.9, 35.0, 28.3; **HRMS (ESI):** Calcd. for $C_{40}H_{45}N_4O_8^+$: *m/z* = 709.3237 [M+H]$^+$, found: *m/z*: 709.3240 [M+H]$^+$.

**Compound 2q**

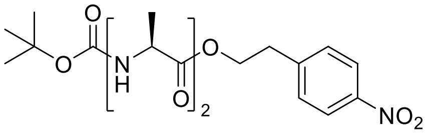

The compound was used after extraction with column purification.

**Yield:** 98%; ***R*f:** 0.44 (95:5 DCM/MeOH); **IR:** $\tilde{\nu}$ (cm$^{-1}$) = 3311 (w), 2980 (w), 2937 (w), 1740 (m), 1656 (s), 1601 (m), 1559 (m), 1541 (m), 1518 (vs), 1497 (s), 1456 (m), 1393 (m), 1365 (m), 1344 (vs), 1319 (m), 1247 (m), 1160 (s), 1109 (m), 1069 (m), 1049 (m), 1017 (m), 855 (m), 747 (m), 698 (s), 631 (m), 617 (m), 533 (m), 502 (m), 493 (m), 480 (m), 473 (m), 448 (m), 420 (m), 410 (m); **$^1$H NMR (500 MHz, CDCl$_3$) δ:** 8.21 – 8.14 (m, 2H), 7.42 – 7.35 (m, 2H), 6.66 (s, 1H), 4.99 (s, 1H), 4.50 (p, *J* = 7.2 Hz, 1H), 4.39 (t, *J* = 6.7 Hz, 2H), 4.15 (s, 1H), 3.07 (t, *J* = 6.7 Hz, 2H), 1.43 (s, 9H), 1.32 (dd, *J* = 7.2, 6.1 Hz, 6H); **$^{13}$C NMR (126 MHz, CDCl$_3$) δ:** 172.7, 172.4, 155.7, 147.1, 145.4, 129.9, 123.9, 80.4, 64.8, 50.0, 48.1, 34.9, 28.4, 18.3, 18.2; **HRMS (ESI):** Calcd. for $C_{19}H_{27}N_3O_7Cl^-$: *m/z* = 444.1538 [M+Cl]$^-$, found: *m/z*: 444.1541 [M+Cl]$^-$.

**Compound 2r**

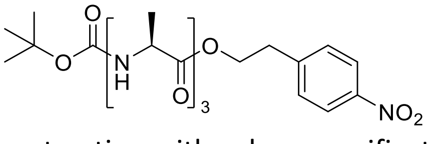

The compound was used after extraction with column purification.

**Yield:** 63%; ***R*$_f$:** 0.40 (95:5 DCM/MeOH); **IR:** $\tilde{\nu}$ (cm$^{-1}$) = 3334 (w), 3304 (w), 3279 (w), 2980 (w), 2932 (w), 1733 (m), 1699 (m), 1670 (m), 1637 (s), 1558 (m), 1541 (m), 1515 (vs), 1490 (m), 1459 (m), 1447 (m), 1393 (m), 1365 (m), 1342 (s), 1318 (m), 1270 (m), 1250 (m), 1239 (m), 1219 (m), 1165 (s), 1117 (m), 1105 (m), 1052 (m), 1000 (m), 948 (w), 855 (m), 782 (w), 749 (w), 713 (m), 696 (m), 657 (m), 536 (m), 418 (m), 401 (m); **$^1$H NMR (500 MHz, DMSO-*d*$_6$) δ**: 8.24 (d, *J* = 7.0 Hz, 1H), 8.19 – 8.13 (m, 2H), 7.78 (d, *J* = 7.6 Hz, 1H), 7.60 – 7.53 (m, 2H), 6.94 (d, *J* = 7.7 Hz, 1H), 4.38 – 4.22 (m, 3H), 4.19 (p, *J* = 7.3 Hz, 1H), 3.98 – 3.90 (m, 1H), 3.04 (t, *J* = 6.4 Hz, 2H), 1.36 (s, 9H), 1.19 (d, *J* = 7.3 Hz, 3H), 1.15 (dd, *J* = 7.1, 4.1 Hz, 6H); **$^{13}$C NMR (126 MHz, DMSO-*d*$_6$) δ:** 172.3, 172.2, 172.1, 155.1, 146.5, 146.3, 130.3, 123.4, 78.1, 64.1, 49.6, 47.5, 34.0, 28.2, 18.3, 18.0, 16.8; **HRMS (ESI):** Calcd. for $C_{22}H_{32}N_4O_8Na^+$: *m/z* = 503.2118 [M+Na]$^+$, found: *m/z*: 503.2107 [M+Na]$^+$.

**Compound 2s**

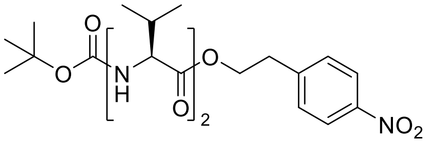

The compound was used after extraction with column purification.

**Yield:** 96%; **$^1$H NMR (500 MHz, CDCl$_3$) δ:** 8.21 – 8.13 (m, 2H), 7.43 – 7.35 (m, 2H), 6.37 (d, *J* = 8.7 Hz, 1H), 5.02 (d, *J* = 8.7 Hz, 1H), 4.48 (dd, *J* = 8.6, 4.9 Hz, 1H), 4.45 – 4.33 (m, 2H), 3.88 (dd, *J* = 8.7, 6.5 Hz, 1H), 3.07 (td, *J* = 6.7, 1.4 Hz, 2H), 2.08 (ddtt, *J* = 15.8, 8.9, 7.0, 3.1 Hz, 2H), 1.43 (s, 9H), 0.93 (dd, *J* = 11.3, 6.8 Hz, 6H), 0.87 (d, *J* = 6.9 Hz, 3H), 0.81 (d, *J* = 6.9 Hz, 3H); **HRMS (ESI):** Calcd. for $C_{23}H_{35}N_3O_7Na^+$: *m/z* = 488.2373 [M+Na]$^+$, found: *m/z*: 488.2362 [M+Na]$^+$.
Literature: ref.[4]

**Compound 2t**

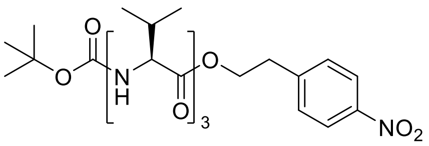

The compound was used after extraction with column purification.

**Yield:** 80%; **$^1$H NMR (500 MHz, CDCl$_3$) δ:** 8.20 – 8.13 (m, 2H), 7.42 – 7.36 (m, 2H), 6.56 (d, *J* = 8.6 Hz, 2H), 5.11 (d, *J* = 8.2 Hz, 1H), 4.46 (dd, *J* = 8.6, 5.1 Hz, 1H), 4.39 (td, *J* = 6.7, 3.8 Hz, 2H), 4.27 (t, *J* = 7.7 Hz, 1H), 3.94 – 3.87 (m, 1H), 3.07 (t, *J* = 6.7 Hz, 2H), 2.21 – 2.01 (m, 3H), 1.43 (s, 9H), 0.96 – 0.89 (m, 12H), 0.85 (d, *J* = 6.8 Hz, 3H), 0.80 (d, *J* = 6.9 Hz, 3H); **HRMS (ESI):** Calcd. for $C_{28}H_{44}N_4O_8Na^+$: *m/z* = 587.3057 [M+Na]$^+$, found: *m/z*: 587.3049 [M+Na]$^+$.
Literature: ref.[4]

**General procedure for the boc-deprotection of 2a-t yielding 4a-t**
The reaction was performed according to our previously published procedure.[1]
The boc- and npe-protected amino acids/peptides **2a-t** (1.0 equiv.) were dissolved in a cooled 4M solution of HCl in dioxane (15 equiv. of HCl) and cooled to 0°C. After 15min the cooling was removed, and the resulting suspension was stirred for additional 2 h at r.t.. Afterwards the solvents were removed *in vacuo*, the resulting crude products were resuspended in diethyl ether and filtered. Washing the filtrates with diethyl ether and drying yielded the target compounds **4a-u** as colourless solids.

**Compound 4a**

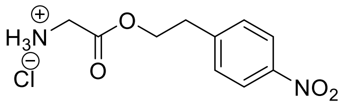

**Yield:** 87%; **$^1$H NMR (500 MHz, DMSO-*d*$_6$) δ:** 8.46 (s, 3H), 8.20 – 8.15 (m, 2H), 7.63 – 7.57 (m, 2H), 4.43 (t, *J* = 6.4 Hz, 2H), 3.75 (s, 2H), 3.09 (t, *J* = 6.4 Hz, 2H); **HRMS (ESI):** Calcd. for $C_{10}H_{13}N_2O_4^+$: *m/z* = 225.0875 [M+H]$^+$, found: *m/z*: 225.0869 [M+H]$^+$.
Literature: ref.[1,2]

**Compound 4b**

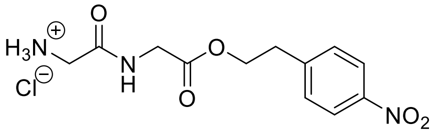

**Yield:** 53% over npe-protection and boc-deprotection steps; **$^1$H NMR (500 MHz, DMSO-*d*$_6$) δ:** 8.93 (t, *J* = 5.9 Hz, 1H), 8.23 (s, 3H), 8.21 – 8.14 (m, 2H), 7.60 – 7.54 (m, 2H), 4.34 (t, *J* = 6.5 Hz, 2H), 3.91 (d, *J* = 5.9 Hz, 2H), 3.59 (s, 2H), 3.06 (t, *J* = 6.5 Hz, 2H); **HRMS (ESI):** Calcd. for $C_{12}H_{16}N_3O_5^+$: *m/z* = 282.1090 [M+H]$^+$, found: *m/z*: 282.1084 [M+H]$^+$.
Literature: ref.[5]

**Compound 4c**

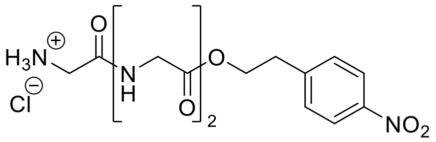

**Yield:** 98%; ***R*$_f$:** (95: 5 DCM/MeOH); **IR:** $\tilde{\nu}$ (cm$^{-1}$) = 3333 (w), 3202 (w), 2870 (m), 1735 (s), 1671 (w), 1640 (w), 1549 (s), 1519 (s), 1474 (m), 1344 (m), 1199 (w), 1030 (w), 957 (m), 907 (m), 858 (m), 696 (m), 612 (m), 594 (m), 503 (w), 460 (w); **$^1$H NMR (500 MHz, DMSO-*d*$_6$) δ:** 8.79 (t, *J* = 5.9 Hz, 1H), 8.53 (t, *J* = 5.9 Hz, 1H), 8.26 – 8.22 (m, 3H), 8.19 – 8.14 (m, 2H), 7.59 – 7.53 (m, 2H), 4.31 (t, *J* = 6.4 Hz, 2H), 3.80 (dd, *J* = 5.9, 4.3 Hz, 4H), 3.59 (s, 2H), 3.04 (t, *J* = 6.4 Hz, 2H); **$^{13}$C NMR (126 MHz, DMSO-*d*$_6$) δ**: 169.6, 168.9, 166.4, 146.5, 146.3, 130.3, 123.4, 64.2, 41.8,

40.6, 40.2, 34.0; **HRMS (ESI):** Calcd. for $C_{14}H_{19}N_4O_6^+$: *m/z* = 339.1305 [M+H]$^+$, found: *m/z*: 339.1299 [M+H]$^+$.

**Compound 4d**

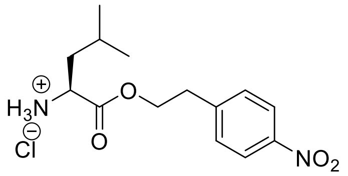


**Yield:** 97%; **$^1$H NMR (500 MHz, DMSO-*d*$_6$) δ:** 8.58 (s, 3H), 8.21 – 8.15 (m, 2H), 7.64 – 7.57 (m, 2H), 4.45 (ddt, *J* = 34.2, 11.0, 6.3 Hz, 2H), 3.83 (t, *J* = 6.6 Hz, 1H), 3.11 (t, *J* = 6.3 Hz, 2H), 1.59 – 1.40 (m, 3H), 0.76 (t, *J* = 6.1 Hz, 6H); **HRMS (ESI):** Calcd. for $C_{14}H_{21}N_2O_4^+$: *m/z* = 281.1501 [M+H]$^+$, found: *m/z*: 281.1494 [M+H]$^+$.
Literature: ref.[5]

**Compound 4f**

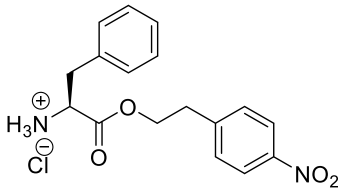


**Yield:** 74% for npe-protection and boc-deprotection step; **$^1$H NMR (500 MHz, DMSO-*d*$_6$) δ:** 8.67 (s, 3H), 8.18 – 8.12 (m, 2H), 7.51 – 7.46 (m, 2H), 7.32 – 7.22 (m, 3H), 7.14 – 7.08 (m, 2H), 4.33 (t, *J* = 6.3 Hz, 2H), 4.20 (dd, *J* = 7.8, 5.6 Hz, 1H), 3.14 (dd, *J* = 14.0, 5.6 Hz, 1H), 3.07 – 2.88 (m, 3H); **HRMS (ESI):** Calcd. for $C_{17}H_{19}N_2O_4^+$: *m/z* = 315.1345 [M+H]$^+$, found: *m/z*: 315.1338 [M+H]$^+$.
Literature: ref.[1]

**Compound 4h**

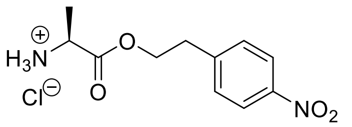


**Yield:** 71% for npe-protection and boc-deprotection step; **$^1$H NMR (500 MHz, $D_2O$) δ:** 8.23 – 8.16 (m, 2H), 7.57 – 7.49 (m, 2H), 4.61 (dt, *J* = 10.9, 6.4 Hz, 1H), 4.53 (dt, *J* = 10.7, 6.2 Hz, 1H), 4.15 (q, *J* = 7.3 Hz, 1H), 3.18 (t, *J* = 6.3 Hz, 2H), 1.46 (d, *J* = 7.3 Hz, 3H); **HRMS (ESI):** Calcd. for $C_{11}H_{15}N_2O_4^+$: *m/z* = 239.1032 [M+H]$^+$, found: *m/z*: 239.1026 [M+H]$^+$.
Literature: ref.[5]

**Compound 4i**

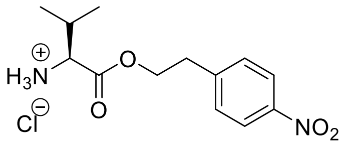


**Yield:** 93% for npe-protection and boc-deprotection step; **$^1$H NMR (500 MHz, DMSO-*d*$_6$) δ:** 8.58 (s, 3H), 8.21 – 8.14 (m, 2H), 7.65 – 7.58 (m, 2H), 4.47 (ddt, *J* = 34.6, 11.0, 6.4 Hz, 2H), 3.78 (d, *J* = 4.6 Hz, 1H), 3.12 (t, *J* = 6.3 Hz, 2H), 2.09 (heptd, *J* = 7.0, 4.6 Hz, 1H), 0.83 (dd, *J* = 6.9, 2.2 Hz, 6H); **HRMS (ESI):** Calcd. for $C_{13}H_{19}N_2O_4^+$: *m/z* = 267.1345 [M+H]$^+$, found: *m/z*: 267.1338 [M+H]$^+$.
Literature: ref.[1]

**Compound 4k**

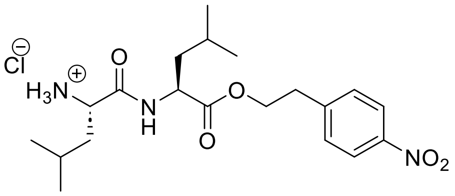

**Yield:** 93%; **IR:** $\tilde{\nu}$ (cm$^{-1}$) = 3384 (w), 2959 (w), 2929 (w), 2874 (w), 1722 (s), 1670 (s), 1605 (w), 1522 (s), 1467 (w), 1393 (w), 1345 (s), 1330 (w), 1319 (w), 1275 (m), 1245 (w), 1196 (w), 1143 (w), 1130 (w), 1109 (w), 1069 (w), 1016 (w), 999 (w), 973 (w), 857 (m), 849 (m), 747 (w), 697 (m), 655 (w), 594 (m), 525 (w), 510 (w), 454 (w); **$^1$H NMR (500 MHz, DMSO-*d*$_6$) δ:** 8.88 (d, *J* = 7.3 Hz, 1H), 8.29 (s, 3H), 8.21 – 8.14 (m, 2H), 7.59 – 7.53 (m, 2H), 4.34 (qt, *J* = 11.0, 6.4 Hz, 2H), 4.23 (ddd, *J* = 10.0, 7.2, 5.1 Hz, 1H), 3.78 (t, *J* = 7.2 Hz, 1H), 3.05 (t, *J* = 6.4 Hz, 2H), 1.75 – 1.28 (m, 6H), 0.87 (dd, *J* = 9.3, 6.5 Hz, 6H), 0.81 (d, *J* = 6.6 Hz, 3H), 0.78 (d, *J* = 6.5 Hz, 3H); **$^{13}$C NMR (126 MHz, DMSO-*d*$_6$) δ:** 171.8, 169.3, 146.5, 146.3, 130.4, 123.4, 64.3, 50.5, 50.5, 40.2, 34.0, 24.0, 23.4, 22.6, 22.6, 22.1, 21.3; **HRMS (ESI):** Calcd. for $C_{20}H_{32}N_3O_5^+$: *m/z* = 394.2342 [M+H]$^+$, found: *m/z*: 394.2336 [M+H]$^+$.

**Compound 4l**

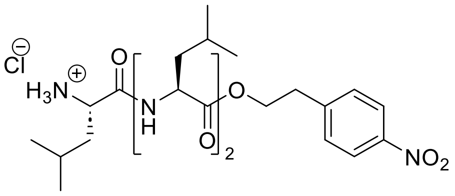

**Yield:** 75%; **IR:** $\tilde{\nu}$ (cm$^{-1}$) = 3340 (w), 2963 (w), 2929 (w), 2877 (w), 1750 (s), 1673 (m), 1644 (s), 1601 (w), 1522 (vs), 1472 (w), 1388 (w), 1345 (vs), 1280 (w), 1250 (w), 1204 (w), 1191 (m), 1157 (m), 1108 (w), 996 (w), 856 (m), 749 (m), 618 (w), 539 (w), 471 (w); **$^1$H NMR (500 MHz, DMSO-*d*$_6$) δ:** 8.66 (d, *J* = 8.3 Hz, 1H), 8.41 (d, *J* = 7.7 Hz, 1H), 8.26 (s, 3H), 8.21 – 8.12 (m, 2H), 7.59 – 7.52 (m, 2H), 4.43 – 4.25 (m, 3H), 4.21 (ddd, *J* = 10.3, 7.7, 4.7 Hz, 1H), 3.76 (t, *J* = 6.9 Hz, 1H), 3.04 (t, *J* = 6.4 Hz, 2H), 1.70 – 1.37 (m, 8H), 1.29 (ddd, *J* = 13.7, 9.1, 4.7 Hz, 1H), 0.93 – 0.82 (m, 12H), 0.80 (d, *J* = 6.5 Hz, 3H), 0.75 (d, *J* = 6.4 Hz, 3H); **$^{13}$C NMR (126 MHz, DMSO-*d*$_6$) δ:** 172.1, 171.5, 168.5, 146.5, 146.2, 130.3, 123.3, 64.1, 50.8, 50.7, 50.1, 40.9, 40.3, 39.4, 34.0, 24.0, 23.9, 23.5, 22.9, 22.7, 22.5, 22.3, 21.9, 21.0; **HRMS (ESI):** Calcd. for $C_{26}H_{43}N_4O_6^+$: *m/z* = 507.3183 [M+H]$^+$, found: *m/z*: 507.3168 [M+H]$^+$.

**Compound 4o**

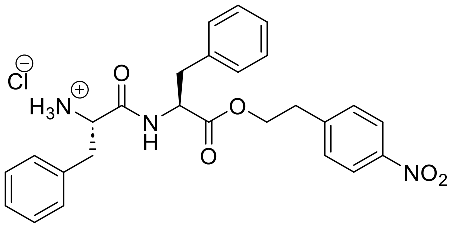

**Yield:** 98%; **IR:** $\tilde{\nu}$ (cm$^{-1}$) = 3347 (w), 3031 (w), 2927 (w), 2857 (w), 1740 (w), 1720 (w), 1686 (m), 1601 (w), 1551 (w), 1515 (s), 1497 (m), 1456 (w), 1344 (s), 1255 (m), 1180 (m), 1109 (w), 1065 (w), 1016 (w), 856 (m), 744 (s), 698 (vs), 619 (w), 595 (w), 529 (w); **$^1$H NMR (500 MHz, DMSO-*d*$_6$) δ:** 9.29 (d, *J* = 7.4 Hz, 1H), 8.30 – 8.21 (m, 3H), 8.17 – 8.11 (m, 2H), 7.55 – 7.49 (m, 2H), 7.33 – 7.15 (m, 10H), 4.53 – 4.44 (m, 1H), 4.36 – 4.21 (m, 2H), 4.07 (s, 1H), 3.13 – 2.85 (m, 6H); **$^{13}$C NMR (126 MHz, DMSO-*d*$_6$) δ:** 170.6, 168.3, 146.4, 146.3, 136.7, 134.8, 130.3,

129.6, 129.1, 128.4, 128.4, 128.3, 127.1, 126.7, 123.4, 64.5, 54.0, 53.1, 36.8, 36.6, 33.9; **HRMS (ESI):** Calcd. for $C_{26}H_{28}N_3O_5^+$: *m/z* = 462.2029 [M+H]$^+$, found: *m/z*: 462.2016 [M+H]$^+$.

**Compound 4p**

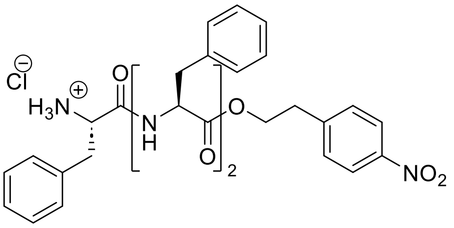


**Yield:** 98%; **IR:** ṽ (cm$^{-1}$) = 3212 (w), 3031 (w), 2929 (w), 2855 (w), 1736 (m), 1656 (s), 1602 (w), 1516 (s), 1497 (m), 1456 (w), 1344 (s), 1181 (m), 1109 (m), 1081 (w), 920 (w), 856 (m), 744 (s), 698 (vs), 540 (w), 484 (w); **$^1$H NMR (500 MHz, DMSO-*d*$_6$) δ:** 8.93 (d, *J* = 8.3 Hz, 1H), 8.76 (d, *J* = 7.5 Hz, 1H), 8.16 – 8.08 (m, 5H), 7.54 – 7.48 (m, 2H), 7.29 – 7.15 (m, 15H), 4.59 (td, *J* = 8.9, 4.8 Hz, 1H), 4.48 (td, *J* = 8.0, 6.3 Hz, 1H), 4.27 (qt, *J* = 11.1, 6.5 Hz, 2H), 3.98 (s, 1H), 3.13 (dd, *J* = 14.2, 4.7 Hz, 1H), 3.06 – 2.83 (m, 6H), 2.82 – 2.71 (m, 1H); **$^{13}$C NMR (126 MHz, DMSO-*d*$_6$) δ:** 171.0, 170.8, 167.8, 146.4, 146.2, 137.4, 137.0, 134.8, 130.3, 129.7, 129.2, 129.0, 128.4, 128.3, 128.1, 127.1, 126.6, 126.4, 123.4, 64.3, 54.0, 53.7, 53.1, 37.6, 36.8, 36.6, 33.9; **HRMS (ESI):** Calcd. for $C_{35}H_{37}N_4O_6^+$: *m/z* = 609.2713 [M+H]$^+$, found: *m/z*: 609.2699 [M+H]$^+$.

**Compound 4q**

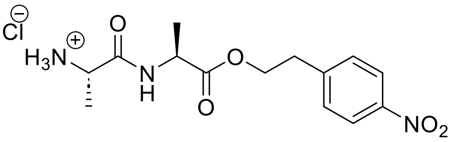


**Yield:** 98%; **IR:** ṽ (cm$^{-1}$) = 3357 (w), 3228 (w), 2848 (w), 2792 (w), 2715 (w), 2513 (w), 1733 (s), 1683 (s), 1599 (m), 1533 (m), 1510 (s), 1454 (w), 1391 (m), 1345 (s), 1319 (w), 1291 (w), 1219 (m), 1176 (m), 1136 (m), 1107 (w), 1068 (w), 1051 (w), 999 (w), 983 (w), 955 (w), 855 (s), 841 (m), 753 (m), 700 (s), 556 (m), 537 (m), 454 (w), 410 (w); **$^1$H NMR (500 MHz, DMSO-*d*$_6$) δ:** 8.92 (d, *J* = 6.9 Hz, 1H), 8.29 (s, 3H), 8.22 – 8.14 (m, 2H), 7.62 – 7.53 (m, 2H), 4.38 – 4.30 (m, 2H), 4.26 (p, *J* = 7.2 Hz, 1H), 3.83 (d, *J* = 8.5 Hz, 1H), 3.06 (t, *J* = 6.4 Hz, 2H), 1.32 (d, *J* = 7.0 Hz, 3H), 1.24 (d, *J* = 7.3 Hz, 3H); **$^{13}$C NMR (126 MHz, DMSO-*d*$_6$) δ:** 171.9, 169.4, 146.4, 146.3, 130.3, 123.4, 64.3, 47.8, 47.7, 34.0, 17.0, 16.7; **HRMS (ESI):** Calcd. for $C_{14}H_{20}N_3O_5^+$: *m/z* = 310.1403 [M+H]$^+$, found: *m/z*: 310.1397 [M+H]$^+$.

**Compound 4r**

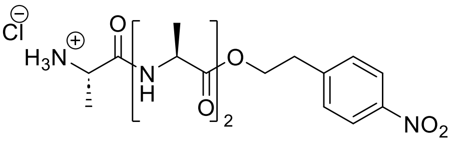


**Yield:** 44%;); **IR:** ṽ (cm$^{-1}$) = 3219 (w), 3057 (w), 2983 (w), 2926 (w), 1736 (m), 1654 (s), 1601 (w), 1542 (w), 1515 (vs), 1456 (m), 1388 (w), 1344 (s), 1259 (w), 1210 (m), 1167 (m), 1109 (m), 1048 (w), 999 (w), 959 (w), 855 (m), 747 (w), 698 (w), 537 (w); **$^1$H NMR (500 MHz, DMSO-*d*$_6$) δ:** 8.63 (d, *J* = 7.6 Hz, 1H), 8.43 (d, *J* = 6.9 Hz, 1H), 8.31 – 8.20 (m, 3H), 8.20 – 8.13 (m, 2H), 7.60 – 7.51 (m, 2H), 4.39 – 4.26 (m, 3H), 4.20 (p, *J* = 7.2 Hz, 1H), 3.84 (p, *J* = 5.9 Hz, 1H), 3.04 (t, *J* = 6.4 Hz, 2H), 1.33 (d, *J* = 6.9 Hz, 3H), 1.20 (dd, *J* = 7.2, 3.4 Hz, 6H); **$^{13}$C NMR (126 MHz, DMSO-*d*$_6$) δ:** 172.3, 171.7, 169.0, 146.5, 146.3, 130.3, 123.4, 64.1, 47.9, 47.6, 34.0, 18.2, 17.2, 16.8; **HRMS (ESI):** Calcd. for $C_{17}H_{25}N_4O_6^+$: *m/z* = 381.1774 [M+H]$^+$, found: *m/z*: 381.1769 [M+H]$^+$.

**Compound 4s**

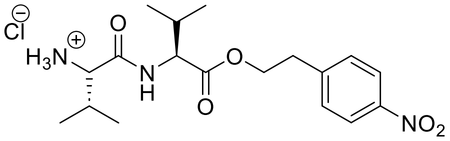


**Yield:** 96%; **$^1$H NMR (500 MHz, DMSO-$d_6$) δ:** 8.64 (d, *J* = 7.4 Hz, 1H), 8.25 (s, 3H), 8.20 – 8.13 (m, 2H), 7.60 – 7.54 (m, 2H), 4.44 – 4.31 (m, 2H), 4.14 (dd, *J* = 7.4, 5.7 Hz, 1H), 3.83 – 3.72 (m, 1H), 3.06 (t, *J* = 6.4 Hz, 2H), 2.11 – 1.94 (m, 2H), 0.91 (dd, *J* = 6.9, 2.6 Hz, 6H), 0.82 (t, *J* = 6.8 Hz, 6H); **HRMS (ESI):** Calcd. for $C_{18}H_{28}N_3O_5^+$: *m/z* = 366.2029 [M+H]$^+$, found: *m/z*: 366.2025 [M+H]$^+$.
Literature: ref.[4]

**Compound 4t**

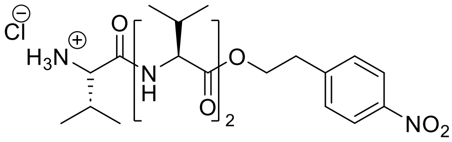


**Yield:** 96%; **$^1$H NMR (500 MHz, DMSO-$d_6$) δ:** 8.42 (d, *J* = 8.6 Hz, 1H), 8.26 (d, *J* = 8.0 Hz, 1H), 8.23 – 8.12 (m, 5H), 7.60 – 7.52 (m, 2H), 4.37 – 4.26 (m, 3H), 4.12 (dd, *J* = 8.0, 6.2 Hz, 1H), 3.72 (d, *J* = 5.6 Hz, 1H), 3.04 (t, *J* = 6.3 Hz, 2H), 1.98 (dddt, *J* = 49.1, 27.8, 13.8, 6.8 Hz, 3H), 0.89 (dd, *J* = 6.9, 3.6 Hz, 6H), 0.85 (dd, *J* = 6.8, 5.2 Hz, 6H), 0.77 (dd, *J* = 6.8, 4.1 Hz, 6H); **HRMS (ESI):** Calcd. for $C_{23}H_{37}N_4O_6^+$: *m/z* = 465.2713 [M+H]$^+$, found: *m/z*: 465.2701 [M+H]$^+$.
Literature: ref.[4]

#### 2.2 Synthesis of 2-thiouridine phosphoramidite

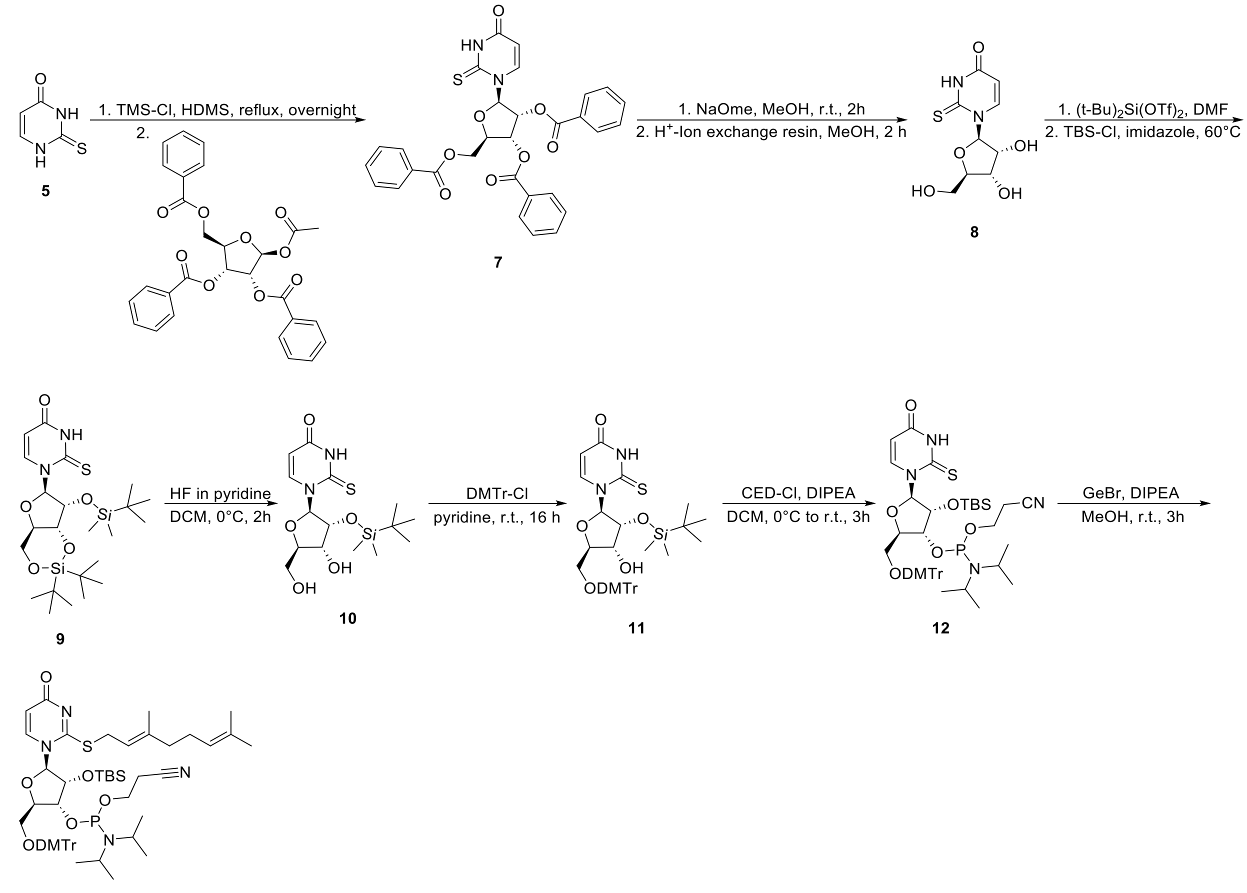


12b

**Scheme S2.** Synthesis of 2-thiouridine **12** and *S*-geranyl-2-thiouridine phosphoramidites.

#### Compound 7

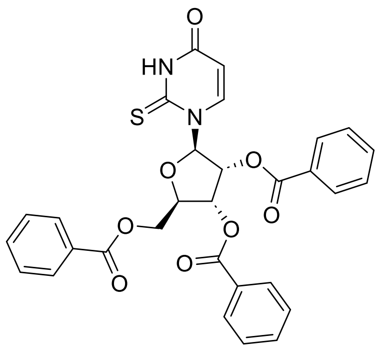

The reaction was performed similar to a procedure in the literature.[6]
**5** (1.0 equiv.) was suspended in HMDS and stirred at r.t.. TMS-Cl (2.0 equiv.) was added dropwise. The reaction was refluxed overnight and subsequently a solution of **6** (0.7 equiv.) in 1,2-dichloroethane was added, so that a 1:1 ratio of HMDS and 1,2-dichloroethane was formed. This mixture was cooled to 0°C and $SnCl_4$ (1.4 equiv. from 1M solution in DCM) was added dropwise. After addition was completed, the cooling was removed, and the reaction was stirred at r.t. overnight. The reaction was quenched by addition of aqueous saturated $NaHCO_3$ solution and stirred for 1h. Afterwards $H_2O$ was added. The organic Phase was separated, and the aqueous phase was extracted with DCM (three times). The combined organic layers were dried over $Na_2SO_4$ and the solvents were removed *in vacuo*. Purification

of the crude product via flash column chromatography (0-1.5% MeOH in DCM) yielded target compound **7**** as a white foam.

**Yield:** 55%; **$^1$H NMR (500 MHz, $CDCl_3$) δ:** 9.90 (s, 1H), 8.10 – 8.06 (m, 2H), 8.02 – 7.98 (m, 2H), 7.95 – 7.90 (m, 2H), 7.70 (d, *J* = 8.3 Hz, 1H), 7.68 – 7.62 (m, 1H), 7.61 – 7.54 (m, 2H), 7.52 (t, *J* = 7.8 Hz, 2H), 7.42 (t, *J* = 7.9 Hz, 2H), 7.38 (t, *J* = 7.8 Hz, 2H), 7.26 (d, *J* = 4.8 Hz, 1H), 5.84 – 5.76 (m, 3H), 4.88 (dd, *J* = 12.6, 2.6 Hz, 1H), 4.78 (dt, *J* = 5.3, 2.8 Hz, 1H), 4.69 (dd, *J* = 12.7, 3.0 Hz, 1H); **HRMS (ESI):** Calcd. for $C_{30}H_{23}N_2O_8S^-$: *m/z* = 571.1175 [M-H]$^-$, found: *m/z*: 571.1183 [M-H]$^-$.

Literature: ref.[6]

**Compound 8**

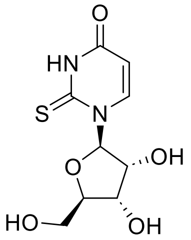

The reaction was performed similar to a procedure published before.[6]

Protected thiouridine **7** (1.0 equiv.) was dissolved in MeOH under $N_2$ atmosphere and the solution was cooled to 0°C. Afterwards $NaOCH_3$ (6.0 equiv.) was added in several portions and the reaction was stirred for 4 h at r.t.. Afterwards freshly protonated ion-exchange resin (Dowex 50WX8) was added until the solution showed neutral pH. Afterwards the resin was filtered, the residue washed with MeOH and the filtrate's solvents were removed *in vacuo*. The crude was suspended in $H_2O$ and the aqueous phase was washed with a small amount of EtOAc (two times). The aqueous phase was lyophilized to afford target compound **8** as a white powder.

**Yield:** 61%; **$^1$H NMR (500 MHz, $CDCl_3$) δ:** 8.44 (s, 1H), 8.17 (d, *J* = 8.1 Hz, 1H), 6.54 (d, *J* = 3.5 Hz, 1H), 5.94 (d, *J* = 8.1 Hz, 1H), 4.05 – 4.00 (m, 1H), 3.97 (t, *J* = 5.4 Hz, 1H), 3.89 (dt, *J* = 5.6, 2.6 Hz, 1H), 3.69 (dd, *J* = 12.4, 2.7 Hz, 1H), 3.60 (dd, *J* = 12.3, 2.7 Hz, 1H); **HRMS (ESI):** Calcd. for $C_9H_{11}N_2O_5S^+$: *m/z* = 259.0389 [M-H]$^-$, found: *m/z*: 259.0326 [M-H]$^-$.

Literature: ref.[6]

**Compound 9**

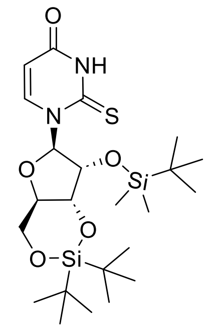

The reaction was performed similar to a procedure published before.

2-Thiouridine **8** (1.0 equiv.) was dissolved in dry DMF under $N_2$ atmosphere and the solution was cooled to 0°C. $(tBu)_2Si(OTf)_2$ (1.2 equiv.) was added dropwise under stirring and the reaction was left to stir for 1 h at 0°C. 1*H*-Imidazole (5.0 equiv.) was then added and the solution was allowed to warm to r.t. over 30 min. Afterwards TBS-Cl (1.2 equiv.) was added and the reaction mixture heated overnight at 60°C. The solution was concentrated *in vacuo*

and diluted with EtOAc. The organic phases were washed with brine (two times) and dried over $Na_2SO_4$. The solvent was removed *in vacuo* and the crude product was purified by flash column chromatography (7:3 Hex/EtOAc) to yield **9** as a white solid.
**Yield:** 80%; **$^1$H NMR (500 MHz, $CDCl_3$) δ:** 10.09 (s, 1H), 7.35 (d, *J* = 8.2 Hz, 1H), 6.46 (s, 1H), 6.03 (dd, *J* = 8.2, 1.8 Hz, 1H), 4.55 (dd, *J* = 9.4, 5.3 Hz, 1H), 4.34 (d, *J* = 4.4 Hz, 1H), 4.25 (td, *J* = 10.2, 5.3 Hz, 1H), 4.01 (dd, *J* = 10.6, 9.5 Hz, 1H), 3.78 (dd, *J* = 9.8, 4.5 Hz, 1H), 1.05 (d, *J* = 1.3 Hz, 9H), 1.02 (s, 9H), 0.95 (s, 9H), 0.23 (s, 3H), 0.16 (s, 3H); **HRMS (ESI):** Calcd. for $C_{23}H_{43}N_2O_5SSi_2^+$: *m/z* = 515.2431 [M+H]$^+$, found: *m/z*: 515.2429 [M+H]$^+$.
Literature: ref.[7]

**Compound 10**

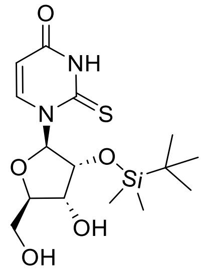


The reaction was performed similar to a procedure published before.[7]
**9** (1.0 equiv.) was dissolved in DCM/pyridine (9:1 v/v) under $N_2$-atmosphere in a plastic reaction vessel. The solution was cooled to 0°C and HF in pyridine (70%, 5.0 equiv.) was slowly added. The reaction was stirred at 0°C for 2h and quenched afterwards by addition of aqueous saturated $NaHCO_3$ solution. The mixture was extracted with DCM (three times), the combined organics dried over $Na_2SO_4$, and the solvents were removed *in vacuo*. Purification of the crude product via flash column chromatography (3% MeOH in DCM) yielded the target compound **10** as a white solid.
**Yield:** 75%; **$^1$H NMR (500 MHz, $CDCl_3$) δ:** 10.39 (s, 1H), 8.22 (d, *J* = 8.2 Hz, 1H), 6.67 (d, *J* = 3.3 Hz, 1H), 6.00 (d, *J* = 8.2 Hz, 1H), 4.31 (dd, *J* = 4.7, 3.3 Hz, 1H), 4.18 (q, *J* = 4.8 Hz, 1H), 4.14 (dt, *J* = 5.7, 2.0 Hz, 1H), 4.07 (dd, *J* = 12.0, 2.1 Hz, 1H), 3.93 (dd, *J* = 12.0, 1.9 Hz, 1H), 2.69 – 2.63 (m, 1H), 2.51 (s, 1H), 0.92 (s, 9H), 0.20 (s, 3H), 0.13 (s, 3H); **HRMS (ESI):** Calcd. for $C_{15}H_{25}N_2O_5SSi^-$: *m/z* = 373.1253 [M-H]$^-$, found: *m/z*: 373.1257 [M-H]$^-$.
Literature: ref.[7]

**Compound 11**

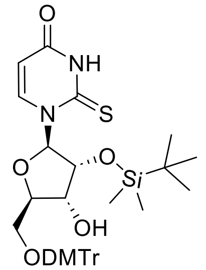


The reaction was performed similar to a procedure published before.[6,7]
Pre-dried compound **10** (1.0 equiv.) and pre-dried DMTr-Cl (1.5 equiv.) were dissolved in pyridine under $N_2$ atmosphere, and the solution was stirred at r.t. overnight. The solvent was removed *in vacuo* and the crude product was purified by flash column chromatography (7:3 Hex/EtOAc, addition of 0.1% pyridine) to afford **11** as a white foam.

**Yield:** 72%; **$^{1}$H NMR (500 MHz, $CD_2Cl_2$) δ:** 9.93 (s, 1H), 8.14 (d, *J* = 8.2 Hz, 1H), 7.42 – 7.36 (m, 2H), 7.36 – 7.19 (m, 8H), 6.90 – 6.83 (m, 4H), 6.69 (d, *J* = 3.6 Hz, 1H), 5.44 (d, *J* = 8.2 Hz, 1H), 4.44 (dd, *J* = 4.7, 3.7 Hz, 1H), 4.39 (t, *J* = 5.0 Hz, 1H), 4.15 (dt, *J* = 4.9, 2.2 Hz, 1H), 3.79 (d, *J* = 0.7 Hz, 6H), 3.50 (qd, *J* = 11.2, 2.3 Hz, 2H), 0.94 (s, 9H), 0.22 (s, 3H), 0.18 (s, 3H); **HRMS (ESI):** Calcd. for $C_{36}H_{43}N_2O_7SSi^-$: *m/z* = 675.2560 [M-H]$^-$, found: *m/z*: 675.2564 [M-H]$^-$.
Literature: ref.[6,7]

**Compound 12**

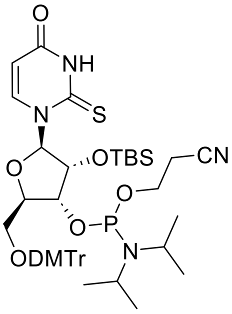


The reaction was performed similar to a procedure published before.[6]
**11** (1.0 equiv.) was pre-dried, set in Ar atmosphere, dissolved in dry DCM and cooled to 0°C. DIPEA (4.0 equiv.) was added dropwise to the solution, followed by addition of CED-Cl (2.5 equiv.). The reaction mixture was stirred at r.t. for 4.5 h. Afterwards the reaction was quenched by addition of saturated aqueous $NaHCO_3$ solution, the mixture was extracted with DCM (three times) and the combined organics were washed with brine and dried over $Na_2SO_4$. After removing of the solvent *in vacuo* flash column chromatography of the residue (EtOAc/*n*Hex 4:6 v/v, addition of 0.1% pyridine) yielded target compound **12** as a white foam.
**Yield:** 82%; **$^{31}$P NMR (202 MHz, $CD_2Cl_2$) δ:** 149.95, 149.87; **HRMS (ESI):** Calcd. for $C_{45}H_{61}N_4O_8PSSi^+$: *m/z* = 877.3795 [M+H]$^+$, found: *m/z*: 877.3804 [M+H]$^+$.
Literature: ref.[6,7]

**Compound 12b**

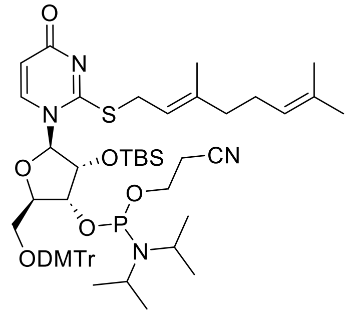


**12b** was synthesised according to literature.[8]
**12** (1.0 equiv.) was dissolved in anhydrous methanol to give a 10 mM concentration, followed by the addition of DIPEA (3.0 equiv.) and geranyl bromide (1.5 equiv.). The reaction was stirred at r.t. under inert atmosphere for 3 h. The reaction mixture was diluted with DCM and washed with brine 3 times. The crude was then purified by flash-column chromatography to afford the geranylated 2-thiouridine phosphoramidite **12b** as a white solid.
**Yield:** 90%; **$^{31}$P NMR (202 MHz, $CD_2Cl_2$) δ:** 151.47, 148.98; ***R*$_f$:** 0.5 (1:1 i-Hex/EtOAc).
Literature: ref.[8]

#### 2.3 Synthesis of 5-aminomethyluridine phosphoramidite

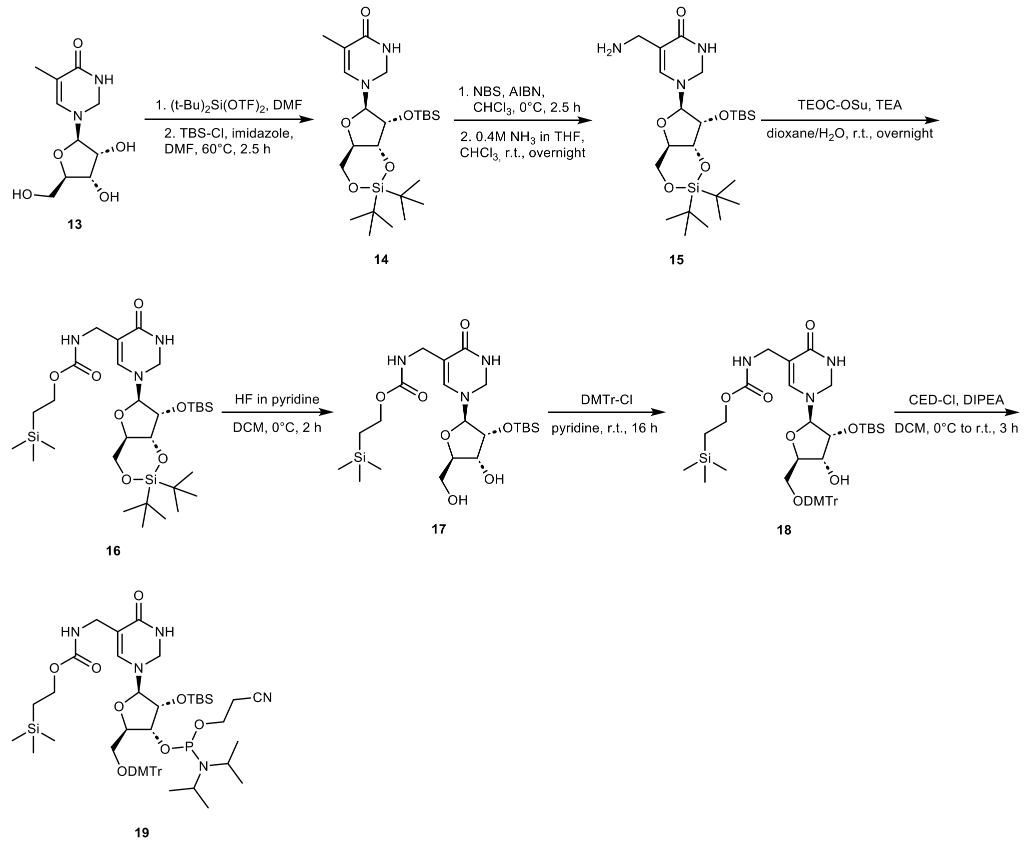

**Scheme S3.** Synthesis of TEOC-protected 5-aminomethyluridine phosphoramidite **19**.

##### Compound 14

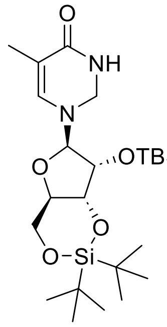

The hydroxy groups of 5-methyluridine **13** were protected according to a procedure published before.[9]

**13** (1.0 equiv.) was dissolved in dry DMF and the solution was cooled to 0°C. $(tBu)_2Si(OTf)_2$ (1.1 equiv.) was added dropwise and the reaction was left to stir for 1 h at 0°C. 1*H*-Imidazole (5.0 equiv.) and TBS chloride (1.2 equiv.) were added and the solution was stirred at 60°C overnight. The solvent was removed *in vacuo* and the remaining solid was dissolved in EtOAc. The organic phase was washed with $H_2O$ and brine and dried over $Na_2SO_4$. The solvent was removed *in vacuo* and the crude product was purified by flash column chromatography (1-2% MeOH in DCM) to yield **14** as a white solid.

**Yield:** 97%; **$^1$H NMR (500 MHz, CDCl$_3$) δ:** 8.78 (s, 1H), 7.01 (q, *J* = 1.2 Hz, 1H), 5.65 (s, 1H), 4.49 (dd, *J* = 9.2, 5.1 Hz, 1H), 4.29 (d, *J* = 4.8 Hz, 1H), 4.12 (td, *J* = 10.1, 5.1 Hz, 1H), 3.98 (dd, *J* = 10.5, 9.3 Hz, 1H), 3.92 (dd, *J* = 9.6, 4.9 Hz, 1H), 1.93 (d, *J* = 1.2 Hz, 3H), 1.06 (s, 9H), 1.02 (s, 9H), 0.92

(s, 9H), 0.16 (s, 3H), 0.13 (s, 3H); **HRMS (ESI):** Calcd. for $C_{24}H_{45}N_2O_6Si_2^+$: *m/z* = 513.2816 [M+H]$^+$, found: *m/z*: 518.2813 [M+H]$^+$.
Literature: ref.[9]

**Compound 15**

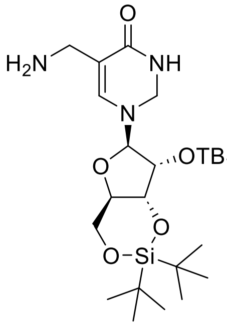

The amine group was inserted according to a procedure published before.[5]
**14** (1.0 equiv.) was heated in dry $CHCl_3$ under reflux and $N_2$ atmosphere. Freshly recrystallized NBS (1.2 equiv.) and AIBN (0.12 equiv.) were slowly added to the mixture and it was stirred for another 2 h. After completion of the bromination the solution was cooled to 0°C, $NH_3$ in dioxane (0.4 M, 5.0 equiv.) was added and the reaction mixture was stirred for 3 h at r.t.. The reaction was quenched by adding saturated aqueous $NaHCO_3$ solution and the aqueous phase was extracted with DCM (three times). The combined organics were dried over $Na_2SO_4$ and the solvents were removed *in vacuo*. The crude product was purified by flash column chromatography (0-6% MeOH in DCM) to yield **15** as a white solid.
**Yield:** 27%; **$^1$H NMR (500 MHz, CDCl$_3$) δ:** 7.33 (s, 1H), 5.69 (s, 1H), 4.48 (dd, *J* = 9.2, 5.0 Hz, 1H), 4.29 (d, *J* = 4.8 Hz, 1H), 4.13 (td, *J* = 10.0, 5.1 Hz, 1H), 4.02 (t, *J* = 9.9 Hz, 1H), 3.93 (dd, *J* = 9.6, 4.8 Hz, 1H), 3.62 (s, 2H), 1.05 (s, 9H), 1.02 (s, 9H), 0.92 (s, 9H), 0.16 (s, 3H), 0.13 (s, 3H); **HRMS (ESI):** Calcd. for $C_{24}H_{46}N_3O_6Si_2^+$: *m/z* = 528.2925 [M+H]$^+$, found: *m/z*: 528.29185 [M+H]$^+$.
Literature: ref.[5]

**Compound 16**

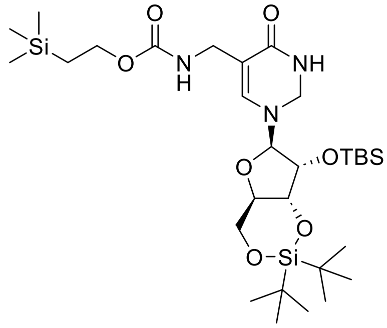

The amine group was TEOC-protected according to a procedure published before.[5]
**15** (1.0 equiv.) was dissolved in a mixture of dioxane and $H_2O$ (1:1 v/v). Teoc-Osu (1.1 equiv.) and subsequently TEA (1.5 equiv.) were added dropwise, and the reaction was stirred at r.t. overnight. $H_2O$ was added, and the aqueous phase was extracted with $Et_2O$ (three times). The combined organic phases were dried over $Na_2SO_4$, the solvents were evaporated, and the crude product was purified by flash column chromatography (0-2% methanol in DCM) yielding the target compound **16** as a white foam.
**Yield:** 74%; **$^1$H NMR (500 MHz, CDCl$_3$) δ:** 8.82 (s, 1H), 7.47 (s, 1H), 5.64 (s, 1H), 5.32 (t, *J* = 6.3 Hz, 1H), 4.50 (dd, *J* = 9.1, 5.0 Hz, 1H), 4.28 (d, *J* = 4.6 Hz, 1H), 4.22 – 4.03 (m, 4H), 3.98 (d, *J* =

6.4 Hz, 2H), 3.91 (dd, *J* = 9.6, 4.6 Hz, 1H), 1.06 (s, 9H), 1.02 (s, 9H), 0.99 – 0.92 (m, 11H), 0.18 (s, 3H), 0.14 (s, 3H), 0.03 (s, 9H); **HRMS (ESI):** Calcd. for $C_{30}H_{57}N_3NaO_8Si_3^+$: *m/z* = 694.3351 [M+Na]+, found: *m/z*: 694.3341 [M+Na]+.
Literature: ref.[5]

**Compound 17**

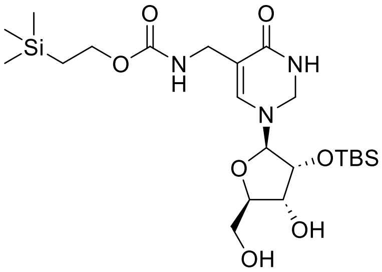


The reaction was performed similar to a procedure published before.[5]
**16** (1.0 equiv.) was dissolved in DCM/pyridine (9:1 v/v) under $N_2$-atmosphere in a plastic reaction vessel. The solution was cooled to 0°C and HF in pyridine (70%, 5.0 equiv.) was slowly added. The reaction was stirred at 0°C for 2h and quenched afterwards by addition of aqueous saturated $NaHCO_3$ solution. The mixture was extracted with DCM (three times), the combined organics dried over $Na_2SO_4$, and the solvents were removed *in vacuo*. Purification of the crude product via flash column chromatography (1-2% MeOH in DCM) yielded the target compound **17** as a white solid.
**Yield:** 90%; **$^1$H NMR (500 MHz, $CDCl_3$) δ:** 8.96 (s, 1H), 8.10 (s, 1H), 5.81 (d, *J* = 4.8 Hz, 1H), 5.46 (t, *J* = 6.4 Hz, 1H), 4.48 (t, *J* = 5.0 Hz, 1H), 4.26 (q, *J* = 4.4 Hz, 1H), 4.15 – 4.07 (m, 3H), 4.01 – 3.93 (m, 3H), 3.81 (d, *J* = 12.6 Hz, 1H), 3.66 (s, 1H), 2.71 (d, *J* = 4.6 Hz, 1H), 0.95 (dd, *J* = 10.1, 7.0 Hz, 2H), 0.90 (s, 9H), 0.09 (s, 6H), 0.02 (s, 9H); **HRMS (ESI):** Calcd. for $C_{22}H_{41}N_3NaO_8Si_2^+$: *m/z* = 554.2330 [M+Na]+, found: *m/z*: 554.2323 [M+Na]+.
Literature: ref.[5]

**Compound 18**

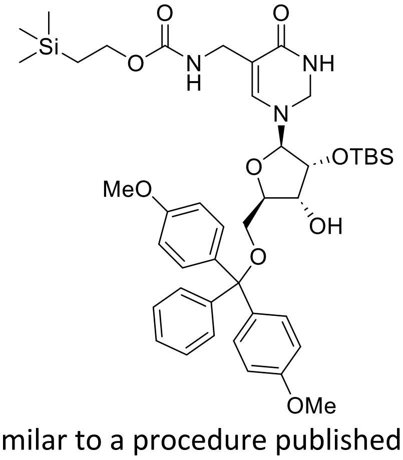


The reaction was performed similar to a procedure published before.[5]
**17** (1.0 equiv.) and DMTr-Cl (1.5 equiv.) were dissolved in pyridine and the solution was stirred at r.t. overnight. The solvent was removed *in vacuo* and the crude product was purified by flash column chromatography (0-1.5% MeOH in DCM, addition of 0.1% pyridine) to afford **18** as a white foam.
**Yield:** 99%; **$^1$H NMR (500 MHz, $CD_2Cl_2$) δ:** 8.67 (s, 1H), 7.70 (s, 1H), 7.49 – 7.42 (m, 2H), 7.38 – 7.27 (m, 6H), 7.26 – 7.22 (m, 1H), 6.90 – 6.83 (m, 4H), 5.91 (d, *J* = 4.8 Hz, 1H), 5.09 (t, *J* = 6.0

Hz, 1H), 4.42 (t, $J$ = 5.0 Hz, 1H), 4.22 (q, $J$ = 4.8 Hz, 1H), 4.18 – 4.01 (m, 3H), 3.78 (s, 6H), 3.56 (dd, $J$ = 14.7, 5.8 Hz, 1H), 3.43 (dd, $J$ = 15.1, 7.0 Hz, 1H), 3.39 (d, $J$ = 2.9 Hz, 2H), 2.62 (d, $J$ = 5.0 Hz, 1H), 0.92 (s, 11H), 0.14 (m, 6H), 0.02 (s, 9H).
; **HRMS (ESI):** Calcd. for $C_{43}H_{59}N_3NaO_{10}Si_2^+$: $m/z$ = 856.3637 $[M+Na]^+$, found: $m/z$: 856.3626 $[M+Na]^+$.
Literature: ref.[5]

**Compound 19**

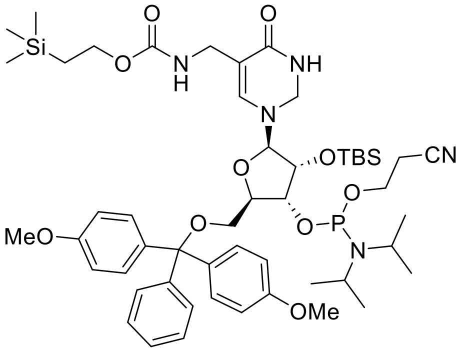


The reaction was performed similar to a procedure published before.[5]
**18** (1.0 equiv.) was pre-dried, set in Ar atmosphere, dissolved in dry DCM and cooled to 0°C. DIPEA (4.0 equiv.) was added dropwise to the solution, followed by addition of CED-Cl (2.5 equiv.). The reaction mixture was stirred at r.t. for 4.5 h. Afterwards the reaction was quenched by addition of saturated aqueous $NaHCO_3$ solution, the mixture was extracted with DCM (three times) and the combined organics were dried over $Na_2SO_4$. After removing of the solvent *in vacuo* flash column chromatography of the residue (EtOAc/*n*Hex 4:6 v/v, addition of 0.1% pyridine) yielded target compound **19** as a white foam.
**Yield:** 92%; **$^{31}$P NMR (202 MHz, $CD_2Cl_2$) δ:** 150.31, 149.20; **HRMS (ESI):** Calcd. for $C_{52}H_{77}N_5O_{11}PSi_2^+$: $m/z$ = 1034.4900 $[M+H]^+$, found: $m/z$: 1034.4887 $[M+H]^+$.
Literature: ref.[5]

#### 2.4 Synthesis of amino acid/peptide-modified $N^6$-carbamoyl adenosine phosphoramidites

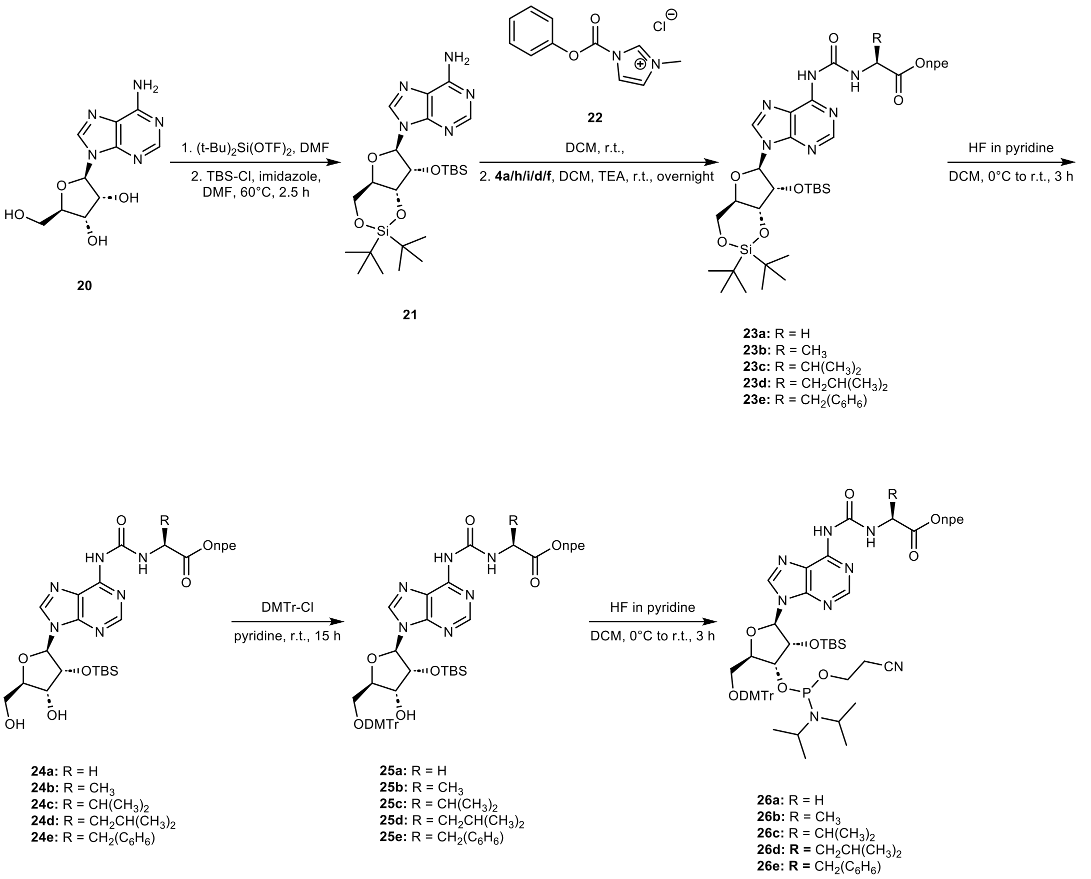


**Scheme S4.** Synthesis of TEOC-protected 5-aminomethyluridine phosphoramidite **26a-h**.

**Compound 21**

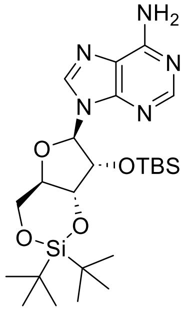

The reaction was performed similar to a procedure published before.[1,10]
Adenosine **20** (1.0 equiv.) was suspended in dry DMF under $N_2$ atmosphere and the solution was cooled to 0°C. $(tBu)_2Si(OTf)_2$ (1.2 equiv.) was added dropwise under stirring and the reaction was left to stir for 1 h at 0°C. 1*H*-Imidazole (5.0 equiv.) was then added and the solution was allowed to warm to r.t. over 30 min. Afterwards TBS-Cl (1.2 equiv.) was added and the reaction mixture was heated overnight at 60°C. Subsequently the reaction mixture was diluted with EtOAc, washed with brine (two times) and dried over $Na_2SO_4$. The solvent was removed *in vacuo* and the crude product was purified by flash column chromatography (2% MeOH in DCM) to yield **21** as a white solid.
**Yield:** 63%; **$^1$H NMR (500 MHz, $CDCl_3$) δ:** 8.26 (s, 1H), 7.82 (s, 1H), 6.93 (s, 2H), 5.89 (s, 1H), 4.59 (d, *J* = 4.7 Hz, 1H), 4.50 (dd, *J* = 9.5, 4.7 Hz, 1H), 4.44 (dd, *J* = 9.1, 5.1 Hz, 1H), 4.17 (td, *J* = 9.9, 5.0 Hz, 1H), 4.02 – 3.96 (m, 1H), 1.02 (s, 9H), 0.99 (s, 9H), 0.88 (s, 9H), 0.12 (s, 3H), 0.10

(s, 3H); **HRMS (ESI):** Calcd. for $C_{24}H_{44}N_5O_4Si_2^+$: *m/z* = 522.2932 [M+H]$^+$, found: *m/z*: 522.2929 [M+H]$^+$.
Literature: ref.[1,10]

**Compound 22**

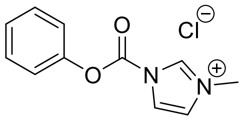

The reaction was performed according to published procedures.[1,11]
Phenyl chloroformate (1.0 equiv.) was dissolved in dry DCM under $N_2$-atmosphere and the solution was cooled to 0°C. Subsequently *N*-methylimidazole (1.0 equiv.) was added dropwise and the mixture was warmed to r.t. and stirred for another 2 h at r.t.. The formed precipitate was separated from solution by filtration and washed with DCM and dried *in vacuo* what yielded the activator **21** as a white crystalline solid.
**Yield:** 95%; **$^1$H NMR (500 MHz, $CDCl_3$) δ:** 9.43 (s, 1H), 7.49 – 7.26 (m, 6H), 7.08 (t, J = 1.7 Hz, 1H), 4.04 (s, 3H).
Literature: ref. [1,11]

**General procedure for the loading of the amino acids onto the adenosine derivative 23a-e**
The reaction was performed similar to a procedure published before.[1]
Silyl-protected adenosine **21** (1.0 equiv.) was dissolved in DCM under $N_2$ atmosphere. Subsequently the activator **22** (2.0 equiv.) was added to the solution and the resulting suspension was stirred overnight at r.t.. Afterwards the protected amino acids **4a/h/i/d/f/v** (2.0 equiv.) were added to the reaction mixture as a solution in DCM together with TEA (2.0 equiv.). The reaction was stirred overnight and quenched by addition of saturated aqueous $NaHCO_3$ solution. The aqueous Phase was separated and extracted with DCM (three times). The combined organic phases were dried over $Na_2SO_4$ and the solvent was removed *in vacuo*. Purification of the crude product by flash column chromatography yielded the target compounds **23a-e** as colorless solids.

**Compound 23a****

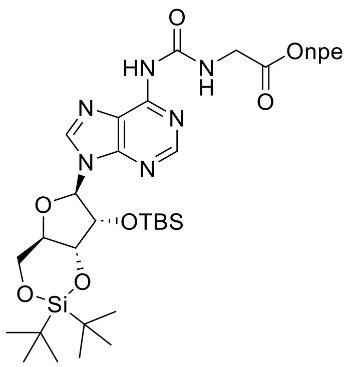


**Yield:** 69%; **$^1$H NMR (500 MHz, $CDCl_3$) δ:** 9.99 (t, *J* = 5.6 Hz, 1H), 8.67 (s, 1H), 8.50 (s, 1H), 8.20 (s, 1H), 8.12 – 8.05 (m, 2H), 7.41 – 7.35 (m, 2H), 5.99 (s, 1H), 4.59 (d, *J* = 4.6 Hz, 1H), 4.53 – 4.41 (m, 4H), 4.25 (td, *J* = 10.1, 5.1 Hz, 1H), 4.19 (d, *J* = 5.6 Hz, 2H), 4.07 (dd, *J* = 10.5, 9.2 Hz, 1H), 3.09 (t, *J* = 6.6 Hz, 2H), 1.08 (s, 9H), 1.05 (s, 9H), 0.94 (s, 9H), 0.17 (s, 3H), 0.15 (s, 3H); **HRMS (ESI):** Calcd. for $C_{35}H_{54}N_7O_9Si_2^+$: *m/z* = 772.3522 [M+H]$^+$, found: *m/z*: 772.3508 [M+H]$^+$.
Literature: ref.[1]

**Compound 23b**

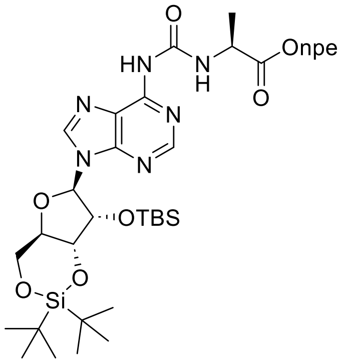

**Yield:** 70%; **$^1$H NMR (500 MHz, $CDCl_3$) δ:** 9.93 – 9.86 (m, 1H), 8.50 (s, 1H), 8.20 (s, 1H), 8.11 – 8.03 (m, 3H), 7.41 – 7.35 (m, 2H), 5.97 (s, 1H), 4.67 – 4.57 (m, 2H), 4.55 – 4.38 (m, 4H), 4.24 (td, *J* = 10.1, 5.1 Hz, 1H), 4.05 (dd, *J* = 10.5, 9.3 Hz, 1H), 3.09 (t, *J* = 6.5 Hz, 2H), 1.49 (d, *J* = 7.2 Hz, 3H), 1.08 (s, 9H), 1.05 (s, 9H), 0.94 (s, 9H), 0.17 (s, 3H), 0.15 (s, 3H); **HRMS (ESI):** Calcd. for $C_{36}H_{56}N_7O_9Si_2^+$: *m/z* = 786.3678 [M+H]$^+$, found: *m/z*: 786.3682 [M+H]$^+$.
Literature: ref.[5]

**Compound 23c**

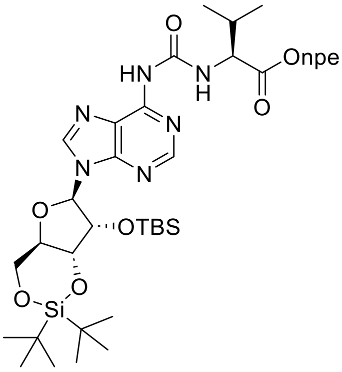

**Yield:** 78%; **$^1$H NMR (500 MHz, $CDCl_3$) δ:** 10.08 (d, *J* = 8.4 Hz, 1H), 8.79 (s, 1H), 8.50 (s, 1H), 8.27 (s, 1H), 8.11 – 8.02 (m, 2H), 7.44 – 7.34 (m, 2H), 5.99 (s, 1H), 4.59 (d, *J* = 4.6 Hz, 1H), 4.58 – 4.37 (m, 5H), 4.24 (td, *J* = 10.0, 5.1 Hz, 1H), 4.08 (dd, *J* = 10.5, 9.2 Hz, 1H), 3.09 (t, *J* = 6.5 Hz, 2H), 2.25 (pd, *J* = 6.9, 4.7 Hz, 1H), 1.08 (s, 9H), 1.04 (s, 9H), 1.00 (d, *J* = 6.9 Hz, 3H), 0.95 (d, *J* = 6.6 Hz, 12H), 0.18 (s, 3H), 0.15 (s, 3H); **HRMS (ESI):** Calcd. for $C_{38}H_{60}N_7O_9Si_2^+$: *m/z* = 814.3991 [M+H]$^+$, found: *m/z*: 814.3976 [M+H]$^+$.
Literature: ref.[1]

**Compound 23d**

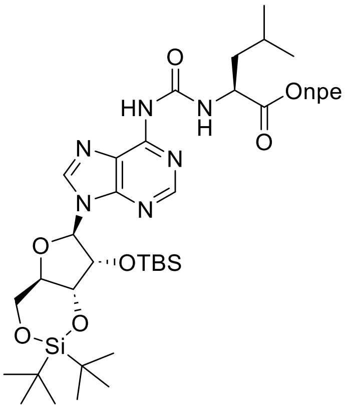

**Yield:** 82%; **$^1$H NMR (500 MHz, CDCl$_3$) δ:** 9.80 (d, *J* = 7.7 Hz, 1H), 8.49 (s, 1H), 8.13 (s, 1H), 8.09 – 8.03 (m, 3H), 7.41 – 7.35 (m, 2H), 5.96 (s, 1H), 4.64 – 4.56 (m, 2H), 4.50 (ddd, *J* = 11.7, 9.4, 4.9 Hz, 2H), 4.43 (ddt, *J* = 11.1, 6.6, 4.5 Hz, 2H), 4.24 (td, *J* = 10.1, 5.1 Hz, 1H), 4.05 (dd, *J* = 10.5, 9.3 Hz, 1H), 3.08 (t, *J* = 6.5 Hz, 2H), 1.78 – 1.63 (m, 3H), 1.08 (s, 9H), 1.05 (s, 9H), 0.94 (s, 15H), 0.18 (s, 3H), 0.16 (s, 3H); **HRMS (ESI):** Calcd. for $C_{39}H_{62}N_7O_9Si_2^+$: *m/z* = 828.4148 [M+H]$^+$, found: *m/z*: 828.4149 [M+H]$^+$.
Literature: ref.[5]

**Compound 23e**

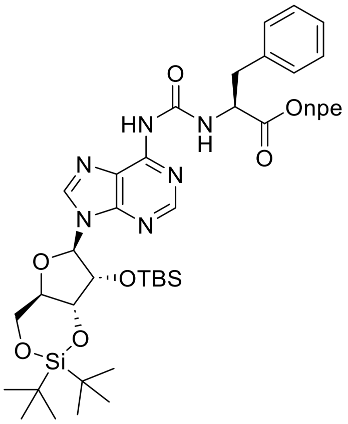

**Yield:** 71%; **$^1$H NMR (500 MHz, CDCl$_3$) δ:** 9.90 (d, *J* = 7.7 Hz, 1H), 8.33 (s, 1H), 8.17 (s, 1H), 8.04 (s, 1H), 8.05 – 7.98 (m, 2H), 7.35 – 7.14 (m, 7H), 5.95 (s, 1H), 4.88 (dt, *J* = 7.5, 6.2 Hz, 1H), 4.62 (d, *J* = 4.6 Hz, 1H), 4.50 (ddd, *J* = 9.1, 4.9, 2.9 Hz, 2H), 4.39 (qt, *J* = 10.9, 6.5 Hz, 2H), 4.24 (td, *J* = 10.1, 5.2 Hz, 1H), 4.06 (dd, *J* = 10.5, 9.3 Hz, 1H), 3.25 – 3.13 (m, 2H), 3.02 – 2.95 (m, 2H), 1.09 (s, 9H), 1.05 (s, 9H), 0.94 (s, 9H), 0.18 (s, 3H), 0.16 (s, 3H); **HRMS (ESI):** Calcd. for $C_{42}H_{60}N_7O_9Si_2^+$: *m/z* = 862.3991 [M+H]$^+$, found: *m/z*: 862.3996 [M+H]$^+$.
Literature: ref.[1]

**General procedure for the removing of the silyl-clamp of 23a-e yielding 24a-e**
The reaction was performed similar to a procedure published before.[1]
Silyl-protected and modified adenosine derivatives **23a-e** (1.0 equiv.) were dissolved in DCM/pyridine (9:1 v/v) under $N_2$-atmosphere in a plastic reaction vessel. The solution was cooled to 0°C and HF in pyridine (70%, 5.0 equiv.) was slowly added. The reaction was stirred at 0°C for 2h and quenched afterwards by addition of aqueous saturated $NaHCO_3$ solution. The mixture was extracted with DCM (three times), the combined organics dried over $Na_2SO_4$,

and the solvents were removed *in vacuo*. Purification of the crude product via flash column chromatography yielded the target compounds **24a-e** as a white solid.

**Compound 24a**

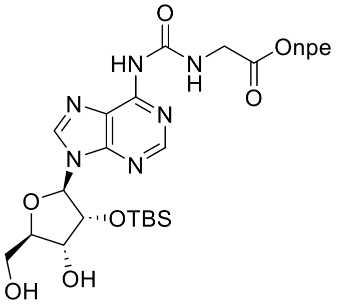

**Yield:** 87%; **$^1$H NMR (500 MHz, CDCl$_3$) δ:** 9.88 (t, *J* = 5.6 Hz, 1H), 8.68 (d, *J* = 3.0 Hz, 1H), 8.54 (s, 1H), 8.18 – 8.11 (m, 3H), 7.44 – 7.36 (m, 2H), 5.88 – 5.81 (m, 2H), 5.08 (dd, *J* = 7.3, 4.8 Hz, 1H), 4.45 (td, *J* = 6.7, 1.5 Hz, 2H), 4.40 – 4.34 (m, 2H), 4.27 – 4.13 (m, 2H), 3.97 (dt, *J* = 12.9, 2.0 Hz, 1H), 3.77 (ddd, *J* = 13.2, 11.8, 1.6 Hz, 1H), 3.10 (t, *J* = 6.7 Hz, 2H), 2.89 (s, 1H), 0.79 (s, 9H), -0.19 (s, 3H), -0.38 (s, 3H); **HRMS (ESI):** Calcd. for $C_{27}H_{38}N_7O_9Si^+$: *m/z* = 632.2500 [M+H]$^+$, found: *m/z*: 632.2492 [M+H]$^+$.
Literature: ref.[1]

**Compound 24b**

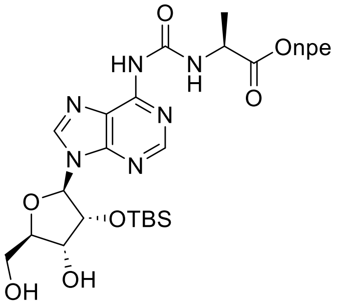

**Yield:** 74%; ***R*$_f$:** 0.38 (95: 5 DCM/MeOH); **IR:** ṽ (cm$^{-1}$) = 2858 (w), 1743 (m), 1698 (s), 1613 (s), 1518 (m), 1470 (m), 1345 (s), 1253 (m), 1212 (w) , 1149 (w), 1064 (w), 837 (m), 780 (s), 696 (m), 494 (s); **$^1$H NMR (500 MHz, CDCl$_3$) δ**: 9.83 (d, *J* = 7.1 Hz, 1H), 8.56 (s, 1H), 8.35 (s, 1H), 8.21 – 8.12 (m, 2H), 8.08 (s, 1H), 7.46 – 7.34 (m, 2H), 5.88 – 5.79 (m, 2H), 5.09 (dd, *J* = 7.3, 4.8 Hz, 1H), 4.62 (p, J = 7.2 Hz, 1H), 4.45 (td, *J* = 6.7, 1.0 Hz, 2H), 4.41 – 4.34 (m, 2H), 4.02 – 3.90 (m, 1H), 3.82 – 3.72 (m, 1H), 3.11 (t, *J* = 6.7 Hz, 2H), 2.85 (s, 1H), 1.51 (d, *J* = 7.2 Hz, 3H), 0.80 (s, 9H), -0.18 (s, 3H), -0.38 (s, 3H); **$^{13}$C NMR (126 MHz, CDCl$_3$) δ**: 173.0, 153.0, 151.1, 150.9, 149.4, 147.1, 145.5, 143.2, 129.9, 123.9, 122.2, 91.4, 87.7, 74.7, 72.9, 64.9, 63.4, 49.3, 35.0, 25.6, 18.5, 18.0, -5.2, -5.3; **HRMS (ESI):** Calcd. for $C_{28}H_{39}N_7O_9SiNa^+$: *m/z* = 668.2476 [M+Na]$^+$, found: *m/z*: 668.2463 [M+Na]$^+$.

**Compound 24c**

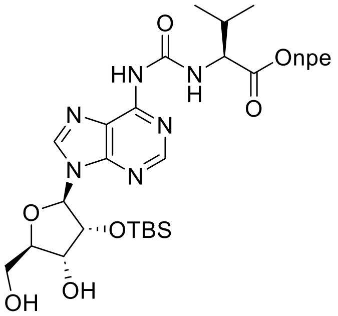

**Yield:** 93%; **$^1$H NMR (500 MHz, $CDCl_3$) δ:** 9.91 (d, J = 8.2 Hz, 1H), 8.57 (s, 1H), 8.44 (s, 1H), 8.18 – 8.11 (m, 2H), 8.09 (s, 1H), 7.44 – 7.37 (m, 2H), 5.89 (d, J = 11.8 Hz, 1H), 5.83 (d, J = 7.3 Hz, 1H), 5.10 (dd, J = 7.3, 4.7 Hz, 1H), 4.53 (dd, J = 8.2, 4.7 Hz, 1H), 4.44 (t, J = 6.6 Hz, 2H), 4.37 (d, J = 5.1 Hz, 2H), 3.95 (d, J = 12.9 Hz, 1H), 3.76 (t, J = 12.2 Hz, 1H), 3.16 – 3.06 (m, 2H), 2.89 (s, 1H), 2.24 (pd, J = 6.9, 4.7 Hz, 1H), 1.00 (d, J = 6.8 Hz, 3H), 0.94 (d, J = 6.9 Hz, 3H), 0.80 (s, 9H), -0.18 (s, 3H), -0.38 (s, 3H); **HRMS (ESI):** Calcd. for $C_{30}H_{44}N_7O_9Si^+$: *m/z* = 674.2970 [M+H]$^+$, found: *m/z*: 674.2974 [M+H]$^+$.
Literature: ref.[1]

**Compound 24d**

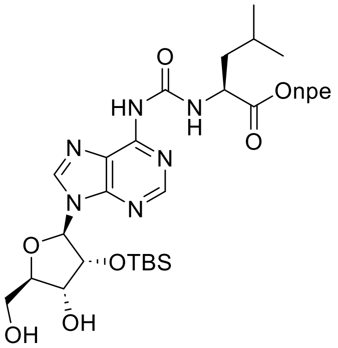

**Yield:** 82%; ***R*$_f$:** 0.56 (95: 5 DCM/MeOH); **IR:** $\tilde{\nu}$ (cm$^{-1}$) = 2955 (w), 2930 (w), 2858 (w), 1742 (w), 1696 (s), 1614 (m), 1589 (m), 1519 (s), 1470 (m), 1345 (s), 1253 (m), 1148 (m), 1089 (m), 1052 (m), 1005 (w), 956 (w), 915 (w), 856 (m), 836 (s), 780 (s), 746 (w), 697 (w), 647 (m), 627 (w), ; **$^1$H NMR (500 MHz, $CDCl_3$) δ:** 9.75 (d, *J* = 7.5 Hz, 1H), 8.55 (s, 1H), 8.31 (s, 1H), 8.18 – 8.11 (m, 2H), 8.05 (s, 1H), 7.45 – 7.37 (m, 2H), 5.88 (dd, *J* = 11.9, 2.1 Hz, 1H), 5.83 (d, *J* = 7.3 Hz, 1H), 5.09 (dd, *J* = 7.3, 4.7 Hz, 1H), 4.60 (td, *J* = 7.9, 5.8 Hz, 1H), 4.49 – 4.37 (m, 2H), 4.39 – 4.34 (m, 2H), 3.95 (dt, *J* = 12.8, 2.0 Hz, 1H), 3.82 – 3.72 (m, 1H), 3.18 – 3.04 (m, 2H), 2.86 (s, 1H), 1.77 – 1.61 (m, 3H), 0.99 – 0.90 (m, 6H), 0.80 (s, 9H), -0.17 (s, 3H), -0.38 (s, 3H); **$^{13}$C NMR (126 MHz, $CDCl_3$) δ:** 173.1, 153.3, 151.0, 151.0, 150.0, 149.3, 147.1, 145.5, 143.1, 130.0, 123.9, 122.1, 91.4, 87.7, 74.7, 72.9, 64.8, 63.4, 52.2, 41.3, 35.0, 26.8, 25.6, 25.2, 23.0, 22.1, 18.0, -5.2, -5.3; **HRMS (ESI):** Calcd. for $C_{31}H_{46}N_7O_9Si^+$: *m/z* = 688.3126 [M+H]$^+$, found: *m/z*: 688.3113 [M+H]$^+$.

**Compound 24e**

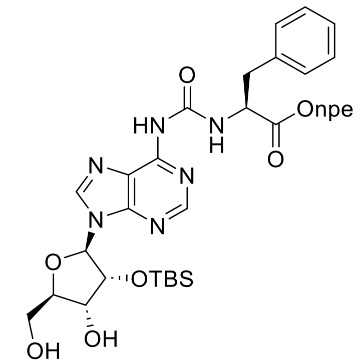


**Yield:** 80%; $^{1}$**H NMR (500 MHz, CDCl$_3$) δ:** 9.80 (d, *J* = 7.5 Hz, 1H), 8.37 (s, 1H), 8.29 (s, 1H), 8.15 – 8.08 (m, 2H), 8.02 (s, 1H), 7.37 – 7.31 (m, 2H), 7.28 (dddd, *J* = 9.2, 7.2, 5.8, 1.7 Hz, 3H), 7.18 – 7.12 (m, 2H), 5.87 (dd, *J* = 12.0, 2.1 Hz, 1H), 5.81 (d, *J* = 7.3 Hz, 1H), 5.07 (dd, *J* = 7.3, 4.8 Hz, 1H), 4.89 (dt, *J* = 7.4, 6.2 Hz, 1H), 4.48 – 4.33 (m, 4H), 3.97 (dt, *J* = 12.8, 1.9 Hz, 1H), 3.82 – 3.73 (m, 1H), 3.18 (d, *J* = 6.2 Hz, 2H), 3.04 (td, *J* = 6.6, 2.4 Hz, 2H), 2.84 (s, 1H), 0.79 (s, 9H), -0.19 (s, 3H), -0.40 (s, 3H); **HRMS (ESI):** Calcd. for $C_{34}H_{44}N_7O_9Si^+$: *m/z* = 722.2970 [M+H]$^+$, found: *m/z*: 722.2971 [M+H]$^+$.
Literature: ref.[1]

**General procedure for the DMTr-protection of the 5'-OH group of compounds 24a-e yielding 25a-e**

The reaction was performed similar to a procedure published before.[1]
3'-5'-deprotected and modified adenosine derivatives **24a-e** (1.0 equiv.) were dissolved in pyridine under $N_2$-atmosphere. Dried DMTr-Cl (1.5 equiv.) was dissolved in pyridine and added to the other solution. The mixture was stirred at r.t. overnight. Afterwards the solvent was removed *in* vacuo and the crude product was purified via flash column chromatography to afford the target compounds **25a-e** as a white solid.

**Compound 25a**

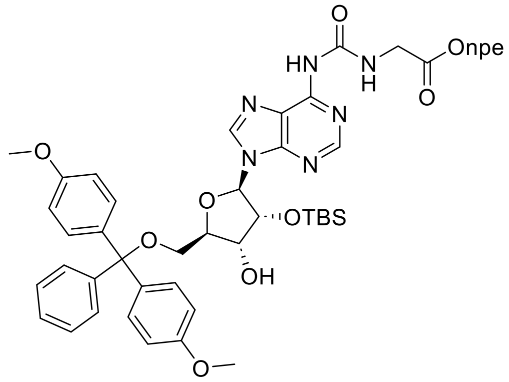


**Yield:** 90%; $^{1}$**H NMR (500 MHz, CD$_2$Cl$_2$) δ:** 9.83 (t, *J* = 5.7 Hz, 1H), 8.44 (s, 1H), 8.19 (s, 1H), 8.14 (s, 1H), 8.14 – 8.08 (m, 2H), 7.48 – 7.39 (m, 4H), 7.37 – 7.30 (m, 4H), 7.32 – 7.24 (m, 2H), 7.25 – 7.20 (m, 1H), 6.85 – 6.78 (m, 4H), 6.05 (d, *J* = 4.9 Hz, 1H), 4.97 (t, *J* = 5.0 Hz, 1H), 4.43 (t, *J* = 6.6 Hz, 2H), 4.38 (q, *J* = 4.6 Hz, 1H), 4.24 (q, *J* = 3.8 Hz, 1H), 4.14 (d, *J* = 5.7 Hz, 2H), 3.77 (s, 6H), 3.51 – 3.35 (m, 2H), 3.09 (t, *J* = 6.6 Hz, 2H), 2.67 (d, *J* = 4.8 Hz, 1H), 0.85 (s, 9H), 0.01 (s, 3H), -0.11 (s, 3H); **HRMS (ESI):** Calcd. for $C_{48}H_{56}N_7O_{11}Si^+$: *m/z* = 934.3807 [M+H]$^+$, found: *m/z*: 934.3786 [M+H]$^+$.
Literature: ref.[1]

**Compound 25b**

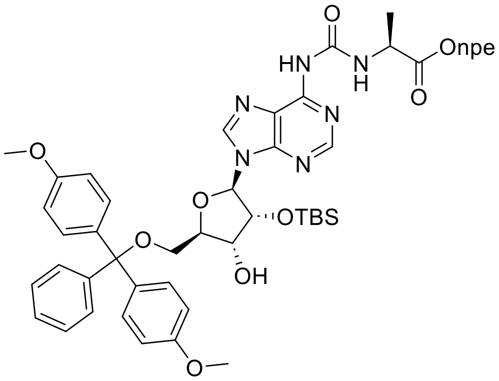


**Yield:** 88%; ***R*$_f$:** 0.45 (95:5 DCM/MeOH); **IR:** $\tilde{\nu}$ (cm$^{-1}$) = = 2856 (w), 1743 (w), 1700 (m), 1608 (m), 1586 (m), 1508 (s), 1468 (m), 1345 (s), 1301 (m), 1248 (s), 1174 (s), 1148 (m), 1112 (m), 1064 (m), 1033 (s), 912 (w), 781 (s), 748 (m), 727 (w), 701 (s), 646 (w) 582 (s), 478 (w), 449 (w); **$^1$H NMR (500 MHz, $CD_2Cl_2$) δ:** 9.79 (d, J = 7.2 Hz, 1H), 8.44 (s, 1H), 8.17 (s, 1H), 8.13 – 8.06 (m, 2H), 7.91 (s, 1H), 7.49 – 7.39 (m, 4H), 7.37 – 7.31 (m, 4H), 7.32 – 7.24 (m, 2H), 7.26 – 7.19 (m, 1H), 6.86 – 6.78 (m, 4H), 6.04 (d, *J* = 5.0 Hz, 1H), 5.01 (t, *J* = 5.0 Hz, 1H), 4.55 (p, *J* = 7.2 Hz, 1H), 4.49 – 4.34 (m, 3H), 4.24 (q, *J* = 3.8 Hz, 1H), 3.77 (s, 6H), 3.48 (dd, *J* = 10.6, 3.2 Hz, 1H), 3.39 (dd, *J* = 10.7, 4.3 Hz, 1H), 3.09 (t, *J* = 6.5 Hz, 2H), 2.66 (d, *J* = 4.7 Hz, 1H), 1.47 (d, *J* = 7.2 Hz, 3H), 0.85 (s, 9H), 0.01 (s, 3H), -0.12 (s, 3H); **$^{13}$C NMR (126 MHz, $CD_2Cl_2$) δ:** 173.1, 159.1, 153.4, 151.5, 150.7, 150.5, 150.1, 147.3, 146.2, 145.3, 142.2, 136.3, 136.1, 130.5, 130.5, 130.3, 128.5, 128.3, 127.3, 123.9, 121.3, 113.5, 89.2, 87.0, 84.6, 75.9, 71.8, 65.0, 63.8, 55.6, 49.6, 35.2, 25.8, 18.4, 18.2, -4.8, -5.0.; **HRMS (ESI):** Calcd. for $C_{49}H_{57}N_7O_{11}SiNa^+$: *m/z* = 970.3783 [M+Na]$^+$, found: *m/z*: 970.3773 [M+Na]$^+$.

**Compound 25c**

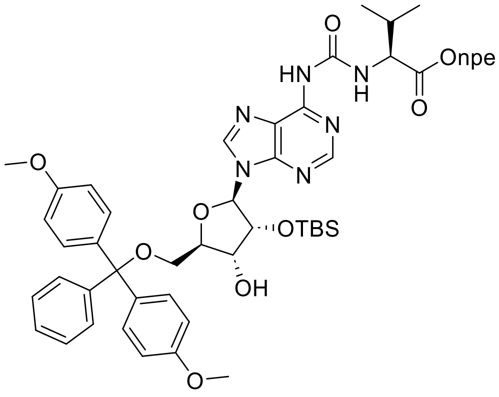


**Yield:** 70%; **$^1$H NMR (500 MHz, $CD_2Cl_2$) δ:** 9.86 (d, *J* = 8.2 Hz, 1H), 8.44 (s, 1H), 8.16 (s, 1H), 8.12 – 8.05 (m, 2H), 7.95 (s, 1H), 7.48 – 7.38 (m, 4H), 7.38 – 7.31 (m, 4H), 7.32 – 7.23 (m, 3H), 7.26 – 7.19 (m, 1H), 6.85 – 6.78 (m, 4H), 6.04 (d, *J* = 4.9 Hz, 1H), 5.01 (t, *J* = 5.0 Hz, 1H), 4.49 – 4.34 (m, 4H), 4.24 (td, *J* = 4.1, 3.1 Hz, 1H), 3.77 (d, *J* = 0.6 Hz, 6H), 3.48 (dd, *J* = 10.7, 3.2 Hz, 1H), 3.40 (dd, *J* = 10.7, 4.3 Hz, 1H), 3.09 (t, *J* = 6.5 Hz, 2H), 2.65 (d, *J* = 4.8 Hz, 1H), 2.23 (pd, *J* = 7.0, 4.9 Hz, 1H), 1.00 (d, *J* = 6.8 Hz, 3H), 0.95 (d, *J* = 6.9 Hz, 3H), 0.86 (s, 9H), 0.02 (s, 3H), -0.10 (s, 3H); **HRMS (ESI):** Calcd. for $C_{51}H_{62}N_7O_{11}Si^+$: *m/z* = 976.4277 [M+H]$^+$, found: *m/z*: 976.4287 [M+H]$^+$.
Literature: ref.[1]

**Compound 25d**

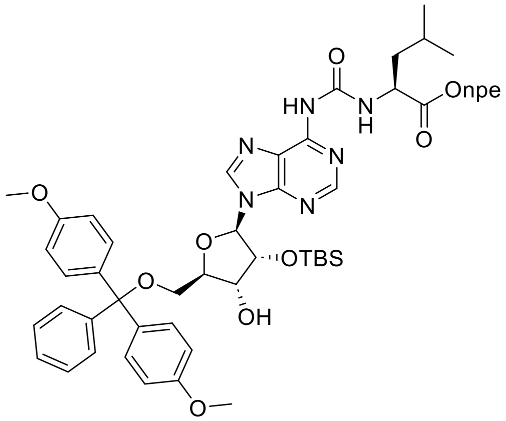


**Yield:** 75%; ***R*f:** 0.52 (8:2 DCM/EtOAc); **IR:** $\tilde{\nu}$ ($cm^{-1}$) = 2955 (w), 2930 (w), 2857 (w), 1742 (w), 1700 (s), 1608 (m), 1587 (w), 1508 (s), 1470 (m), 1445 (w), 1345 (s), 1301 (w), 1248 (s), 1176 (m), 1150 (w), 1112 (w), 1065 (w), 1033 (m), 913 (w), 834 (s), 782 (m), 750 (w), 700 (m), 647 (w), 582 (m); **$^{1}$H NMR (500 MHz, $CD_2Cl_2$) δ:** 9.76 (d, *J* = 7.5 Hz, 1H), 8.43 (s, 1H), 8.20 (s, 1H), 8.11 (s, 1H), 8.11 – 8.05 (m, 2H), 7.50 – 7.38 (m, 4H), 7.35 (dq, *J* = 8.8, 1.9 Hz, 4H), 7.32 – 7.23 (m, 2H), 7.26 – 7.18 (m, 1H), 6.85 – 6.78 (m, 4H), 6.04 (d, *J* = 4.9 Hz, 1H), 5.03 (t, *J* = 5.0 Hz, 1H), 4.54 (ddd, *J* = 8.6, 7.5, 5.8 Hz, 1H), 4.48 – 4.34 (m, 3H), 4.25 (q, *J* = 3.8 Hz, 1H), 3.77 (d, *J* = 0.8 Hz, 6H), 3.48 (dd, *J* = 10.7, 3.2 Hz, 1H), 3.40 (dd, *J* = 10.7, 4.3 Hz, 1H), 3.09 (t, *J* = 6.5 Hz, 2H), 2.71 (d, *J* = 4.8 Hz, 1H), 1.79 – 1.60 (m, 3H), 0.95 (s, 3H), 0.93 (s, 3H), 0.85 (s, 9H), 0.01 (s, 3H), -0.10 (s, 3H); **$^{13}$C NMR (126 MHz, $CD_2Cl_2$) δ:** 173.1, 159.1, 153.7, 151.4, 150.7, 150.5, 150.2, 147.2, 146.3, 145.2, 142.3, 136.2, 136.0, 136.0, 130.5, 130.3, 128.5, 128.3, 127.2, 123.9, 113.5, 89.2, 87.0, 84.5, 75.8, 71.7, 64.9, 63.8, 55.6, 52.4, 41.4, 35.2, 29.5, 25.7, 25.4, 23.0, 22.0, 18.2, -4.8, -5.0; **HRMS (ESI):** Calcd. for $C_{52}H_{64}N_7O_{11}Si^+$: *m/z* = 990.4433 $[M+H]^+$, found: *m/z*: 990.4417 $[M+H]^+$.

**Compound 25e**

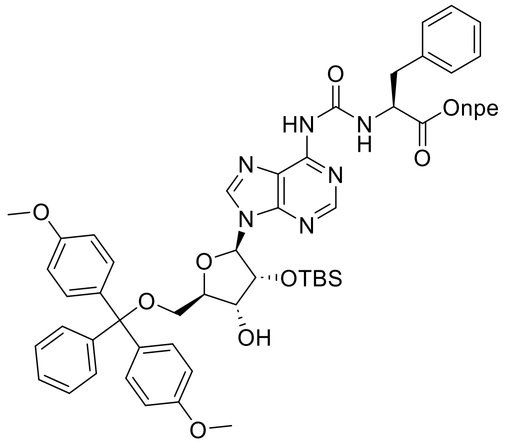


**Yield:** 85%; **$^{1}$H NMR (500 MHz, $CD_2Cl_2$) δ:** 9.80 (d, *J* = 7.6 Hz, 1H), 8.29 (s, 1H), 8.15 (s, 1H), 8.09 – 8.02 (m, 2H), 7.87 (s, 1H), 7.49 – 7.42 (m, 2H), 7.38 – 7.31 (m, 6H), 7.33 – 7.19 (m, 6H), 7.20 – 7.14 (m, 2H), 6.88 – 6.76 (m, 4H), 6.02 (d, *J* = 5.0 Hz, 1H), 5.00 (t, *J* = 5.0 Hz, 1H), 4.81 (dt, *J* = 7.4, 6.2 Hz, 1H), 4.44 – 4.32 (m, 3H), 4.24 (q, *J* = 3.8 Hz, 1H), 3.77 (s, 6H), 3.48 (dd, *J* = 10.7, 3.2 Hz, 1H), 3.40 (dd, *J* = 10.7, 4.3 Hz, 1H), 3.19 – 3.14 (m, 2H), 3.03 (t, *J* = 6.5 Hz, 2H), 2.64 (d, *J* = 4.8 Hz, 1H), 0.85 (s, 9H), 0.01 (s, 3H), -0.11 (s, 3H); **HRMS (ESI):** Calcd. for $C_{55}H_{62}N_7O_{11}Si^+$: *m/z* = 1024.4277 $[M+H]^+$, found: *m/z*: 1024.4291 $[M+H]^+$.
Literature: ref.[1]

**General procedure for the final phosphoramidite synthesis of 26a-e**

The reaction was performed similar to a procedure published before.[1]

**25a-e** (1.0 equiv.) were dissolved in dry DCM under Ar-atmosphere. The solution was cooled to 0°C and DIPEA (4.0 equiv.) were added dropwise followed by the addition of CED-Cl (2.6 equiv.). The reaction progress was monitored by TLC. After reaction completion (average around 3h) the reaction was quenched by addition of aq. sat. solution of $NaHCO_3$. The aqueous phase was extracted with DCM (three times). The combined organic phases were dried over $Na_2SO_4$ and the solvents were removed *in vacuo*. The crude product was purified by flash column chromatography using a mixture of *n*-hexane and EtOAc with addition of 0.1% pyridine to afford the final phosphoramidites **26a-e** as a white solid and a mixture of two diastereoisomers.

**Compound 26a**

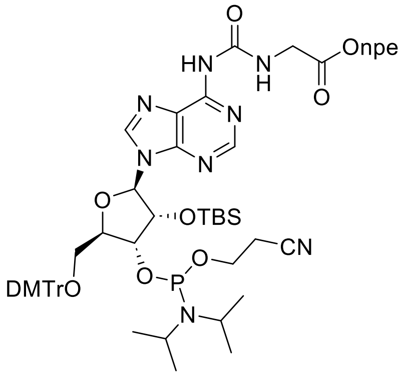


**Yield:** 82%; **$^{31}$P NMR (202 MHz, $CD_2Cl_2$) δ:** 150.66, 149.18; **HRMS (ESI):** Calcd. for $C_{57}H_{73}N_9O_{12}PSi^+$: *m/z* = 1134.4886 $[M+H]^+$, found: *m/z*: 1134.4879 $[M+H]^+$.
Literature: ref.[1]

**Compound 26b**

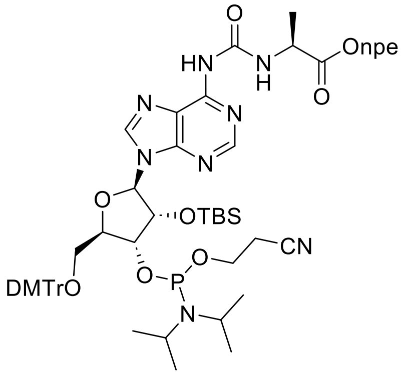


**Yield:** 74%; ***R*$_f$:** 0.27 (97: 3 DCM/MeOH); **$^{31}$P NMR (202 MHz, $CD_2Cl_2$) δ:** 150.77, 149.17; **HRMS (ESI):** Calcd. for $C_{58}H_{74}N_9O_{12}PSiNa^+$: *m/z* = 1170.4862 $[M+Na]^+$, found: *m/z*: 1170.4836 $[M+Na]^+$.

**Compound 26c**

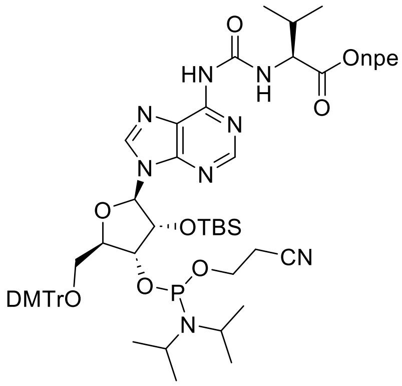


**Yield:** 87%; **$^{31}$P NMR (202 MHz, $CD_2Cl_2$) δ:** 150.76, 149.18; **HRMS (ESI):** Calcd. for $C_{60}H_{79}N_9O_{12}PSi^+$: *m/z* = 1176.5355 [M+H]$^+$, found: *m/z*: 1176.5359 [M+H]$^+$.
Literature: ref.[1]

**Compound 26d**

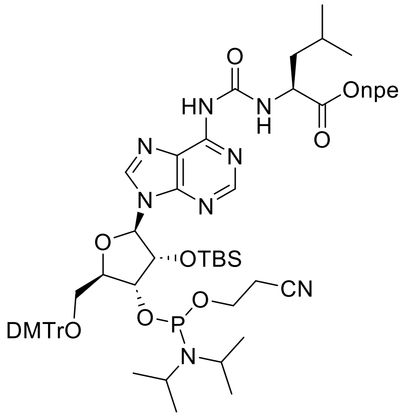


**Yield:** 85%; **$^{31}$P NMR (202 MHz, $CD_2Cl_2$) δ:** 150.70, 149.15; **HRMS (ESI):** Calcd. for $C_{61}H_{81}N_9O_{12}PSi^+$: *m/z* = 1190.5512 [M+H]$^+$, found: *m/z*: 1190.5508 [M+H]$^+$.

**Compound 26e**

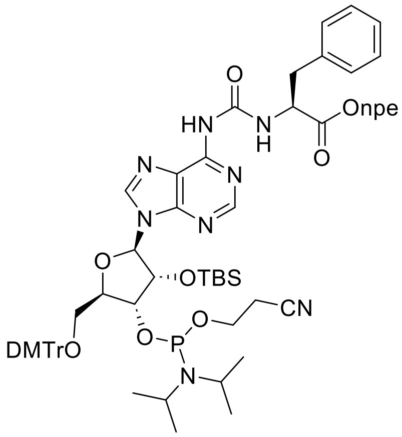


**Yield:** 67%; **$^{31}$P NMR (202 MHz, $CD_2Cl_2$) δ:** 150.77, 149.17; **HRMS (ESI):** Calcd. for $C_{64}H_{79}N_9O_{12}PSi^+$: *m/z* = 1224.5355 [M+H]$^+$, found: *m/z*: 1224.5374 [M+H]$^+$.
Literature: ref.[1]

#### 3. General information and instruments for oligonucleotides

##### 3.1 Synthesis and purification of oligonucleotides

Phosphoramidites of canonical nucleotides (Bz-rA-CE, Dmf-rG-CE, Ac-rC-CE and rU-CE) were purchased from LinkTech (LGC Biosearch Tech.) or Sigma-Aldrich. Oligonucleotides (ONs) consists of only canonical nucleotides and 6-FAM labels were ordered from Ella Biotech (Germany). Otherwise, ONs were synthesised on the corresponding 1.0 µmol CPG cartridges from LinkTech (LGC Biosearch Tech.). High Load Glen UnySupport™ was used as solid supports for strands containing non-canonical nucleotides at the 3' end. RNAs were synthesised on an Applied Biosystems 394 DNA/RNA Synthesizer with a standard phosphoramidite chemistry with a 'DMT-OFF' setting. DCA was used as a deblocking agent in $CH_2Cl_2$, BTT or Activator 42® as activator in MeCN, $Ac_2O$ as capping reagent in pyridine/THF and $I_2$ as oxidizer in pyridine/$H_2O$.

<u>Deprotection of npe and teoc groups</u>

We deprotect ONs containing amino acid-modified carbamoyl adenosine derivatives, and 5-aminomethyl uridine derivatives but not *S*-geranyl-2-thiouridine according to our previously reported methods.[5]

For the deprotection of the *para*-nitrophenylethyl (npe) group in ONs containing amino acid-modified carbamoyl adenosine derivatives, the solid support beads were suspended in 1 mL 10% DBU in THF at r.t. for 2 h. After that, the supernatant was removed and the beads were washed with 1 mL THF thrice.

For the deprotection of the 2-(trimethylsilyl)ethoxycarbonyl (teoc) group in ONs containing 5-(methyl)aminomethyl uridine derivatives but not 2-thiouridine, the solid support beads were suspended in a saturated solution of $ZnBr_2$ in 1mL of 1:1 $MeNO_2$/IPA and incubated at r.t. overnight. After that, the supernatant was removed and the beads were washed with 0.1 M EDTA in water (1 mL) and water (1 mL), once each.

<u>Cleavage from beads, deprotection of TBS groups and precipitation of the synthesized ONs</u>

The solid support beads were suspended in a 1:1 aqueous solution mixture (0.8 mL) of 30% $NH_4OH$ and 40% $MeNH_2$. The suspension was heated at 65˚C (10 min for SynBase™ CPG 1000/110 and 60 min for High Load Glen UnySupport™). Subsequently, the supernatant was collected and the beads were washed with 0.2 mL water once. The combined aqueous solutions were concentrated under reduced pressure using a SpeedVac concentrator. After that, the crude was dissolved in DMSO (100 µL) and triethylamine trihydrofluoride (125 µL) was added. The solution was heated at 65ºC for 90 mins. Finally, the ON was precipitated by adding 3 M NaOAc in water (25 µL) and *n*-butanol (1 mL). The mixture was kept at -80˚C for 2 h and centrifuged at 4˚C for 1 h. The supernatant was removed and the white precipitate was lyophilized.

For ON1-geS, the solid supports was suspended with (0.8 mL) of 30% $NH_4OH$ at r.t. for 14 h. The supernatant was collected and the beads were washed and dried similarly as above.

Followed by that, the crude was reacted with triethylamine trihydrofluoride (125 µL) in DMSO (100 µL) overnight. The ON was then precipitated, centrifuged and lyophilised as above.

Purification of the synthesized ONs by HPLC and desalting

The crude was purified by semi-preparative HPLC (1260 Infinity II Manual Preparative LC System from Agilent equipped with a G7114A detector) using a reverse-phase (RP) VP 250/10 Nucleodur 100-5 C18ec column from Macherey-Nagel. Buffers: A) 0.1 M AcOH/$Et_3N$ in $H_2O$ at pH 7 and B) 0.1 M AcOH/$Et_3N$ in 80% (v/v) MeCN in $H_2O$. Gradient: unless otherwise stated, 0-40% of B for ONs without *S*-geranyl-2-thiouridine and 0-80% of B for ONs with *S*-geranyl-2-thiouridine in 45 min. Flow rate = 5 $mL·min^{-1}$. The purified ON was analyzed by RP-HPLC (1260 Infinity II LC System from Agilent equipped with a G7165A detector) using an EC 250/4 Nucleodur 100-3 C18ec from Macherey-Nagel. Gradient: 0-40% or 0-80% of B in 45 min, similar as above. Flow rate = 1 $mL·min^{-1}$. Finally, the triethylammonium acetate was removed using a Sep-Pak® Plus Short C18 cartridge from Waters and dried with a SpeedVac concentrator.

Determination of the concentration and the mass of the synthesized ONs

The absorbance of the synthesized ON in $H_2O$ solution was measured using an IMPLEN NanoPhotometer® N60/N50 at 260 nm. The extinction coefficient of the single stranded ONs was calculated using the OligoAnalyzer Version 3.0 from Integrated DNA Technologies. For ONs incorporating non-canonical bases, the extinction coefficients were assumed to be identical to their closest canonical counterparts.

The synthesized ON (2-3 µL) was desalted on a ø 0.025 µm VSWP filter (Millipore), co-crystallized in a 3-hydroxypicolinic acid matrix (HPA, 1 µL) and analyzed by MALDI-TOF mass spectrometry (negative mode).

##### 3.2 Modification of oligonucleotides through post-synthetic reactions

Post-synthetic geranylation for 2-thiouridine containing ONs

*S*-geranyl-2-thiouridine containing ONs were geranylated from their corresponding 2-thiouridine ONs according to a reported method with minor modifications.[12] Briefly, dried and desalted ON was dissolved in 66% EtOH to achieve a final concentration of 2 mM. Geranyl bromide (250 eq.) was added to the solution and the mixture was vortexed thoroughly to fully dissolve the geranyl bromide. After that, $Et_3N$ (250 eq.) was added and the solution was incubated at r.t. with 1400 rpm on a thermoshaker for 6 h. The reaction was then quenched by adding 5X volume of 20% EtOH in HPLC buffer A and dried with a SpeedVac concentrator. The crude was then resuspended in water, filtered, purified by HPLC and desalted according to the above method.

Post-geranylation teoc-deprotection for ONs containing both $ges^2U$ and $nm^5U$

The geranylated, HPLC purified, desalted and dried ON was cooled on ice and cold 1M TBAF in THF solution was added to get a 0.5 - 1mM solution of the oligo. The solution was then incubated at 10 ˚C for 1.5 h. After that, it was cooled in an ice bath for 5 min and diluted with 5 mL cold HPLC buffer A. The solution was lyophilized overnight and purified by HPLC.

Coupling of amino acids and peptides to ONs on the solid support

Oligos with more than one amino acids attached to the *aa*$^6$A modification were prepared according to our previously reported method.[5] Briefly, the Npe group was first removed as described above. The beads were then suspended in 0.6 mL DMF solution containing the corresponding Npe protected amino acid or peptide (100 µmol), DMTMM•$BF_4$ (100 µmol) and DIPEA (200 µmol) at r.t. for 1 h. It was then washed with DMF twice and proceed to another Npe deprotection protocol. It was then carried on by the aforementioned oligo purification process.

#### 4. Characterisation data of synthesised oligonucleotides

##### 4.1 Comparison of ges$^2$U incorporation with 2 different methods

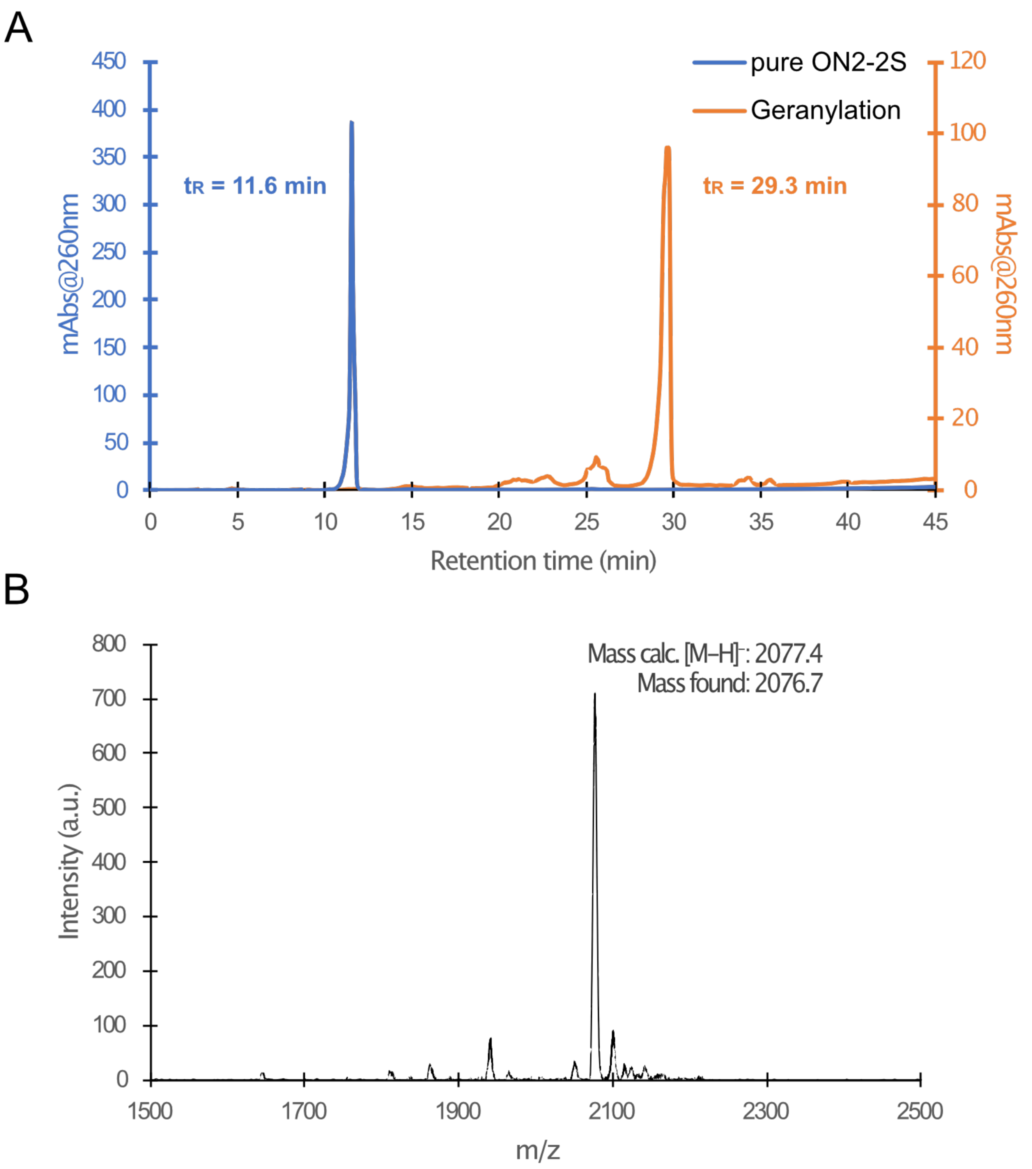


**Figure S1.** (A) HPL-chromatograms (0 → 80% B) of pure ON2-2S (blue) and its following geranylation (orange), which showed minimal amount of side products. (B) MALDI-TOF mass spectrum of purified ON2-2geS.

A

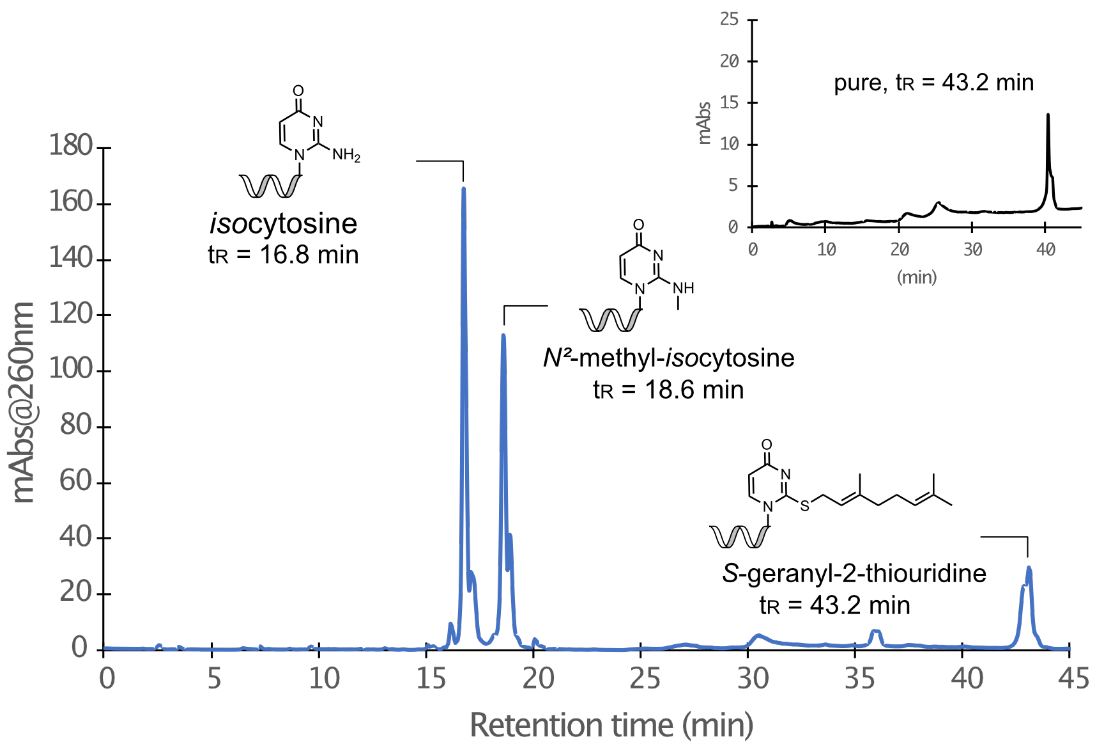


B

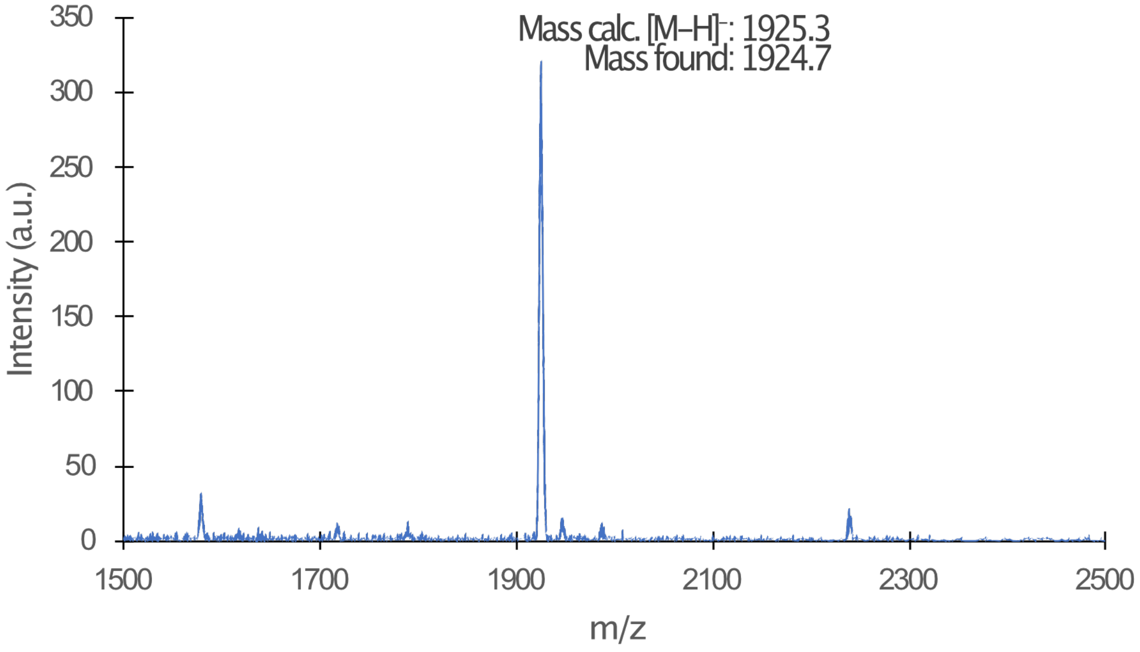


**Figure S2.** (A) HPL-chromatograms (0 → 40% B) of ON1-geS after AMA solution cleavage and 2'O-TBS deprotection, showing the side products of *iso*cytosine and $N^2$-methyl-*iso*cytosine containing oligos and the intact ON1-geS. The small chromatogram shows purified ON1-geS. (B) MALDI-TOF mass spectrum of purified ON1-geS.

#### 4.2. HPLCs and MALDI-TOFs of post-synthetic geranylations

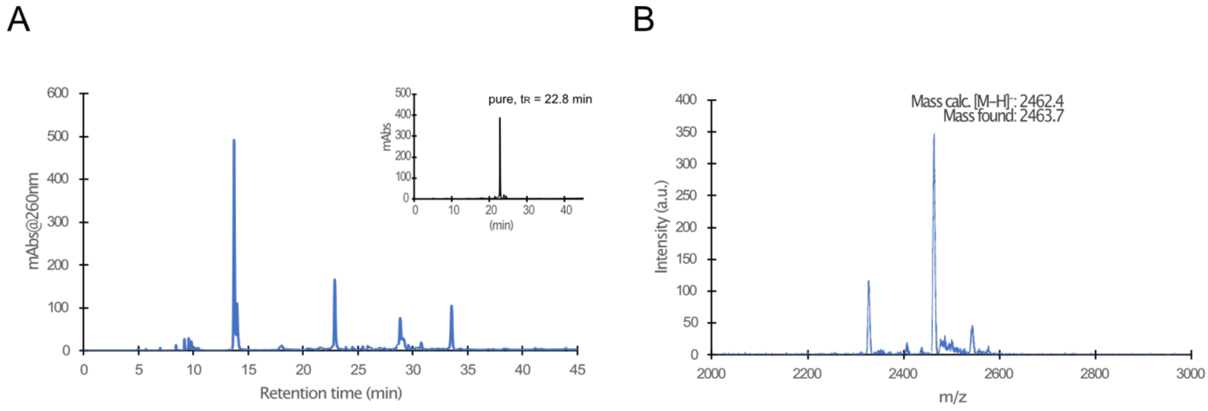


**Figure S3.** (A) HPLCgrams (0 → 80% B) of ON1-5FAM-geS after geranylation. The small chromatogram shows purified ON1-5FAM-geS. (B) MALDI-TOF mass spectrum of purified ON1-5FAM-geS.

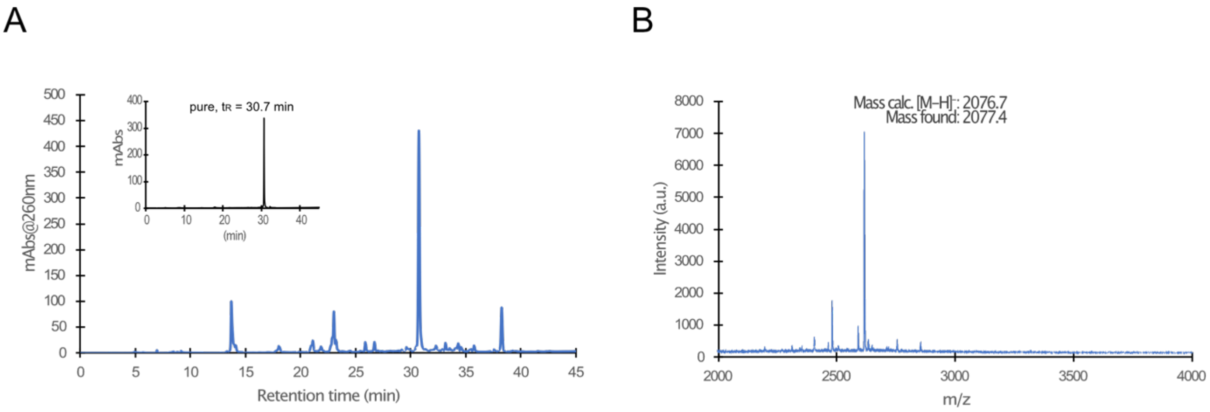


**Figure S4.** (A) HPLCgrams (0 → 80% B) of ON2-5FAM-2geS after geranylation. The small chromatogram shows purified ON2-5FAM-2geS. (B) MALDI-TOF mass spectrum of purified ON2-5FAM-2geS.

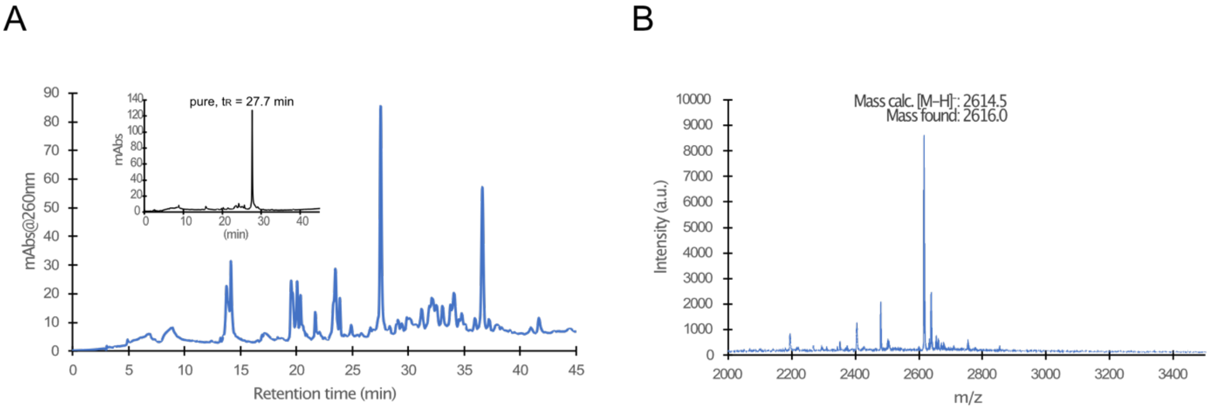


**Figure S5.** (A) HPLCgrams (0 → 80% B) of ON3-5FAM-2geS after geranylation. The small chromatogram shows purified ON3-5FAM-2geS. (B) MALDI-TOF mass spectrum of purified ON3-5FAM-2geS.

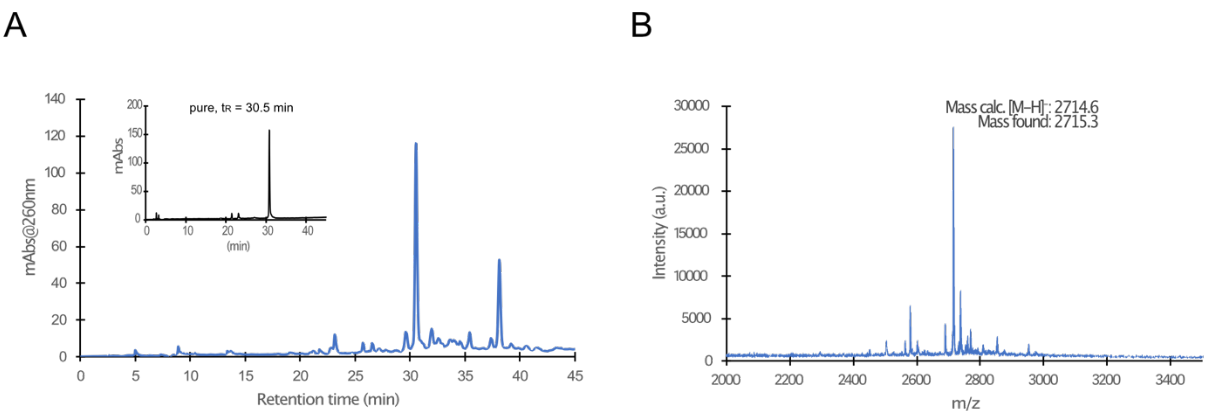


**Figure S6.** (A) HPLCgrams (0 → 80% B) of ON4-5FAM-2geS after geranylation. The small chromatogram shows purified ON4-5FAM-2geS. (B) MALDI-TOF mass spectrum of purified ON4-5FAM-2geS.

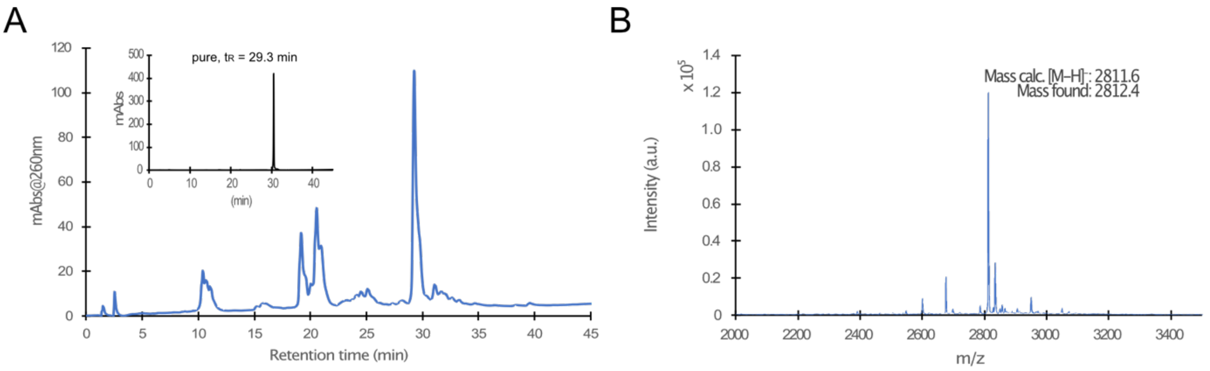


**Figure S7.** (A) HPLCgrams (0 → 80% B) of ON5-2geS after geranylation. The small chromatogram shows purified ON5-2geS. (B) MALDI-TOF mass spectrum of purified ON5-2geS.

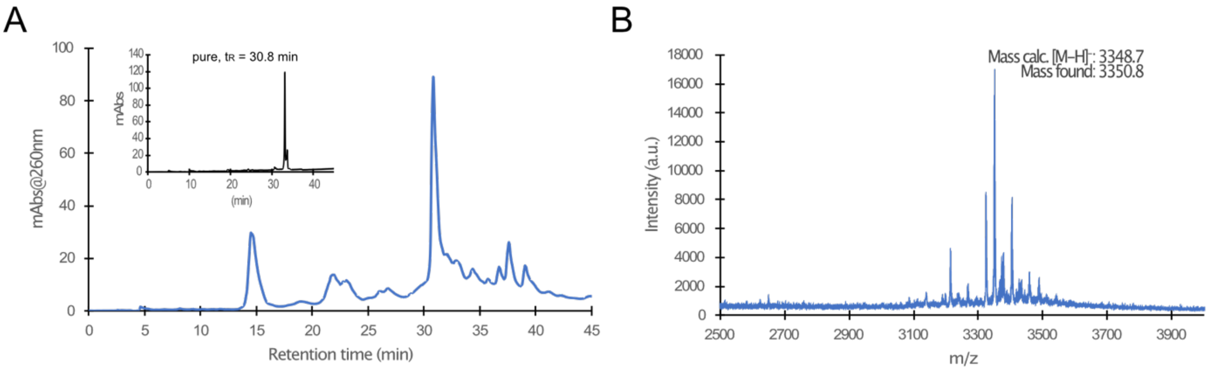


**Figure S8.** (A) HPLCgrams (0 → 80% B) of ON5-5FAM-2geS after geranylation. The small chromatogram shows purified ON5-5FAM-2geS. (B) MALDI-TOF mass spectrum of purified ON5-5FAM-2geS.

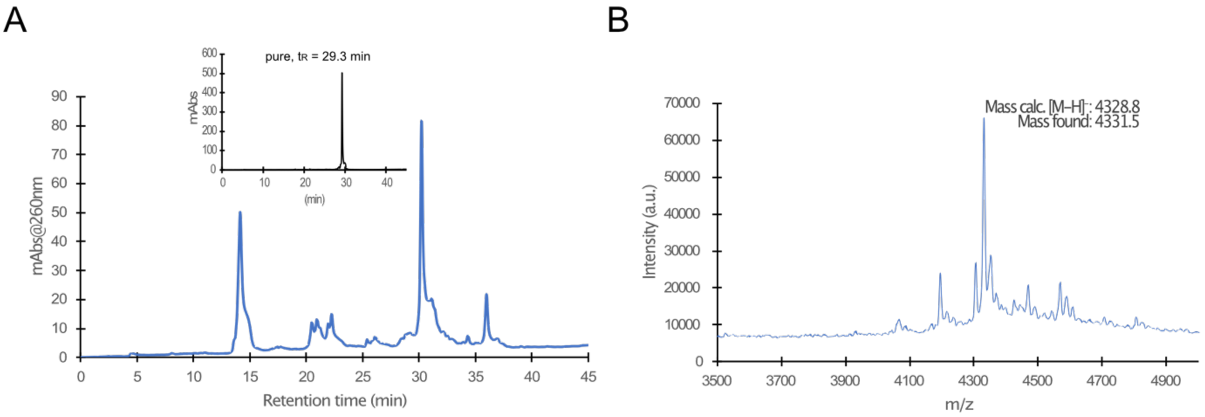


**Figure S9.** (A) HPLCgrams (0 → 80% B) of ON6-5FAM-2geS after geranylation. The small chromatogram shows purified ON6-5FAM-2geS. (B) MALDI-TOF mass spectrum of purified ON6-5FAM-2geS.

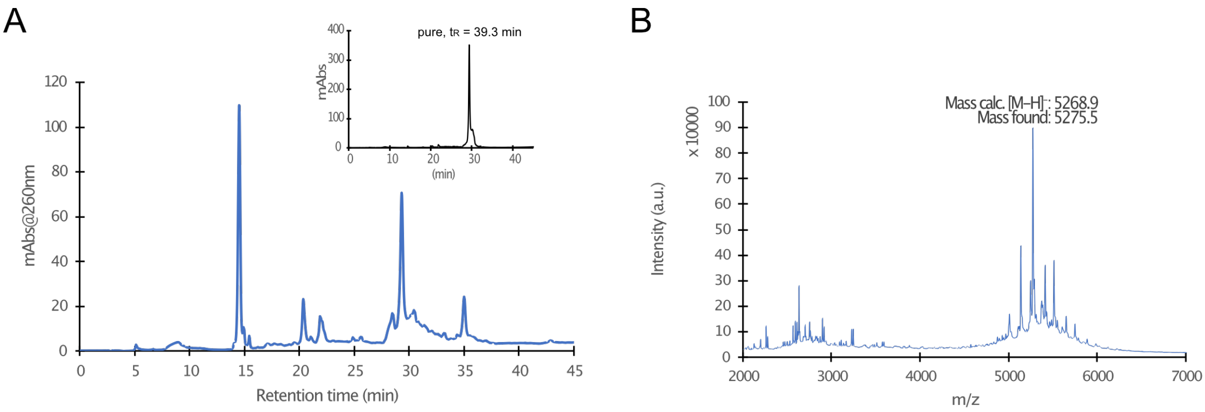


**Figure S10.** (A) HPLCgrams (0 → 80% B) of ON7-5FAM-2geS after geranylation. The small chromatogram shows purified ON7-5FAM-2geS. (B) MALDI-TOF mass spectrum of purified ON7-5FAM-geS.

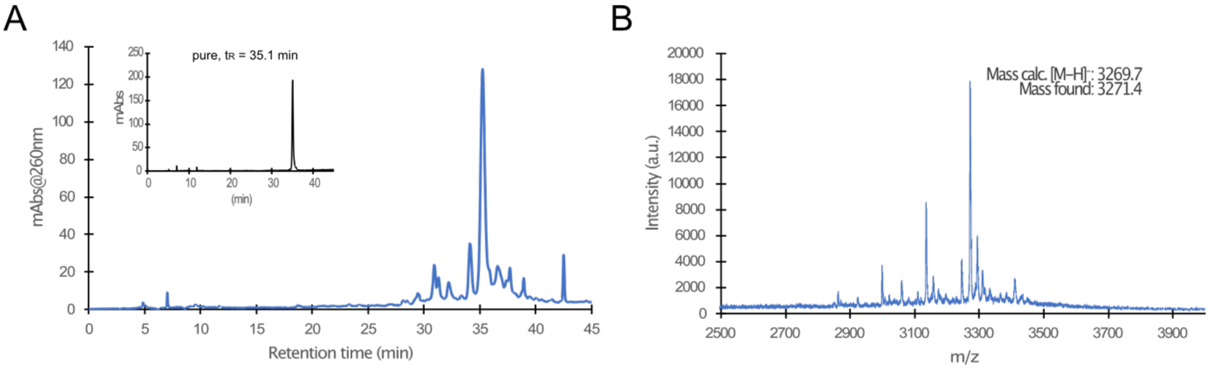


**Figure S11.** (A) HPLCgrams (0 → 80% B) of ON9-3geS after geranylation. The small chromatogram shows purified ON9-3geS. (B) MALDI-TOF mass spectrum of purified ON9-3geS.

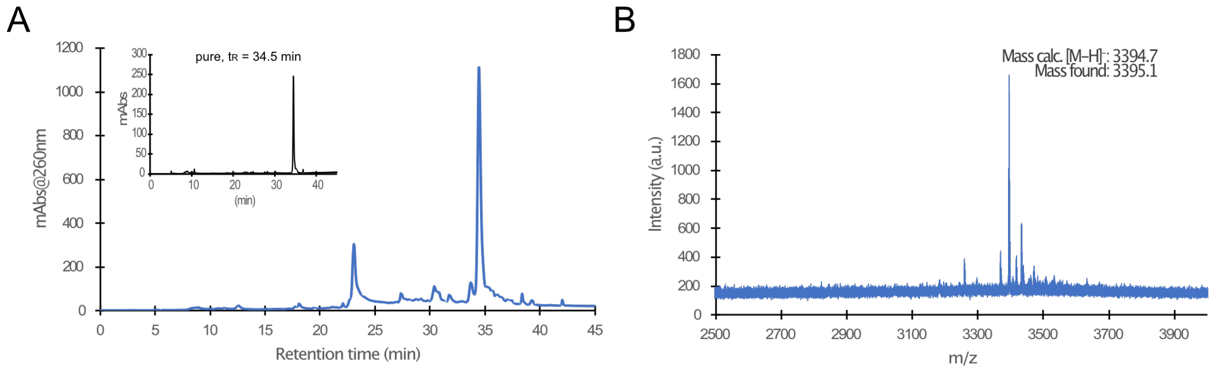


**Figure S12.** (A) HPLCgrams (0 → 80% B) of ON15-g$^6$A-3geS after geranylation. The small chromatogram shows purified ON15-g$^6$A-3geS. (B) MALDI-TOF mass spectrum of purified ON15-g$^6$A-3geS.

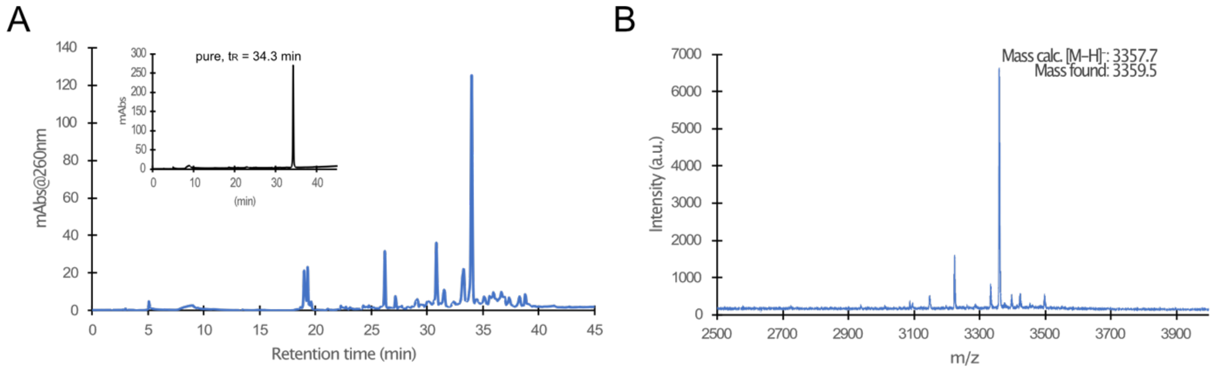


**Figure S13.** (A) HPL-chromatograms (0 → 80% B) of ON18-teocnm$^5$U-3geS after geranylation. The small chromatogram shows purified ON18-teocnm$^5$U-3geS. (B) MALDI-TOF mass spectrum of purified ON18-teocnm$^5$U-3geS.

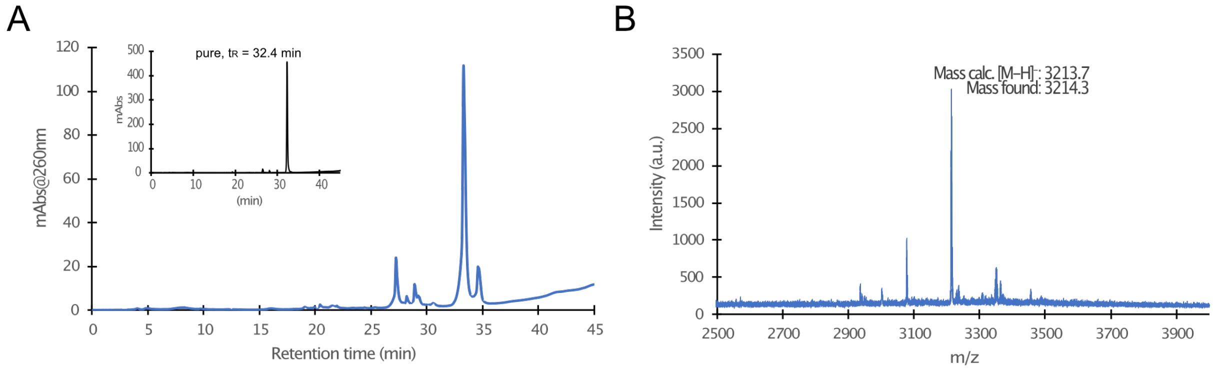


**Figure S14.** (A) HPL-chromatograms (0 → 80% B) of ON18-nm$^5$U-3geS after teoc-deprotection. The small chromatogram shows purified ON18-nm$^5$U-3geS. (B) MALDI-TOF mass spectrum of purified ON18-nm$^5$U-3geS.

#### 4.3. HPLCs and MALDI-TOFs of coupling of amino acids and peptides

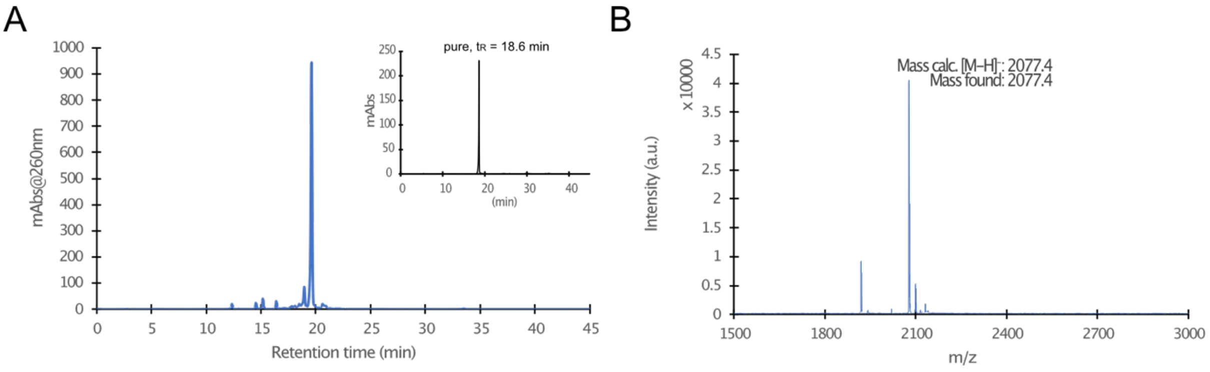


**Figure S15.** (A) HPLCgrams (0 → 40% B) of ON17-2g$^6$A after on-bead coupling of ON17-g$^6$A with Gly-Npe (compound **4a**). The small chromatogram shows purified ON17-2g$^6$A. (B) MALDI-TOF mass spectrum of purified ON17-2g$^6$A.

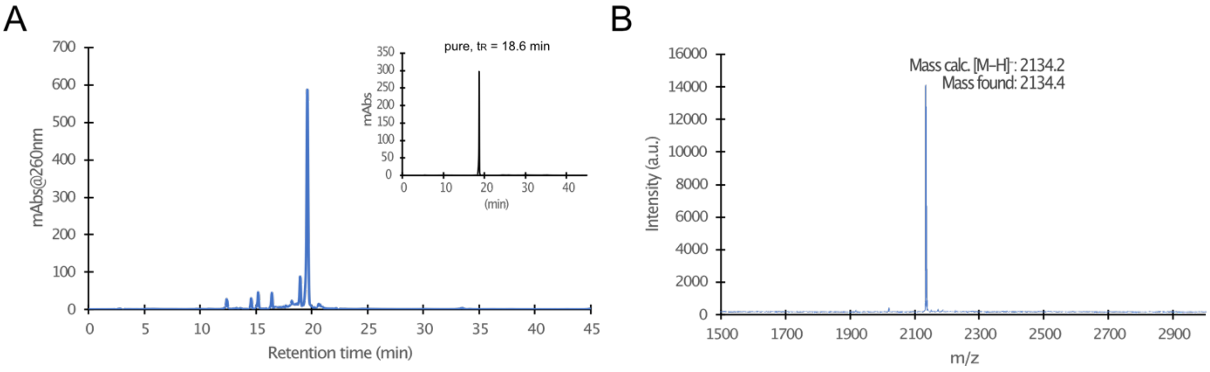


**Figure S16.** (A) HPLCgrams (0 → 40% B) of ON17-3g$^6$A after on-bead coupling of ON17-g$^6$A with Gly-Gly-Npe (compound **4b**). The small chromatogram shows purified ON17-3g$^6$A. (B) MALDI-TOF mass spectrum of purified ON17-3g$^6$A.

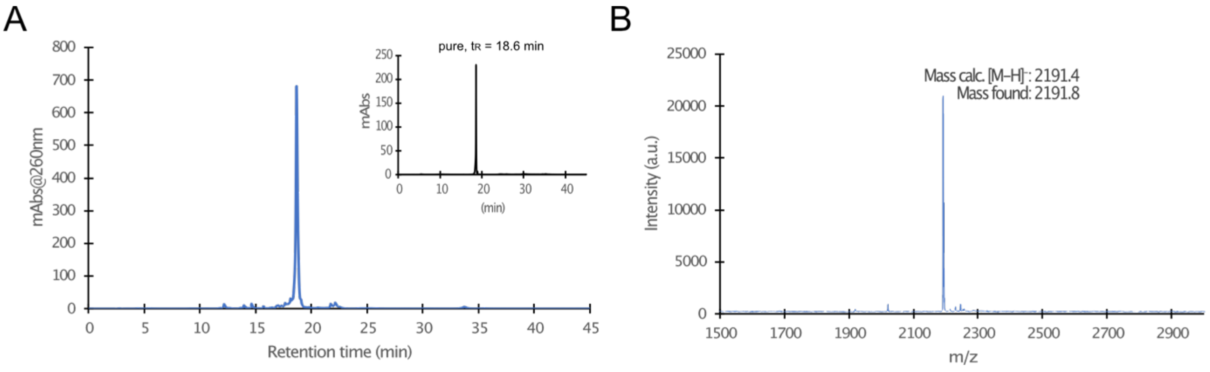


**Figure S17.** (A) HPLCgrams (0 → 40% B) of ON17-4g$^6$A after on-bead coupling of ON17-g$^6$A with Gly-Gly-Gly-Npe (compound **4c**). The small chromatogram shows purified ON17-4g$^6$A. (B) MALDI-TOF mass spectrum of purified ON17-4g$^6$A.

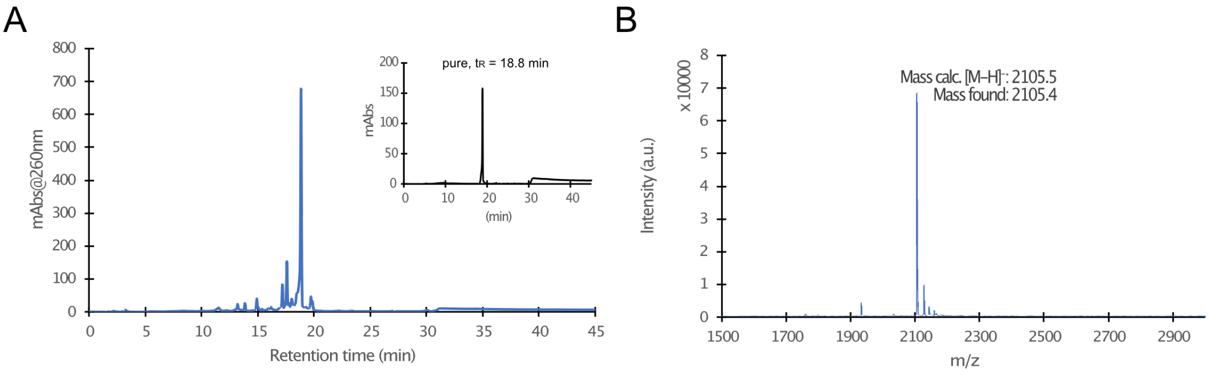


**Figure S18.** (A) HPLCgrams (0 → 40% B) of ON17-2a$^6$A after on-bead coupling of ON17-a$^6$A with Ala-Npe (compound **4h**). The small chromatogram shows purified ON17-2a$^6$A. (B) MALDI-TOF mass spectrum of purified ON17-2a$^6$A.

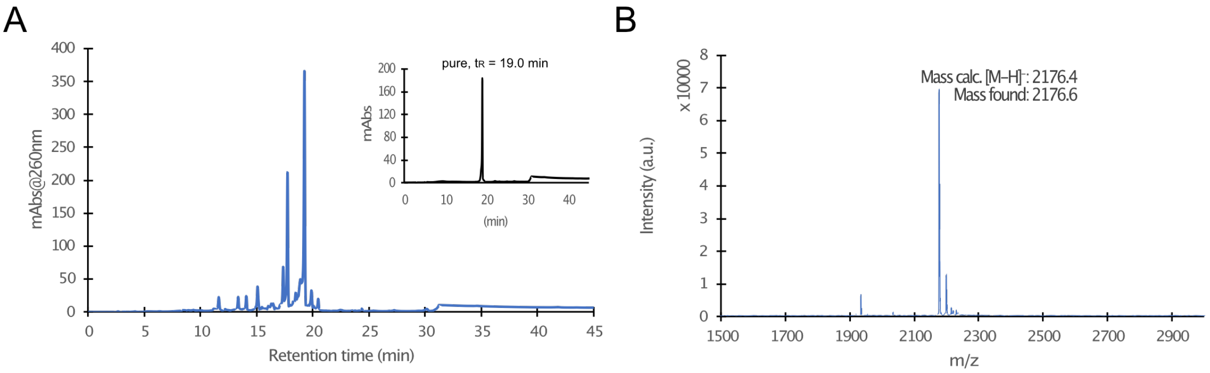


**Figure S19.** (A) HPLCgrams (0 → 40% B) of ON17-3a$^6$A after on-bead coupling of ON17-a$^6$A with Ala-Ala-Npe (compound **4q**). The small chromatogram shows purified ON17-3a$^6$A. (B) MALDI-TOF mass spectrum of purified ON17-3a$^6$A.

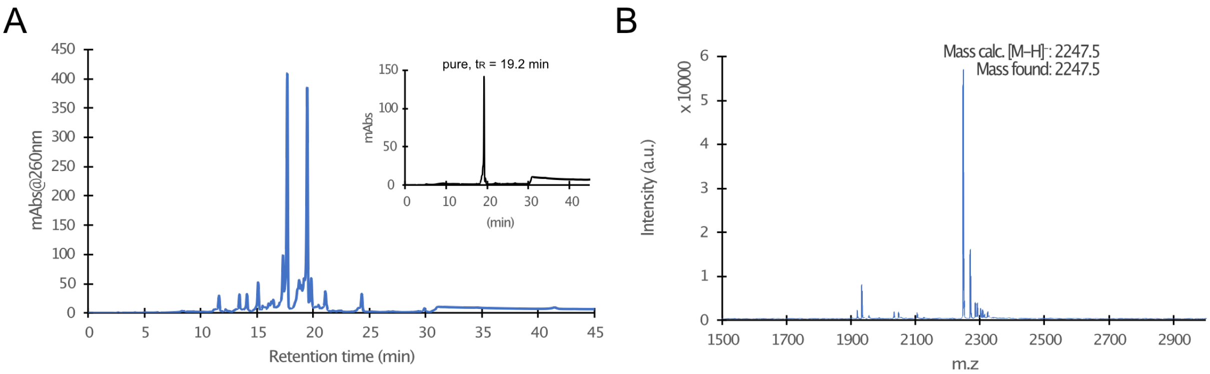


**Figure S20.** (A) HPLCgrams (0 → 40% B) of ON17-4a$^6$A after on-bead coupling of ON17-a$^6$A with Ala-Ala-Ala-Npe (compound **4r**). The small chromatogram shows purified ON17-4a$^6$A. (B) MALDI-TOF mass spectrum of purified ON17-4a$^6$A.

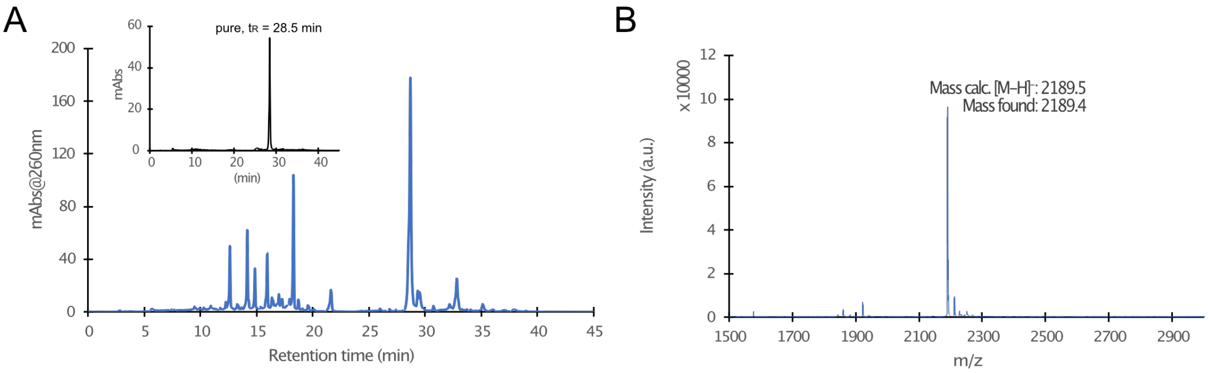


**Figure S21.** (A) HPLCgrams (0 → 40% B) of ON17-2I$^6$A after on-bead coupling of ON17-I$^6$A with Leu-Npe (compound **4d**). The small chromatogram shows purified ON17-2I$^6$A. (B) MALDI-TOF mass spectrum of purified ON17-2I$^6$A.

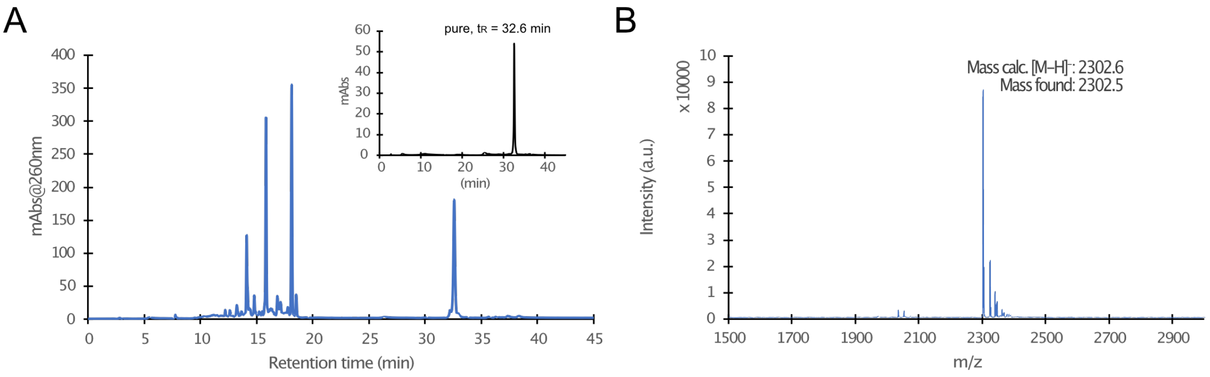


**Figure S22.** (A) HPLCgrams (0 → 40% B) of ON17-3I$^6$A after on-bead coupling of ON17-I$^6$A with Leu-Leu-Npe (compound **4k**). The small chromatogram shows purified ON17-3I$^6$A. (B) MALDI-TOF mass spectrum of purified ON17-3I$^6$A.

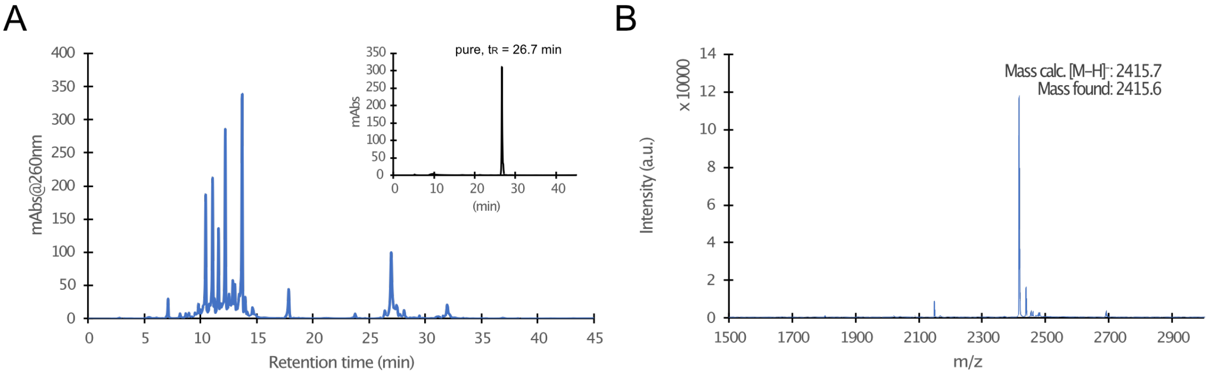


**Figure S23.** (A) HPLCgrams (0 → 60% B) of ON17-4I$^6$A after on-bead coupling of ON17-I$^6$A with Leu-Leu-Leu-Npe (compound **4l**). The small chromatogram shows purified ON17-4I$^6$A. (B) MALDI-TOF mass spectrum of purified ON17-4I$^6$A.

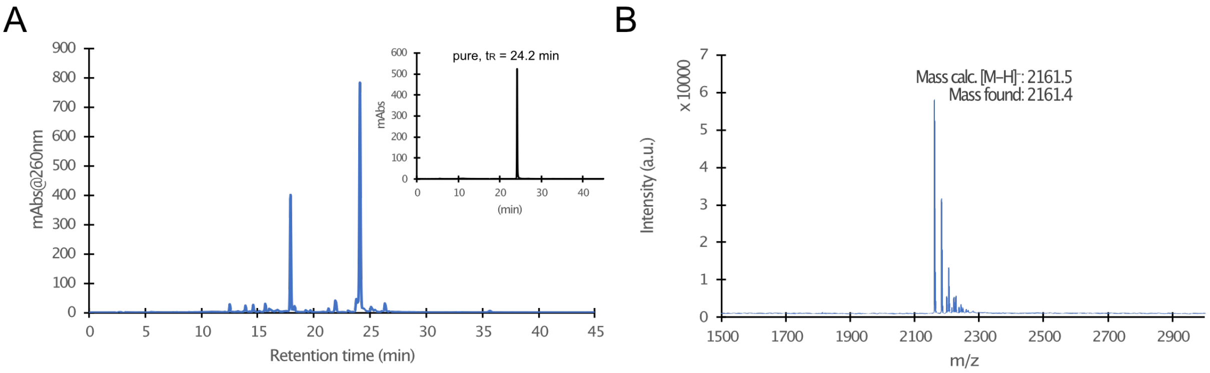


**Figure S24.** (A) HPLCgrams (0 → 40% B) of ON17-2v$^6$A after on-bead coupling of ON17-v$^6$A with Val-Npe (compound **4i**). The small chromatogram shows purified ON17-2v$^6$A. (B) MALDI-TOF mass spectrum of purified ON17-2v$^6$A.

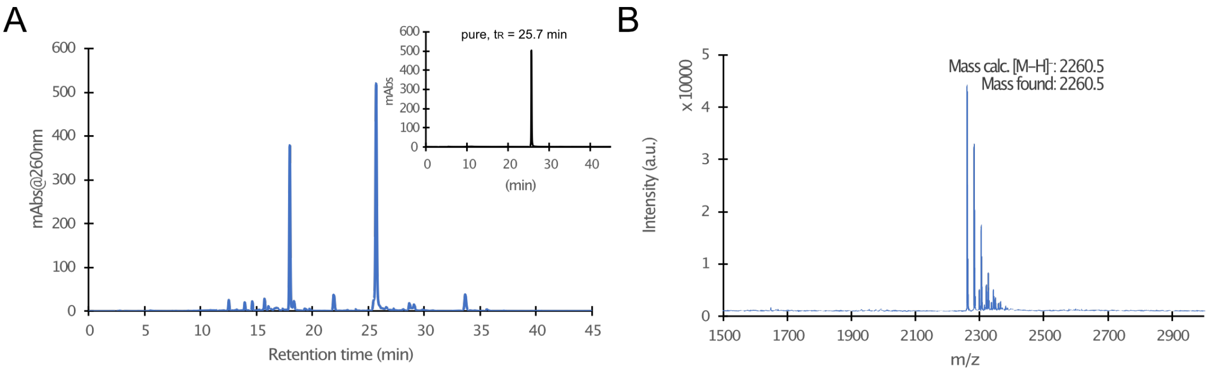


**Figure S25.** (A) HPLCgrams (0 → 40% B) of ON17-3v$^6$A after on-bead coupling of ON17-v$^6$A with Val-Val-Npe (compound **4s**). The small chromatogram shows purified ON17-3v$^6$A. (B) MALDI-TOF mass spectrum of purified ON17-3v$^6$A.

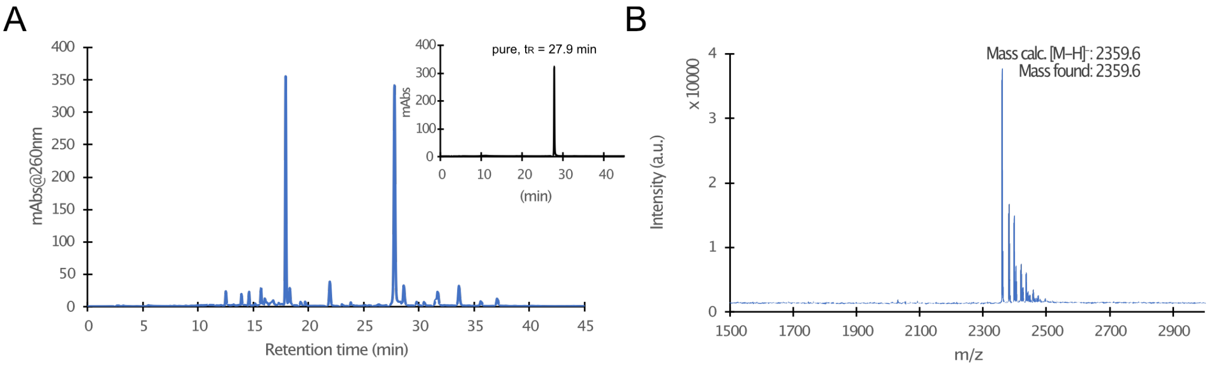


**Figure S26.** (A) HPLCgrams (0 → 40% B) of ON17-4v$^6$A after on-bead coupling of ON17-v$^6$A with Val-Val-Val-Npe (compound **4t**). The small chromatogram shows purified ON17-4v$^6$A. (B) MALDI-TOF mass spectrum of purified ON17-4v$^6$A.

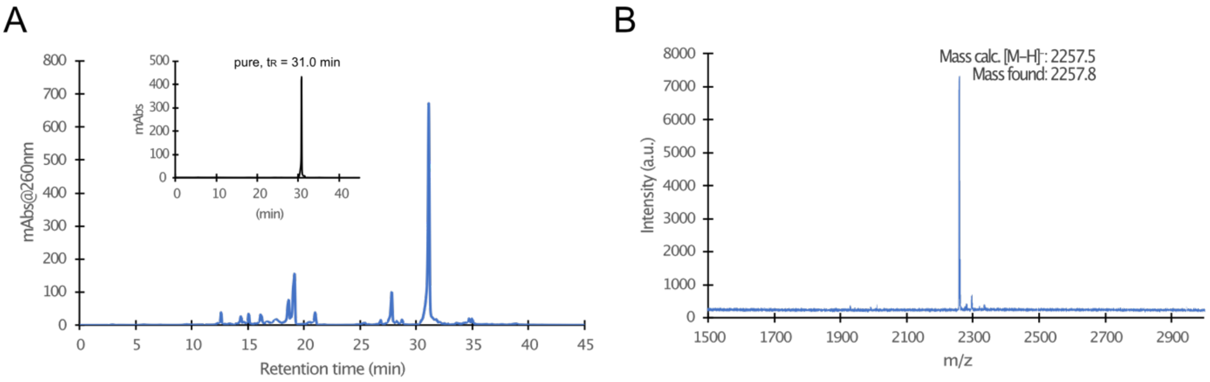


**Figure S27.** (A) HPLCgrams (0 → 40% B) of ON17-2f$^6$A after on-bead coupling of ON17-f$^6$A with Phe-Npe (compound **4f**). The small chromatogram shows purified ON17-2f$^6$A. (B) MALDI-TOF mass spectrum of purified ON17-2f$^6$A.

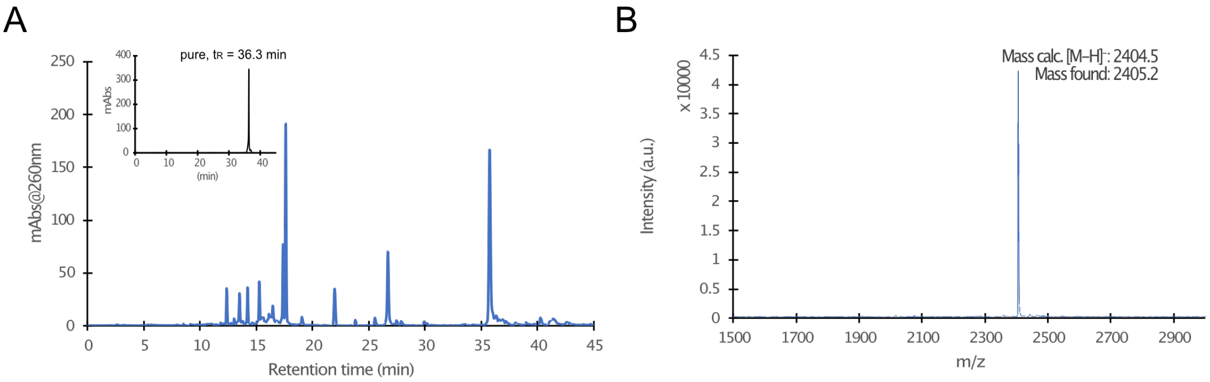


**Figure S28.** (A) HPL-chromatograms (0 → 40% B) of ON13-3f$^6$A after on-bead coupling of ON13-f$^6$A with Phe-Phe-Npe (compound **4o**). The small chromatogram shows purified ON13-3f$^6$A. (B) MALDI-TOF mass spectrum of purified ON13-3f$^6$A.

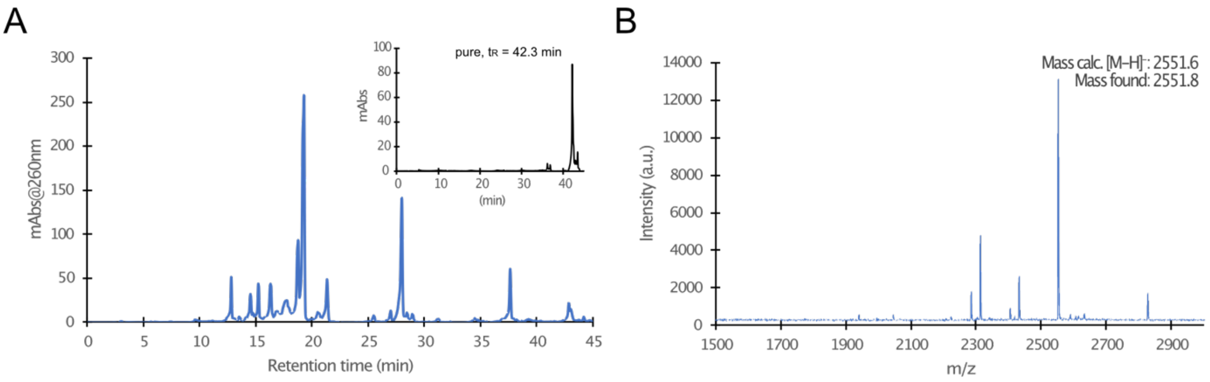


**Figure S29.** (A) HPLCgrams (0 → 40% B) of ON17-4f$^6$A after on-bead coupling of ON17-f$^6$A with Phe-Phe-Phe-Npe (compound **4p**). The small chromatogram shows purified ON17-4f$^6$A. (B) MALDI-TOF mass spectrum of purified ON17-4f$^6$A.

#### 5. General information for liposome preparation

Egg PC (Avanti Polar Lipids) was dissolved in chloroform to achieve a concentration of 20 mM. This solution was prepared as stock and stored in -20 ˚C. To produce liposome suspension, a required volume of the chloroform solution was taken, and a lipid film can be formed by vaporizing the chloroform in a round bottom flask under reduced pressure. Unless otherwise stated, the film was then rehydrated by a 2X reaction buffer and suspended completely in the solution through thorough vortexing.

The liposome suspension was then pressed through a mini-extruder (Avanti Polar Lipids) equipped with a ø 100 nm Nuclepore™ Track-Etched membrane (Whatman) for 15 times and a small sample of the suspension was analysed by dynamic light scattering (DLS, Wyatt DyanPro NanoStar). For confocal microscopy, the extrusion and DLS steps were skipped. For flow cytometry and confocal microscopy, 1% and 0.1 % mol of Nile Red (Sigma-Aldrich) was added to stain the liposome respectively. The suspension was then diluted to a final concentration of 10 mM Egg PC, 100 mM NaCl and 100 mM MES buffer pH6.0, plus the desired concentration of ONs as stated at each experiment.

#### 5.1 Dynamic Light Scattering (DLS) of extruded liposome suspension

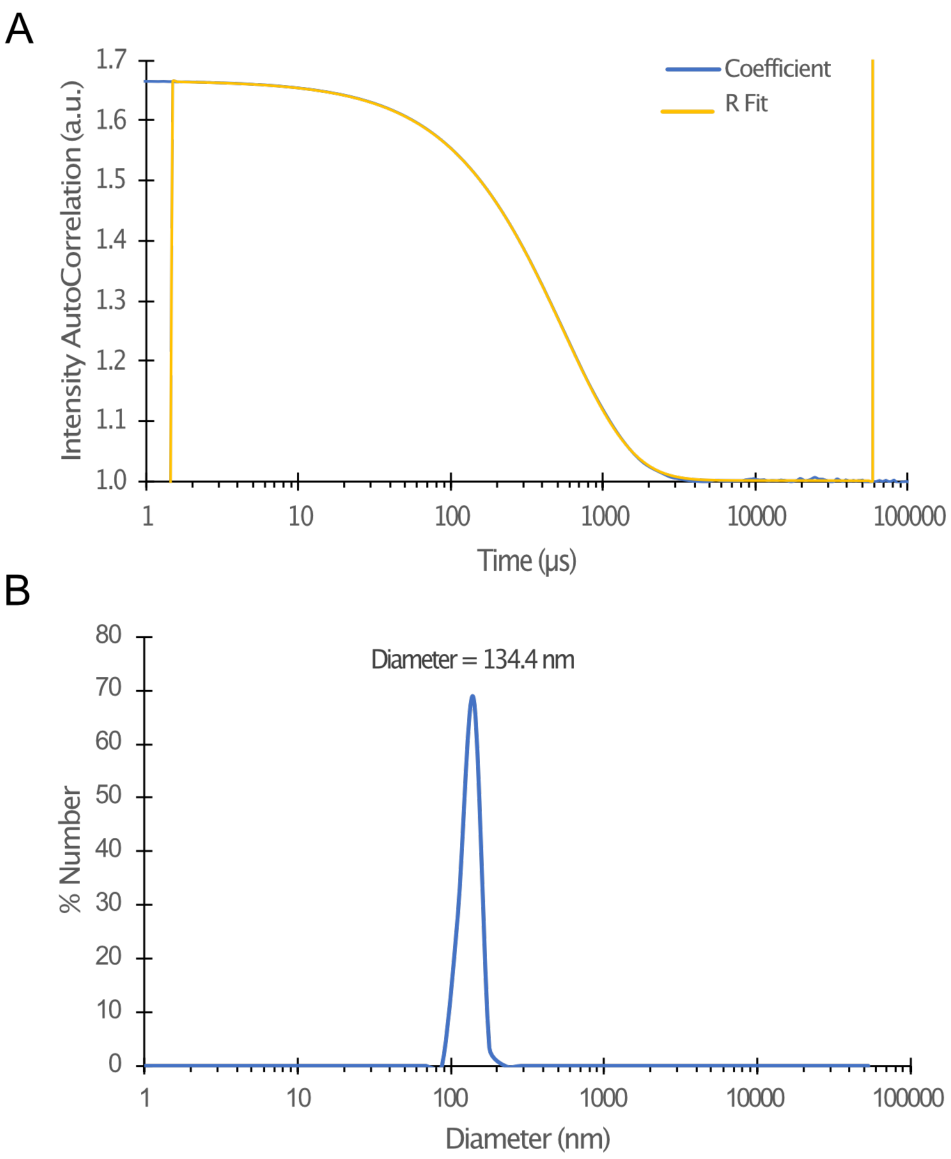


**Figure S30.** A representative dynamic light scattering analysis of one of the prepared extruded Egg PC liposome samples. (A) Raw data of the autocorrelation coefficient of scattering intensity over time (blue) and the selected regularisation fitting region (yellow). (B) The calculated size distribution (number %) of the liposome, with the peak at 134.4 µm diameter.

#### 6. General information for confocal microscopy and FACS

##### 6.1 General information for confocal microscopy

Liposomes were prepared according to the method described above with 10 µM of FAM-labeled ONs. Without further incubation, 10 µL of the sample was then transferred to a sample slide.

The slides were then imaged on a Leica TCS SPE confocal microscope with an inverted microscopic stage. For the detection of Nile Red stained liposome, images were acquired with laser excitation of 532 nm and emission from 550-600 nm. For the detection of FAM-labelled ONs, images were acquired with laser excitation of 488 nm and emission from 500-550 nm.

##### 6.2 General information for flow cytometry

Extruded liposomes were prepared according to the method described above, with a final concentration of 2.5 µM of each of the RNAs in 400 µL. Samples with liposome, were then spun down by a centrifugal force of 21.130 *g* for 3 h in 8 ˚C. The supernatant was carefully removed and 400 µL of fresh buffer (100 mM NaCl and 100 mM MES buffer pH6.0) was used to resuspend the liposome. This washing process was repeated once before the cytometry measurement. Apart from samples containing RNA, controls of pure buffer, unstained liposome, negative control of stained liposome (marked as '-ve' below), and stained liposome + 0.1% Triton™ X-100 were prepared in a similar manner for gate selections (see section X.3).

Flow cytometry experiments had been performed on a LSR Fortessa™ cell analyzer (Becton Dickinson). Buffer only and Triton™ controls had been run for 5 min to ensure proper exclusion of background. In all samples at least 5000 events have been recorded. Nile Red fluorescence was measured on the 'PE-Cy5-5' setting and the fluorescein signal was measured on the 'FITC' setting provided by the instrument software. Gates 1 & 2 were created based on the background of the controls as shown below (see Figure S31) and population within Gate 2 were analyzed with FlowJo™ version 10.9.0 (Becton Dickinson).

#### 6.3 Controls for flow cytometry data

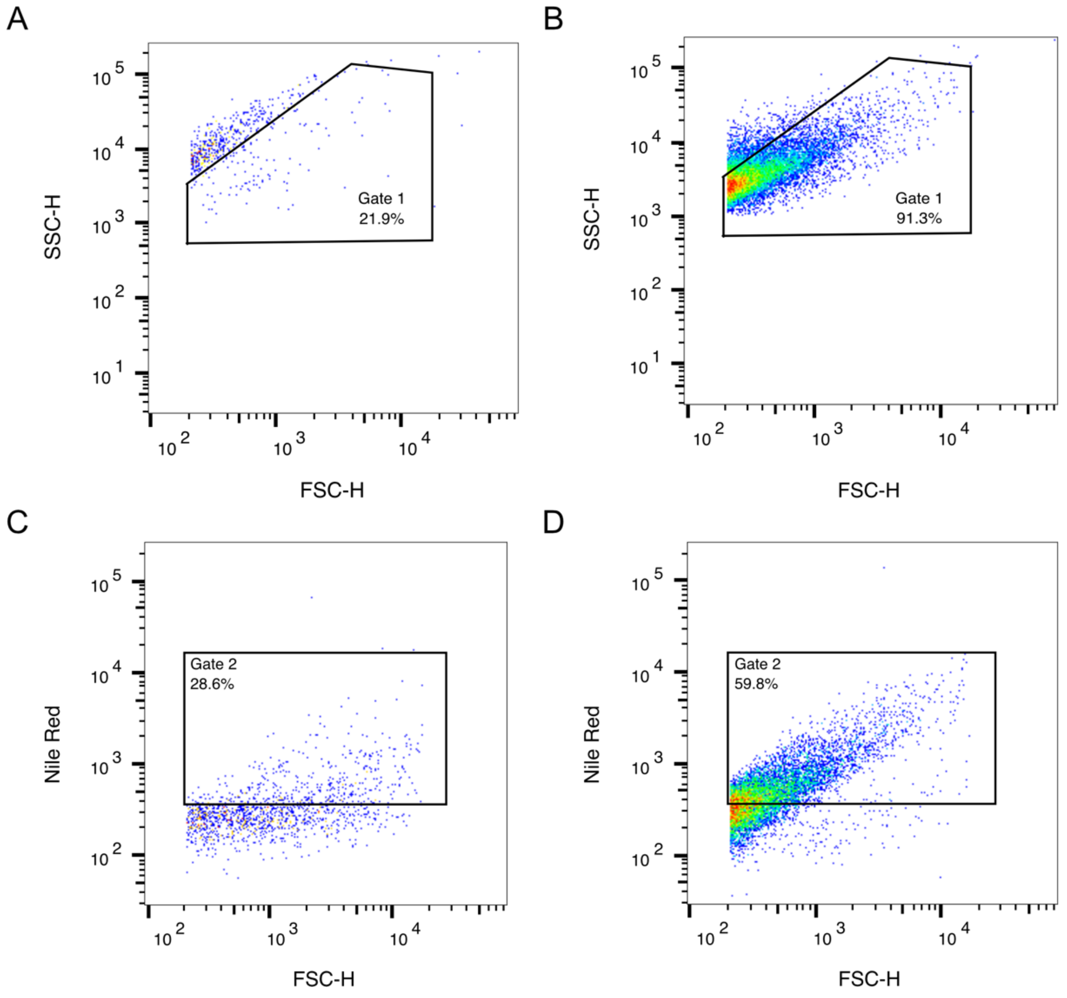


**Figure S31.** Flow cytometry data of (A) Buffer control, (B) Unstained liposome, (C) Nile Red stained liposome + 0.1% Triton™ X-100, and (D) Nile Red stained liposome (-ve). (A)&(B): Gate 1 was selected by excluding the background events caused by the buffer (100 mM NaCl and 100 mM MES buffer pH 6.0). (C)&(D): Within Gate 1, Gate 2 was created by excluding background Nile Red signal that is not caused by its localization in the lipid phase.

##### 6.4 Confocal images and cytometry data

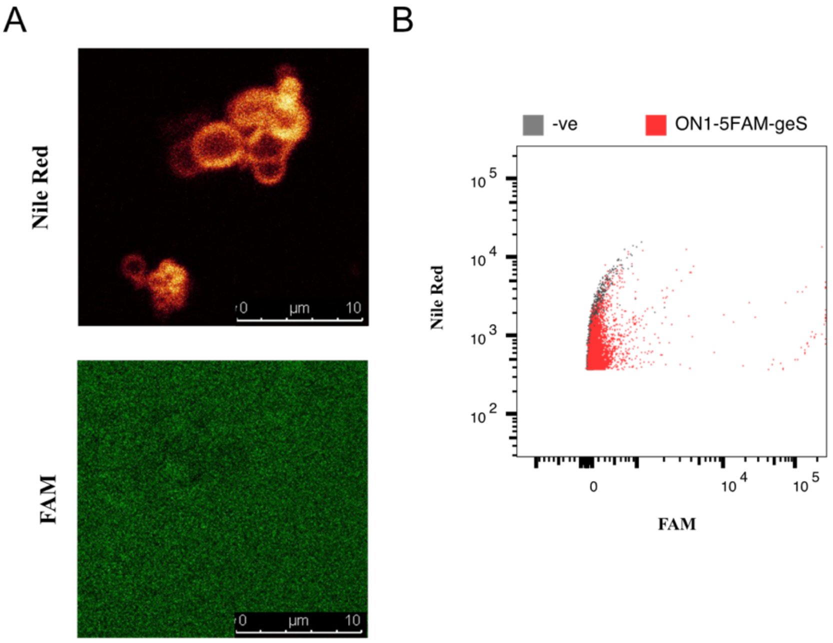


**Figure S32.** (A) Confocal images and (B) cytometry data of ON1-5FAM-geS. The Nile Red and FAM images suggested that the ssRNA does not bind to the liposome. The cytometry data supports this result by showing 2 overlapping populations between the -ve control and the regarding RNA-liposome mixture, with the FAM emission medians of 72 for -ve and 116 for the sample.

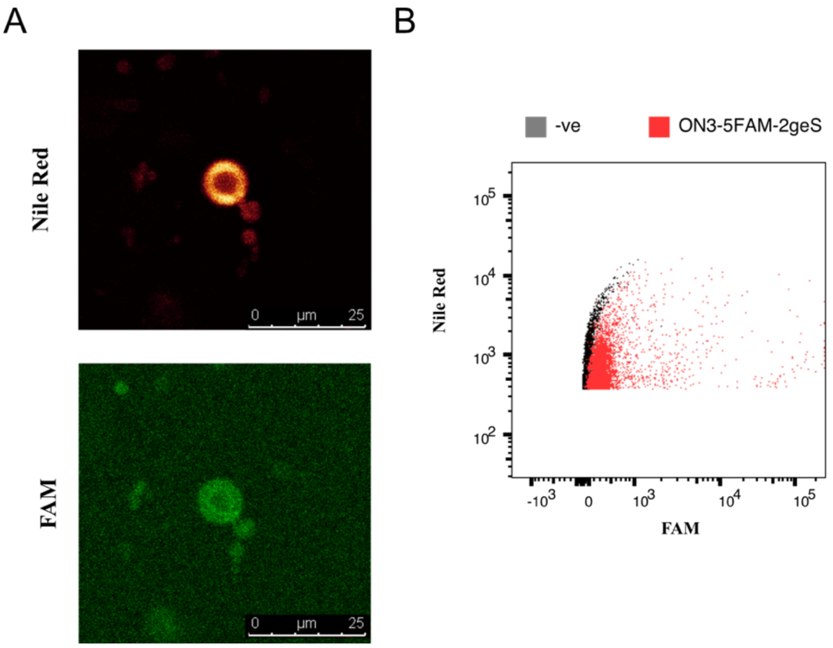


**Figure S33.** (A) Confocal images and (B) cytometry data of ON3-5FAM-2geS. The Nile Red and FAM images suggested that the ssRNA weakly binds to the liposome. The cytometry data supports this result by showing the ON3-5FAM-2geS population slightly shifted from the -ve control, with the FAM emission medians of 72 for -ve and 289 for the sample.

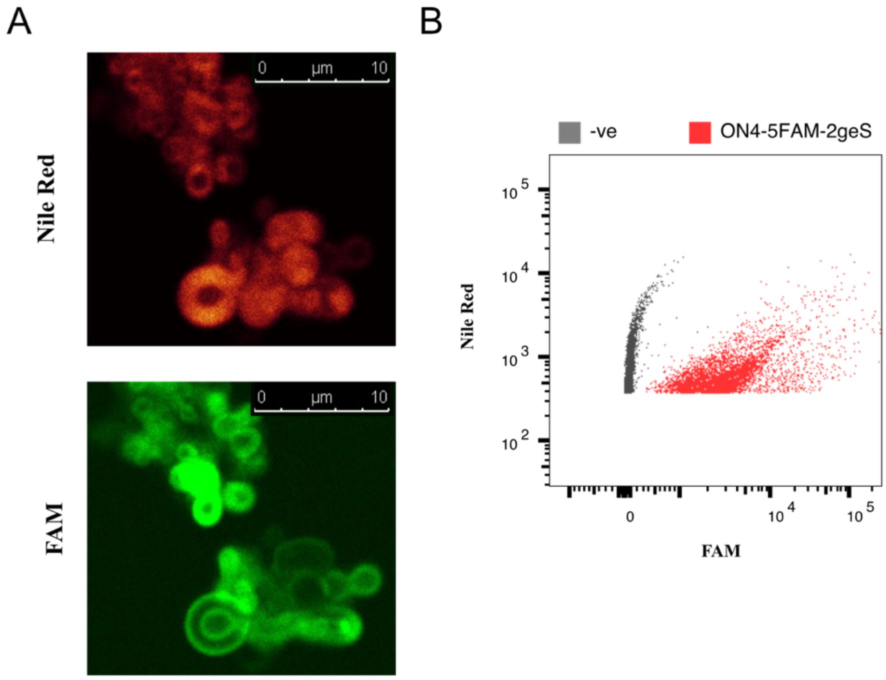


**Figure S34.** (A) Confocal images and (B) cytometry data of ON4-5FAM-2geS. The Nile Red and FAM images suggested that the ssRNA binds to the liposome. The cytometry data supports this result by showing a shifted population of the regarding RNA-liposome sample from the -ve control, with the FAM emission medians of 72 for -ve and 3010 for the sample.

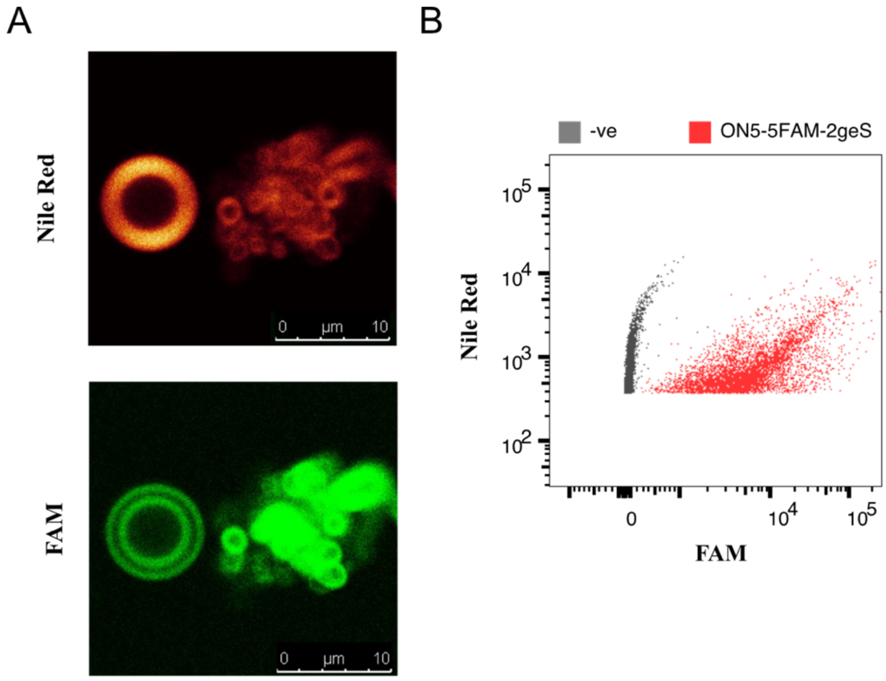


**Figure S35.** (A) Confocal images and (B) cytometry data of ON5-5FAM-2geS. The Nile Red and FAM images suggested that the ssRNA binds to the liposome. The cytometry data supports this result by showing a shifted population of the regarding RNA-liposome sample from the -ve control, with the FAM emission medians of 72 for -ve and 5191 for the sample.

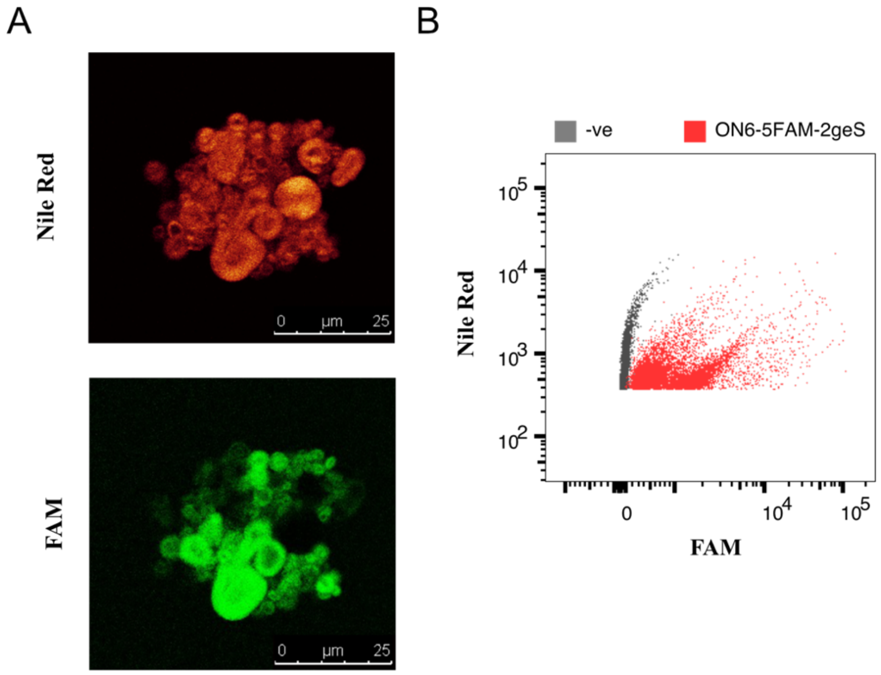


**Figure S36.** (A) Confocal images and (B) cytometry data of ON6-5FAM-2geS. The Nile Red and FAM images suggested that the ssRNA binds to the liposome. The cytometry data supports this result by showing a shifted population of the regarding RNA-liposome sample from the -ve control, with the FAM emission medians of 72 for -ve and 1002 for the sample.

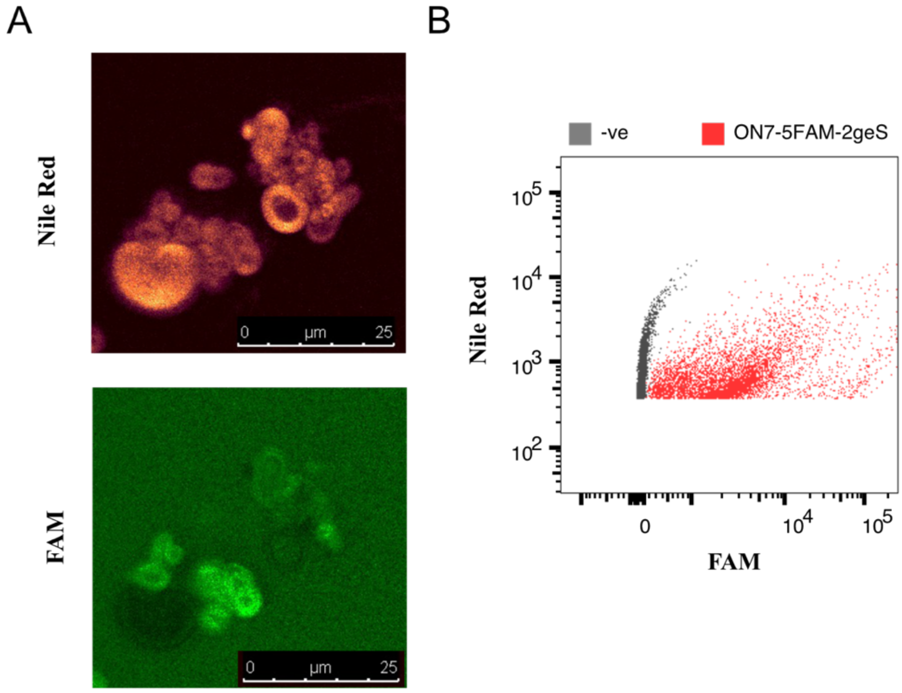


**Figure S37.** (A) Confocal images and (B) cytometry data of ON7-5FAM-2geS. The Nile Red and FAM images suggested that the ssRNA does not bind to the liposome. However, the cytometry data still showed a shifted population of the regarding RNA-liposome sample from the -ve control, with the FAM emission medians of 72 for -ve and 2438 for the sample.

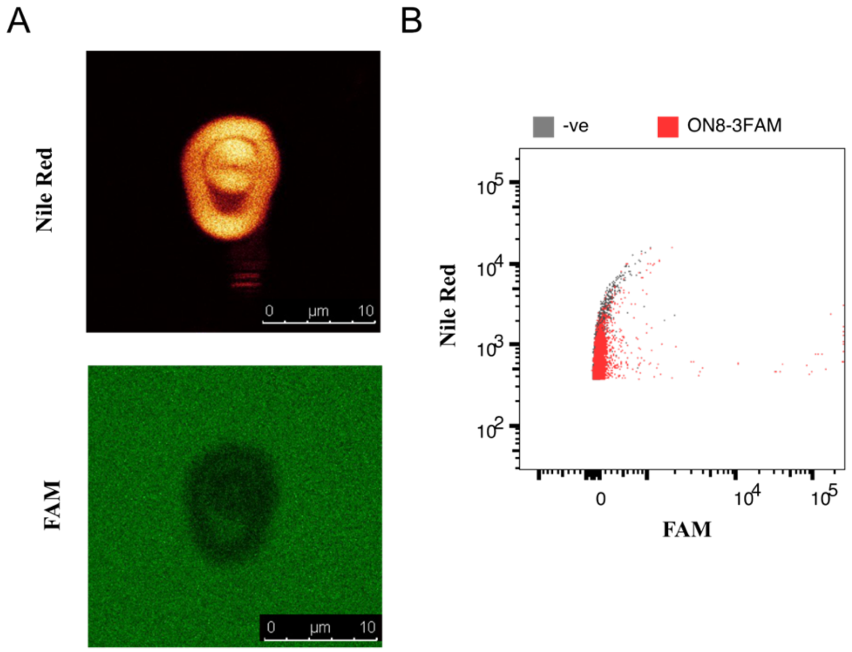


**Figure S38.** (A) Confocal images and (B) cytometry data of ON8-3FAM. The Nile Red and FAM images suggested that the ssRNA does not bind to the liposome. The cytometry data supports this result by showing 2 overlapping populations between the -ve control and the regarding RNA-liposome mixture, with the FAM emission medians of 72 for -ve and 82 for the sample.

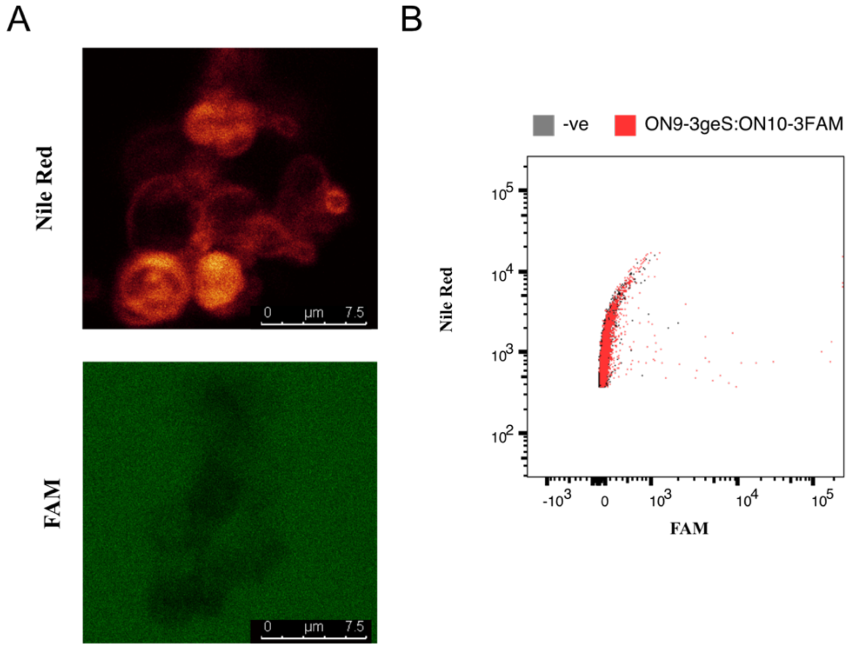


**Figure S39.** (A) Confocal images and (B) cytometry data of ON9-3geS:ON10-3FAM dsRNA. The Nile Red and FAM images suggested that the ssRNA does not bind to the liposome. The cytometry data supports this result by showing 2 overlapping populations between the -ve control and the regarding RNA-liposome mixture, with the FAM emission medians of 72 for -ve and 96 for the sample.

#### 7. General information for liposom-catalysed geranylation

##### 7.1 Liposome-assisted geranylation of s$^2$U ONs

Vesicle solution was prepared by thin film rehydration with an aqueous solution containing NaCl (100 mM) and borate buffer (pH = 10.0, 100 mM) giving a final EggPC concentration of 40 mM. The ONs for the reaction (2 nmol each) were put into a 1.5 mL eppendorf tube and the ONs were dried *in vacuo*. Vesicle solution (100 µl) was added to the dried ONs and geBr (2.0 µmol, 0.397 µL) was added to the mixtures followed by vortexing. The reaction mixtures were incubated at 1000 rpm in r.t. over a period of 2 h. It was then injected to HPLC, and peaks were collected and analysed by MALDI-TOF mass spectrometry.

##### 7.2 HPLCs for liposome-catalysed geranylation

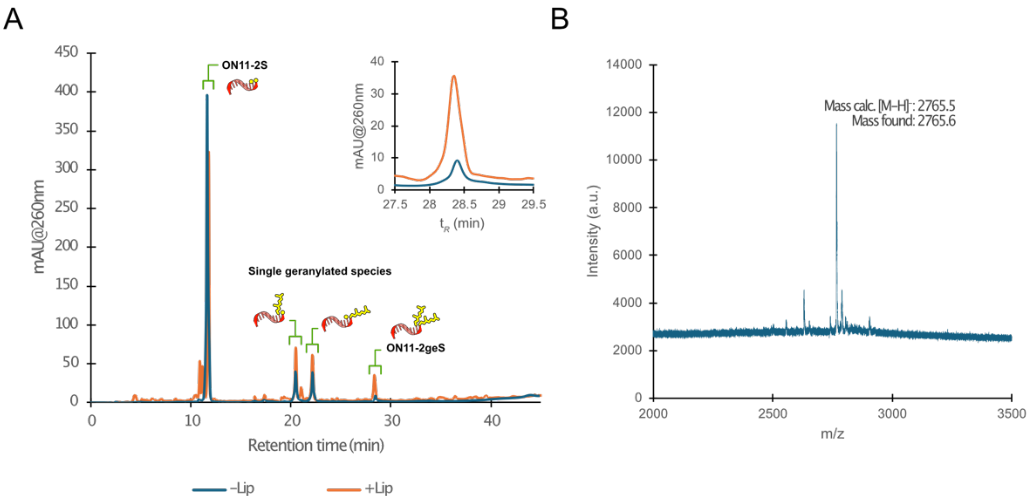


**Figure S40.** (A) Overlay of HPLCgrams between the -Lip (blue) and +Lip (orange, 40 mM Egg PC) geranylation for ON11-2S with 20 mM geBr in pH10. The yield of fully geranylated ON11-2geS increased from 3.1±0.4% in -Lip to 10.5±1.8% in +Lip. Top right showing the magnified region of the ON11-2geS peak. (B) MALDI-TOF mass spectrum of the collected ON11-2geS.

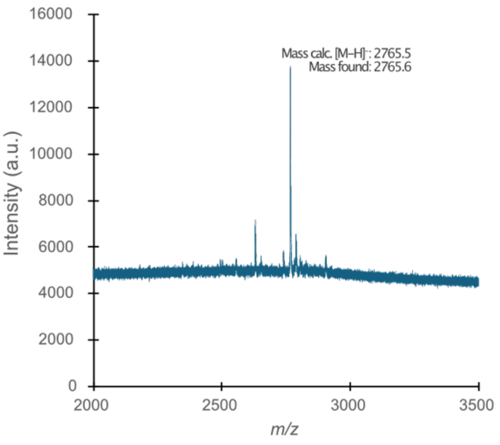


**Figure S41.** MALDI-TOF mass spectrum of the collected ON12-2geS. HPLCgrams are shown in Ext. Data Fig. 1.

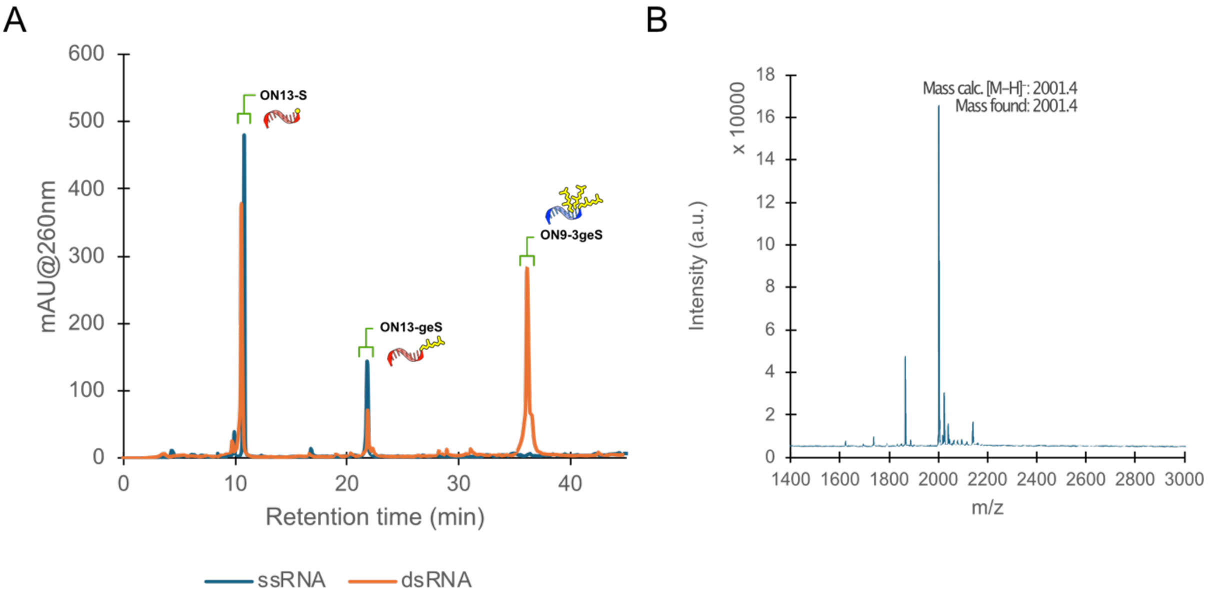


**Figure S42.** (A) Overlay of HPLCgrams between ssRNA ON13-S (blue) and dsRNA ON13-S:ON9-3geS (orange) geranylation in 20 mM geBr and 40 mM Egg PC, pH10. The yield of geranylated ON-13-geS decreased from 10.5±1.8% in ssRNA to 3.1±0.4% in dsRNA. (B) MALDI-TOF mass spectrum of the collected ON13-geS.

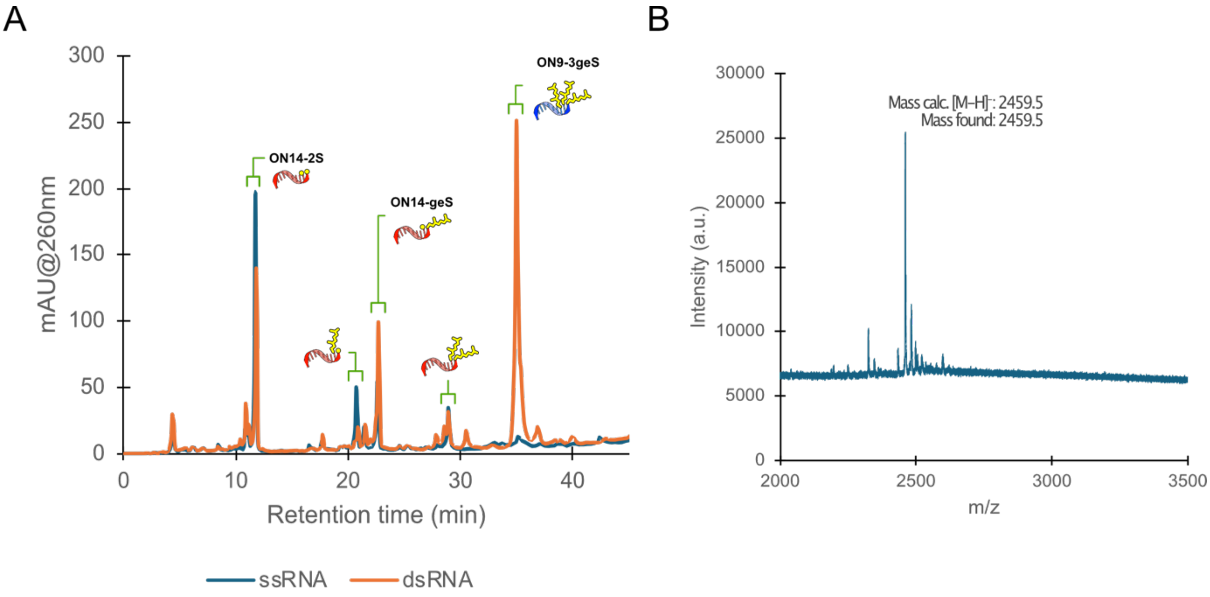


**Figure S43.** (A) Overlay of HPLCgrams between ssRNA ON14-2S (blue) and dsRNA ON14-2S:ON9-3geS (orange) geranylation in 20 mM geBr and 40 mM Egg PC, pH10.0. The yield of partially geranylated ON14-geS increased from 19.6±1.6% in ssRNA to 36.4±2.9% in dsRNA. (B) MALDI-TOF mass spectrum of the collected ON14-geS.

#### 8. General information for ges$^2$U hydrolysis

Stability study of ges$^2$U hydrolysis in the presence or absence of liposome

Liposome solution was prepared as mentioned above containing NaCl (200 mM) and acetate (pH=7.0, 100 mM) or glycine buffer (pH=10, 100 mM), giving a final Egg PC concentration of 20 mM. The ONs for the reaction (5.5 nmol each) were put into a 1.5mL Eppendorf tube and the ONs were dried in vacuo. Vesicle solution (275 µl) was added to the dried ONs followed by addition of rehydration buffer (275 µl), to achieve a final concentration of 100 mM NaCl, 50 mM buffer and 10 mM Egg PC. For the -Lip setup, Egg PC addition was excluded. The samples were vortexed and shaken with a Thermomixer (1000 rpm) at r.t. for 4 days. After 0, 24, 48, 72 and 96 h an aliquot of 100 µl were removed from the samples and analysed by HPLC.

##### 8.1 HPLCs of ges$^2$U hydrolysis

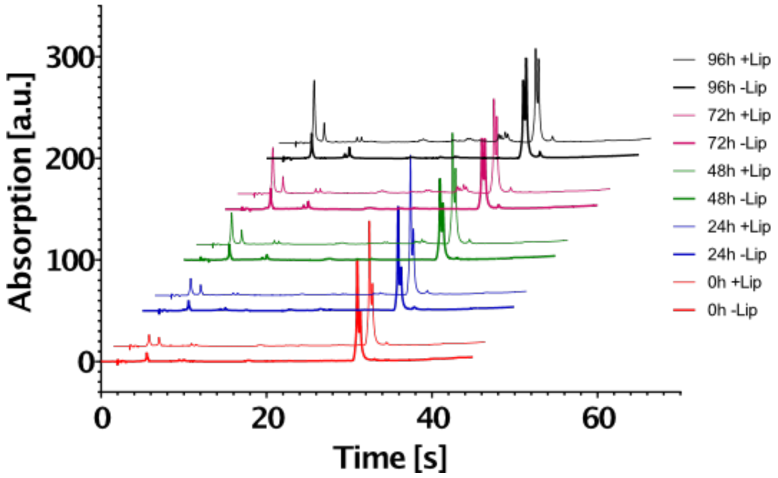


**Figure S44.** HPLCgrams of ON12-2geS incubation in acetate buffer (pH = 7.0) over 4 d at r.t. The integrity of ON12-geS was summarized in Ext. Data Fig. 2.

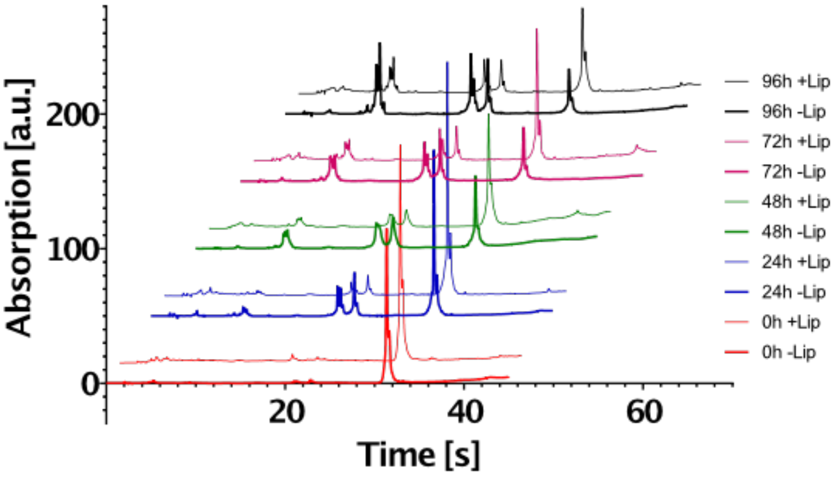


**Figure S45.** HPLCgrams of ON12-2geS incubation in glycine buffer (pH = 10.0) over 4 d at r.t. The integrity of ON12-2geS was summarized in Ext. Data Fig. 2.

#### 9. General information for UV-Vis melting

Samples were prepared to a concentration of 100 mM Tris-HCl buffer, 100 mM NaCl and 10 µM dsRNA, with a total volume of 400 µL. UV melting experiments were performed on an Jasco V-650 UV-Vis Spectrophotometer, using 1-cm path length quartz cuvette. The samples were scanned from 5 to 85 °C with a temperature incremental rate of 1.0 ˚C/min. The temperature was hold at 85 °C for 5-minutes before a reversed scan was performed, scanning from 85 to 5 °C with the same rate. The denaturing transition was monitored at 260 nm. Baseline corrected data was then smoothed over 5 points and its first derivative was plotted in Microsoft Excel. The final melting temperature was obtained by averaging the melting temperatures in the 5 repeated forward scans.

##### 9.1. UV-Vis melting of ON15-g$^6$A-3S and ON16-nm$^5$U

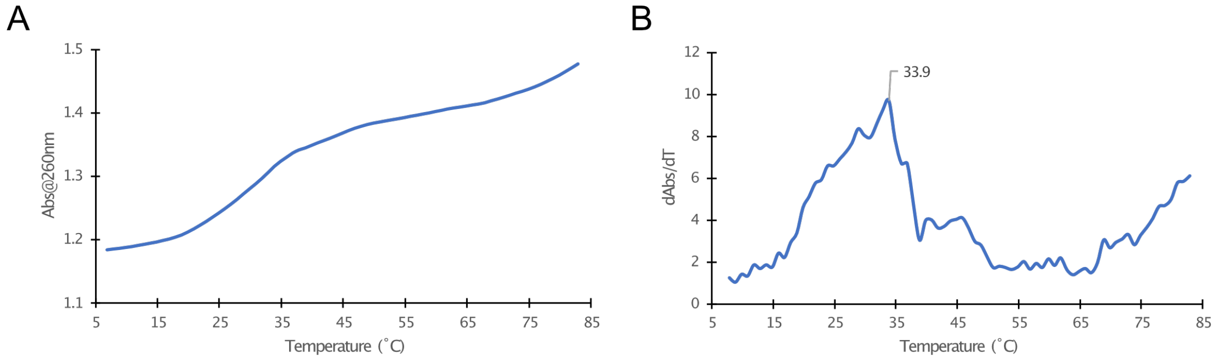


**Figure S46.** (A) The UV melting curve of the 5 basepairs dsRNA between ON15-g$^6$A-3S and ON16-nm$^5$U, monitored by the 260 nm absorption. (B) The derivative plot from the melting curve in (A) indicating the measured melting temperature of the dsRNA is 33.9 ˚C.

#### 10. General information of coupling reactions between *aa*$^6$A and nm$^5$U ONs

Stock solutions of MES pH=6.0 buffer (400 mM), NaCl (1 M) and DMTMM•Cl solution (500 mM) were prepared in water in advance. For setting up a reaction, equimolar of *aa*$^6$A and nm$^5$U containing ONs (1 nmol each) were mixed in water and dried in a SpeedVac concentrator. Subsequently, the oligos were re-dissolved in a 2-fold concentrated reaction buffer (i.e. 200 mM NaCl, 200 mM MES pH=6.0 buffer), with or without 10 mM Egg PC liposome suspension (see above section). The reaction was then diluted to a final concentration of 100 mM NaCl, 100 mM MES pH=6.0 and 10 µM dsRNA, and 50 mM DMTMM•Cl was added from a fresh solution to achieve a final concentration of 50 µM. It was then incubated in a thermomixer at 25°C for 16 h and then analysed by HPLC. Samples with liposome were injected directly to the HPLC without additional treatment.
For the competition experiment, the conditions were the same except that a 1:1:1 mixture of ON17-4a$^6$A, ON17-4l$^6$A and ON18-nm$^5$U-3geS (10 µM each) was employed.

<u>Analysis of coupling reactions by HPLC and MALDI-TOF mass spectrometry</u>

The crudes of the coupling reactions were analyzed by RP-HPLC using an EC 250/4 Nucleodur 100-3 C18ec column from Macherey-Nagel. Buffers: A) 0.1 M AcOH/Et$_3$N in $H_2O$ at pH 7 and

B) 0.1 M AcOH/$Et_3N$ in 80% (v/v) MeCN in $H_2O$. Gradient: unless otherwise stated, 0-80% of B in 45 min. Flow rate = 1 mL·min$^{-1}$. Injection: 100 µL (1 nmol). The yields of the reactions were calculated by integration of the chromatographic peaks of the products and the use of the calibration curves of the corresponding canonical ONs (see 10.1). To simplify the calculations, we assumed that the formed products and the canonical oligonucleotides used for calibration featured identical extinction coefficients, which were calculated for single stranded RNAs. It was expected that double strands and/or secondary structures were disrupted under the HPLC conditions used.

Relative yields of coupling reactions on liposome surface were calculated as follow:

$$Relative\ yield = \frac{[Peak\ area\ of\ RNA\ peptide\ hairpin\ in + Lip]}{[Peak\ area\ of\ RNA\ peptide\ hairpin\ in - Lip]} \times 100\%$$

The the isolated products were desalted with a Sep-Pak tC18 1 cc Vac Cartridge (Waters) and analysed by MALDI-TOF mass spectrometry.

#### 10.1. HPLC calibrations using canonical oligonucleotides

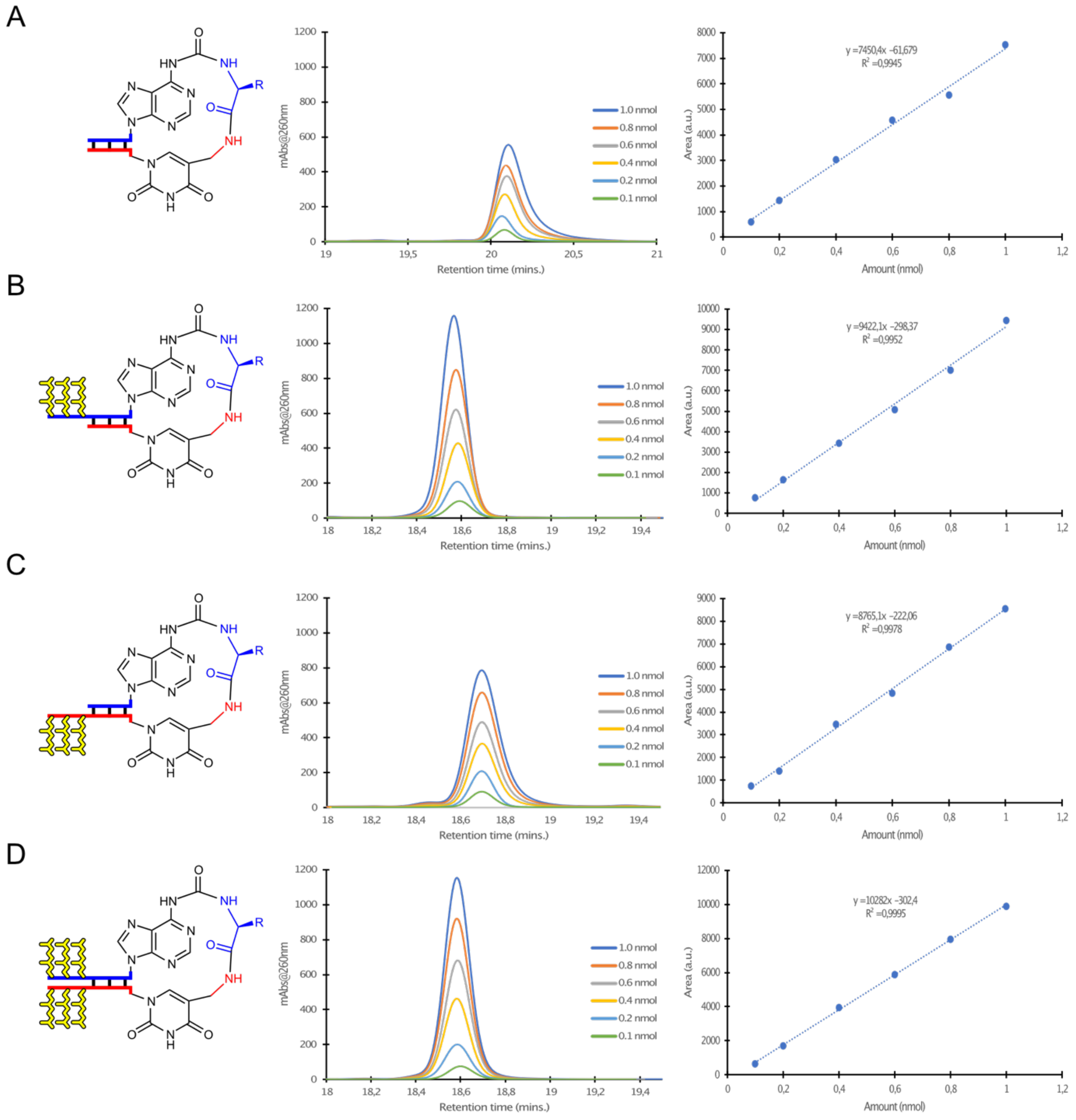


**Figure S47.** Left: The corresponding hairpin-like products that the canonical strands are trying to resemble. Middle: Selected regions of HPL-chromatograms upon the injection of incremental amounts of the corresponding ONs. The peaks are aligned to the 1.0 nmol injection. Right: The linear fit of the area of the peaks on left, the equations are used for quantifying the yield of the peptide coupling reactions. (A) CON-1, (B) CON-2, (C) CON-3 and (D) CON-4.

#### 10.2. HPLCs and MALDI-TOFs of RNA peptide coupling reactions

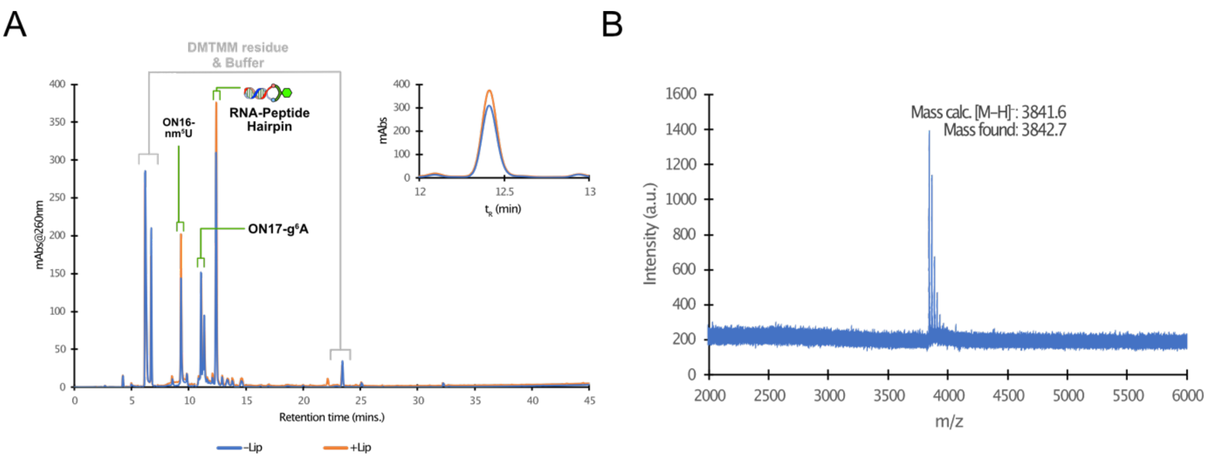


**Figure S48.** (A) Overlay of HPLCgrams between the aqueous solution (blue) and liposome suspension (orange) peptide coupling setups between ON16-nm$^5$U and ON17-g$^6$A. A small window shows the magnified region of the RNA-peptide hairpin formed. (B) MALDI-TOF mass spectrum of the purified RNA-peptide hairpin.

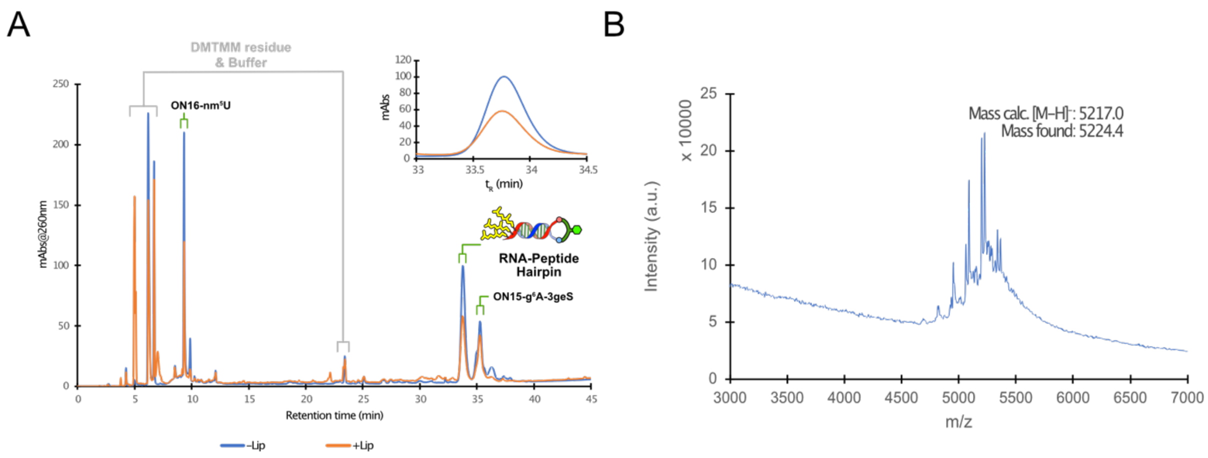


**Figure S49.** (A) Overlay of HPLCgrams between the aqueous solution (blue) and liposome suspension (orange) peptide coupling setups between ON15-g$^6$A-3geS and ON16-nm$^5$U. A small window shows the magnified region of the RNA-peptide hairpin formed. (B) MALDI-TOF mass spectrum of the purified RNA-peptide hairpin.

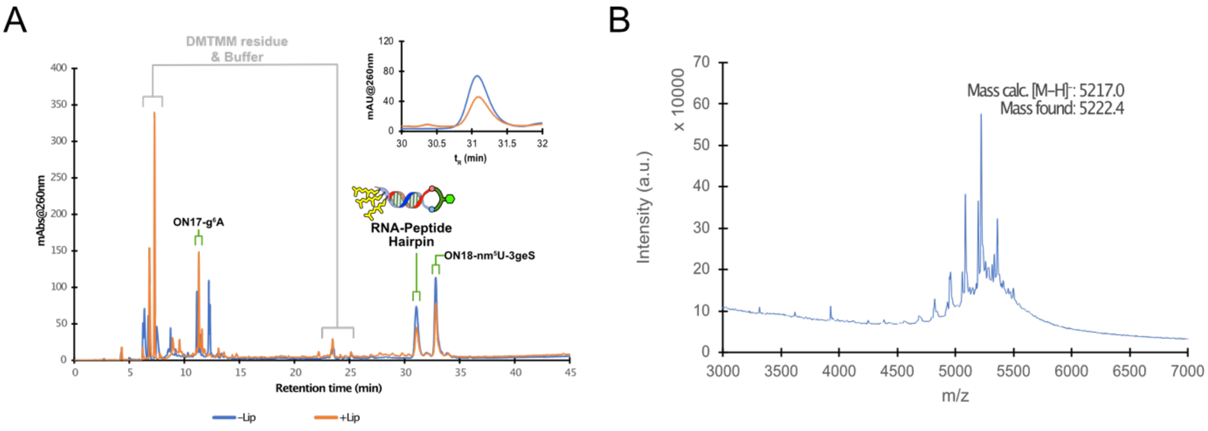


**Figure S50.** (A) Overlay of HPLCgrams between the aqueous solution (blue) and liposome suspension (orange) peptide coupling setups between ON17-g$^6$A and ON18-nm$^5$U-3geS. A small window shows the magnified region of the RNA-peptide hairpin formed. (B) MALDI-TOF mass spectrum of the purified RNA-peptide hairpin.

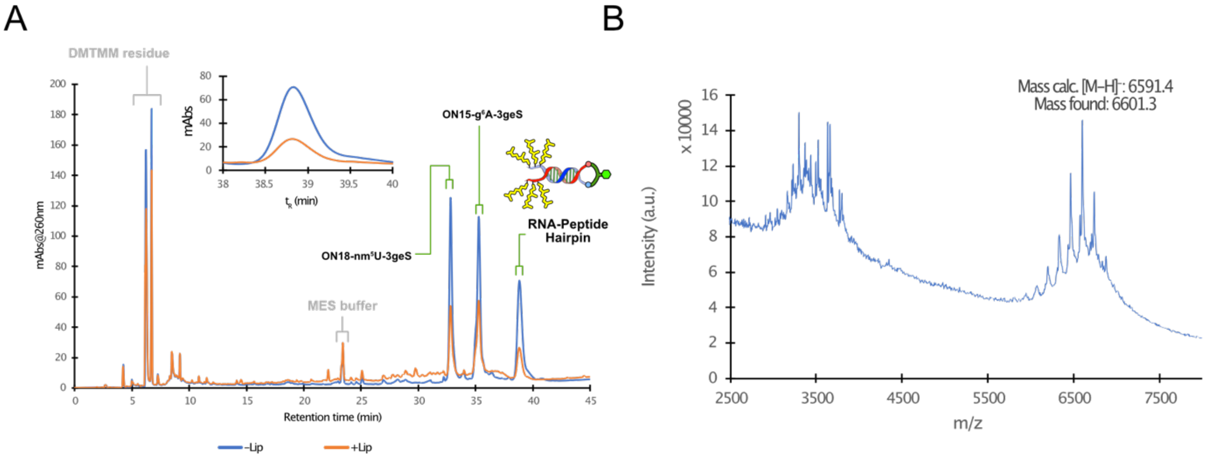


**Figure S51.** (A) Overlay of HPLCgrams between the aqueous solution (blue) and liposome suspension (orange) peptide coupling setups between ON15-g$^6$A-3geS and ON18-nm$^5$U-3geS. A small window shows the magnified region of the RNA-peptide hairpin formed. (B) MALDI-TOF mass spectrum of the purified RNA-peptide hairpin.

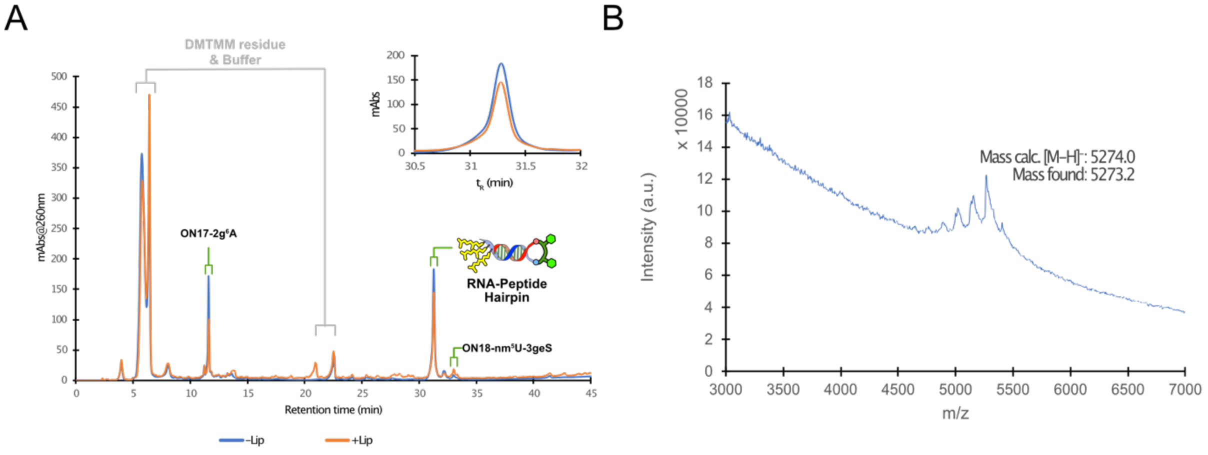


**Figure S52.** (A) Overlay of HPLCgrams between the aqueous solution (blue) and liposome suspension (orange) peptide coupling setups between ON17-2g$^6$A and ON18-nm$^5$U-3geS. A small window shows the magnified region of the RNA-peptide hairpin formed. (B) MALDI-TOF mass spectrum of the purified RNA-peptide hairpin.

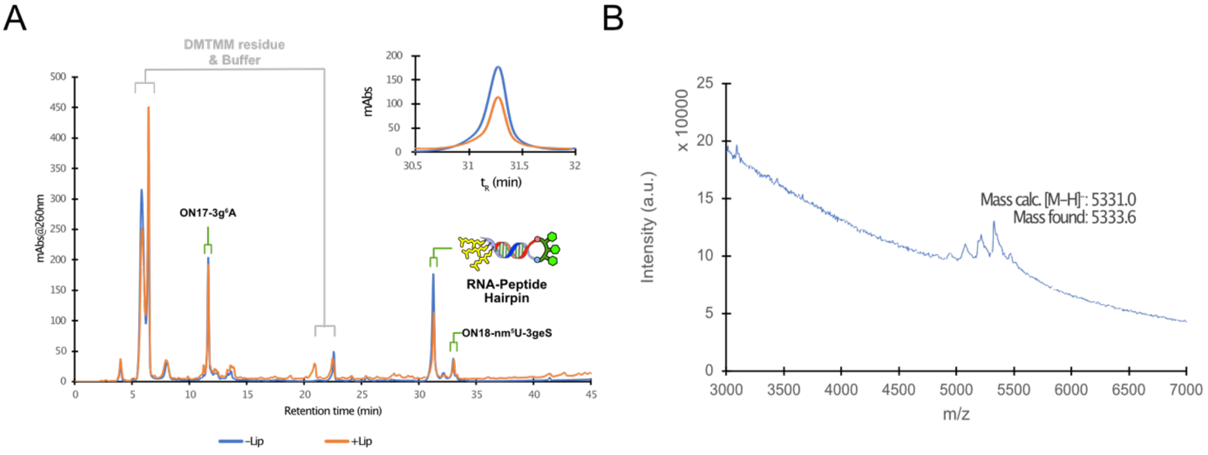


**Figure S53.** (A) Overlay of HPLCgrams between the aqueous solution (blue) and liposome suspension (orange) peptide coupling setups between ON17-3g$^6$A and ON18-nm$^5$U-3geS. A small window shows the magnified region of the RNA-peptide hairpin formed. (B) MALDI-TOF mass spectrum of the purified RNA-peptide hairpin.

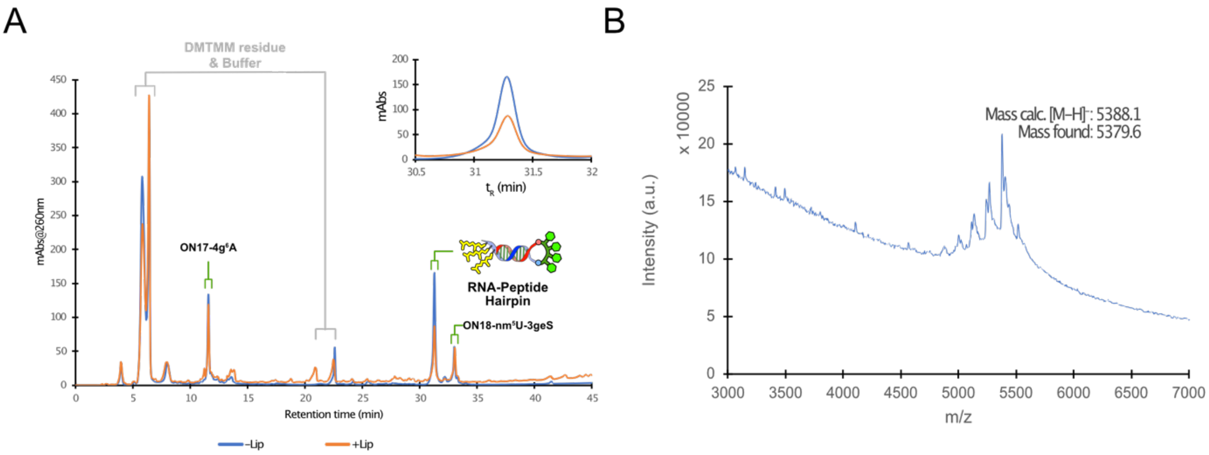


**Figure S54.** (A) Overlay of HPLCgrams between the aqueous solution (blue) and liposome suspension (orange) peptide coupling setups between ON17-4g$^6$A and ON18-nm$^5$U-3geS. A small window shows the magnified region of the RNA-peptide hairpin formed. (B) MALDI-TOF mass spectrum of the purified RNA-peptide hairpin.

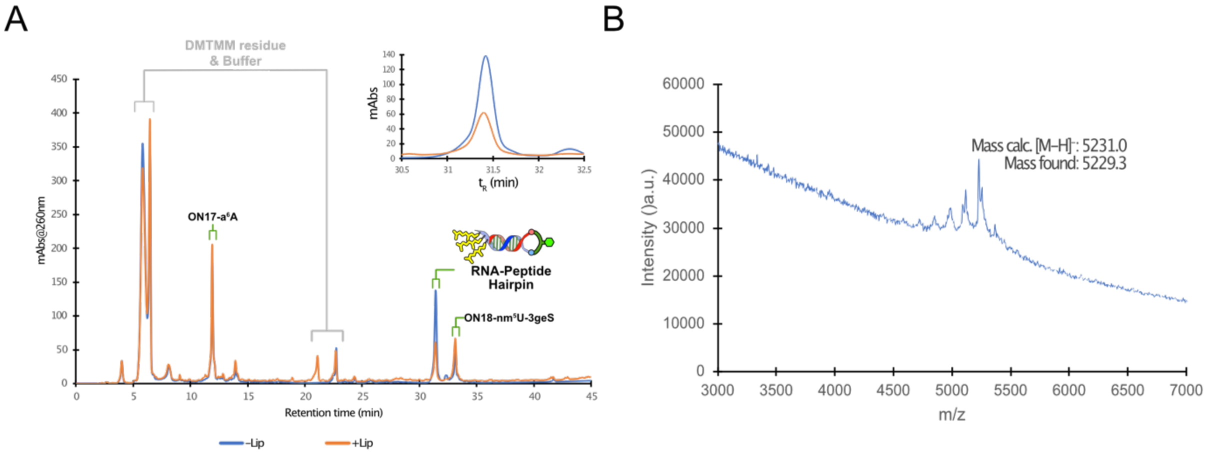


**Figure S55.** (A) Overlay of HPLCgrams between the aqueous solution (blue) and liposome suspension (orange) peptide coupling setups between ON17-a$^6$A and ON18-nm$^5$U-3geS. A small window shows the magnified region of the RNA-peptide hairpin formed. (B) MALDI-TOF mass spectrum of the purified RNA-peptide hairpin.

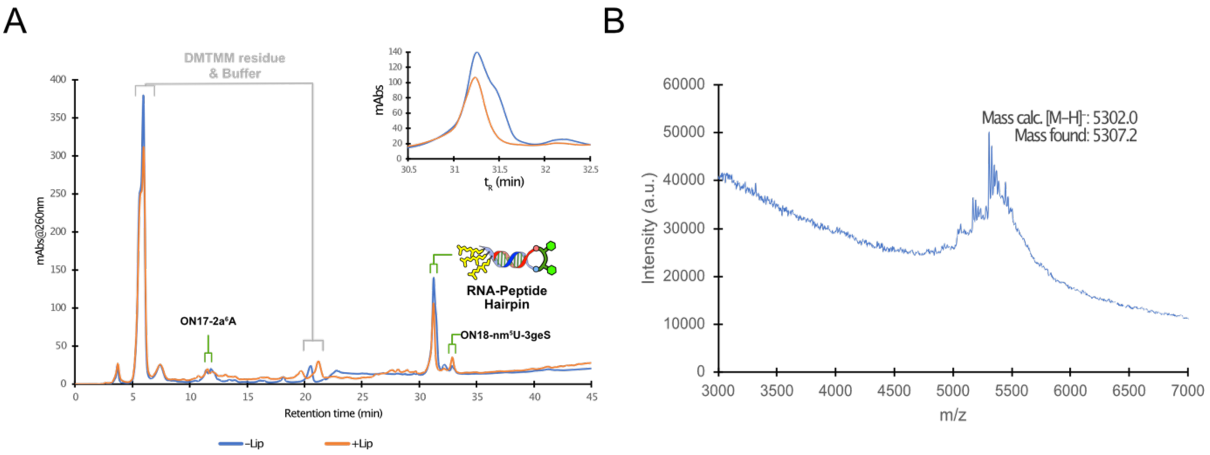


**Figure S56.** (A) Overlay of HPLCgrams between the aqueous solution (blue) and liposome suspension (orange) peptide coupling setups between ON17-2a$^6$A and ON18-nm$^5$U-3geS. A small window shows the magnified region of the RNA-peptide hairpin formed. (B) MALDI-TOF mass spectrum of the purified RNA-peptide hairpin.

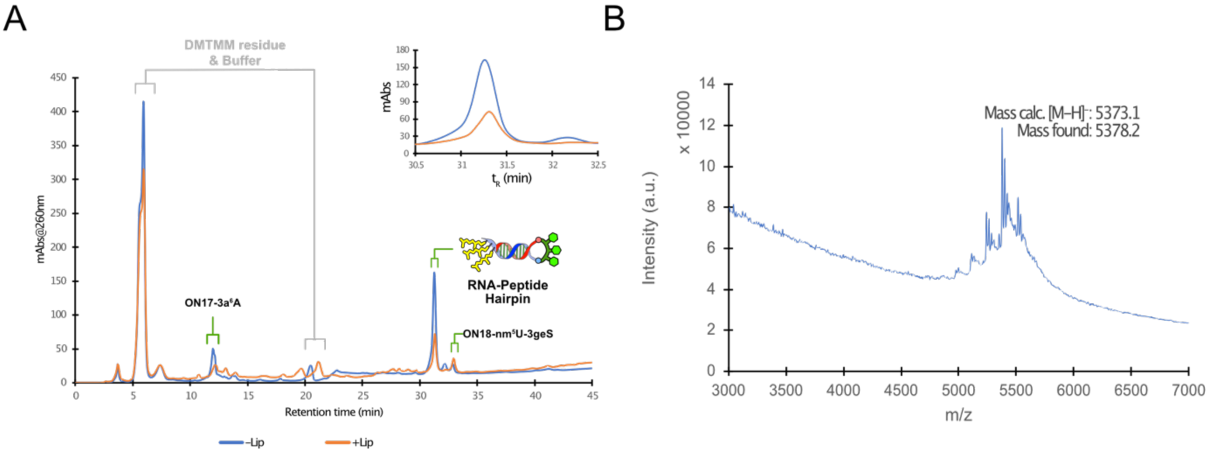


**Figure S57.** (A) Overlay of HPLCgrams between the aqueous solution (blue) and liposome suspension (orange) peptide coupling setups between ON17-3a$^6$A and ON18-nm$^5$U-3geS. A small window shows the magnified region of the RNA-peptide hairpin formed. (B) MALDI-TOF mass spectrum of the purified RNA-peptide hairpin.

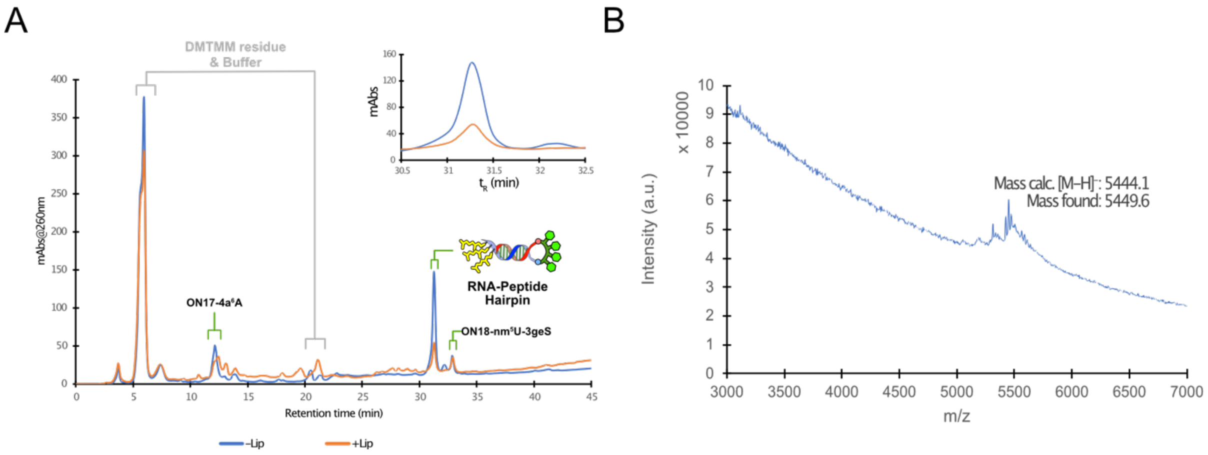


**Figure S58.** (A) Overlay of HPLCgrams between the aqueous solution (blue) and liposome suspension (orange) peptide coupling setups between ON17-4a$^6$A and ON18-nm$^5$U-3geS. A small window shows the magnified region of the RNA-peptide hairpin formed. (B) MALDI-TOF mass spectrum of the purified RNA-peptide hairpin.

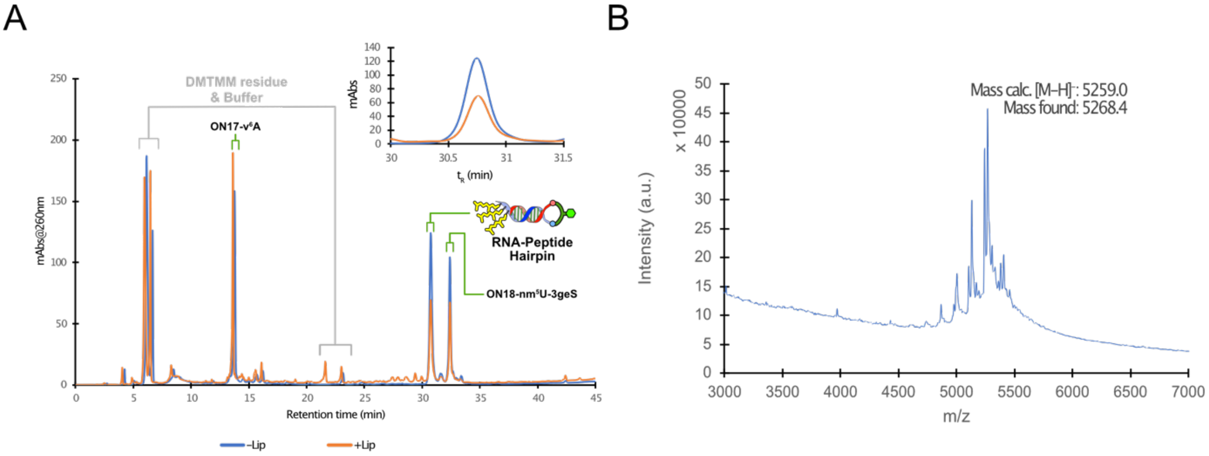


**Figure S59.** (A) Overlay of HPLCgrams between the aqueous solution (blue) and liposome suspension (orange) peptide coupling setups between ON17-v$^6$A and ON18-nm$^5$U-3geS. A small window shows the magnified region of the RNA-peptide hairpin formed. (B) MALDI-TOF mass spectrum of the purified RNA-peptide hairpin.

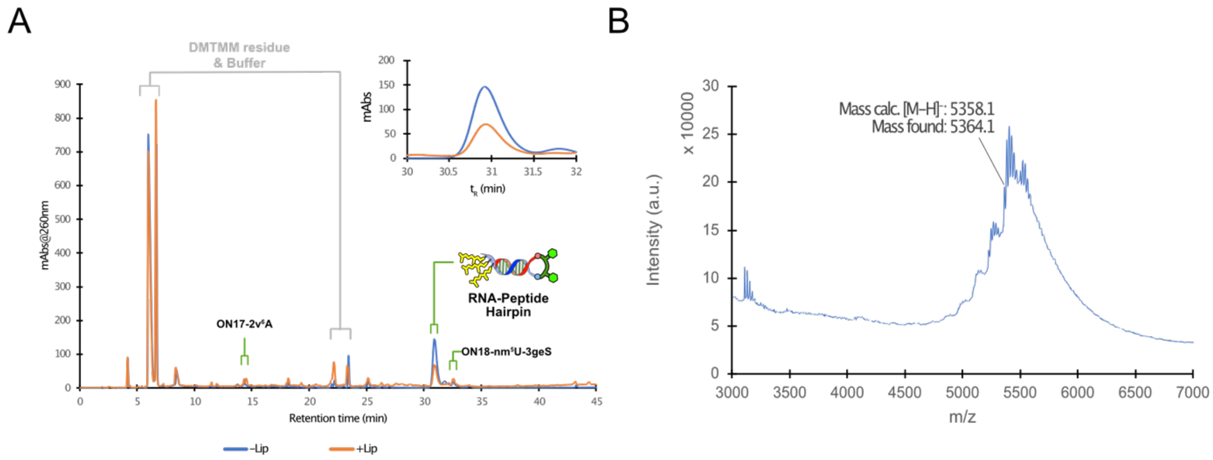


**Figure S60.** (A) Overlay of HPLCgrams between the aqueous solution (blue) and liposome suspension (orange) peptide coupling setups between ON17-2v$^6$A and ON18-nm$^5$U-3geS. A small window shows the magnified region of the RNA-peptide hairpin formed. (B) MALDI-TOF mass spectrum of the purified RNA-peptide hairpin.

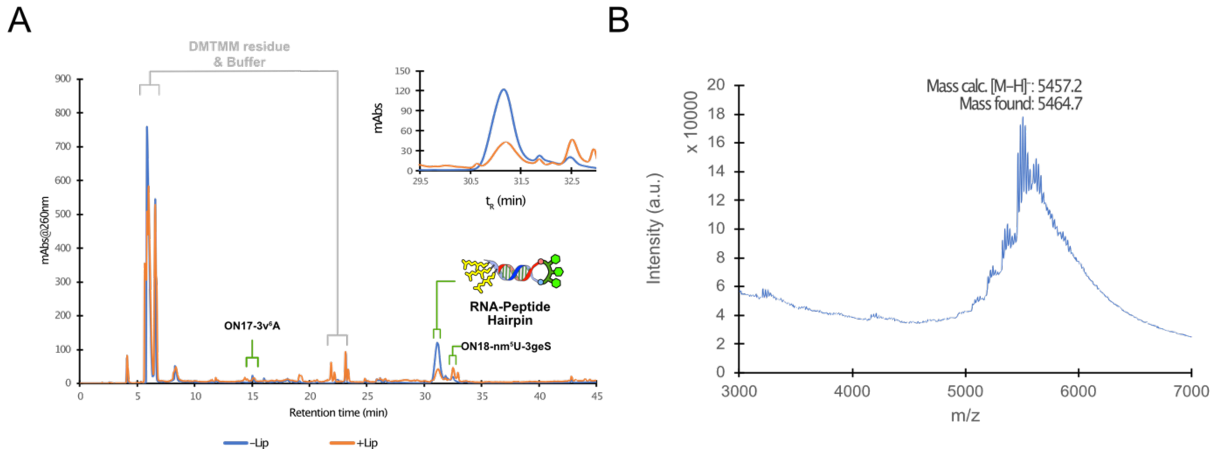


**Figure S61.** (A) Overlay of HPLCgrams between the aqueous solution (blue) and liposome suspension (orange) peptide coupling setups between ON17-3v$^6$A and ON18-nm$^5$U-3geS. A small window shows the magnified region of the RNA-peptide hairpin formed. (B) MALDI-TOF mass spectrum of the purified RNA-peptide hairpin.

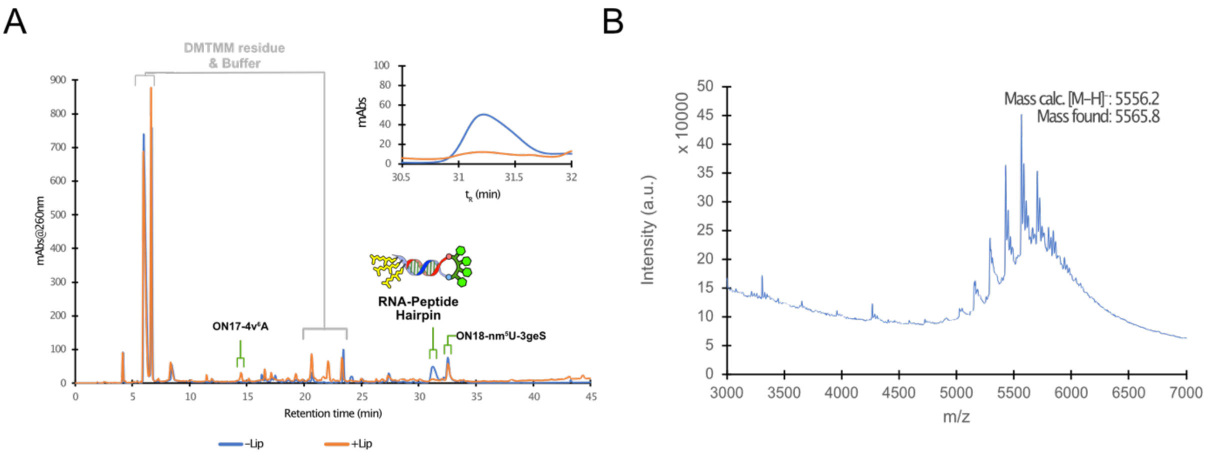


**Figure S62.** (A) Overlay of HPLCgrams between the aqueous solution (blue) and liposome suspension (orange) peptide coupling setups between ON17-4v$^6$A and ON18-nm$^5$U-3geS. A small window shows the magnified region of the RNA-peptide hairpin formed. (B) MALDI-TOF mass spectrum of the purified RNA-peptide hairpin.

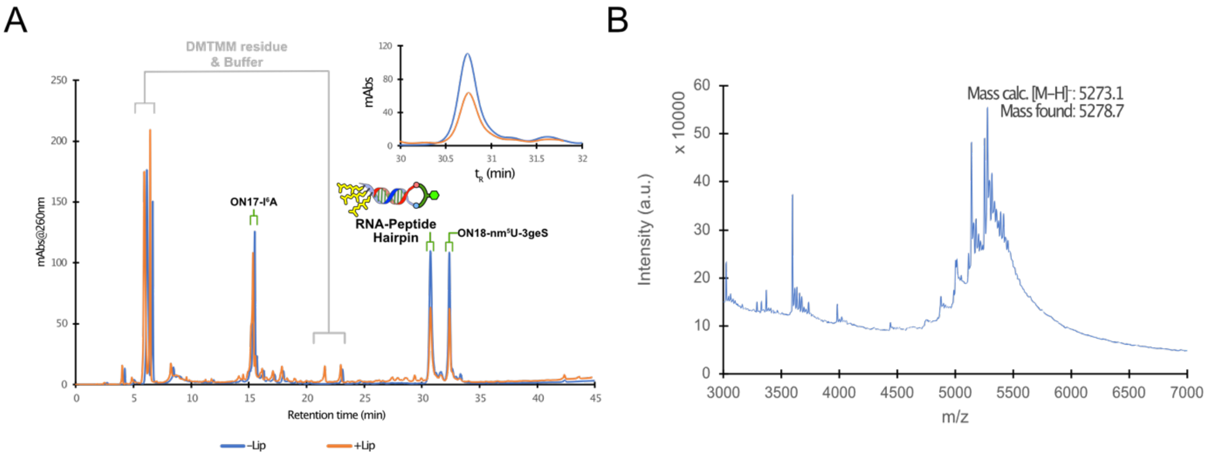


**Figure S63.** (A) Overlay of HPLCgrams between the aqueous solution (blue) and liposome suspension (orange) peptide coupling setups between ON17-I$^6$A and ON18-nm$^5$U-3geS. A small window shows the magnified region of the RNA-peptide hairpin formed. (B) MALDI-TOF mass spectrum of the purified RNA-peptide hairpin.

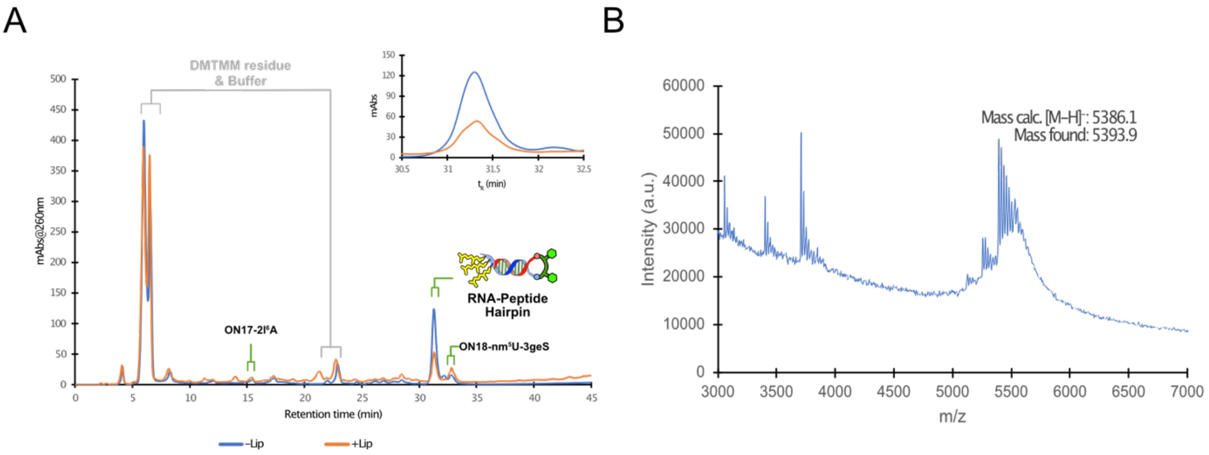


**Figure S64.** (A) Overlay of HPLCgrams between the aqueous solution (blue) and liposome suspension (orange) peptide coupling setups between ON17-2I$^6$A and ON18-nm$^5$U-3geS. A small window shows the magnified region of the RNA-peptide hairpin formed. (B) MALDI-TOF mass spectrum of the purified RNA-peptide hairpin.

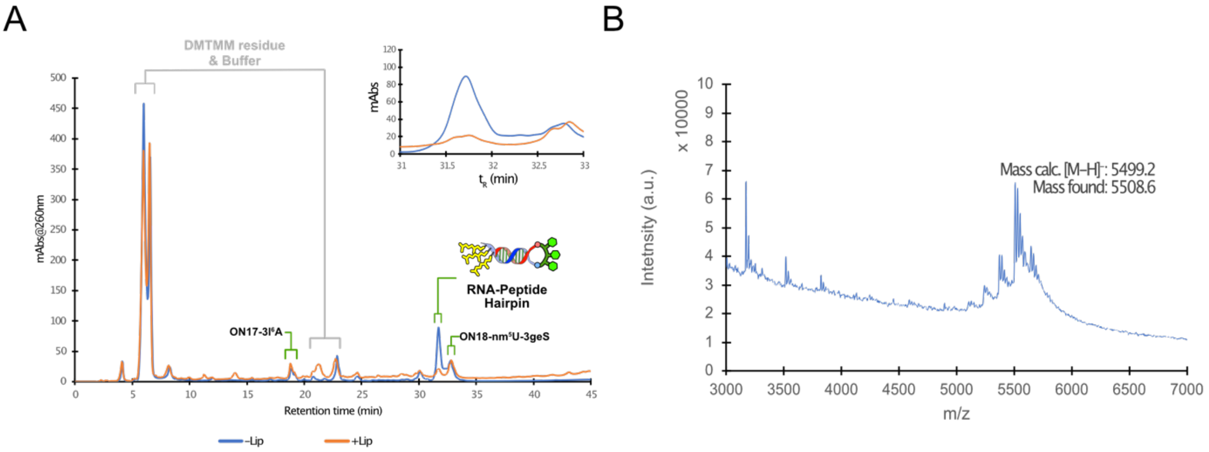


**Figure S65.** (A) Overlay of HPLCgrams between the aqueous solution (blue) and liposome suspension (orange) peptide coupling setups between ON17-3I$^6$A and ON18-nm$^5$U-3geS. A small window shows the magnified region of the RNA-peptide hairpin formed. (B) MALDI-TOF mass spectrum of the purified RNA-peptide hairpin.

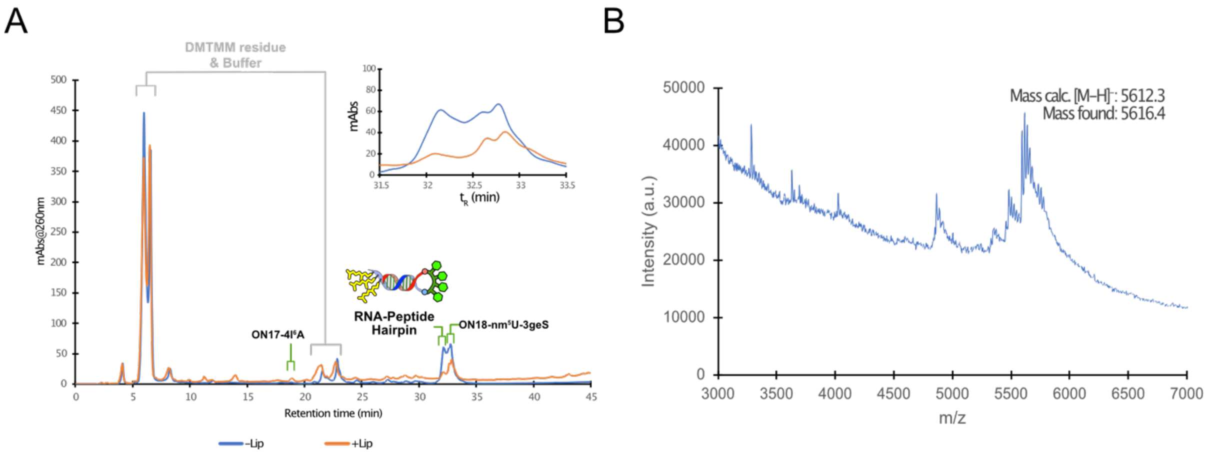


**Figure S66.** (A) Overlay of HPLCgrams between the aqueous solution (blue) and liposome suspension (orange) peptide coupling setups between ON17-4I$^6$A and ON18-nm$^5$U-3geS. A small window shows the magnified region of the RNA-peptide haiprin formed. (B) MALDI-TOF mass spectrum of the purified RNA-peptide hairpin.

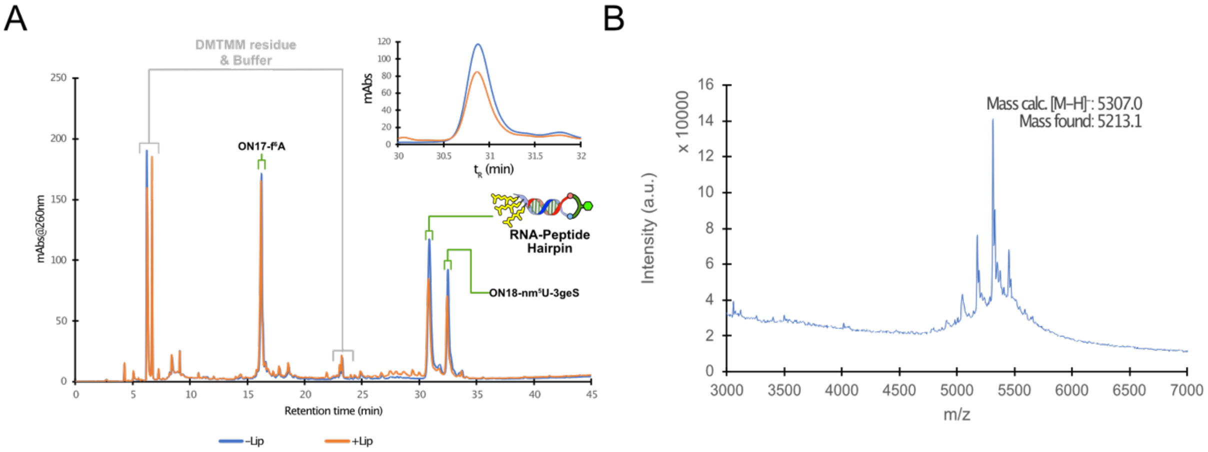


**Figure S67.** (A) Overlay of HPLCgrams between the aqueous solution (blue) and liposome suspension (orange) peptide coupling setups between ON17-f$^6$A and ON18-nm$^5$U-3geS. A small window shows the magnified region of the RNA-peptide haiprin formed. (B) MALDI-TOF mass spectrum of the purified RNA-peptide hairpin.

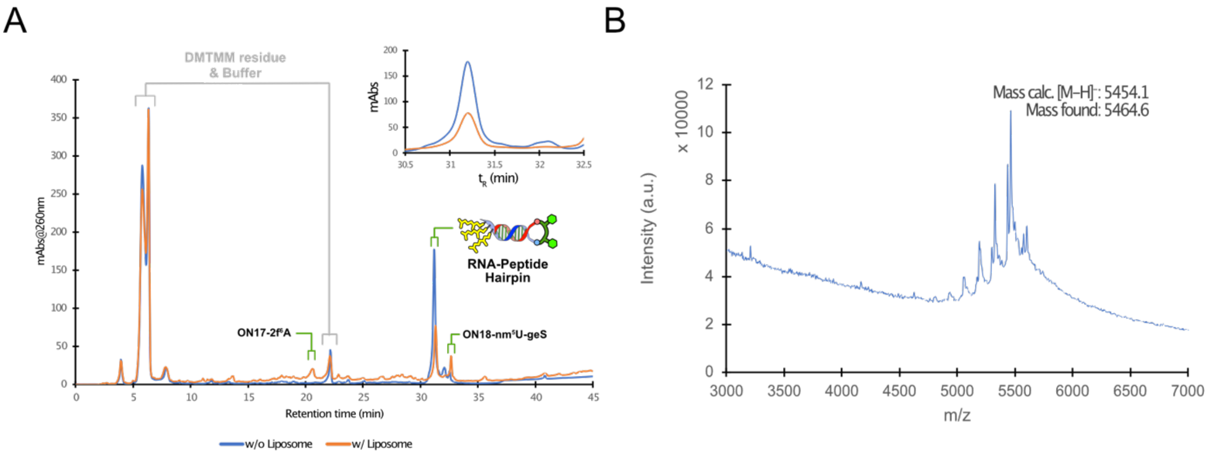


**Figure S68.** (A) Overlay of HPLCgrams between the aqueous solution (blue) and liposome suspension (orange) peptide coupling setups between ON17-2f$^6$A and ON18-nm$^5$U-geS. A small window shows the magnified region of the RNA-peptide hairpin formed. (B) MALDI-TOF mass spectrum of the purified RNA-peptide hairpin.

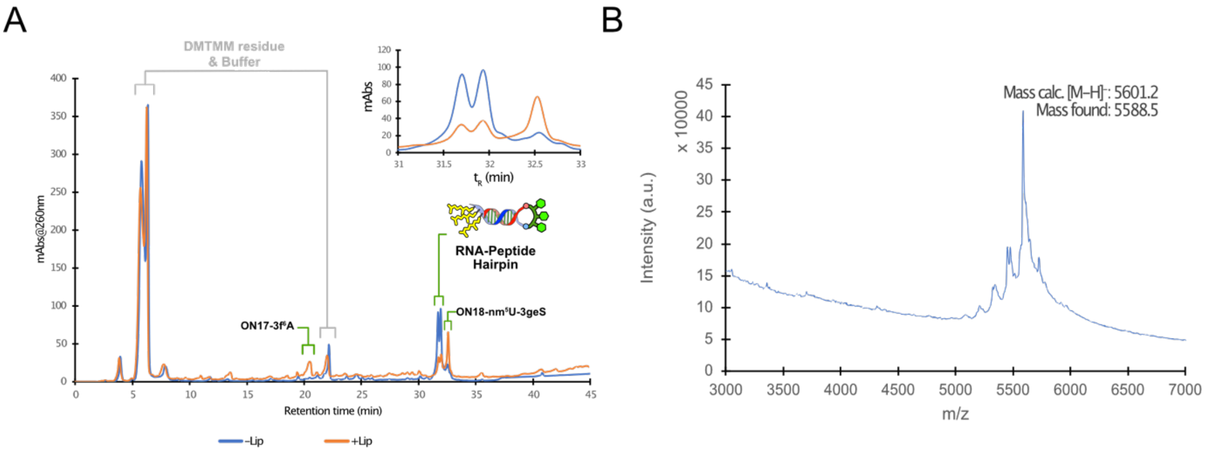


**Figure S69.** (A) Overlay of HPLCgrams between the aqueous solution (blue) and liposome suspension (orange) peptide coupling setups between ON17-3f$^6$A and ON18-nm$^5$U-geS. A small window shows the magnified region of the RNA-peptide hairpin formed. (B) MALDI-TOF mass spectrum of the purified RNA-peptide hairpin.

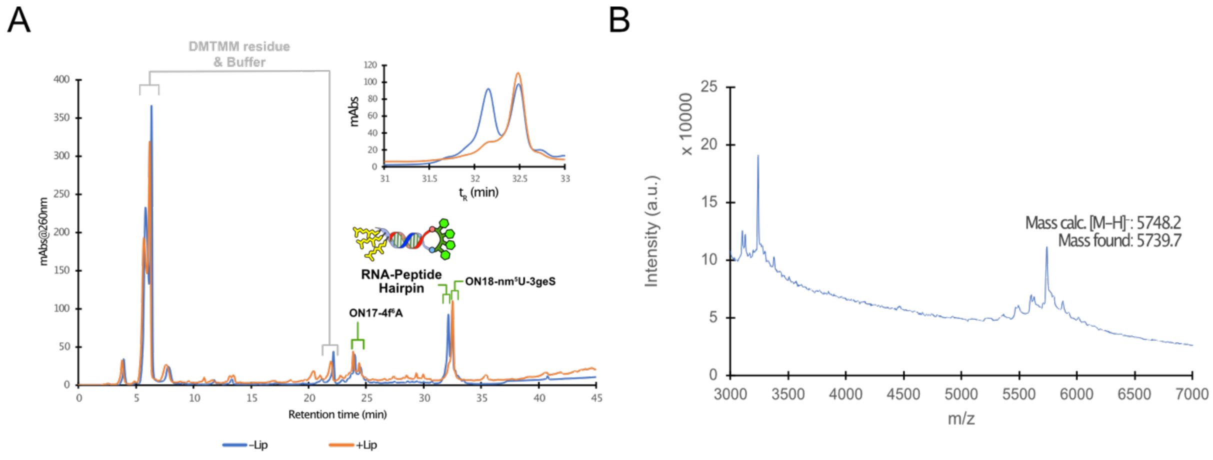


**Figure S70.** (A) Overlay of HPLCgrams between the aqueous solution (blue) and liposome suspension (orange) peptide coupling setups between ON17-4f$^6$A and ON18-nm$^5$U-3geS. A small window shows the magnified region of the RNA-peptide hairpin formed. (B) MALDI-TOF mass spectrum of the purified RNA-peptide hairpin.

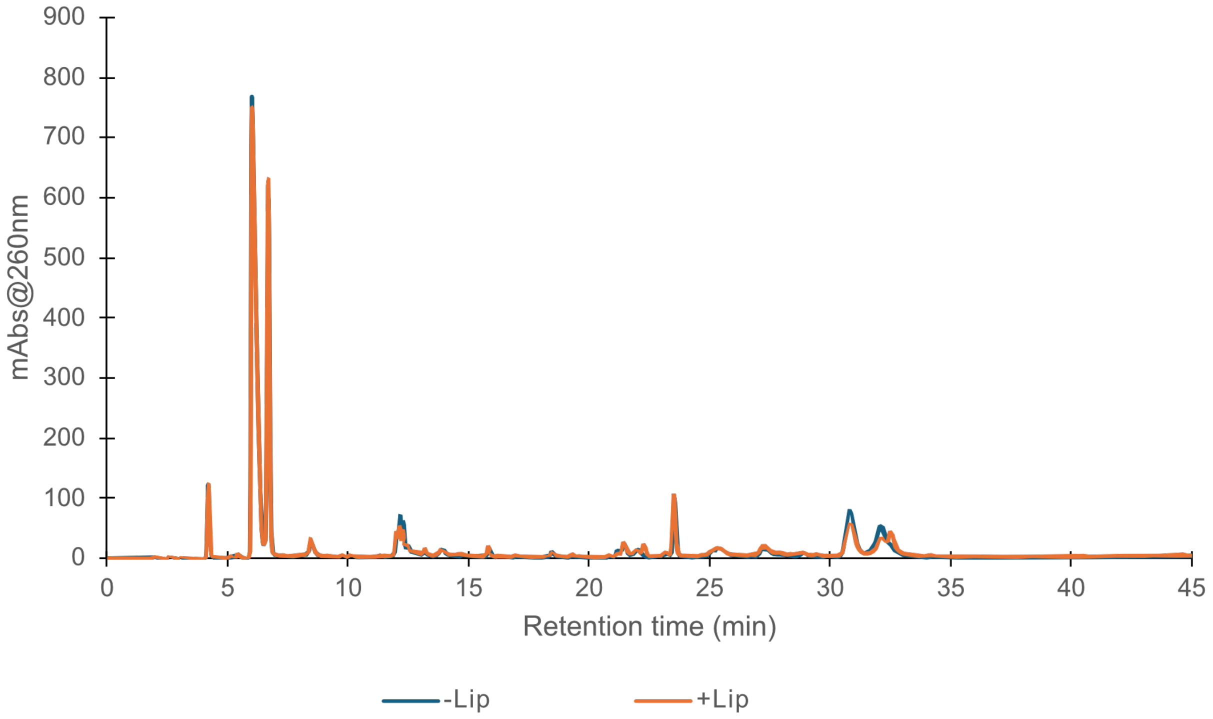


**Figure S71.** Overlay of full HPLCgrams of the competition experiment with ON17-4a$^6$A:ON17-4l$^6$A:ON18-nm$^5$U-3geS, 1:1:1. The yields of RNA-peptides hairpins formed by ON17-4a$^6$A and -4l$^6$A in -Lip were 24.4±0.3% and 19.2±0.4%, respectively. In +Lip, they were 18.3±0% and 11.3±0.1%, respectively.

#### NMR Spectra

##### $^1$H NMR spectrum of compound 2a

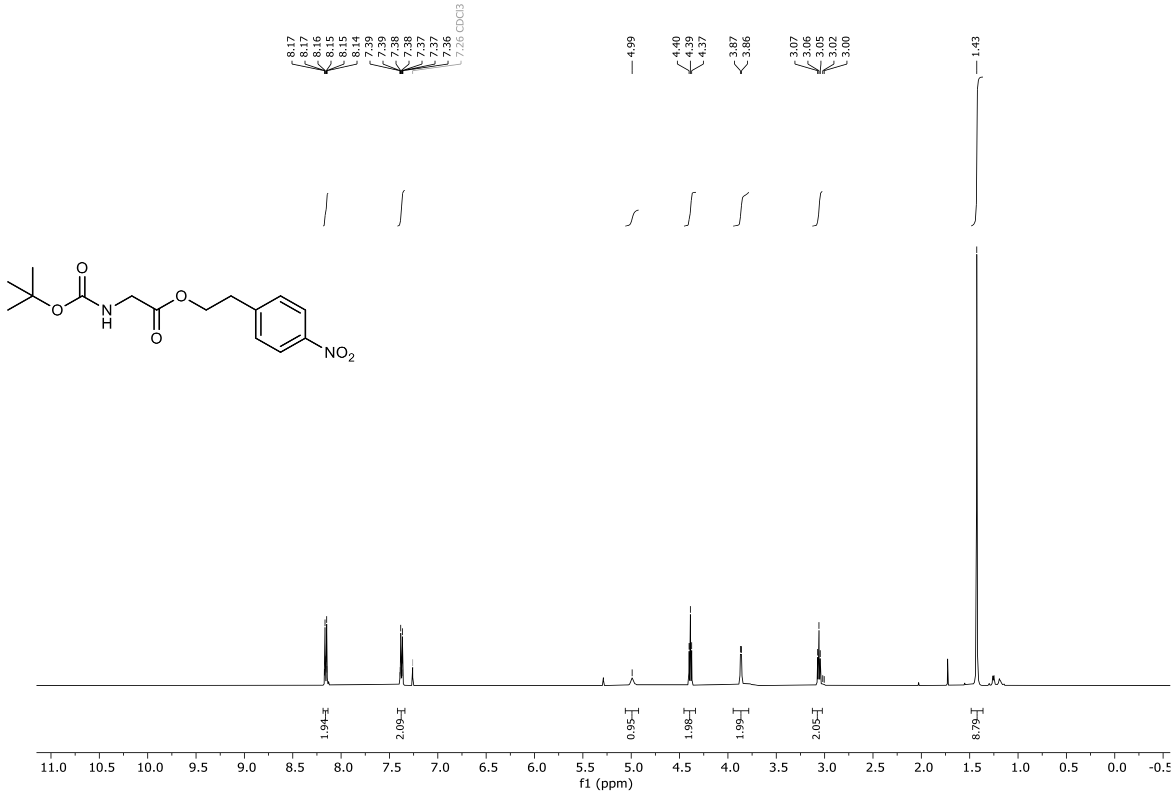


##### $^1$H NMR spectrum of compound 2b

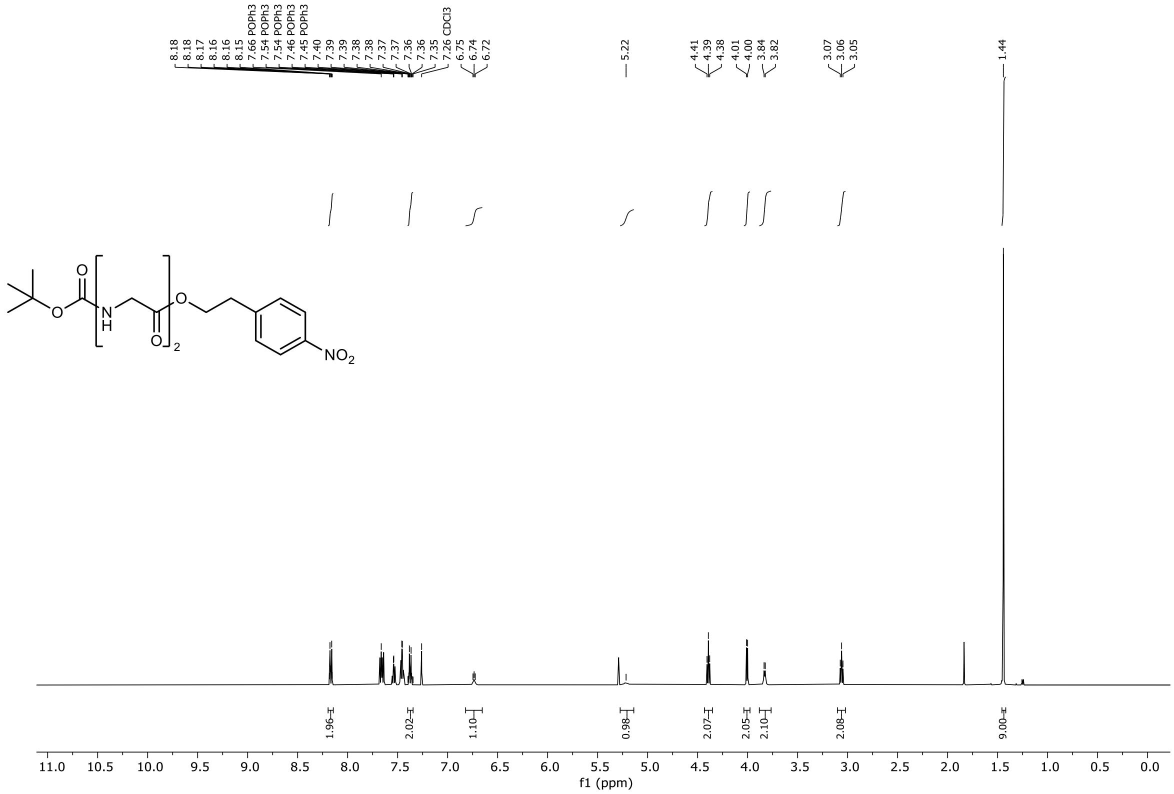

#### $^{13}$C NMR spectrum of compound 2b

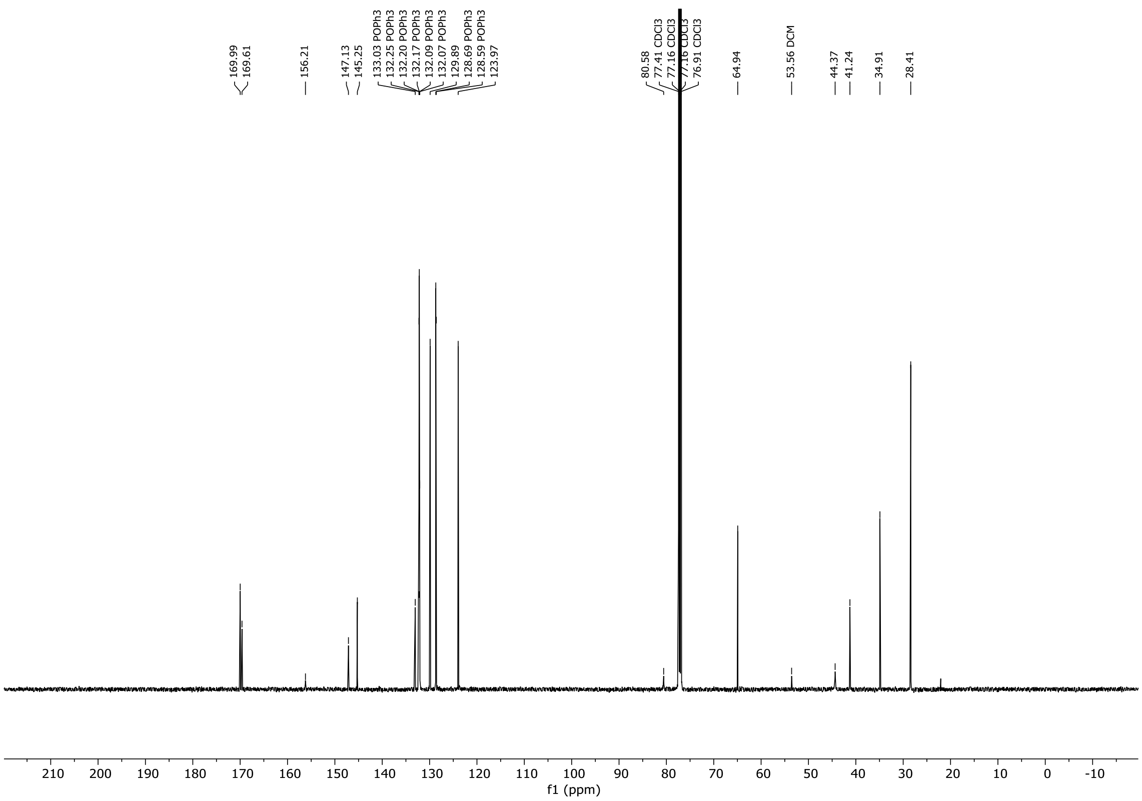

#### $^{1}$H NMR spectrum of compound 2c

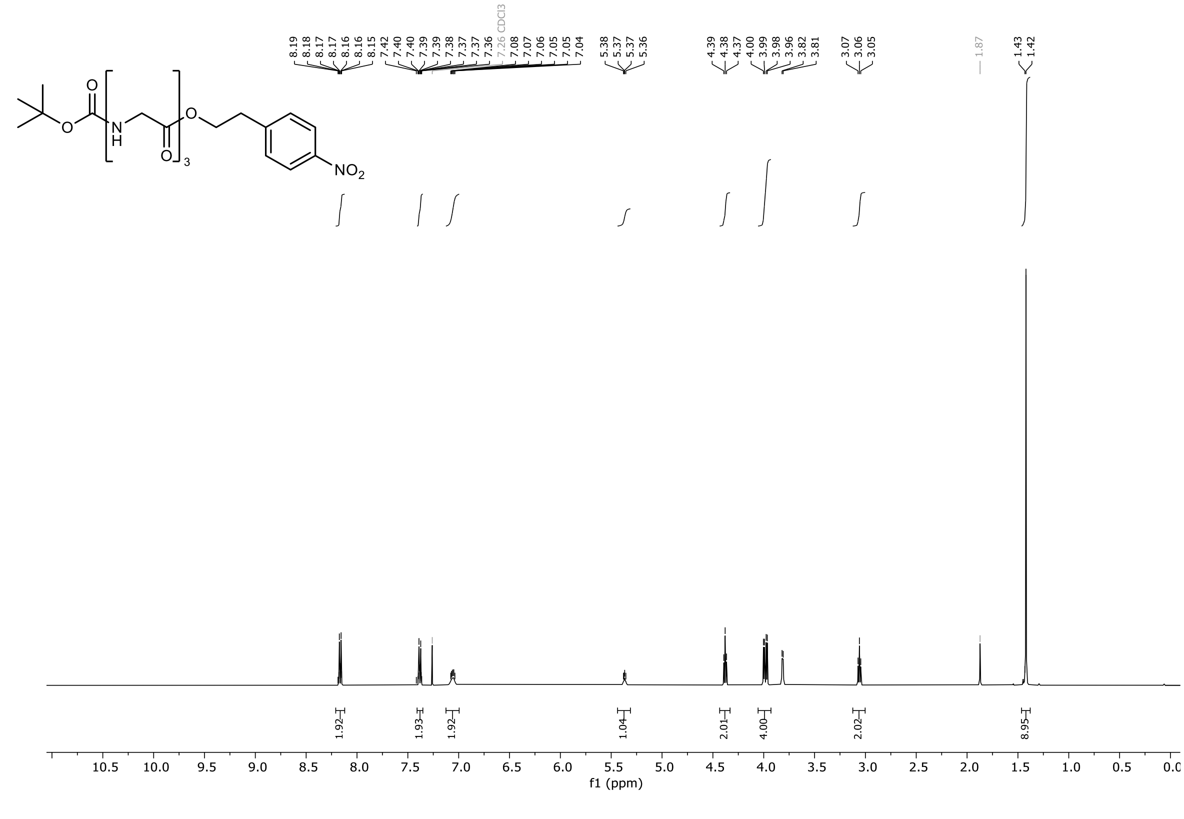

#### ¹³C NMR spectrum of compound 2c

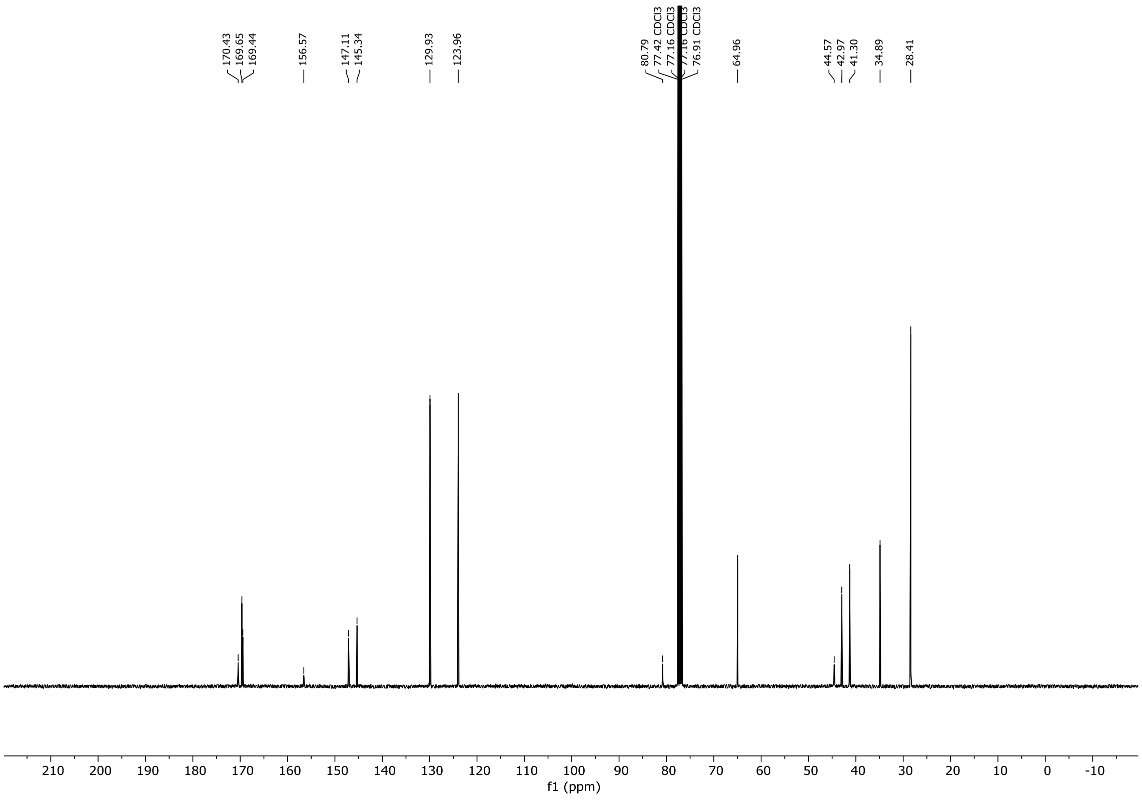


#### ¹H NMR spectrum of compound 2d

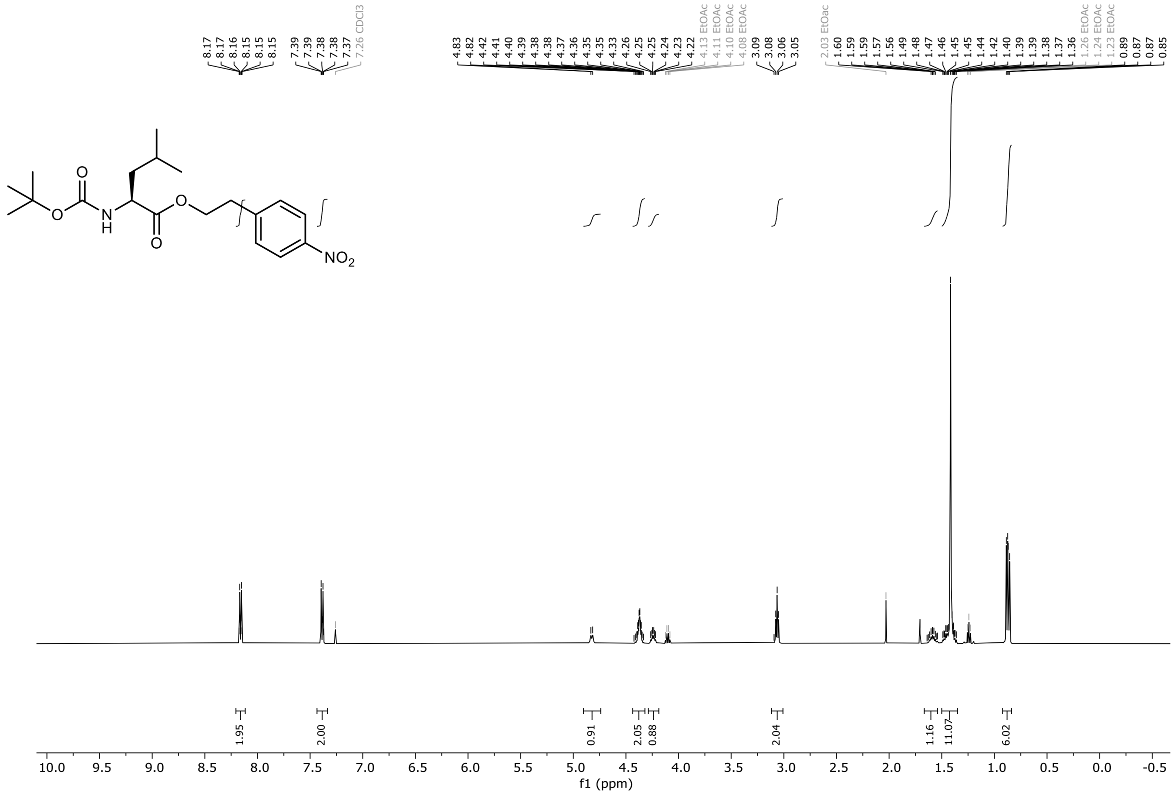

#### $^{13}$C NMR spectrum of compound 2d

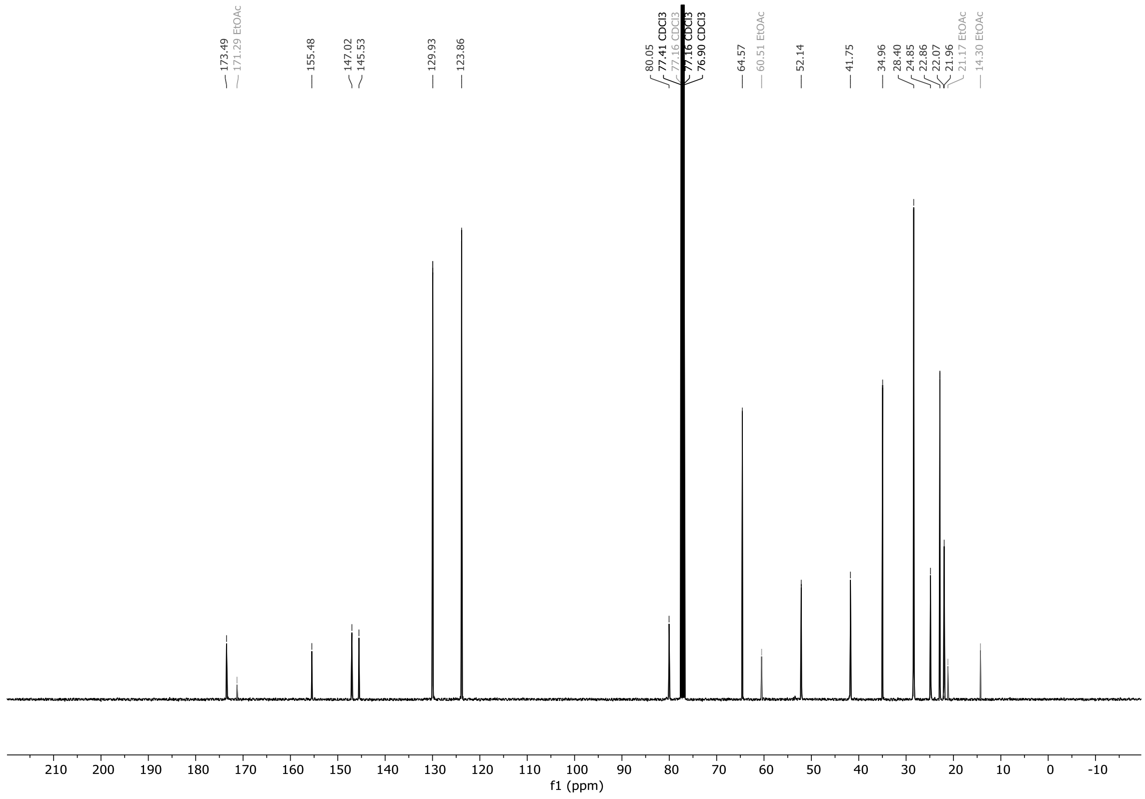


#### $^1$H NMR spectrum of compound 2k

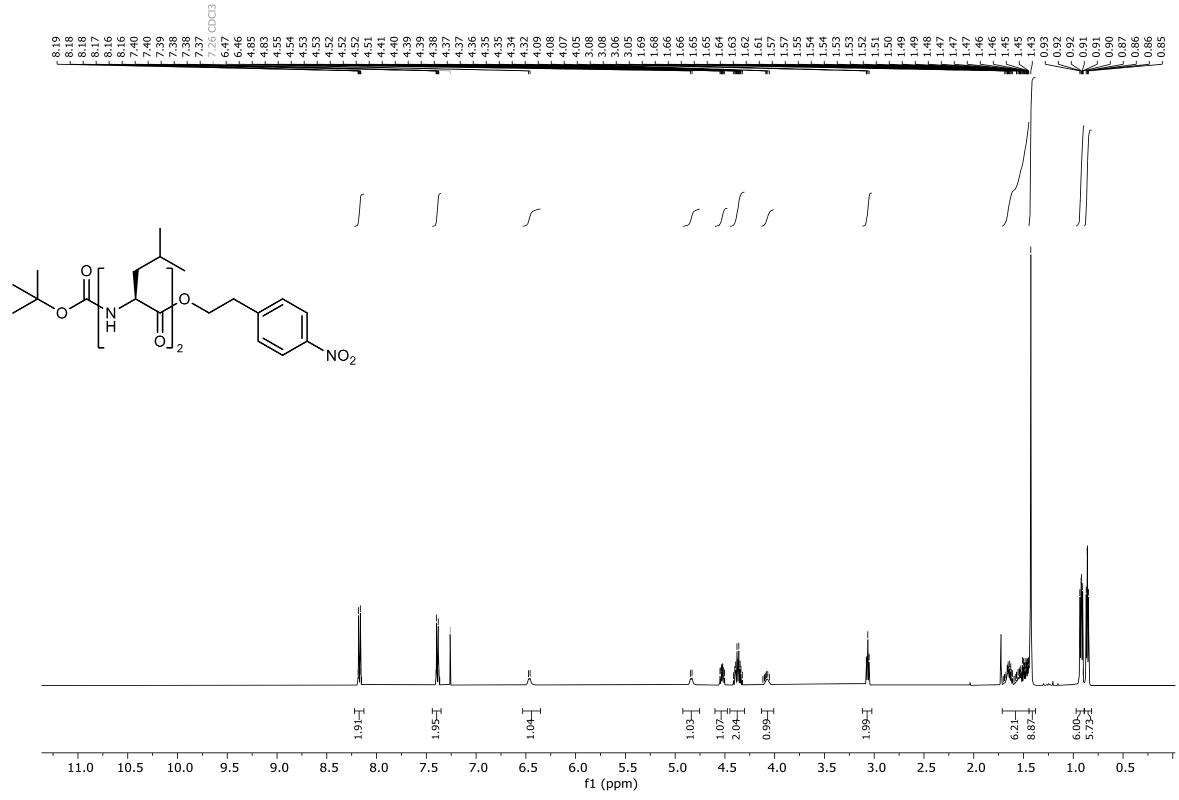

#### $^{13}$C NMR spectrum of compound 2k

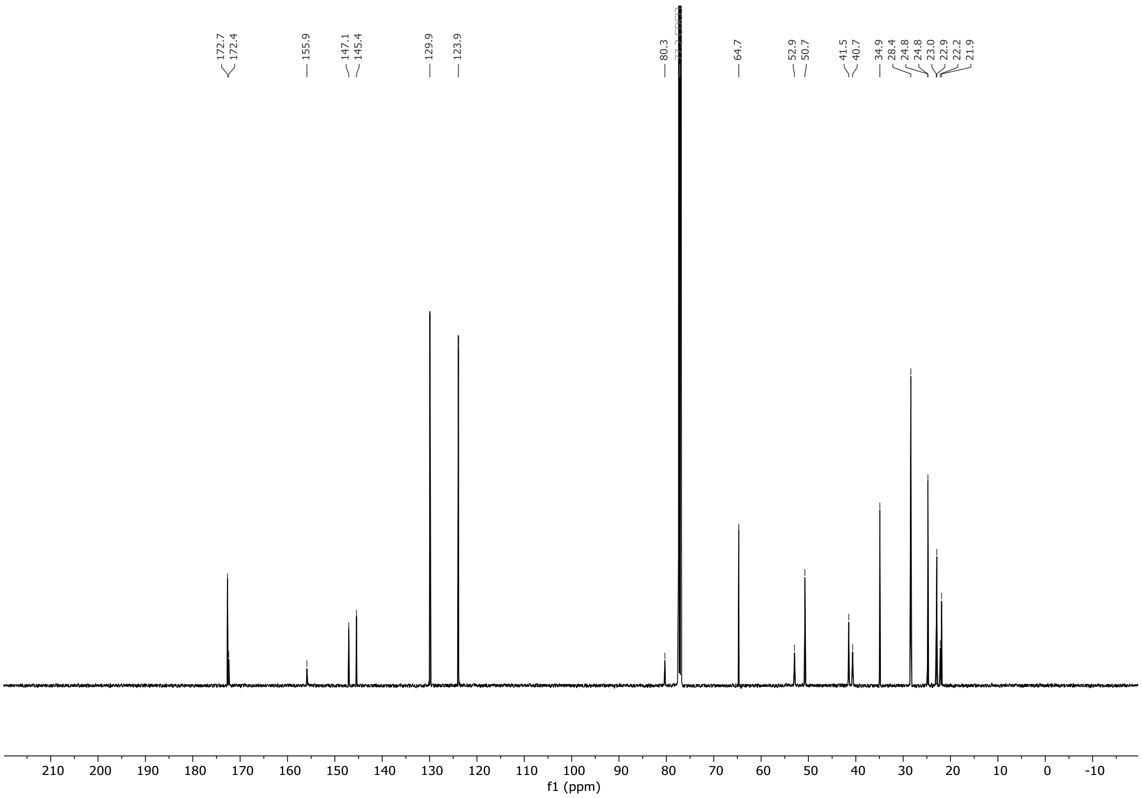

#### $^{1}$H NMR spectrum of compound 2l

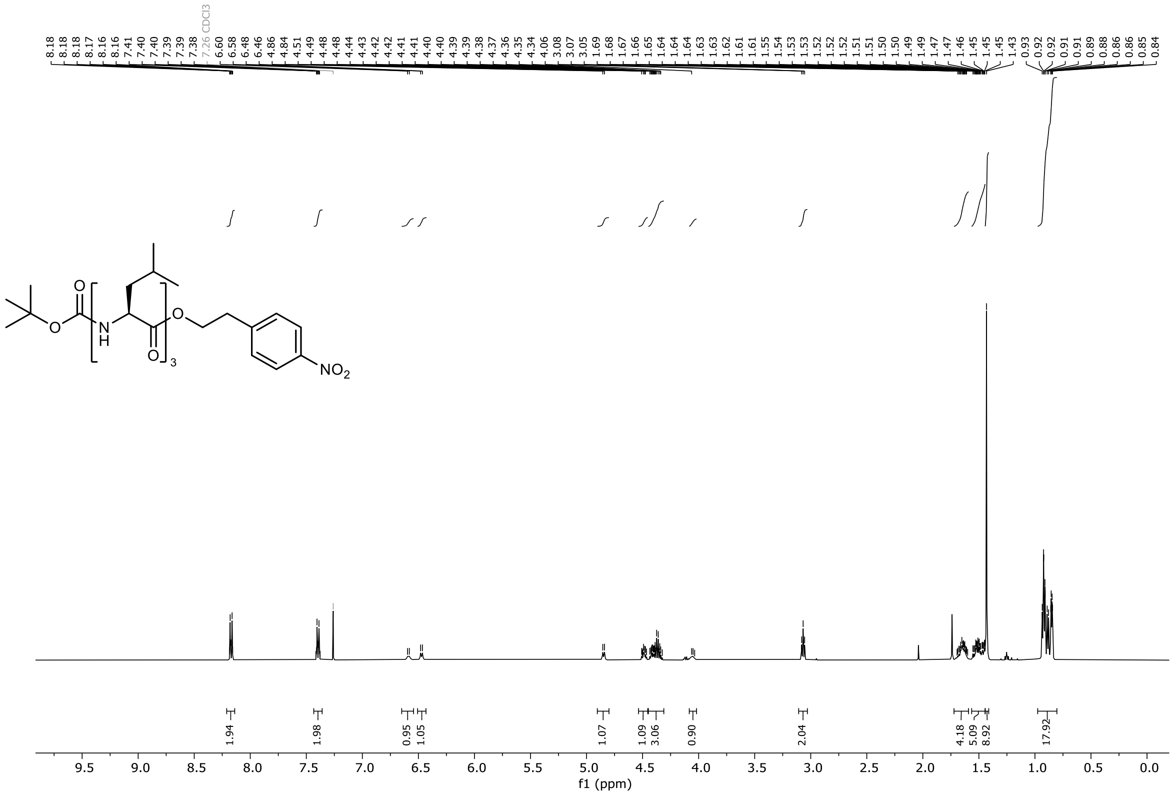

#### $^{13}$C NMR spectrum of compound 2l

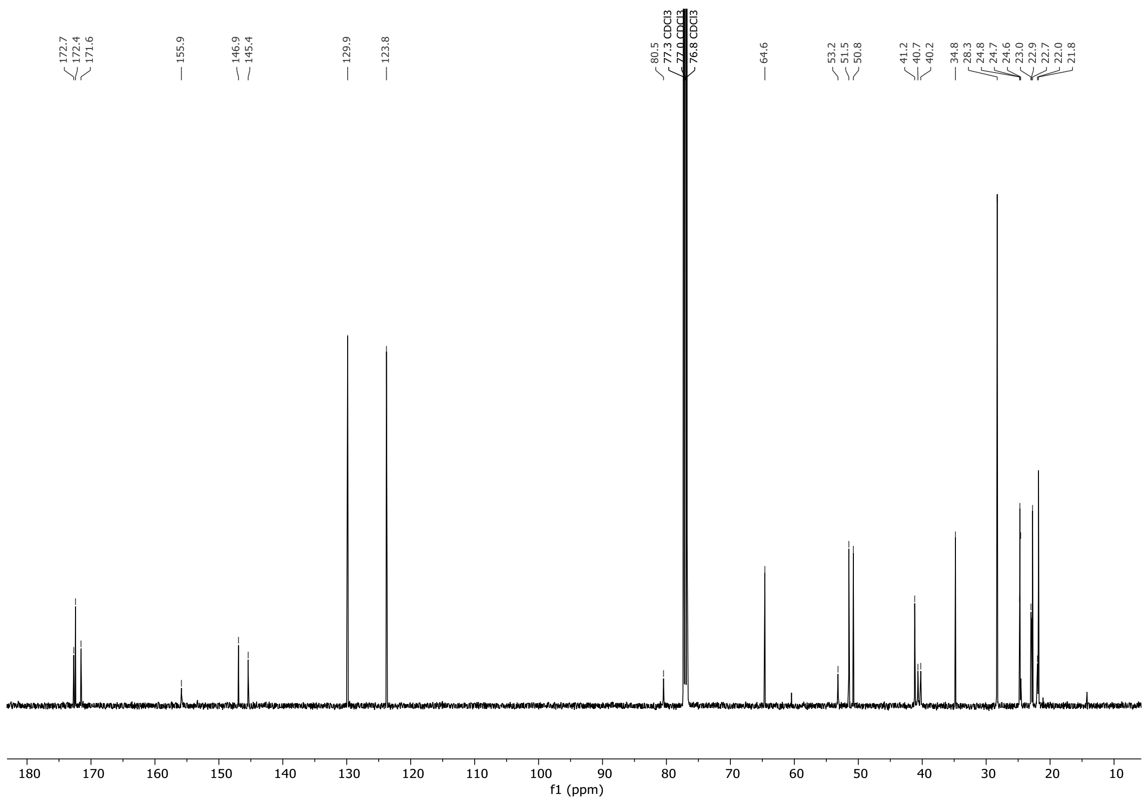

#### $^{1}$H NMR spectrum of compound 2o

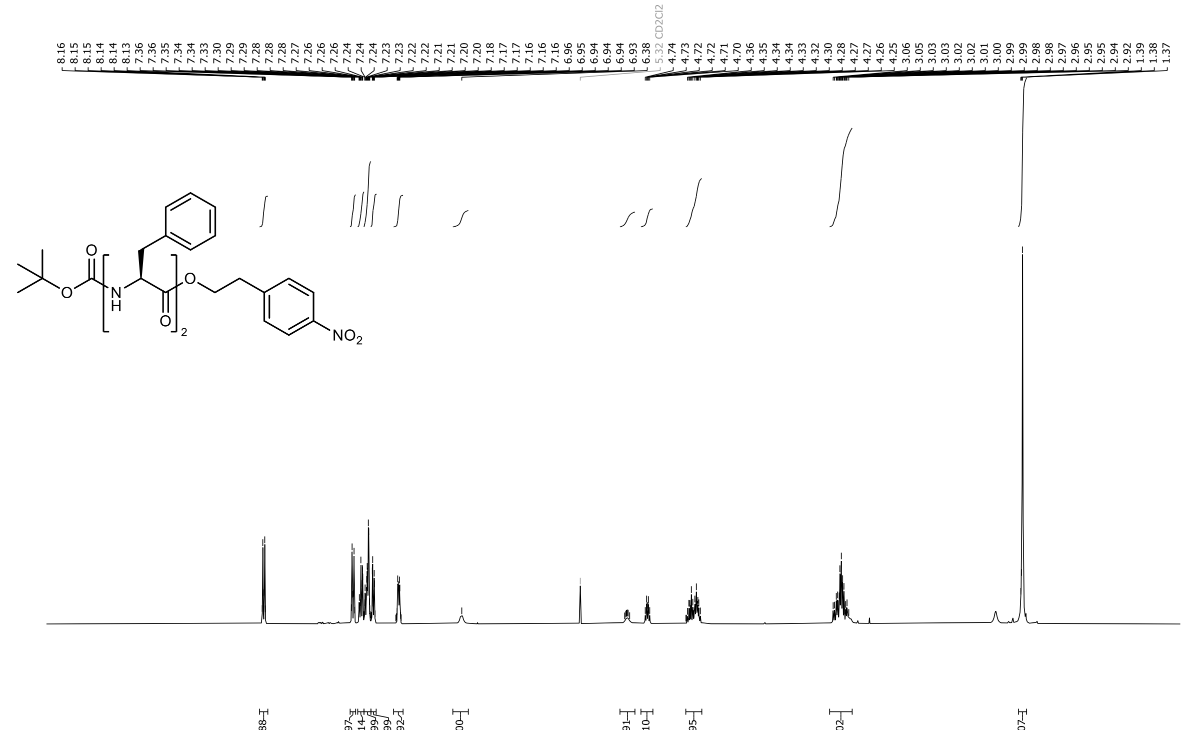

#### $^{13}$C NMR spectrum of compound 2o

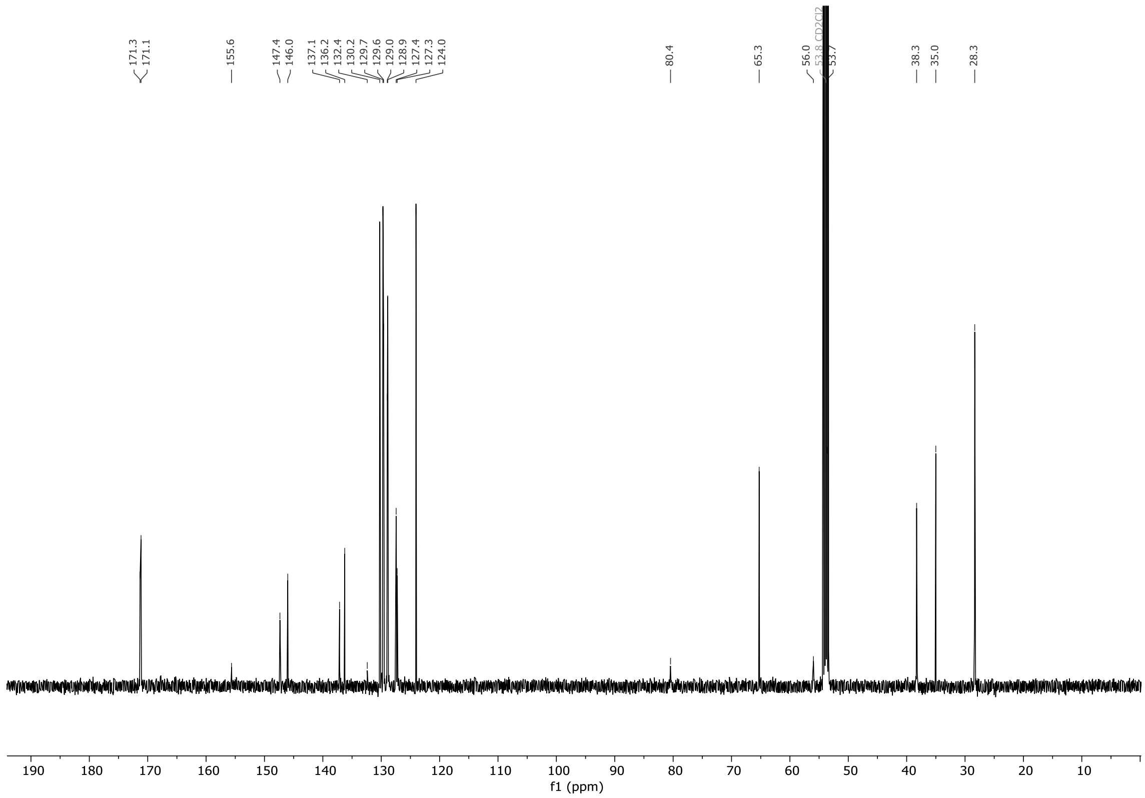


#### $^{1}$H NMR spectrum of compound 2p

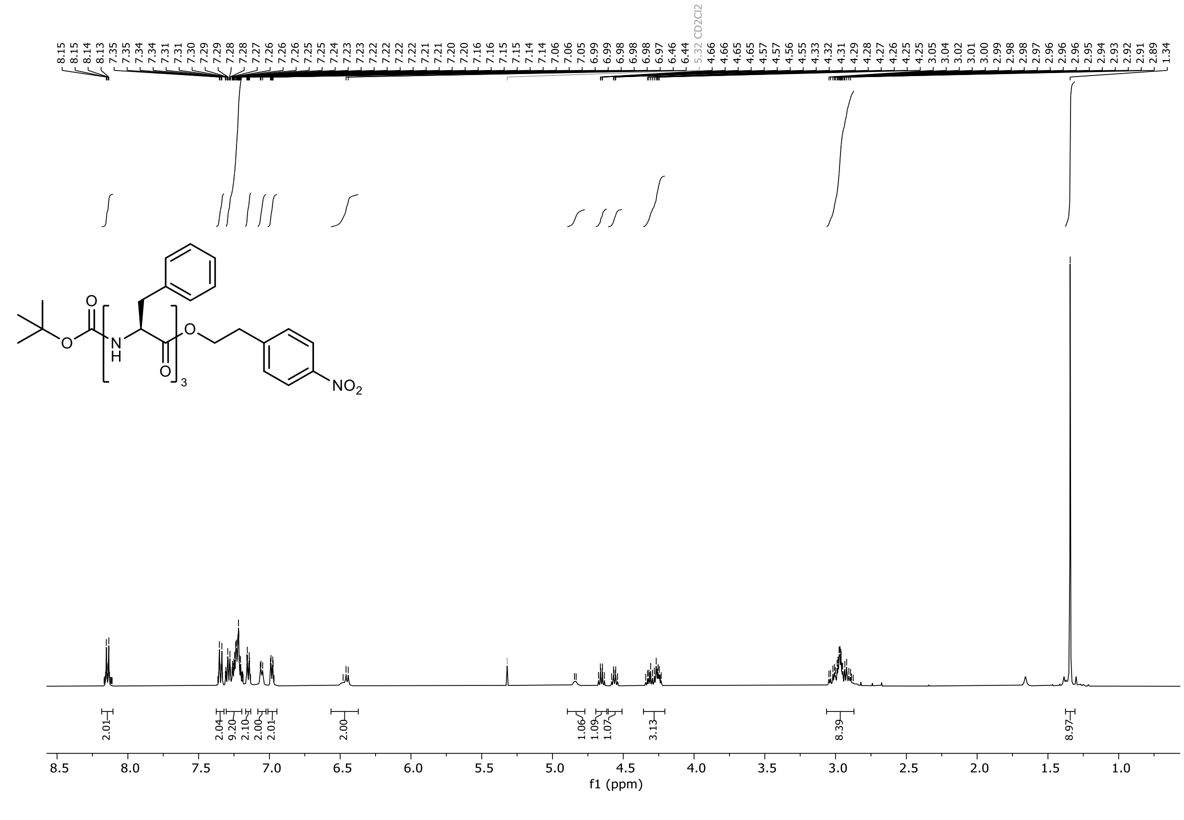

#### $^{13}C$ NMR spectrum of compound 2p

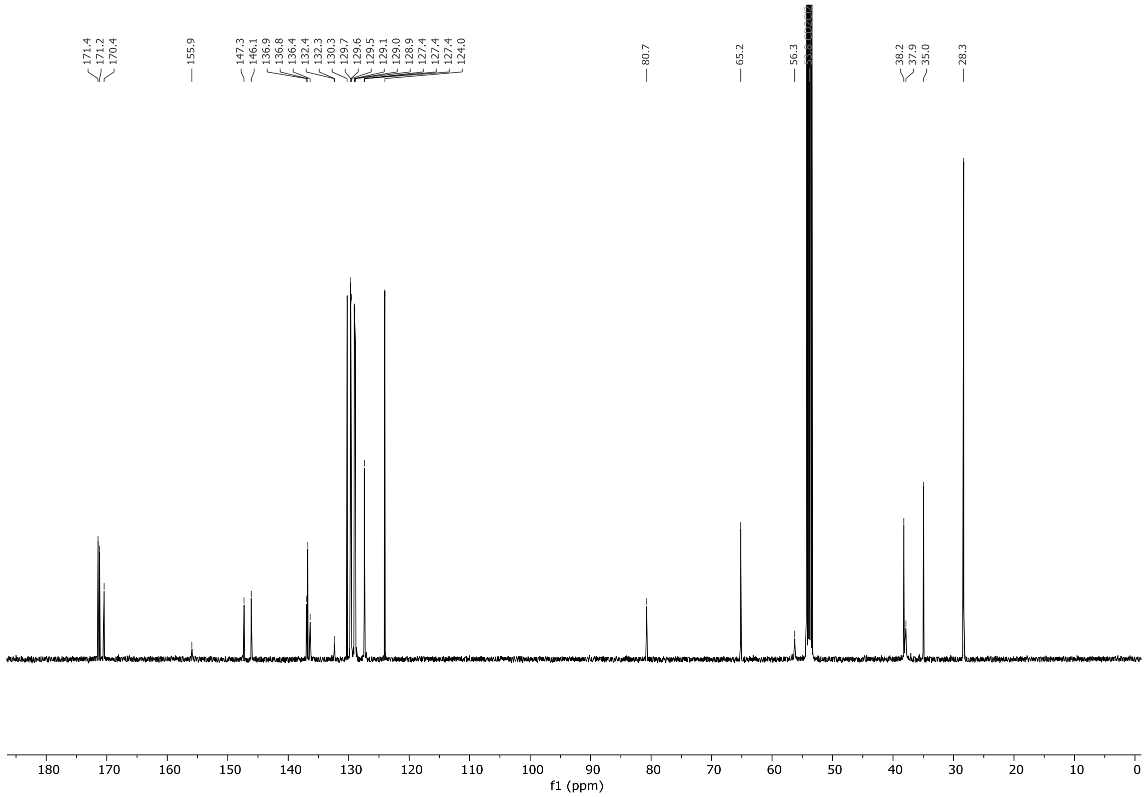

#### $^{1}H$ NMR spectrum of compound 2q

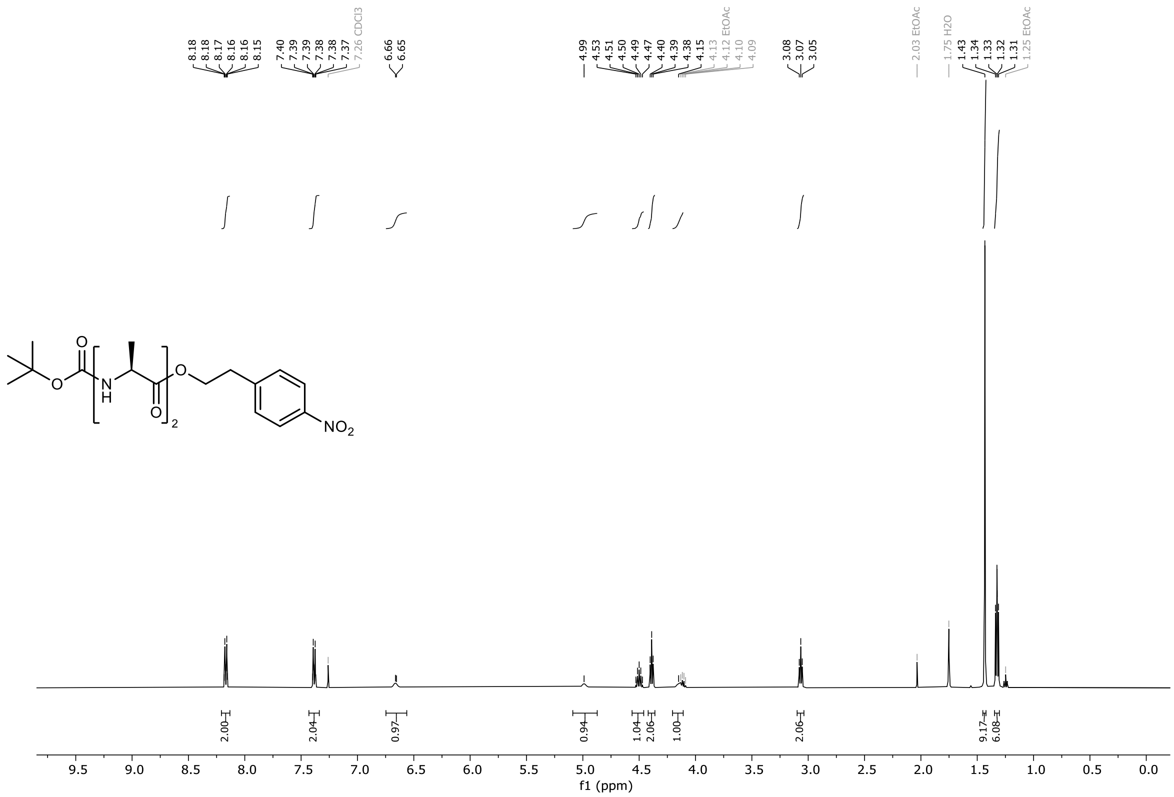

#### $^{13}C$ NMR spectrum of compound 2q

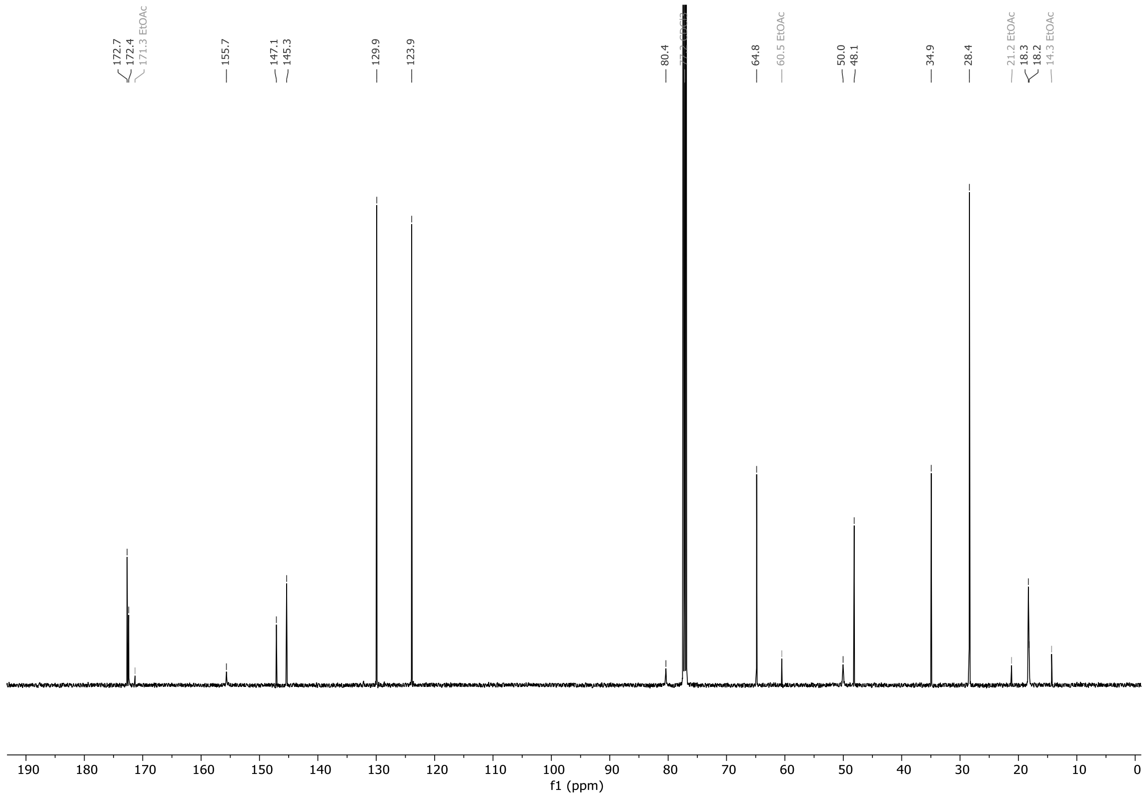


#### $^{1}H$ NMR spectrum of compound 2r

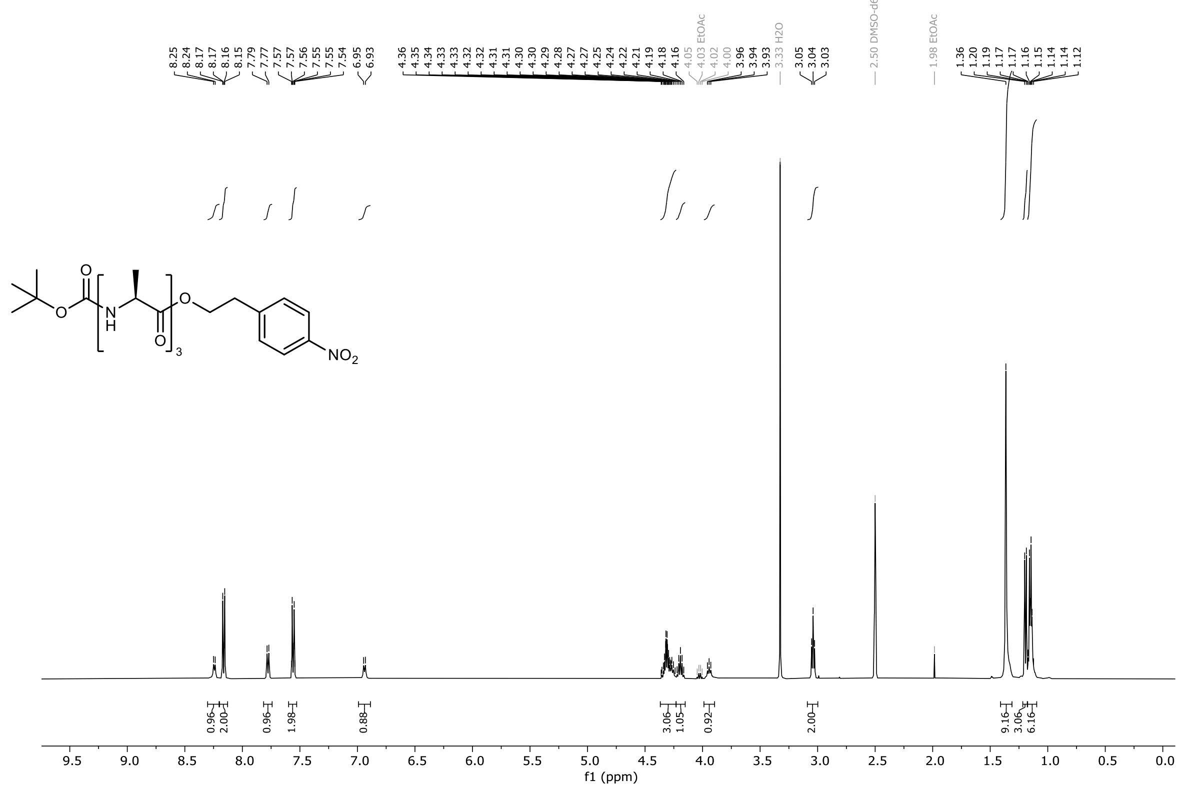

#### $^{13}$C NMR spectrum of compound 2r

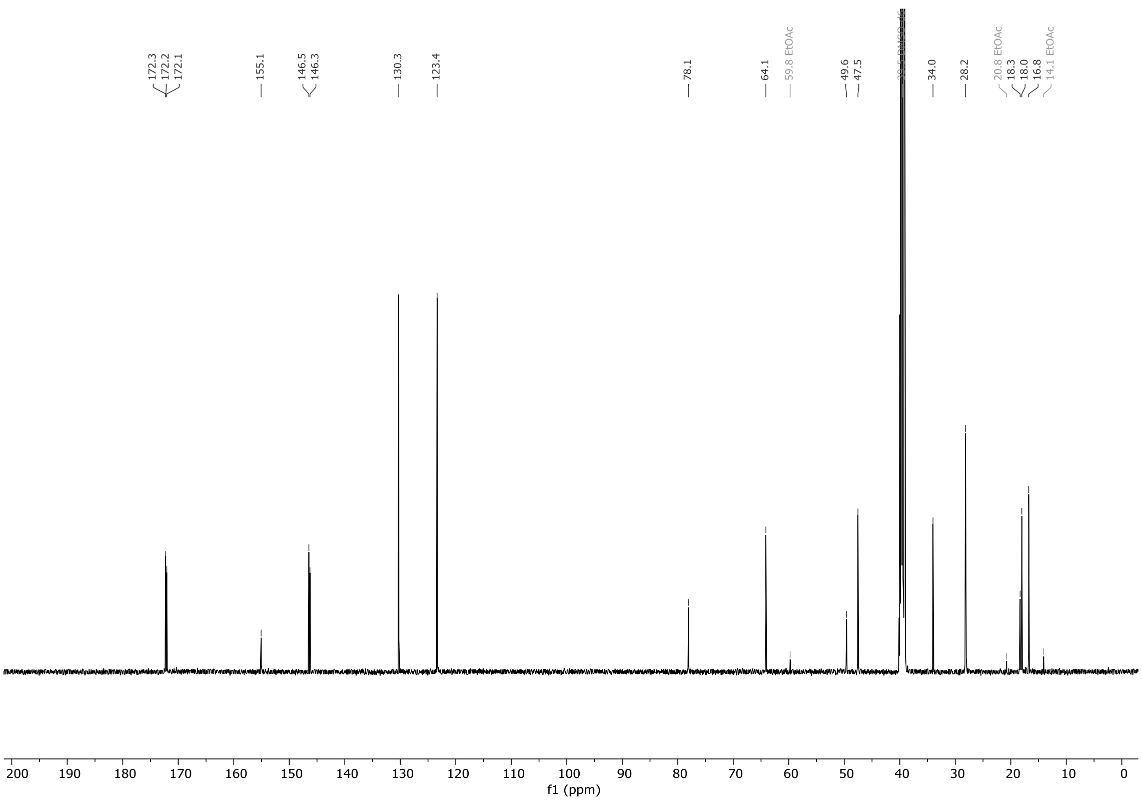


#### $^{1}$H NMR spectrum of compound 2s

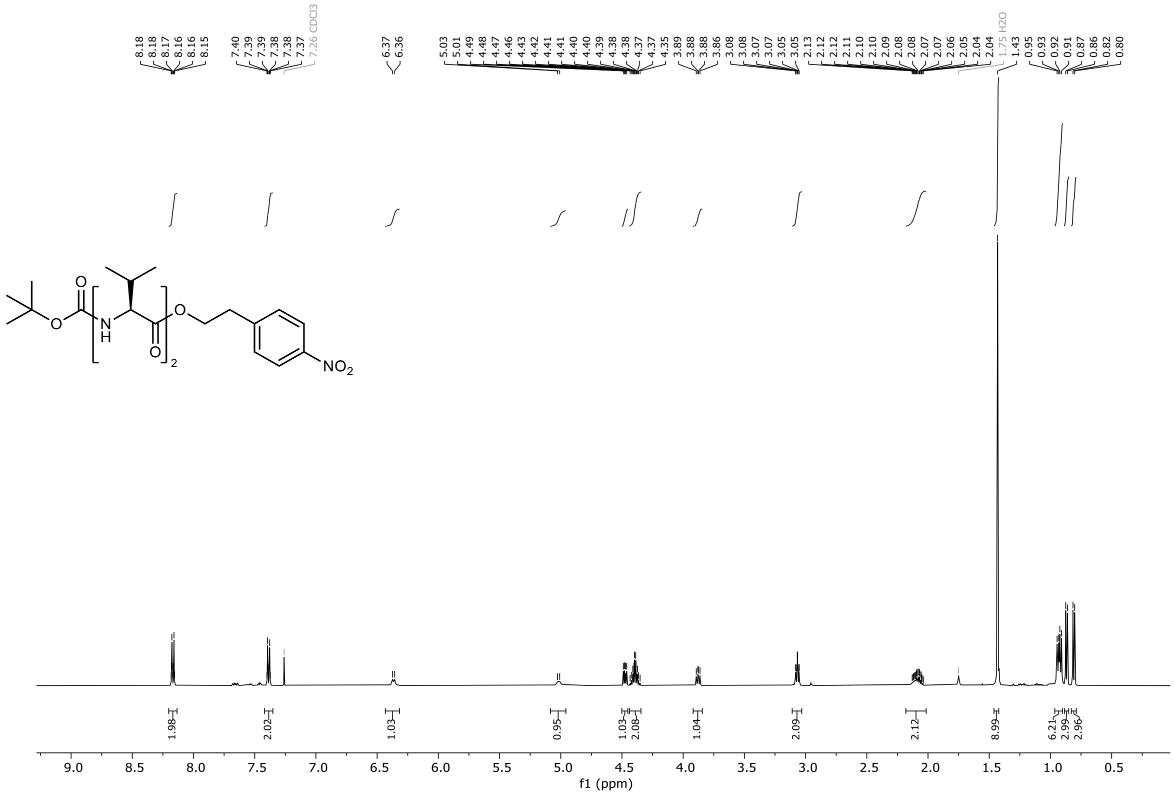

#### $^1$H NMR spectrum of compound 2t

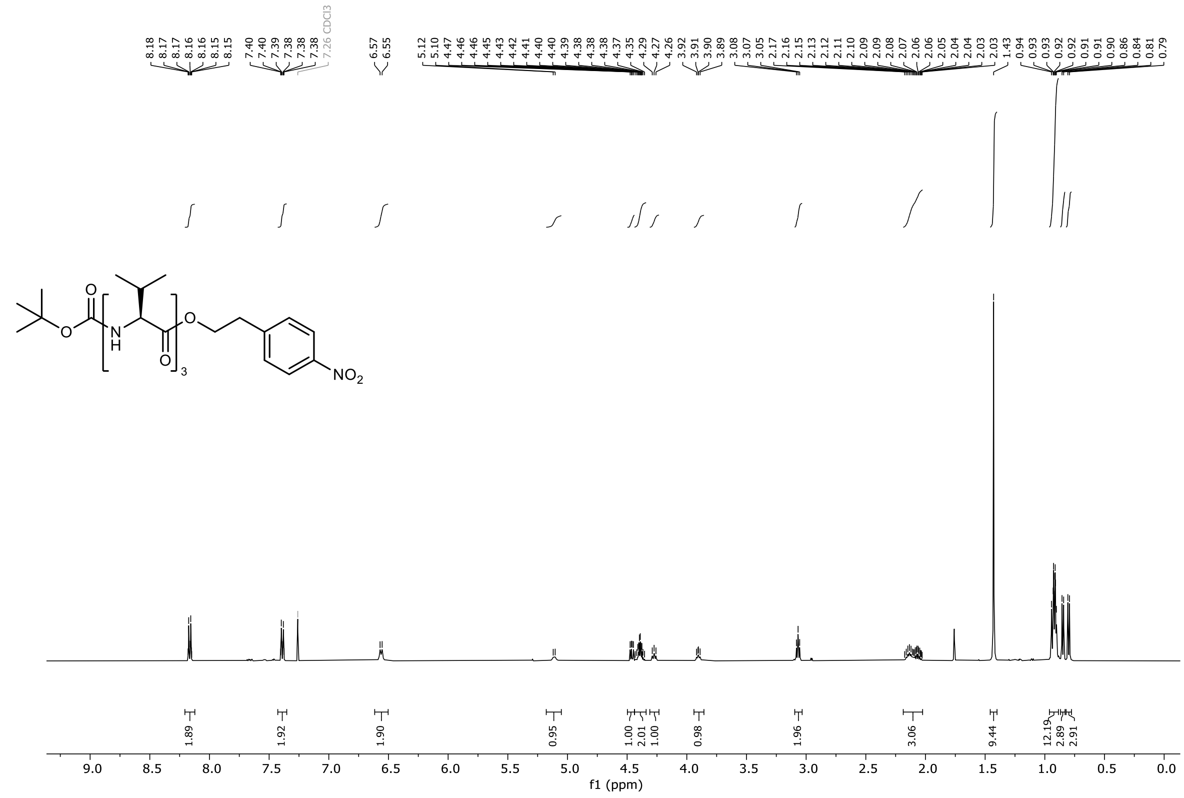

#### $^1$H NMR spectrum of compound 4a

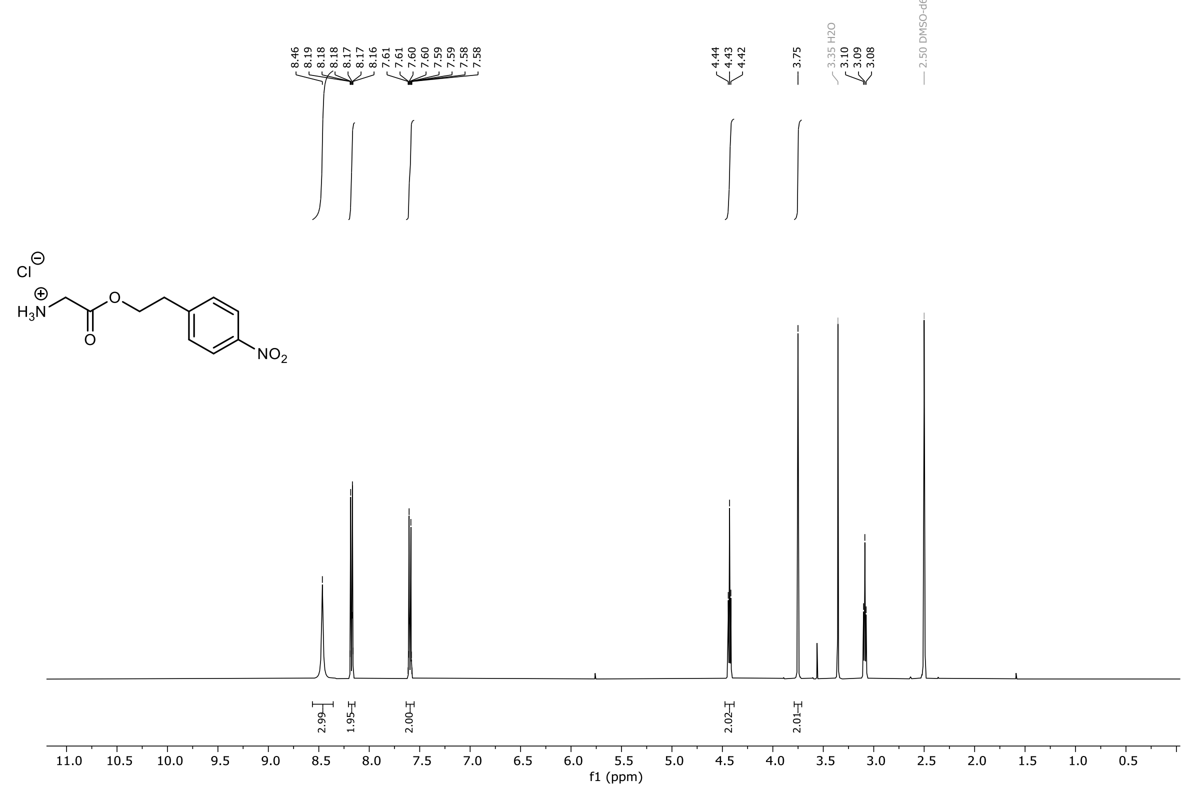

#### $^1$H NMR spectrum of compound 4b

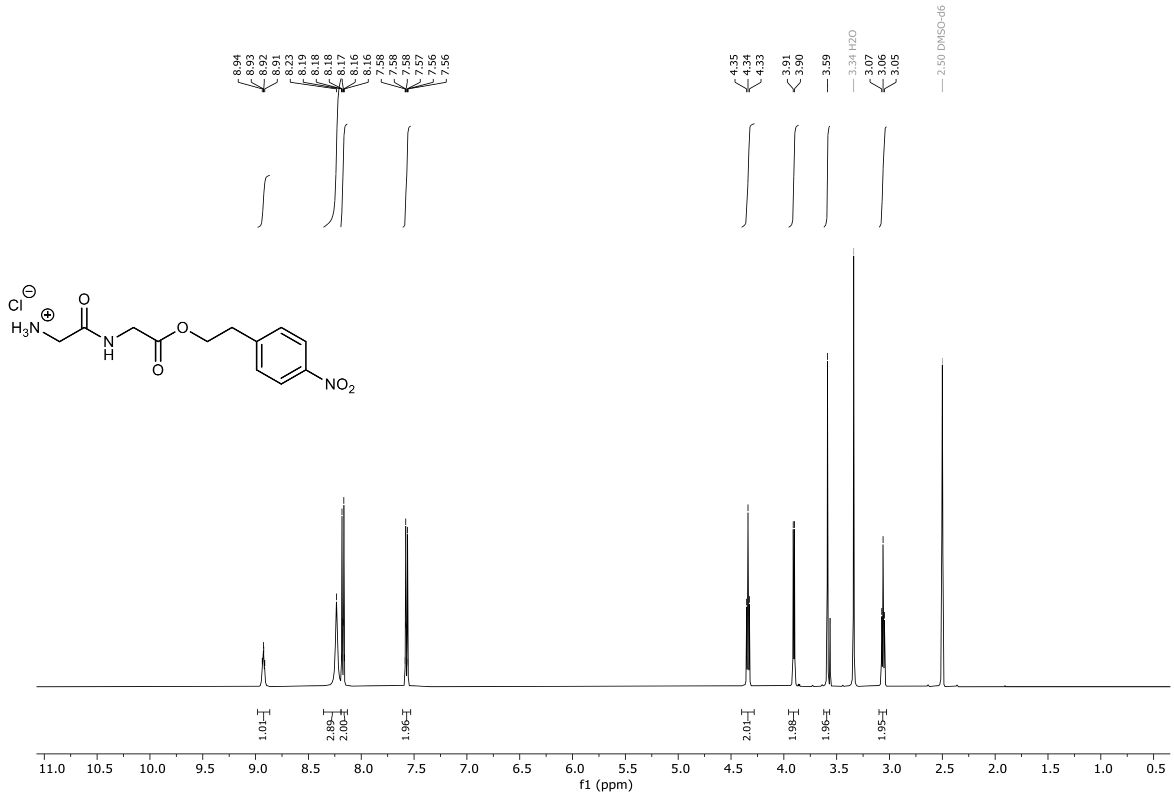

#### $^1$H NMR spectrum of compound 4c

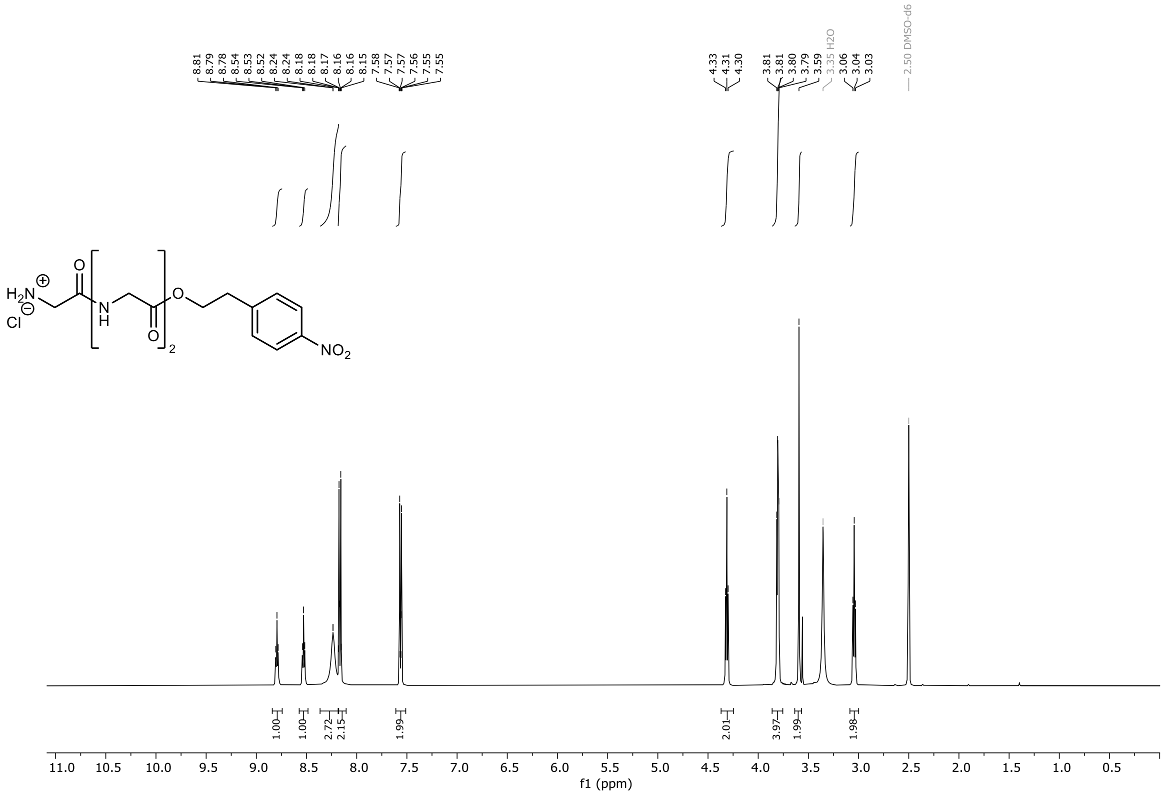

#### $^{13}$C NMR spectrum of compound 4c

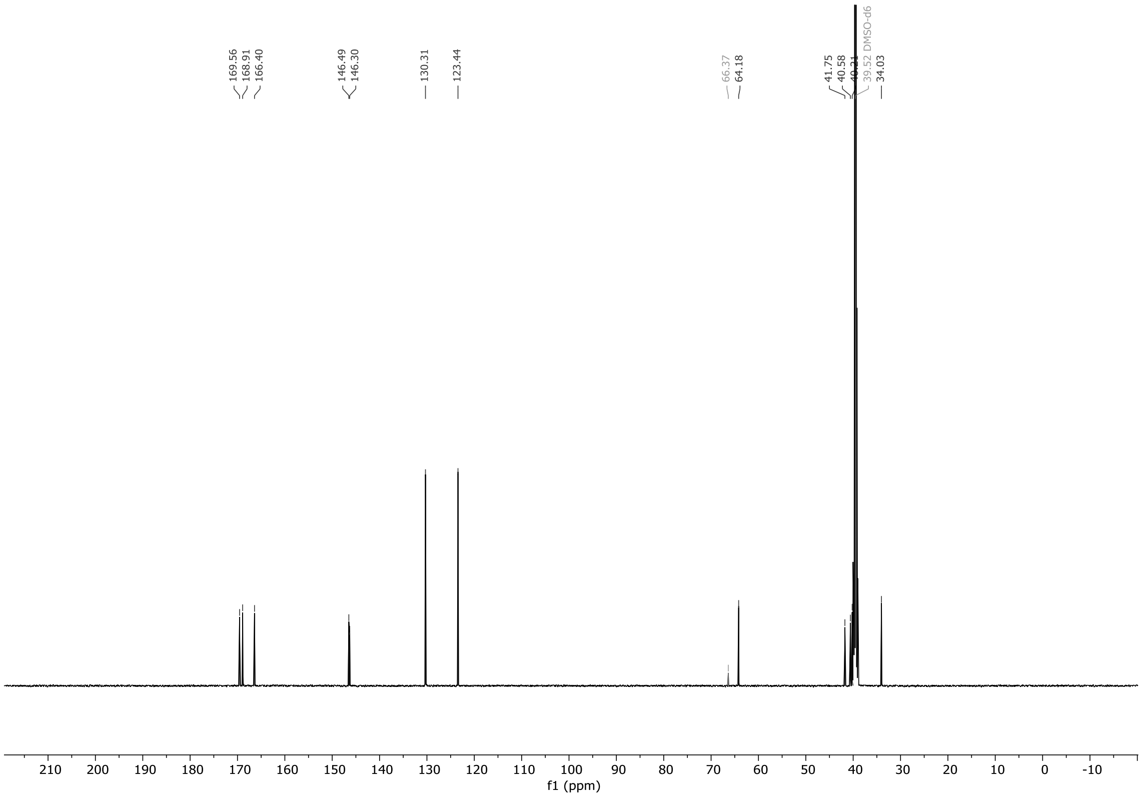


#### $^{1}$H NMR spectrum of compound 4d

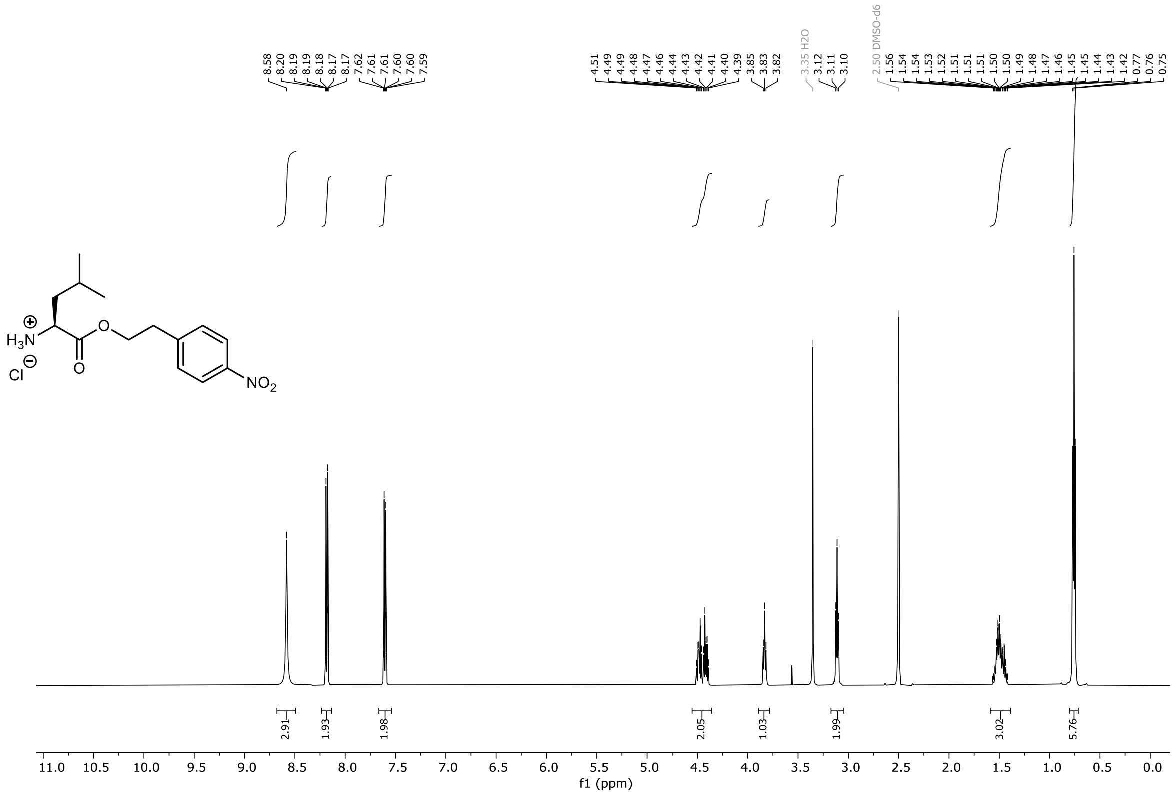

### $^1$H NMR spectrum of compound 4f

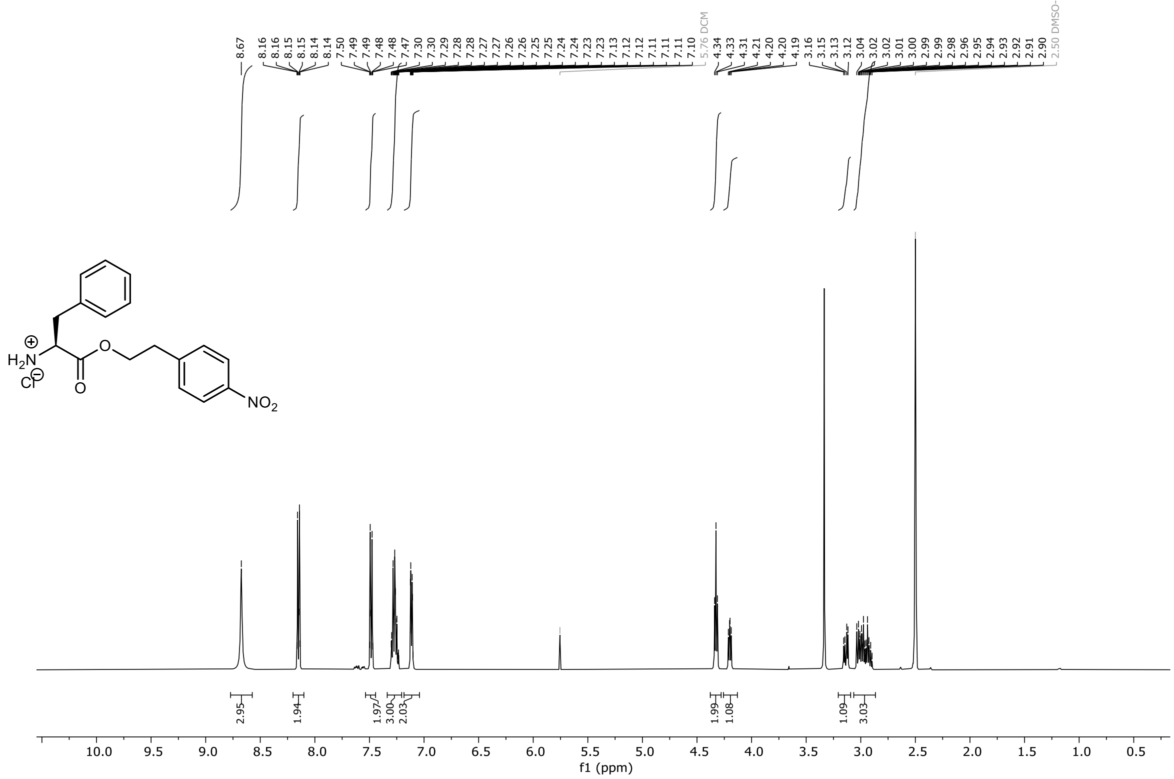


### $^1$H NMR spectrum of compound 4h

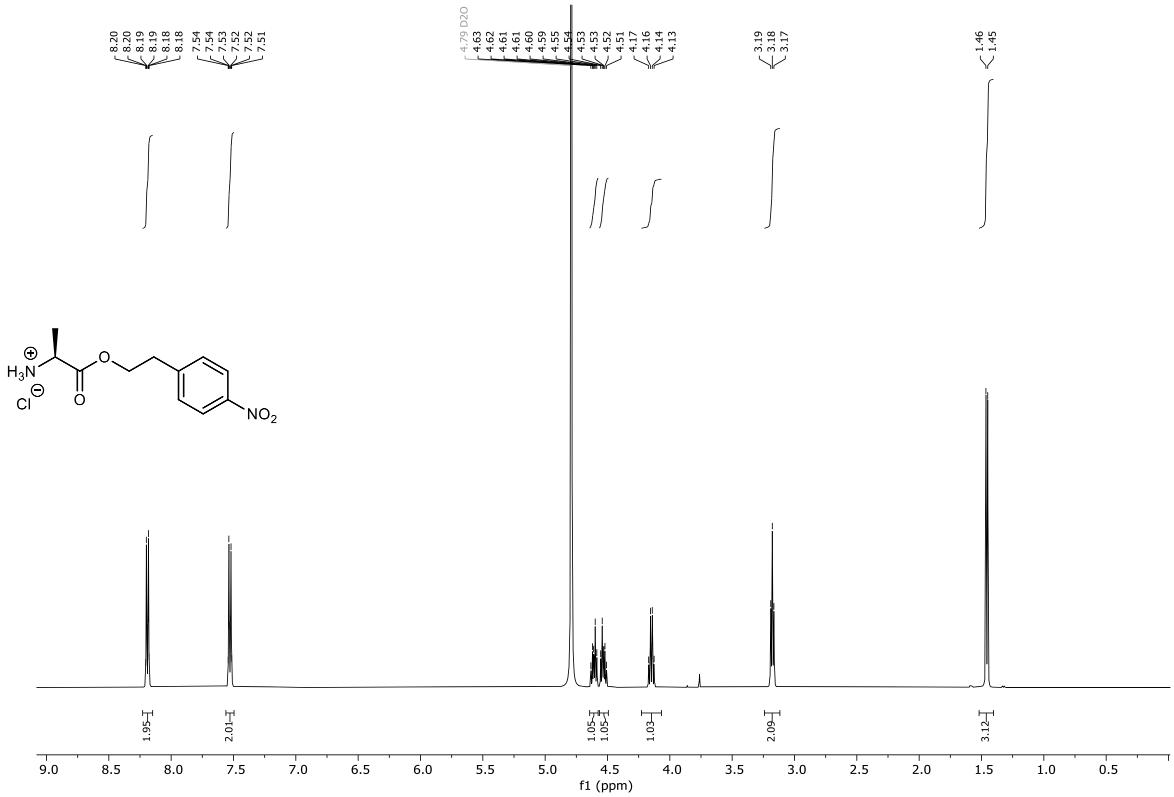

#### $^1$H NMR spectrum of compound 4i

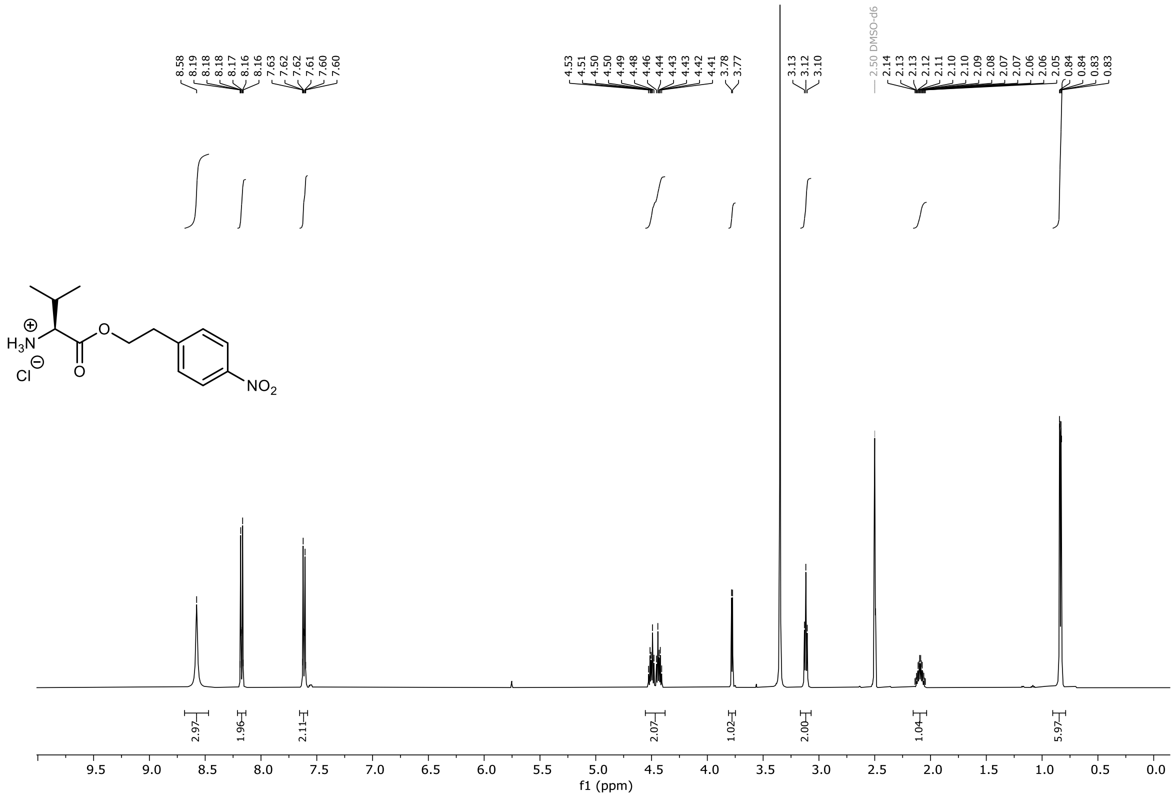

#### $^1$H NMR spectrum of compound 4k

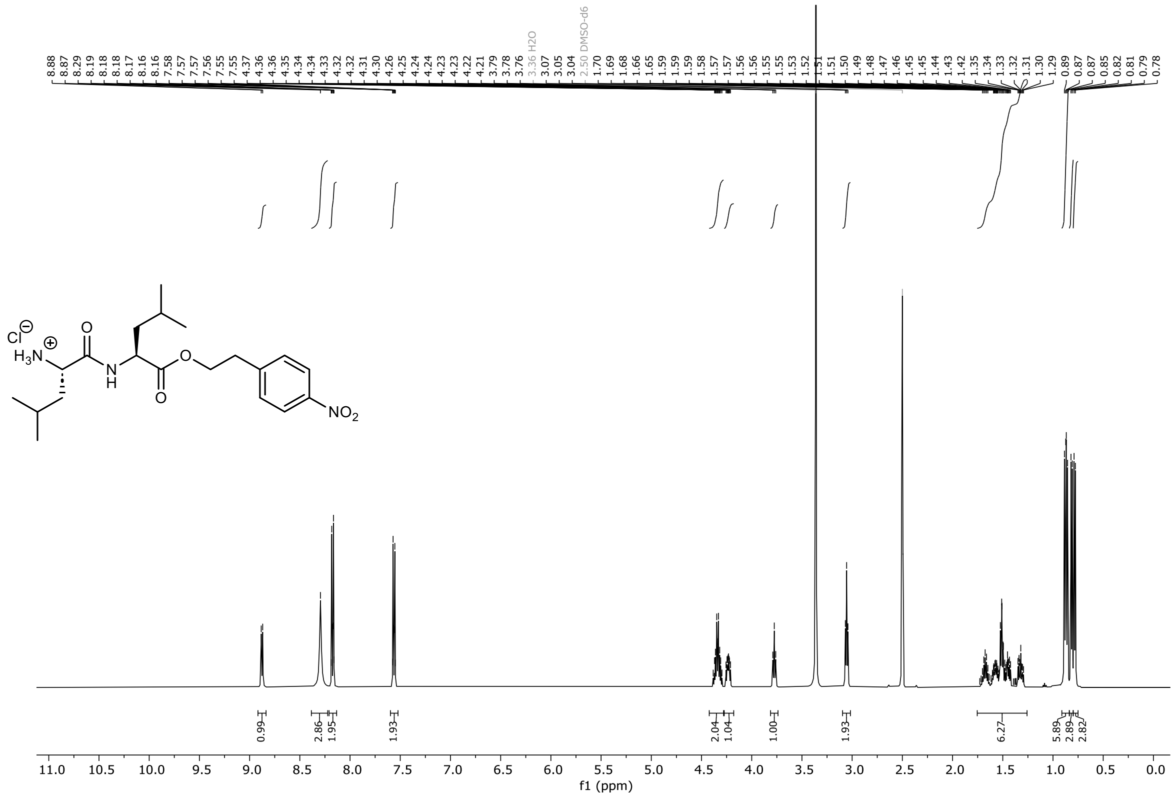

#### $^{13}$C NMR spectrum of compound 4k

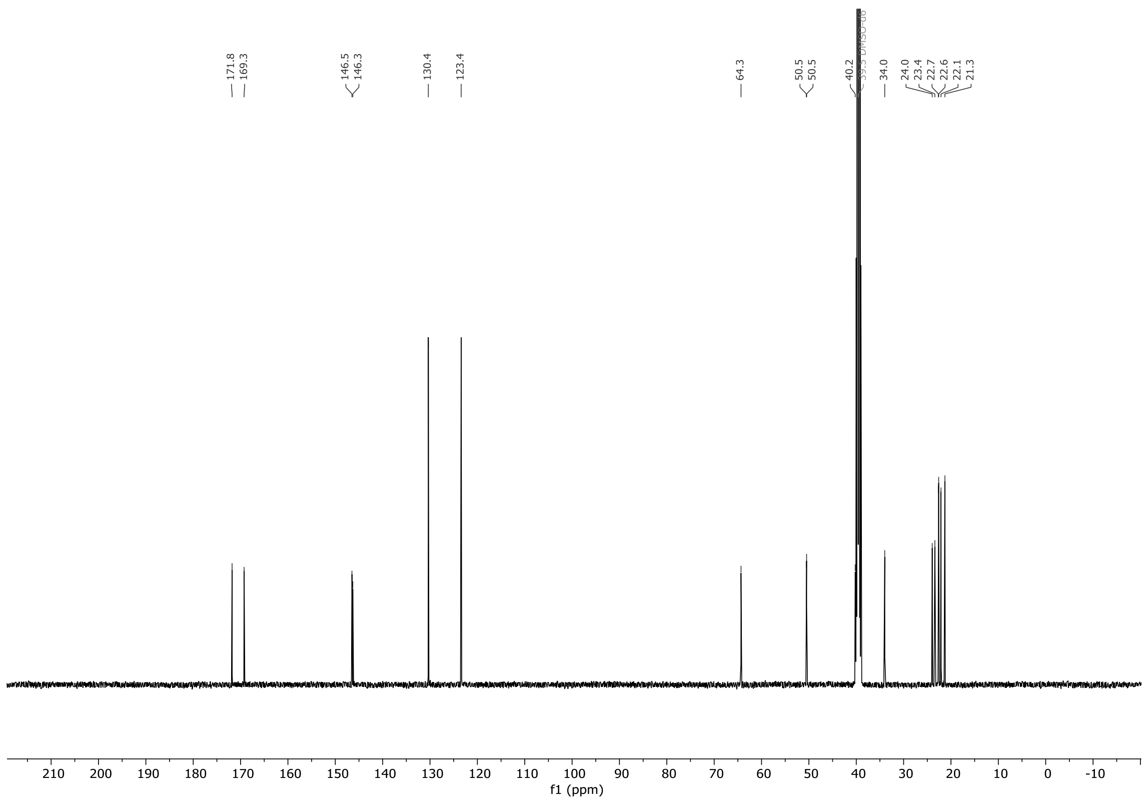


#### $^{1}$H NMR spectrum of compound 4l

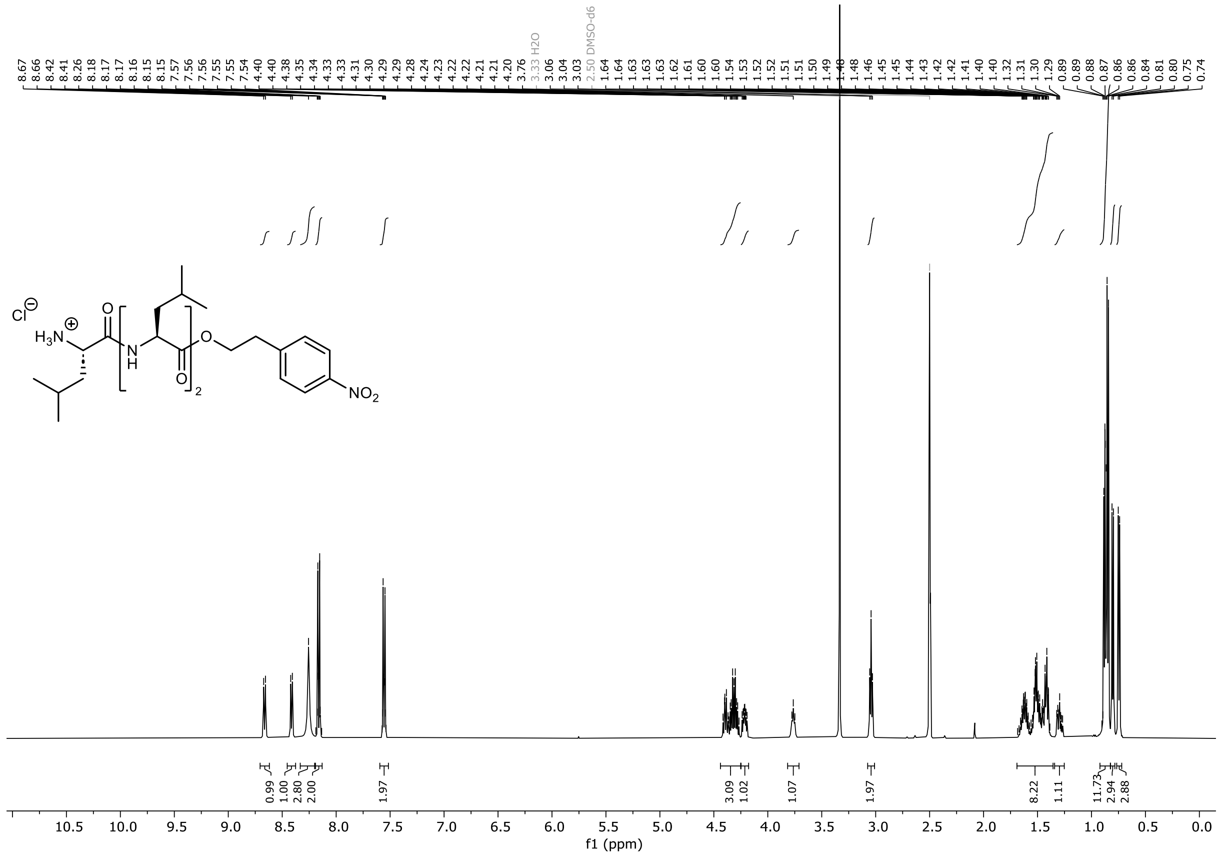


#### $^{13}$C NMR spectrum of compound 4l

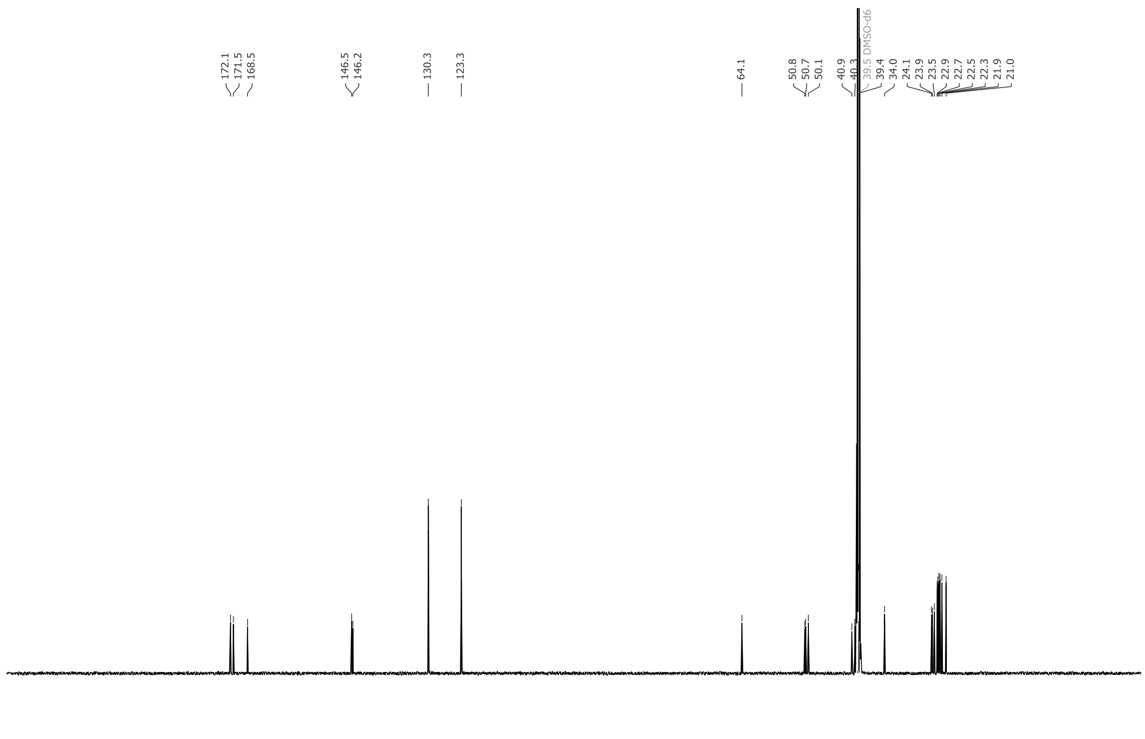

#### [1]H NMR spectrum of compound 4o

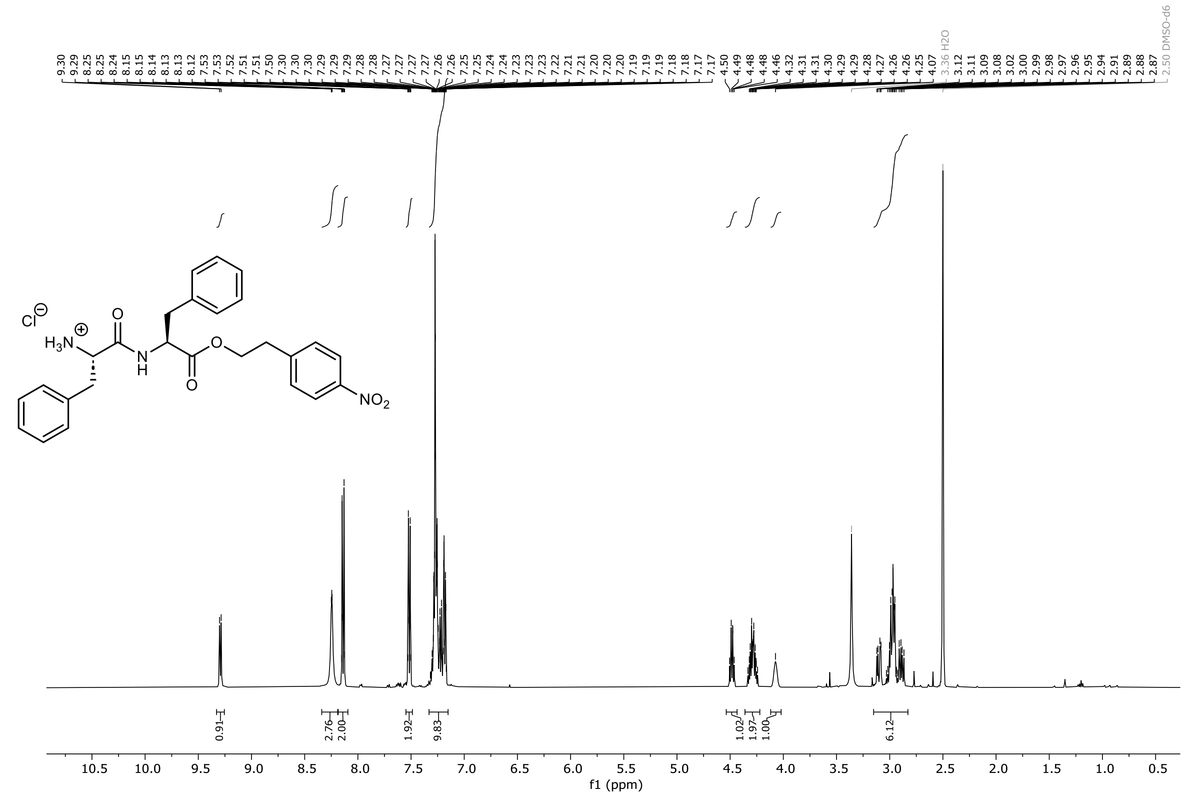

#### [13]C NMR spectrum of compound 4o

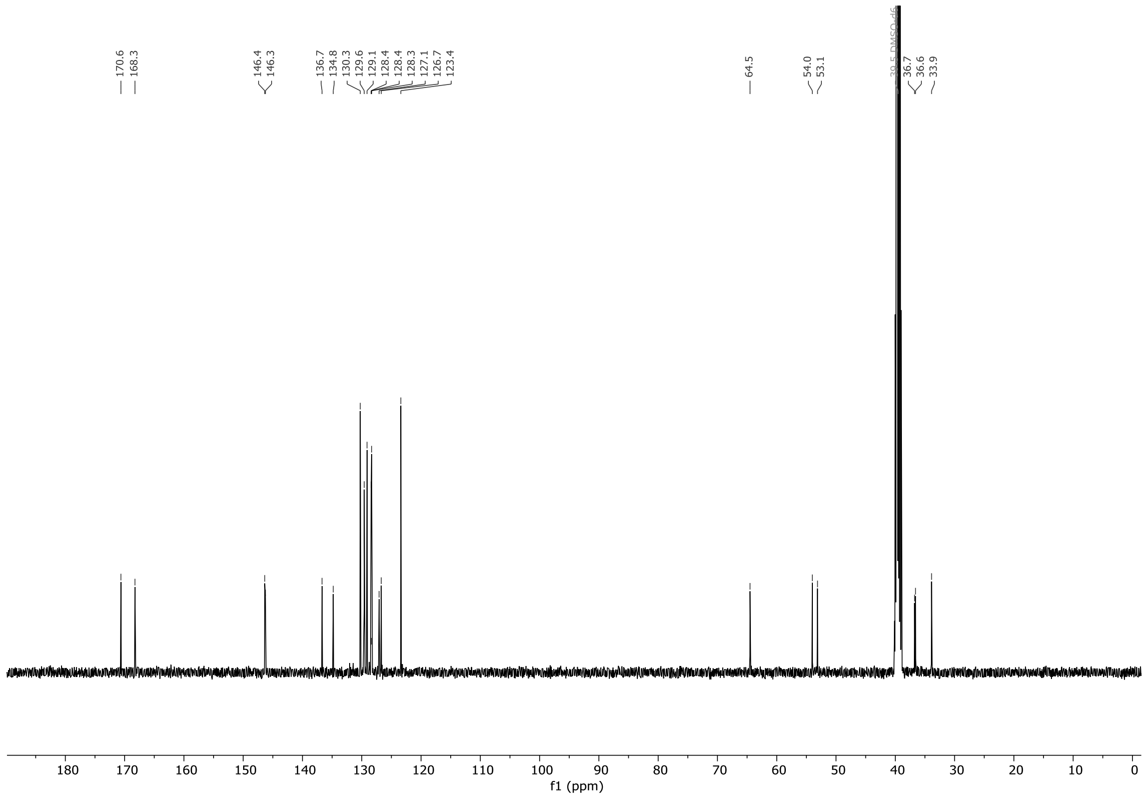

#### $^1$H NMR spectrum of compound 4p

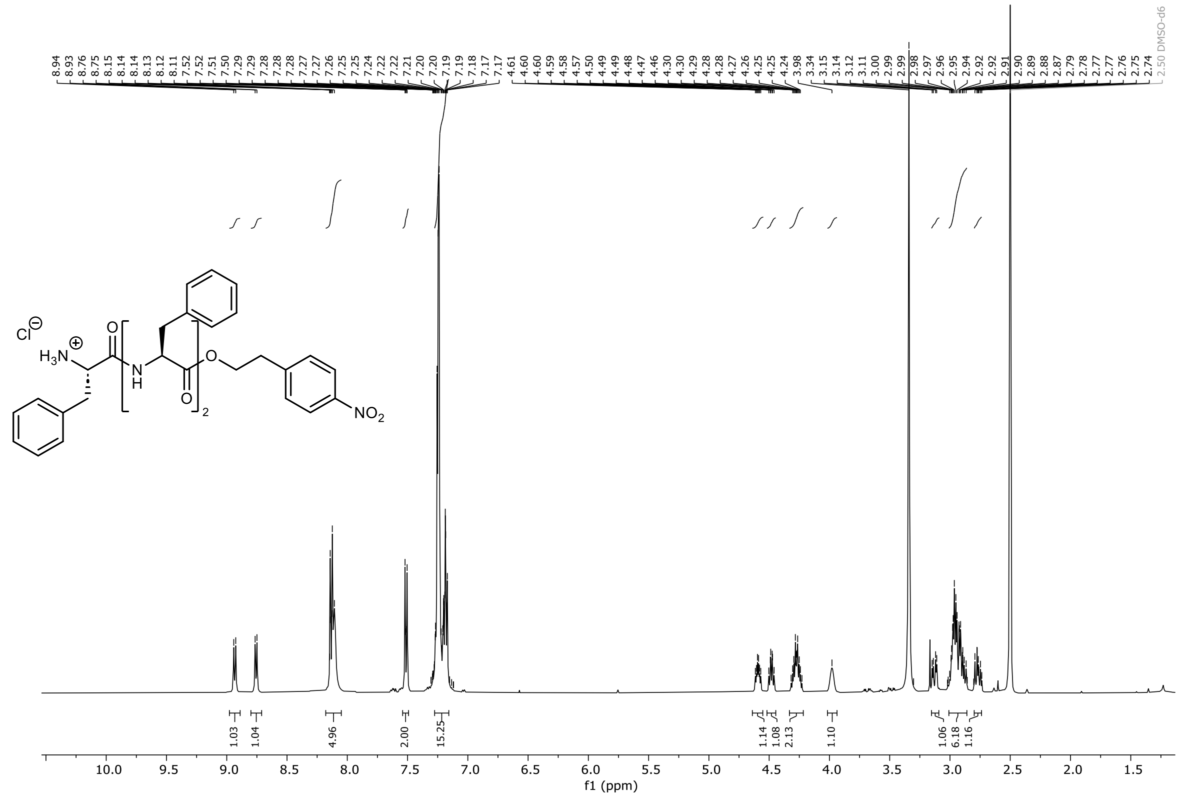


#### $^{13}$C NMR spectrum of compound 4p

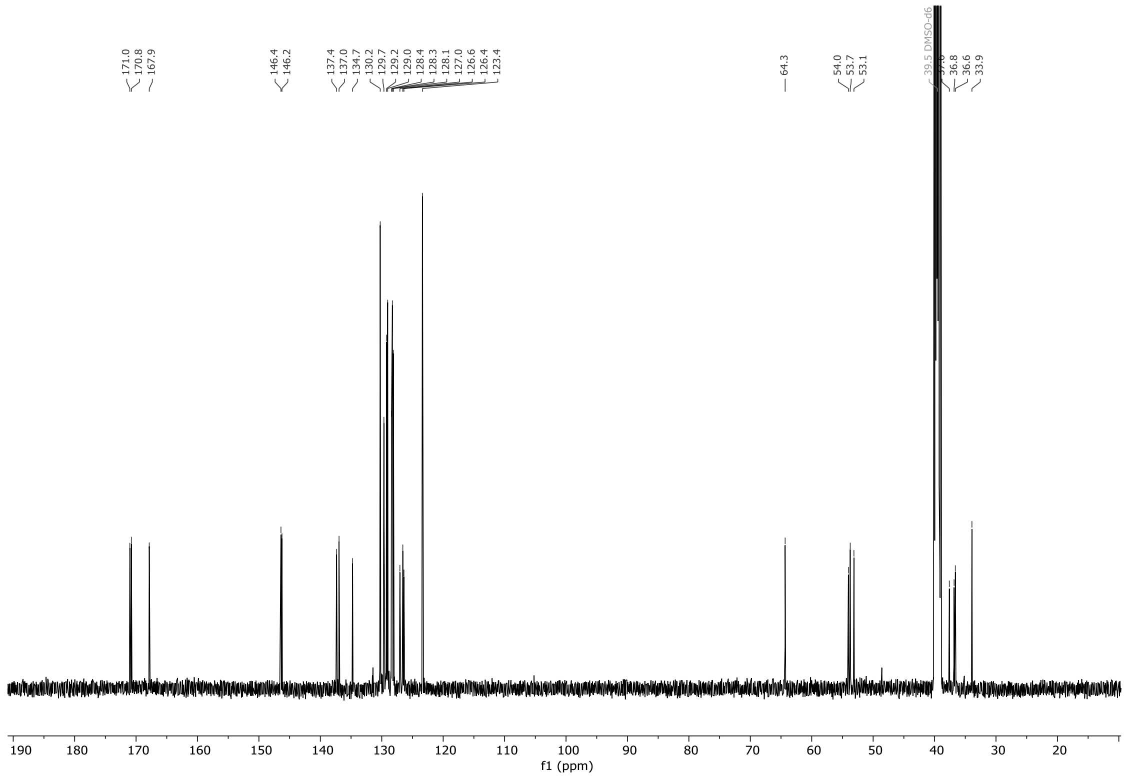


#### $^1$H NMR spectrum of compound 4q

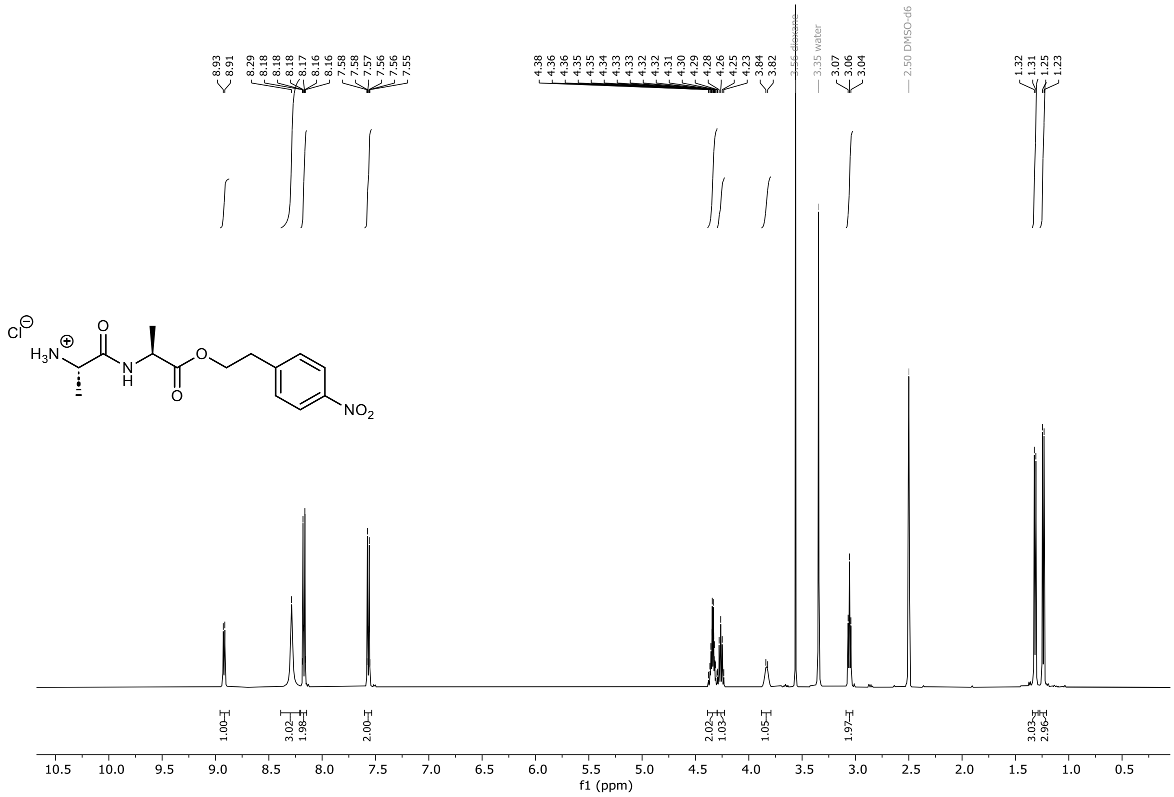


#### $^{13}$C NMR spectrum of compound 4q

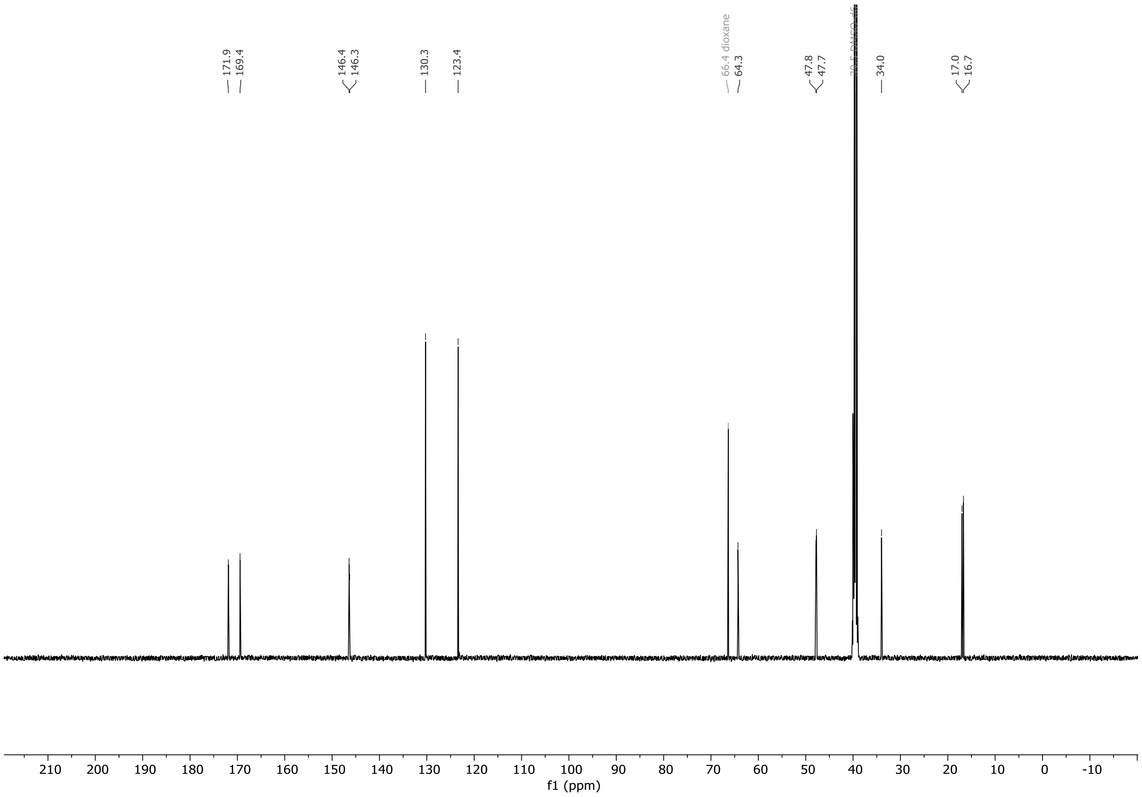

#### $^{1}$H NMR spectrum of compound 4r

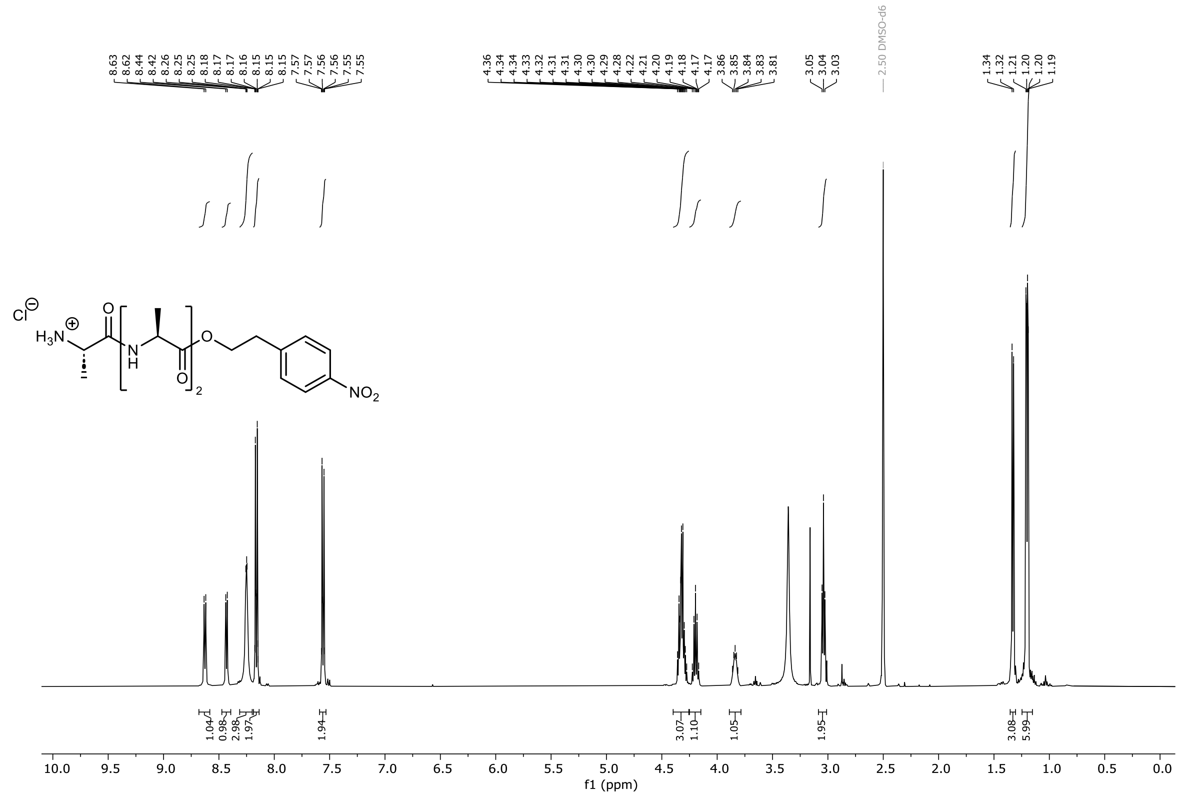


#### $^{13}$C NMR spectrum of compound 4r

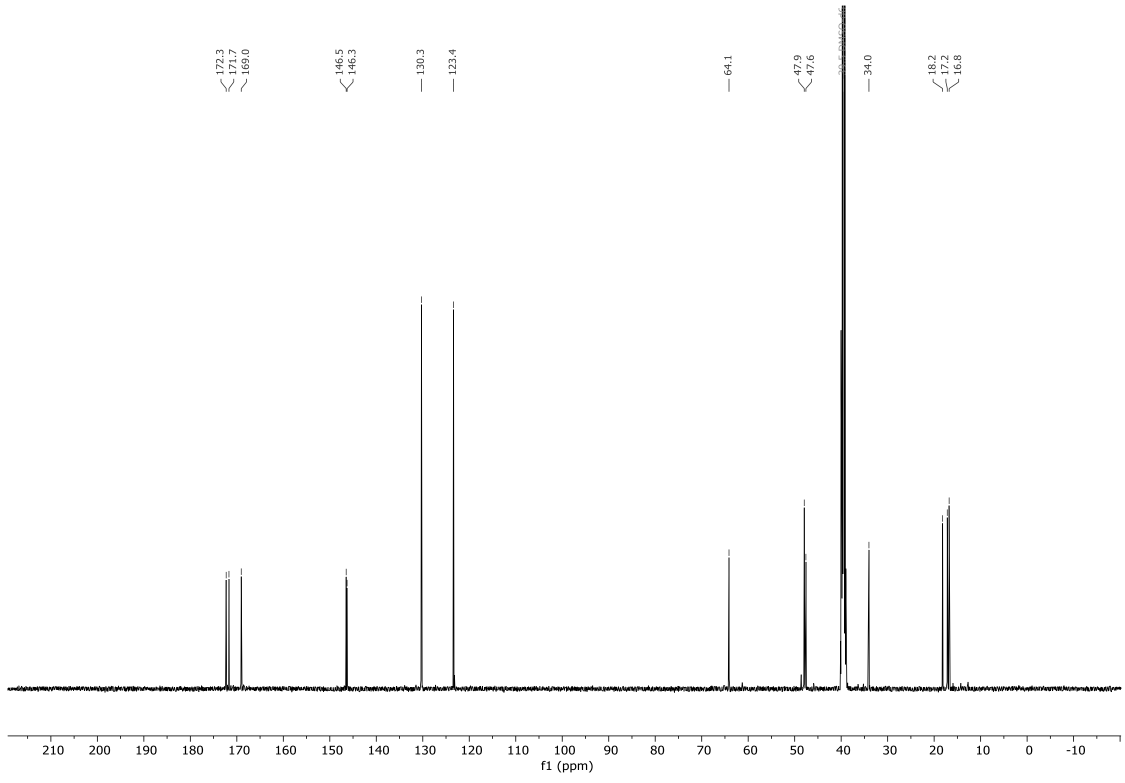


#### $^{1}$H NMR spectrum of compound 4s

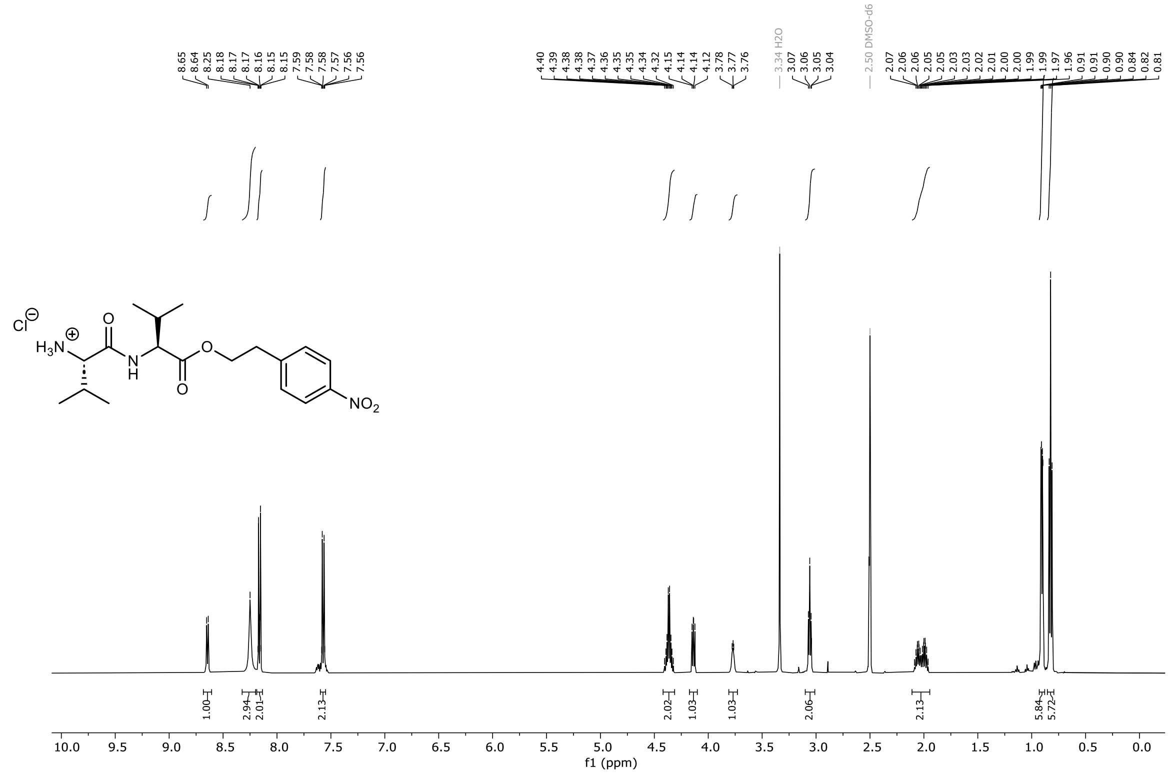


**$^{1}$H NMR spectrum of compound 4t**

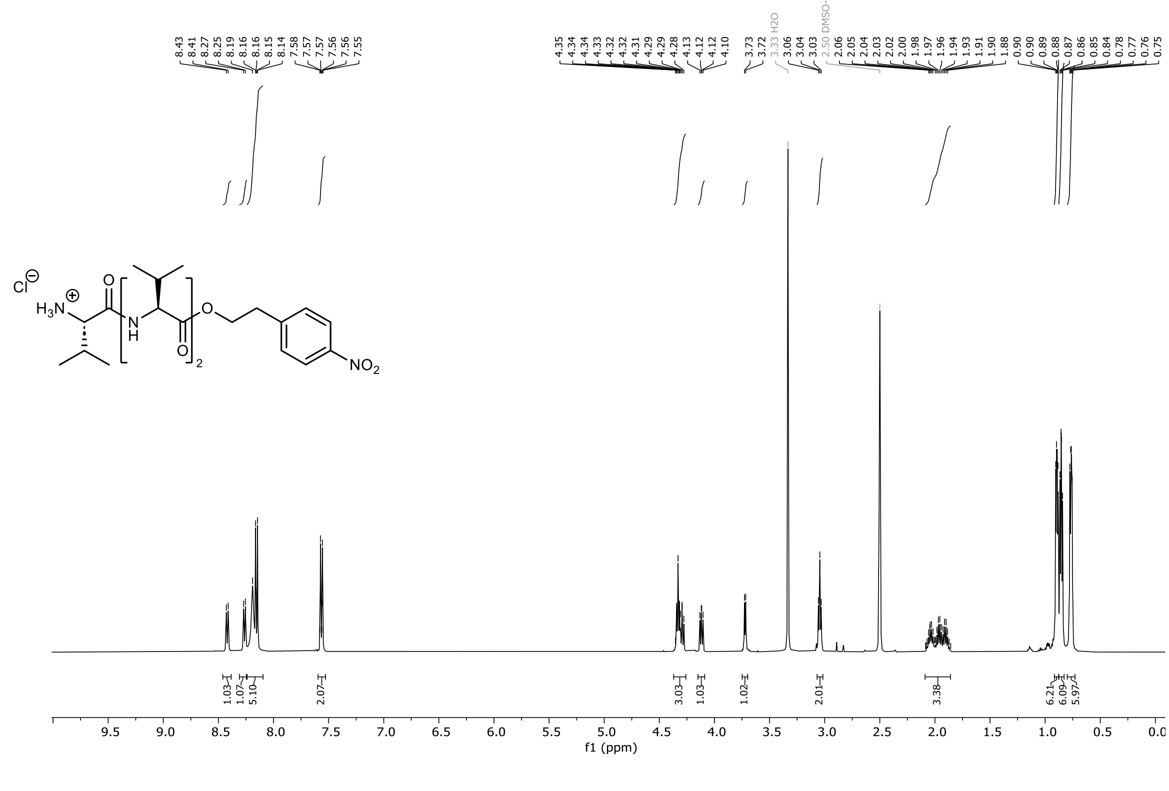


**$^{1}$H NMR spectrum of compound 7**

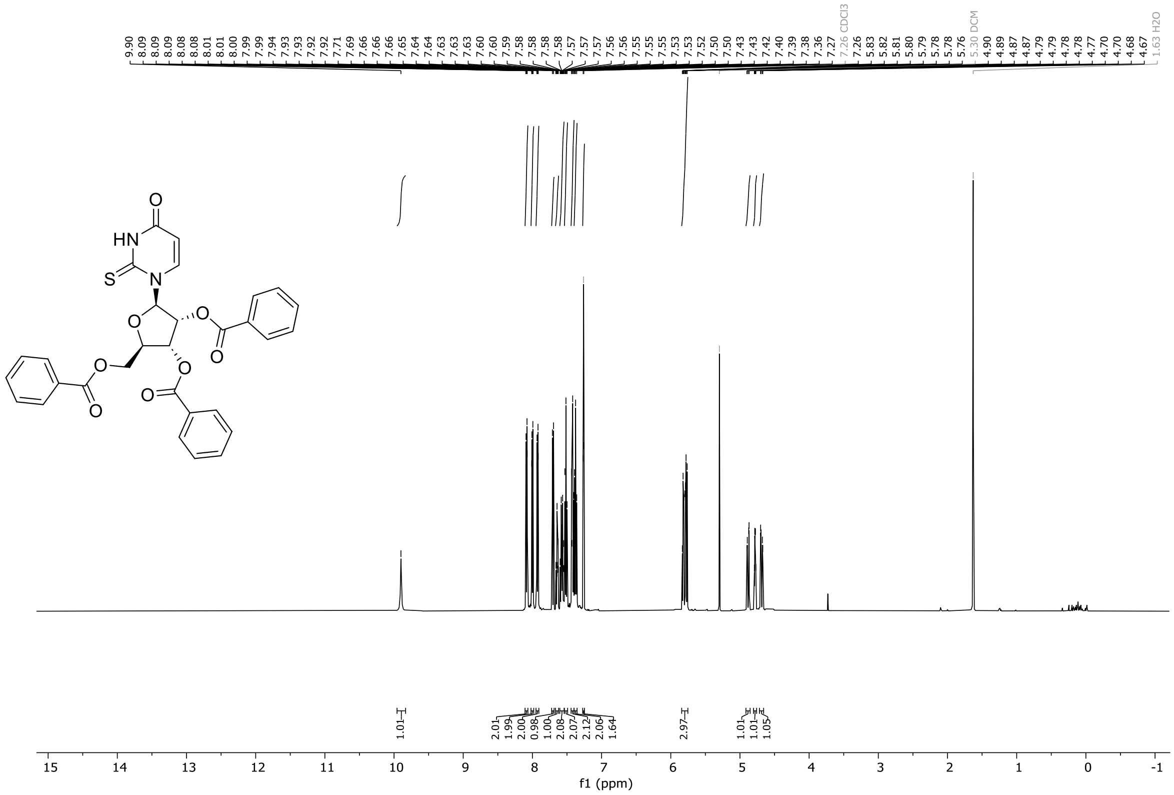


**$^1$H NMR spectrum of compound 8**

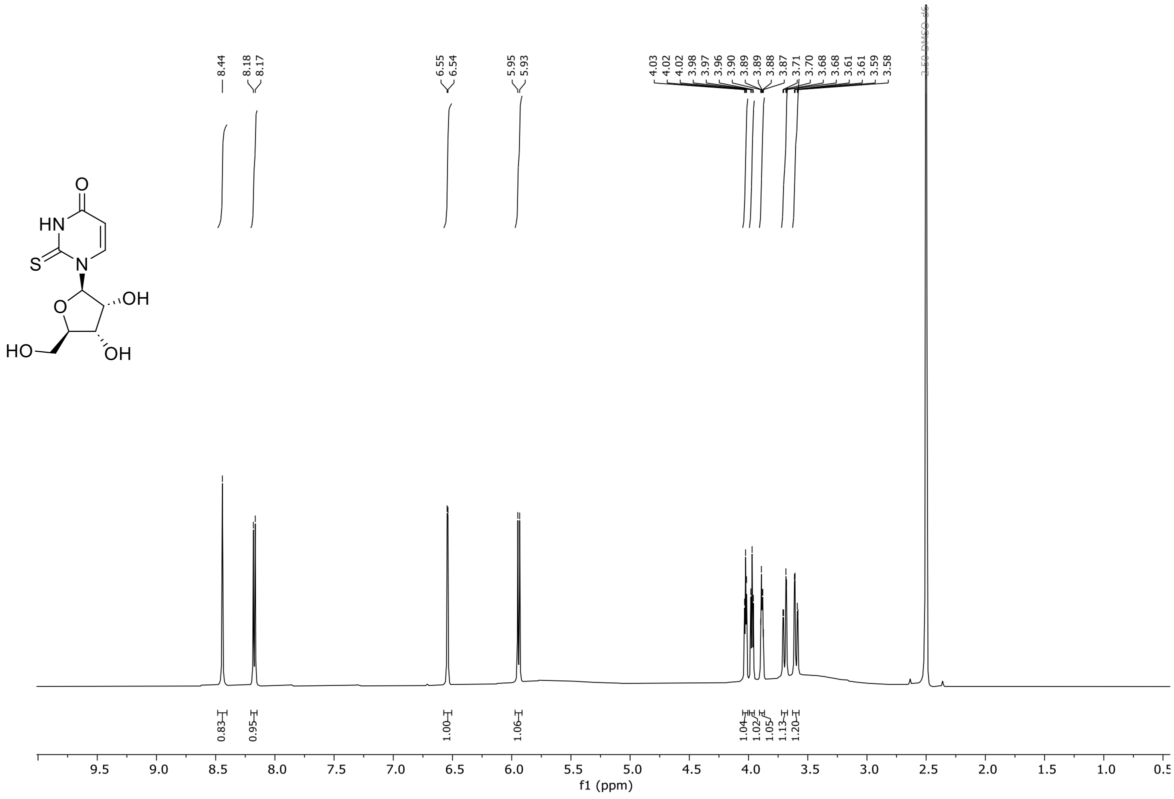


**$^1$H NMR spectrum of compound 9**

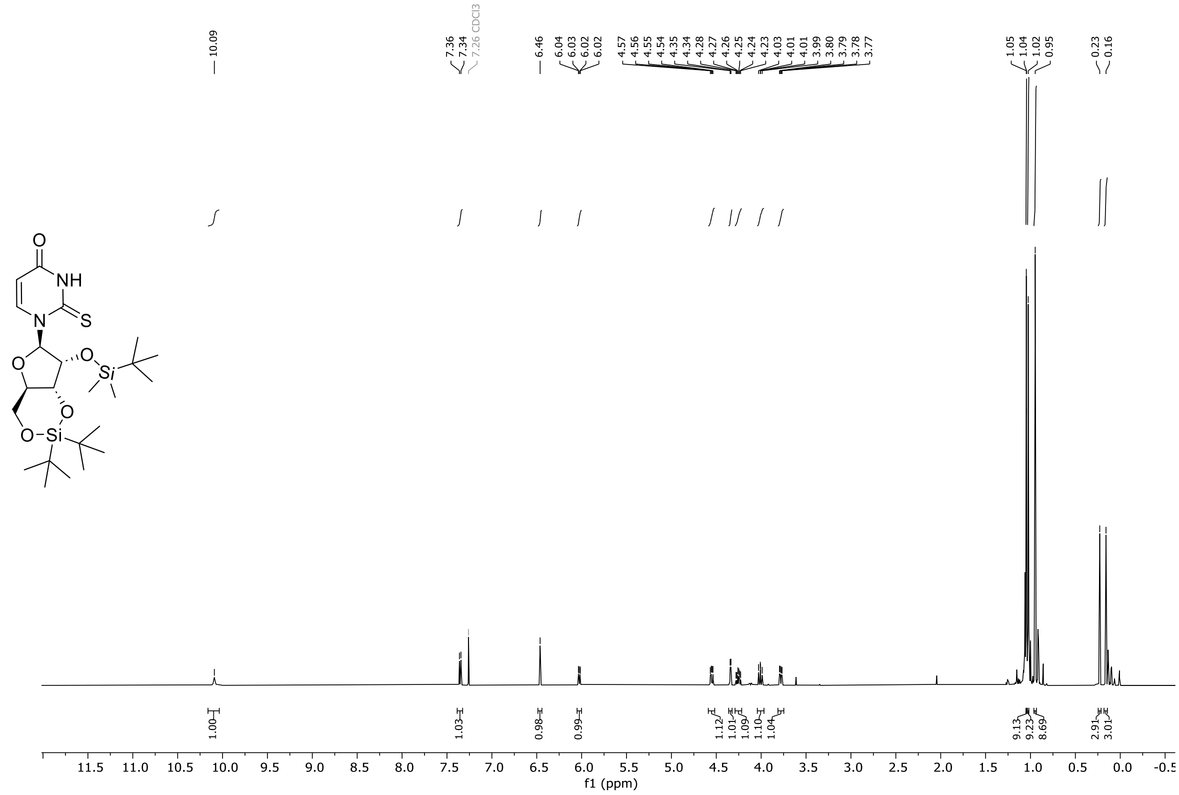

**[1]H NMR spectrum of compound 10**

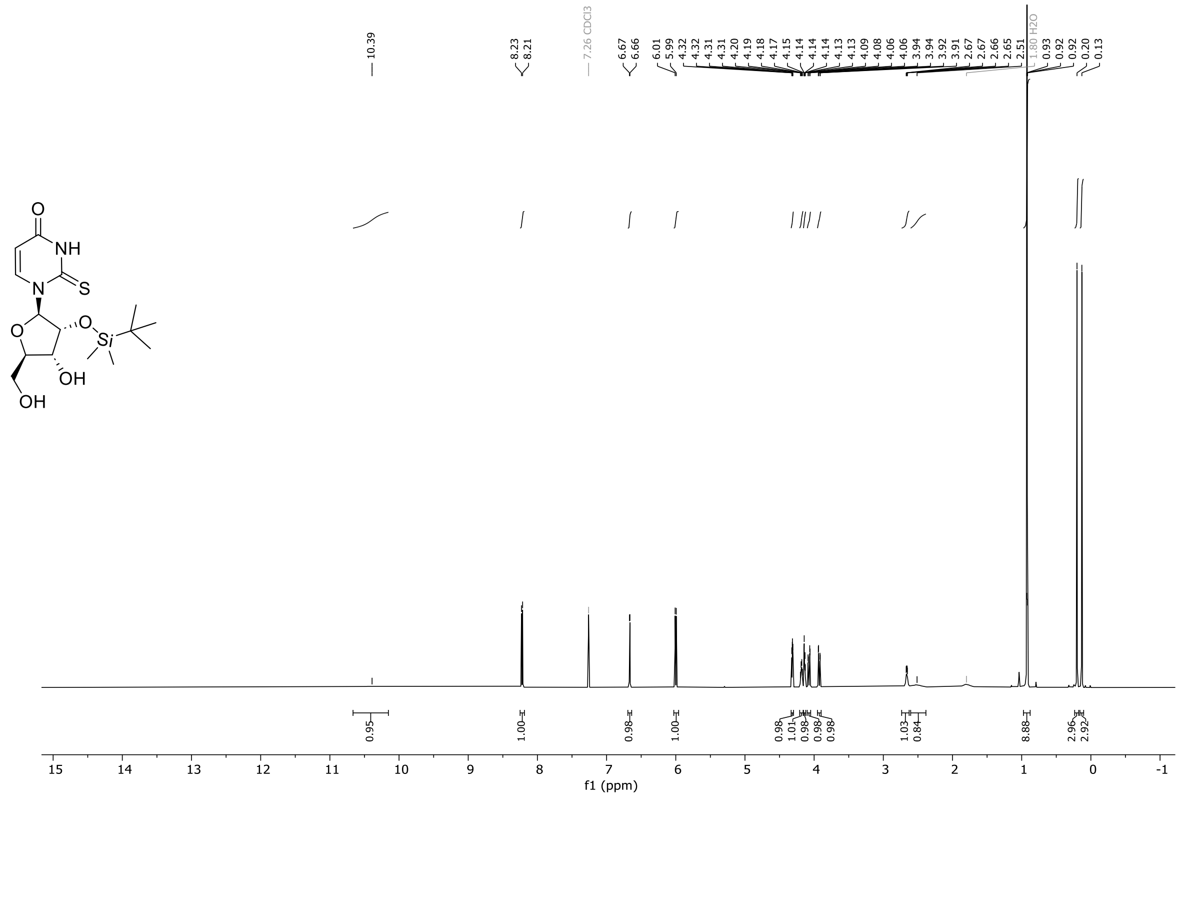

#### $^{1}$H NMR spectrum of compound 11

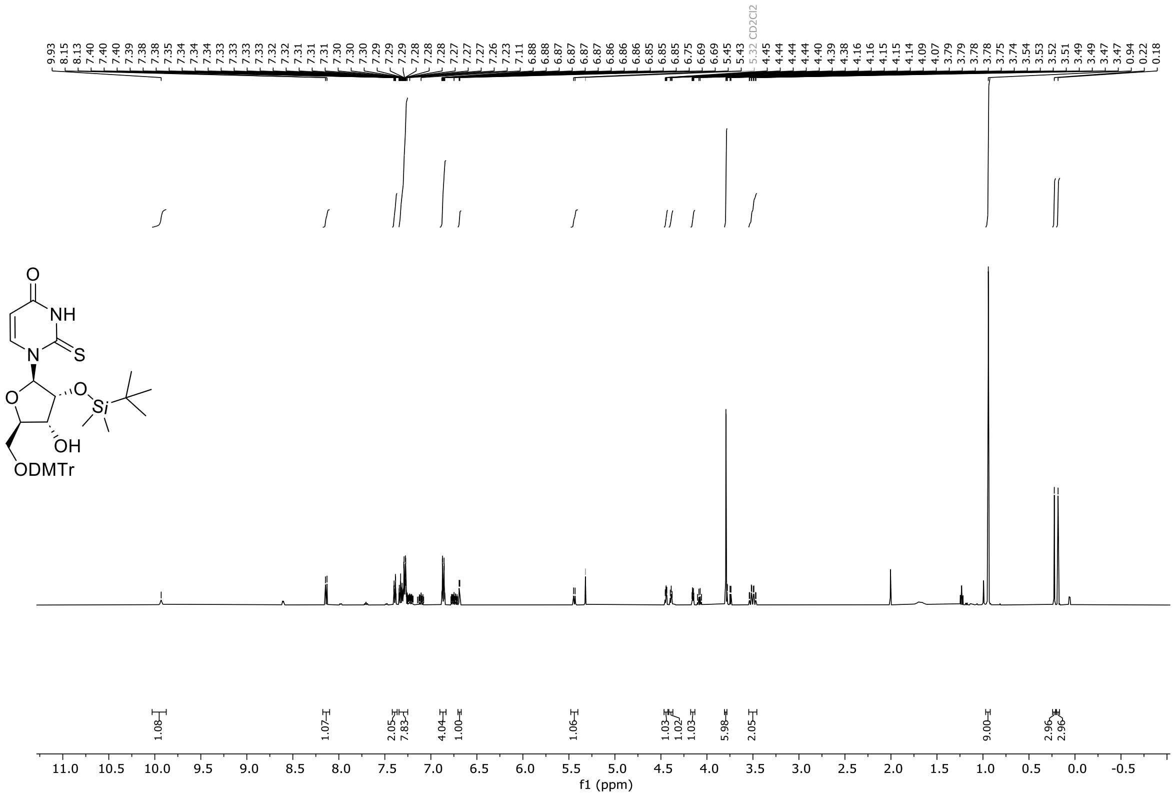


#### $^{31}$P NMR spectrum of compound 12

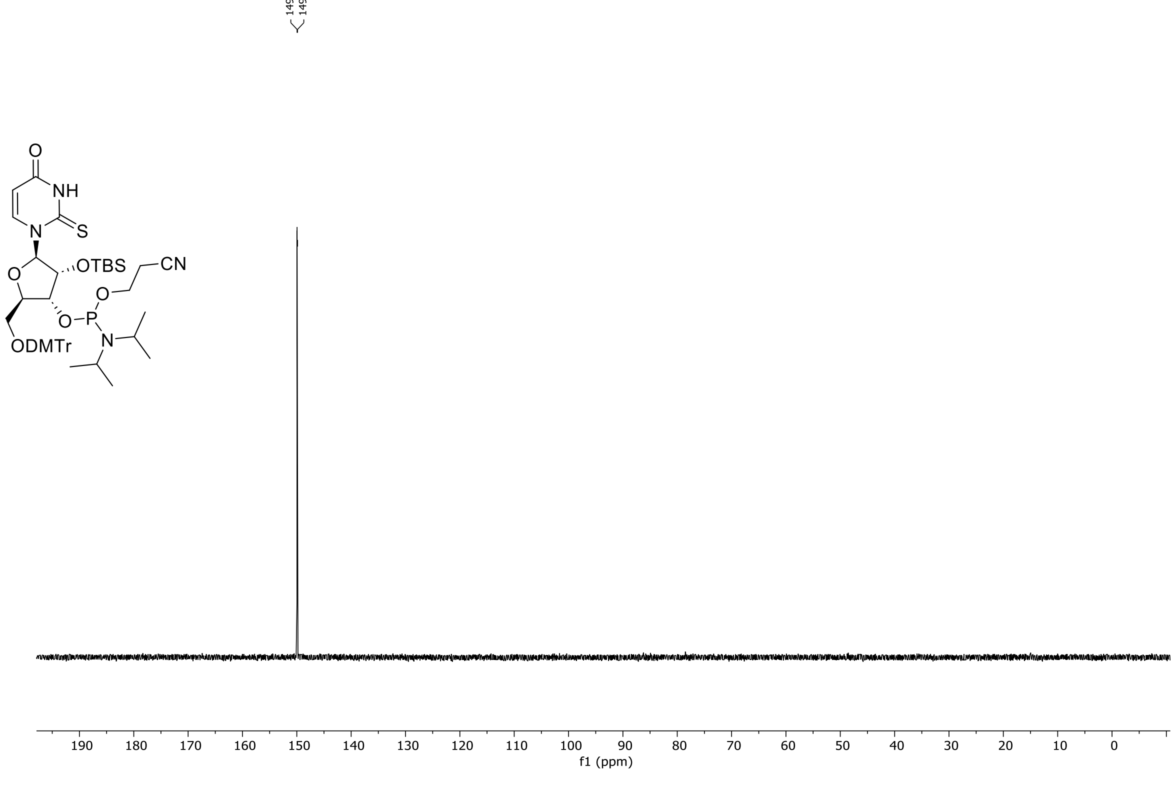

#### $^{31}$P NMR spectrum of compound 12b

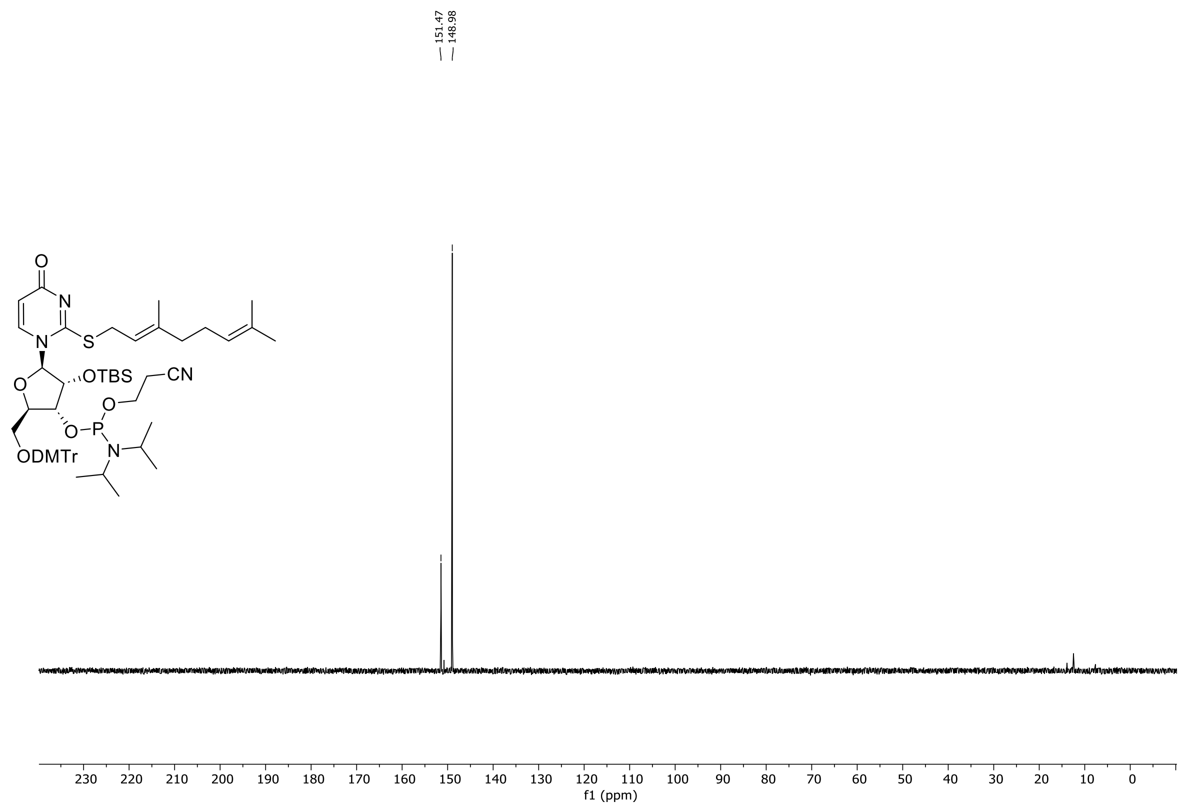


#### $^{1}$H NMR spectrum of compound 14

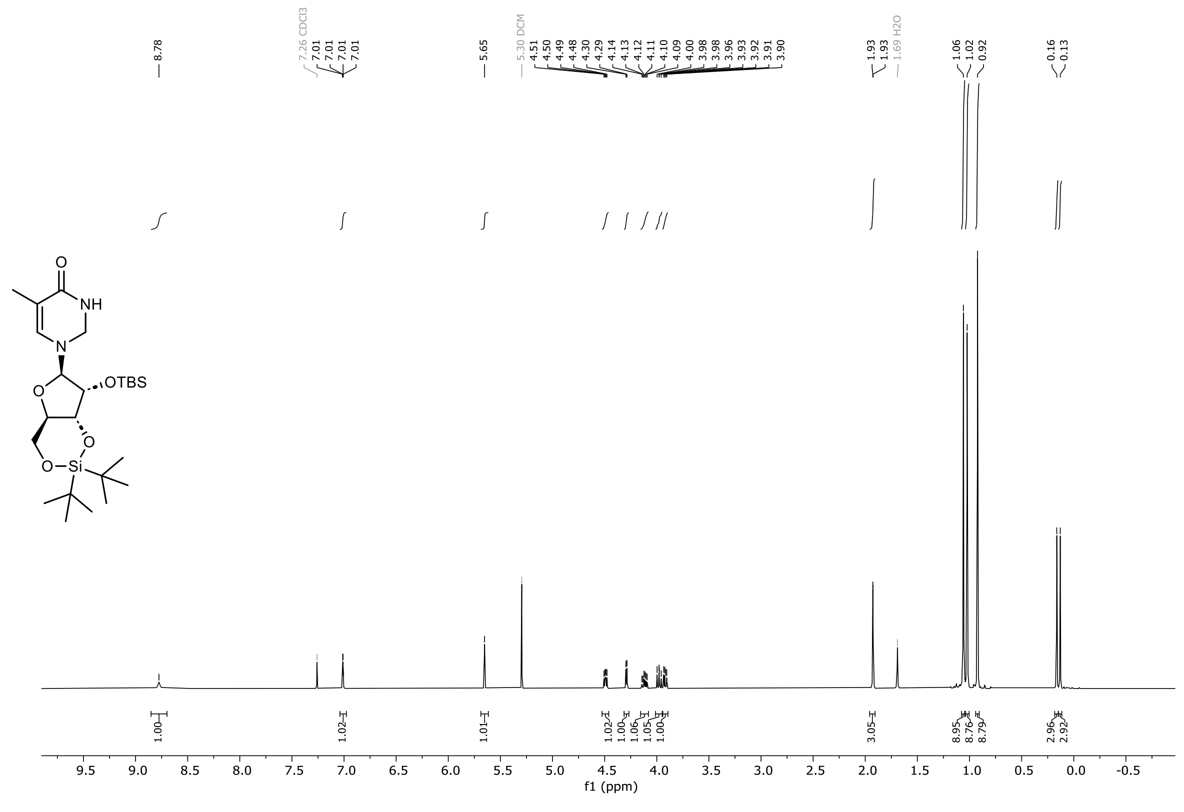

#### $^1$H NMR spectrum of compound 15

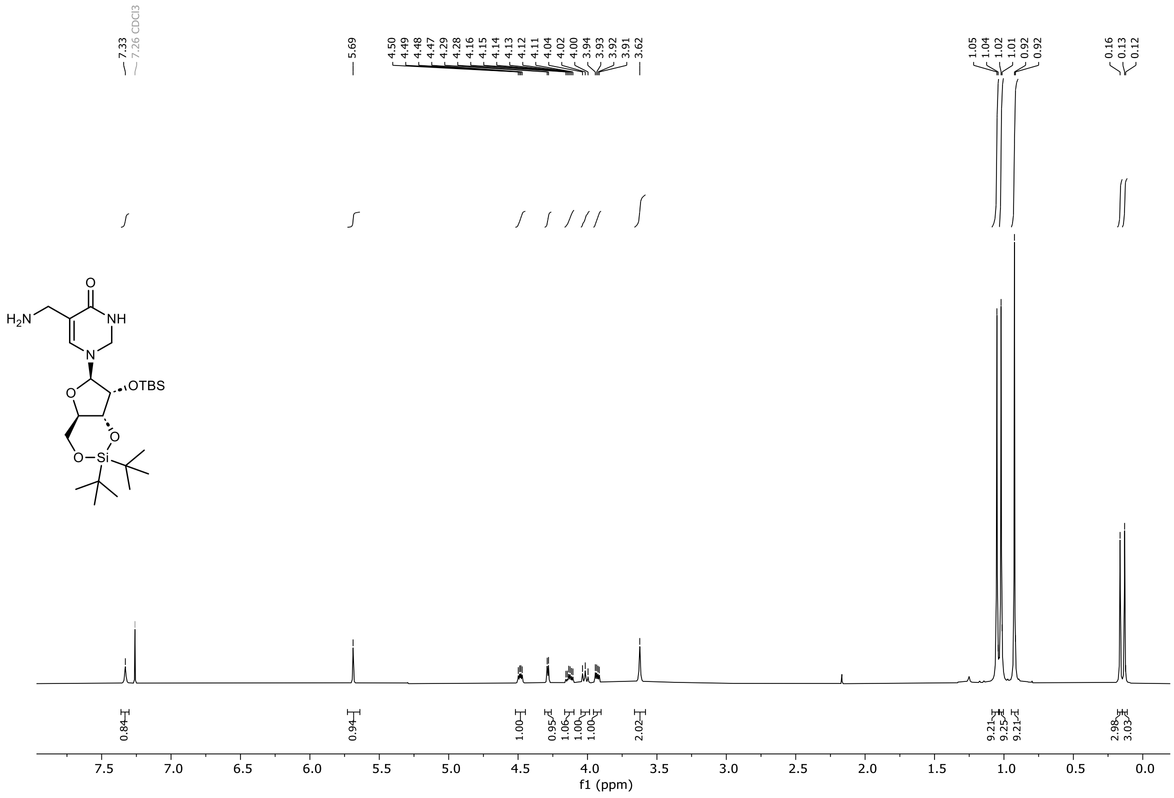


#### $^1$H NMR spectrum of compound 16

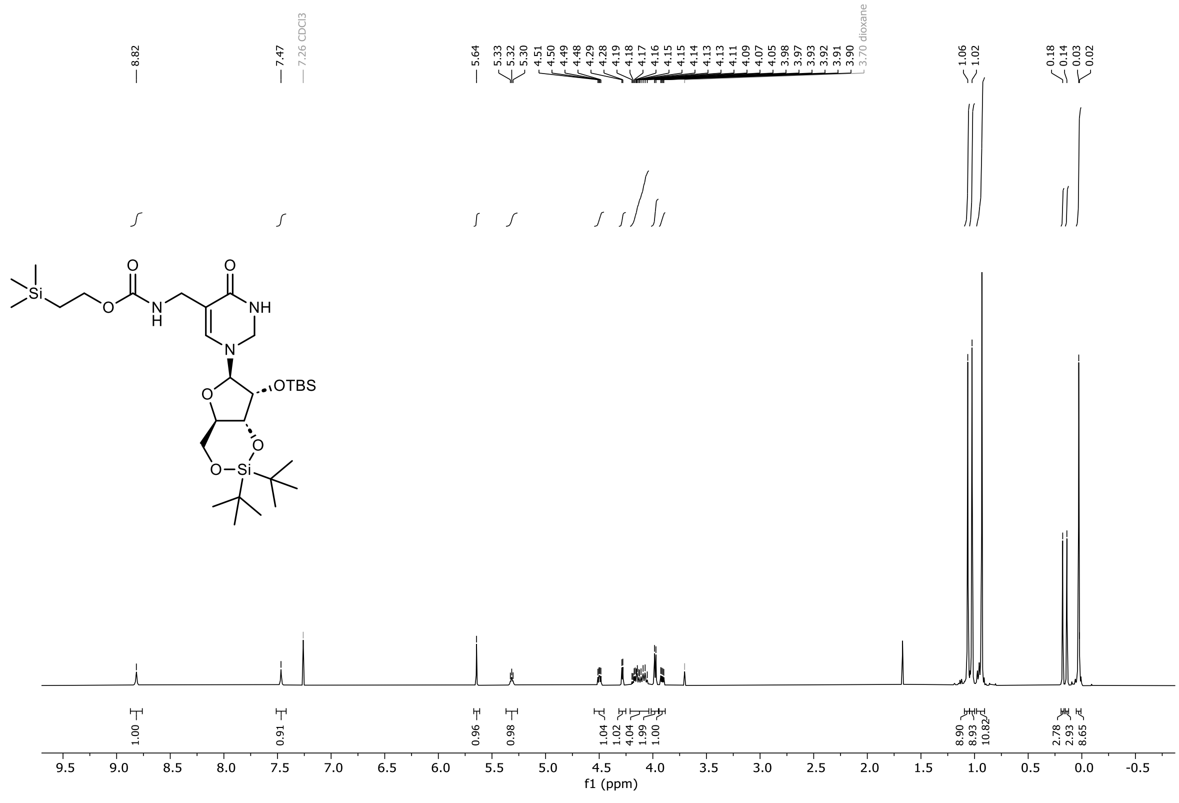

### $^1$H NMR spectrum of compound 17

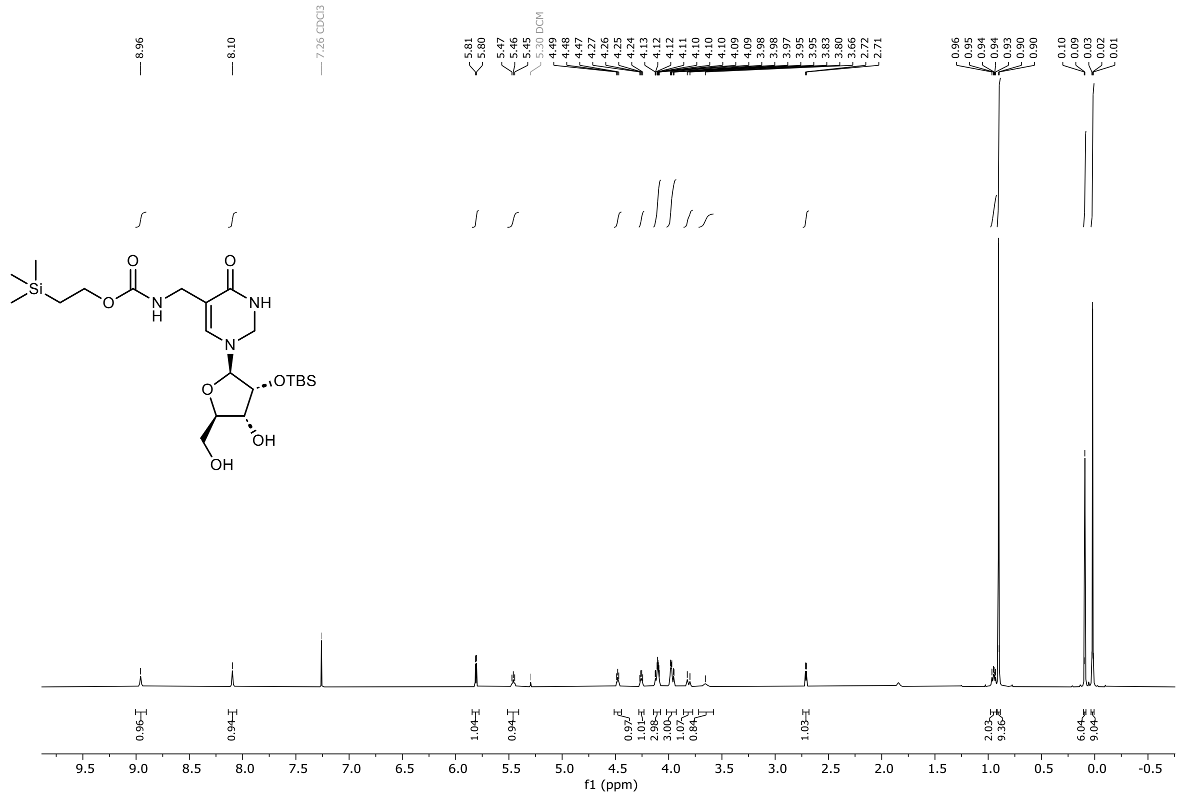

### $^1$H NMR spectrum of compound 18

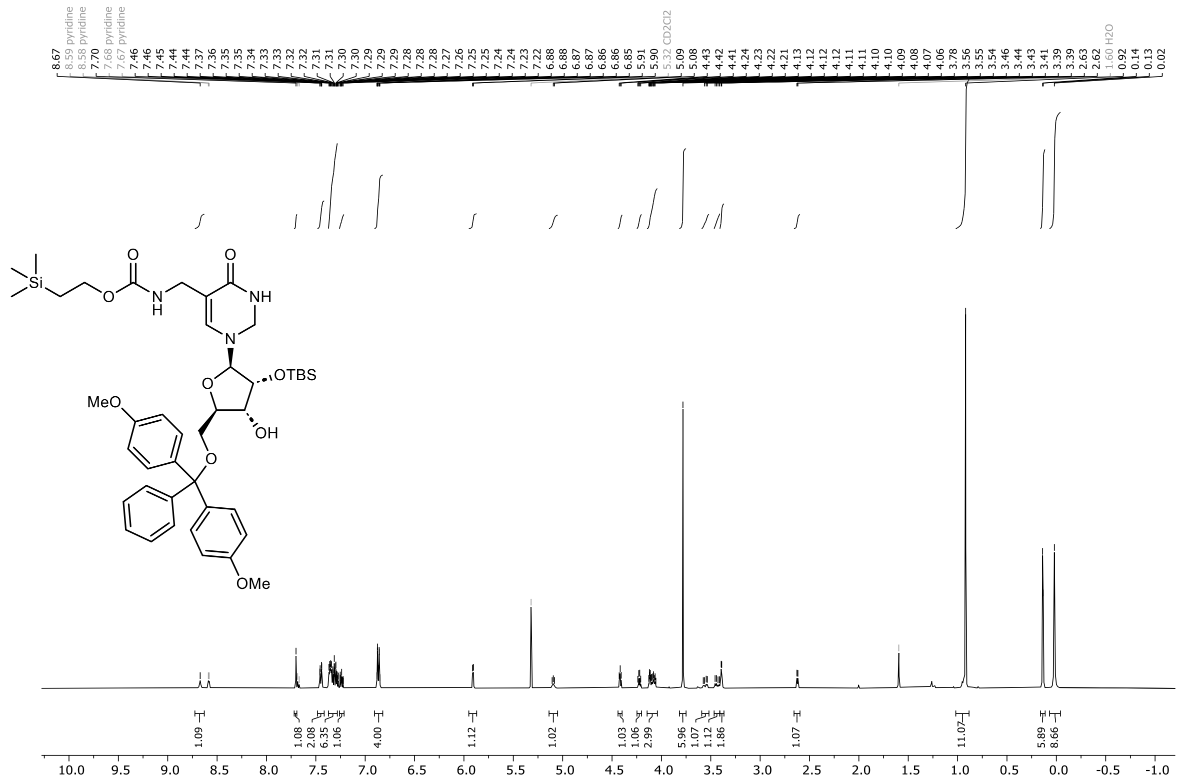

#### $^{31}$P NMR spectrum of compound 19

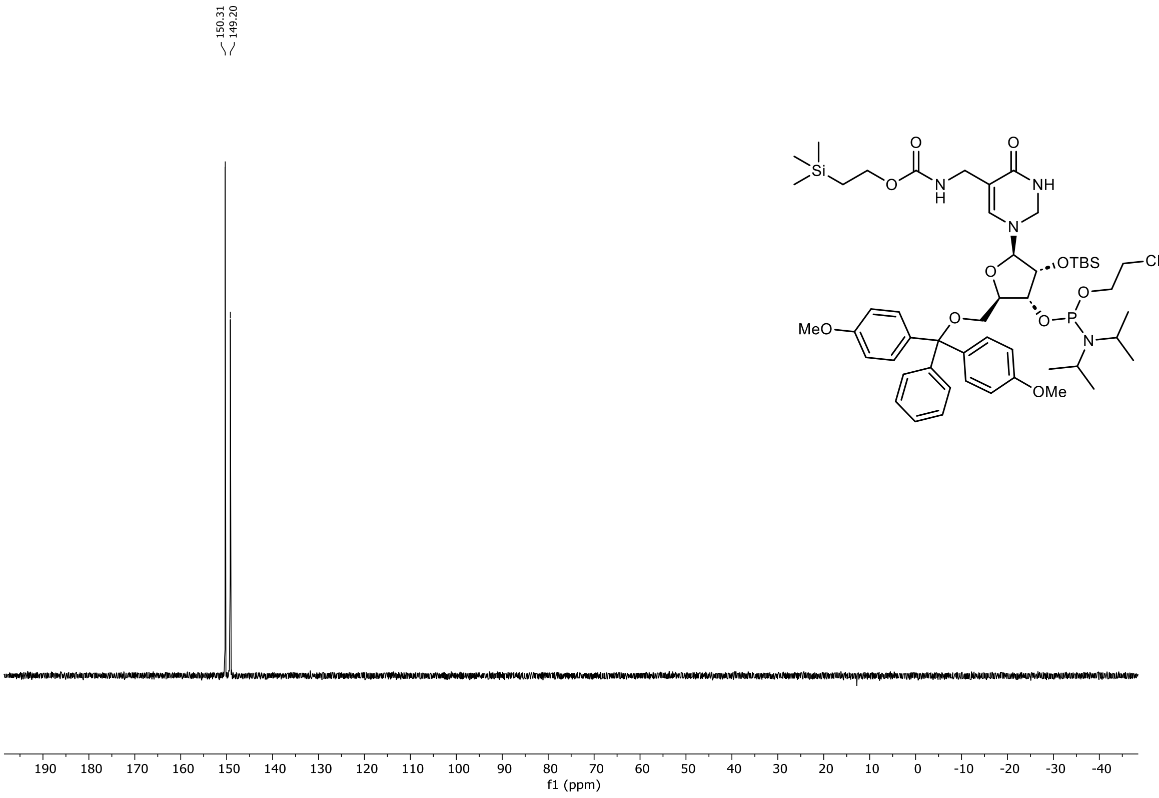


#### $^{1}$H NMR spectrum of compound 21

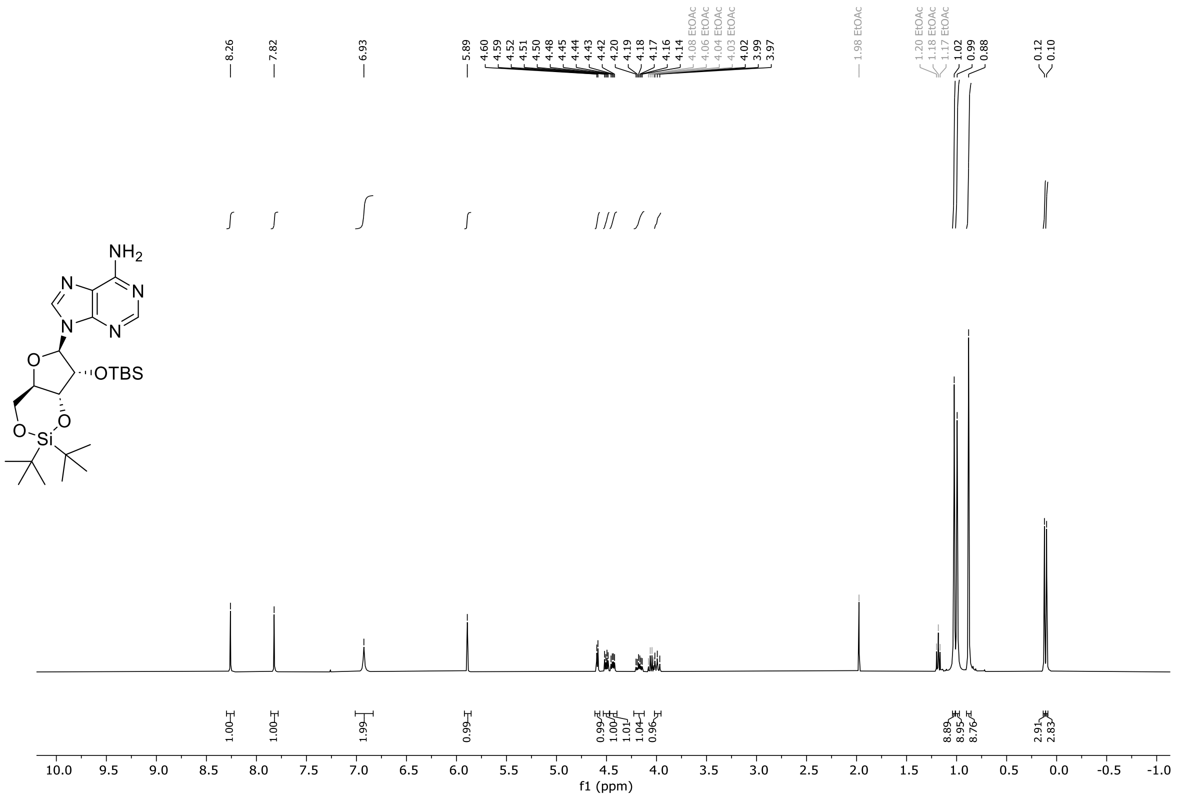

#### $^1$H NMR spectrum of compound 22

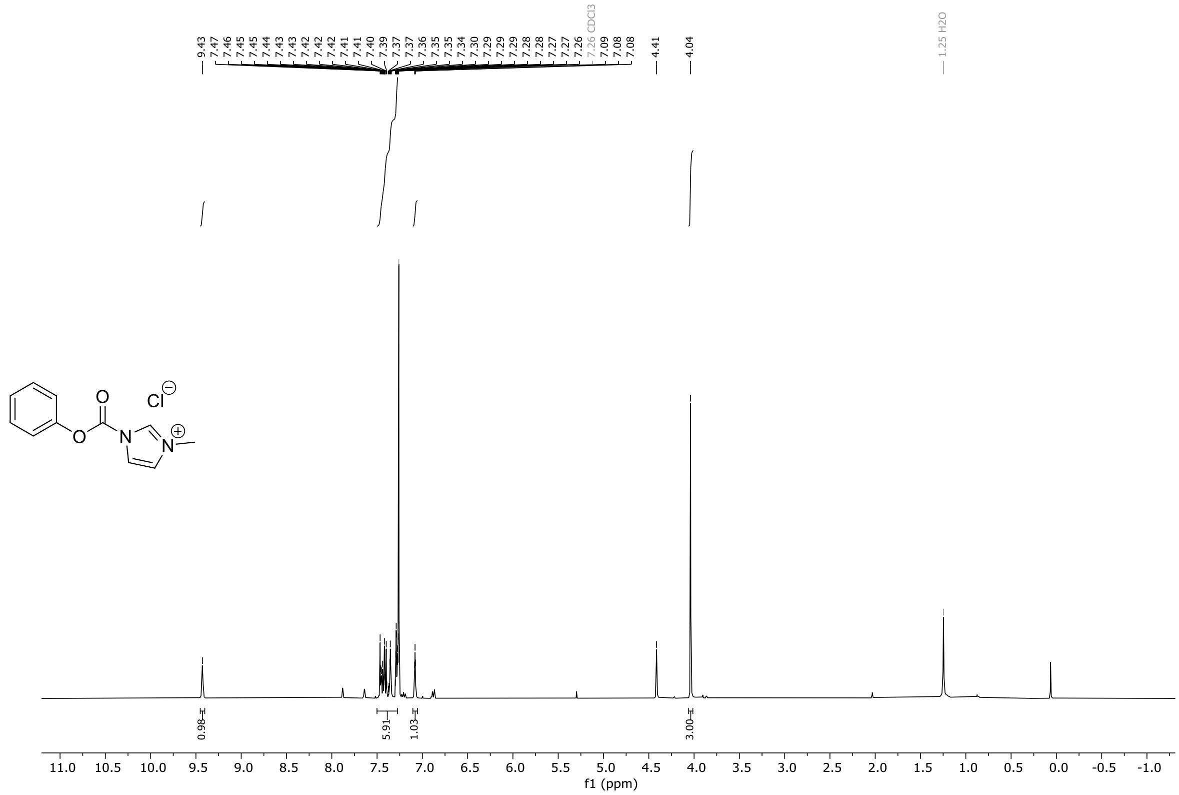


#### $^1$H NMR spectrum of compound 23a

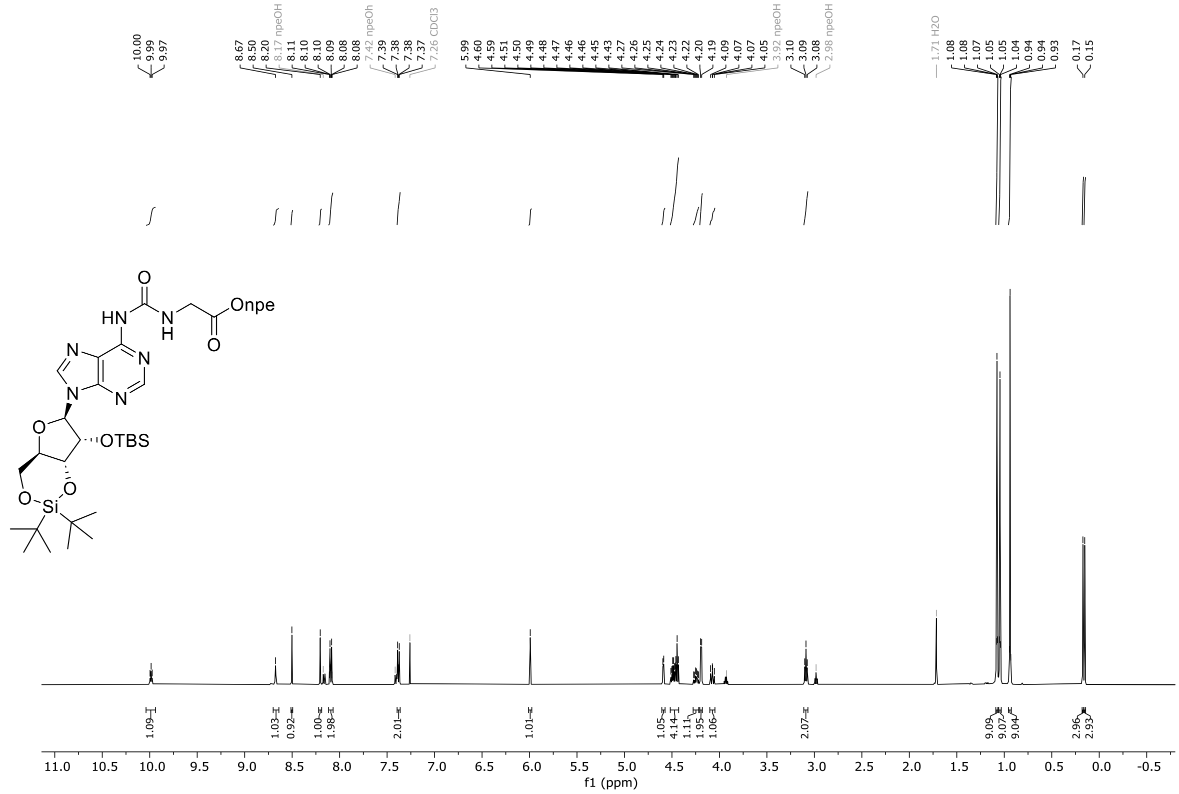

#### $^1$H NMR spectrum of compound 23b

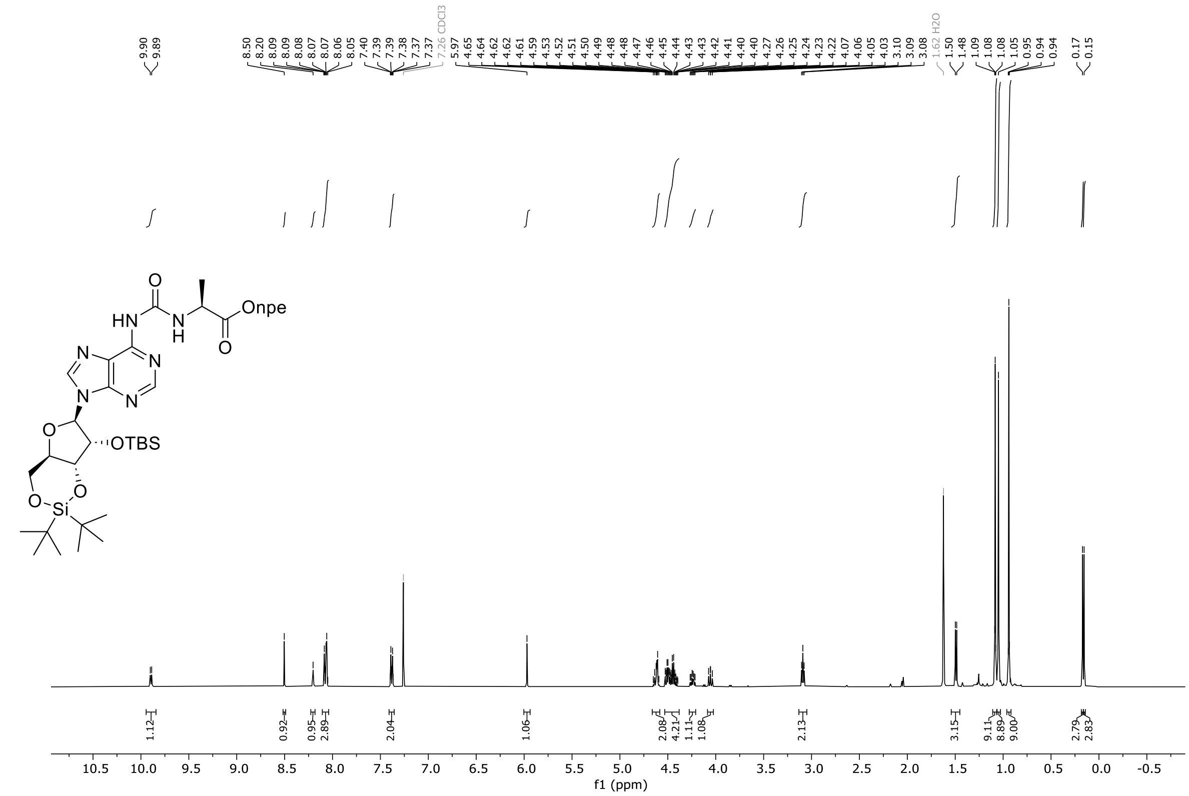


#### $^1$H NMR spectrum of compound 23c

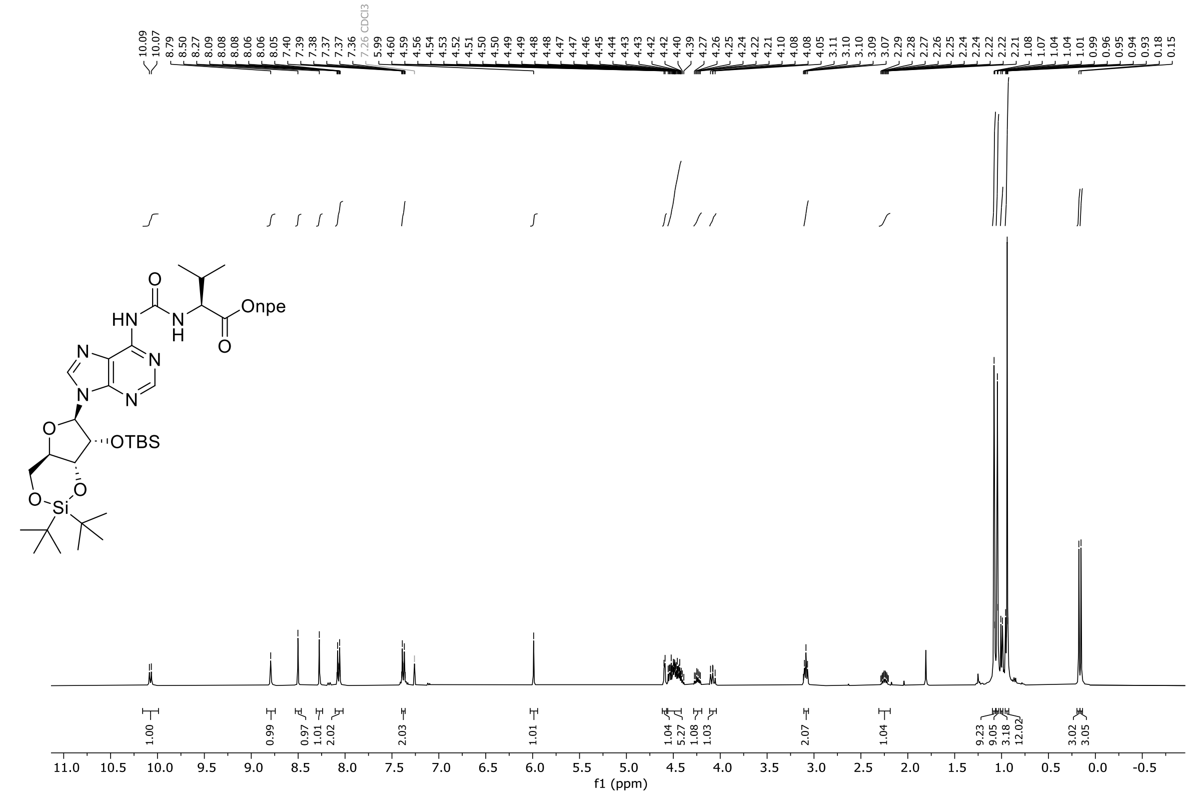

#### $^1$H NMR spectrum of compound 23d

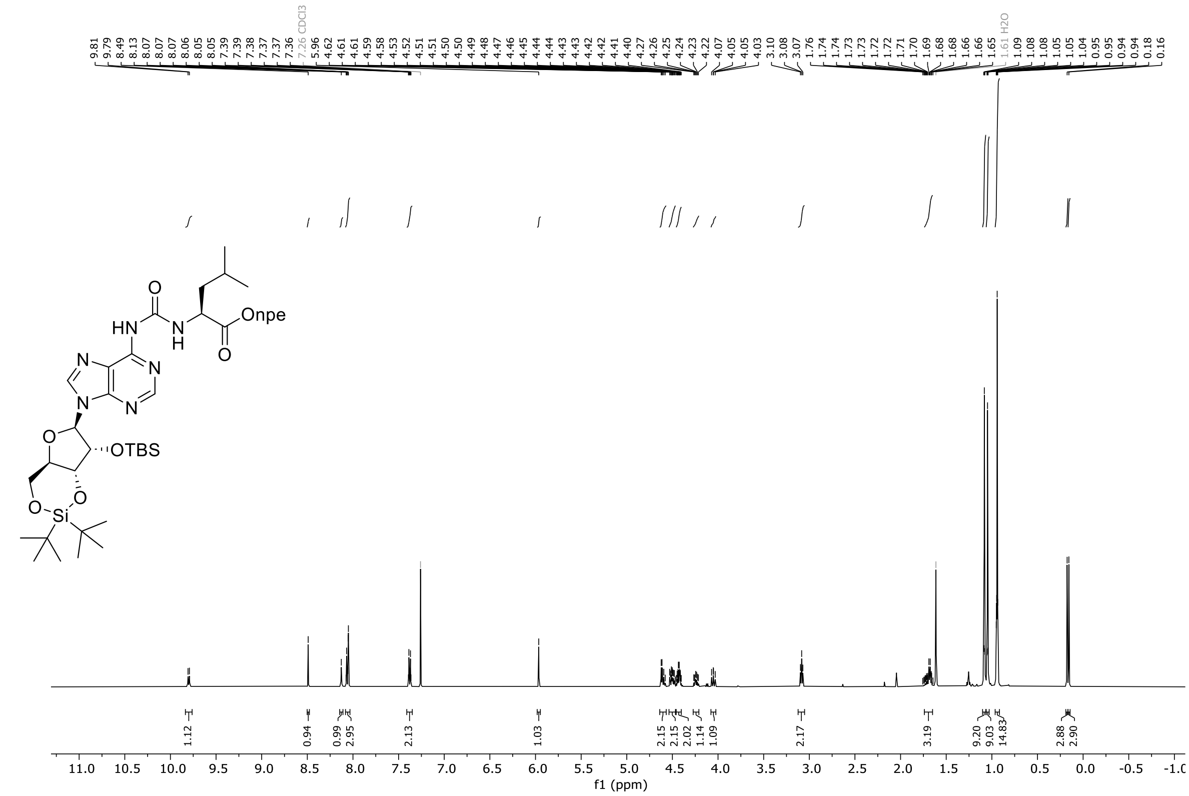

#### $^1$H NMR spectrum of compound 23e

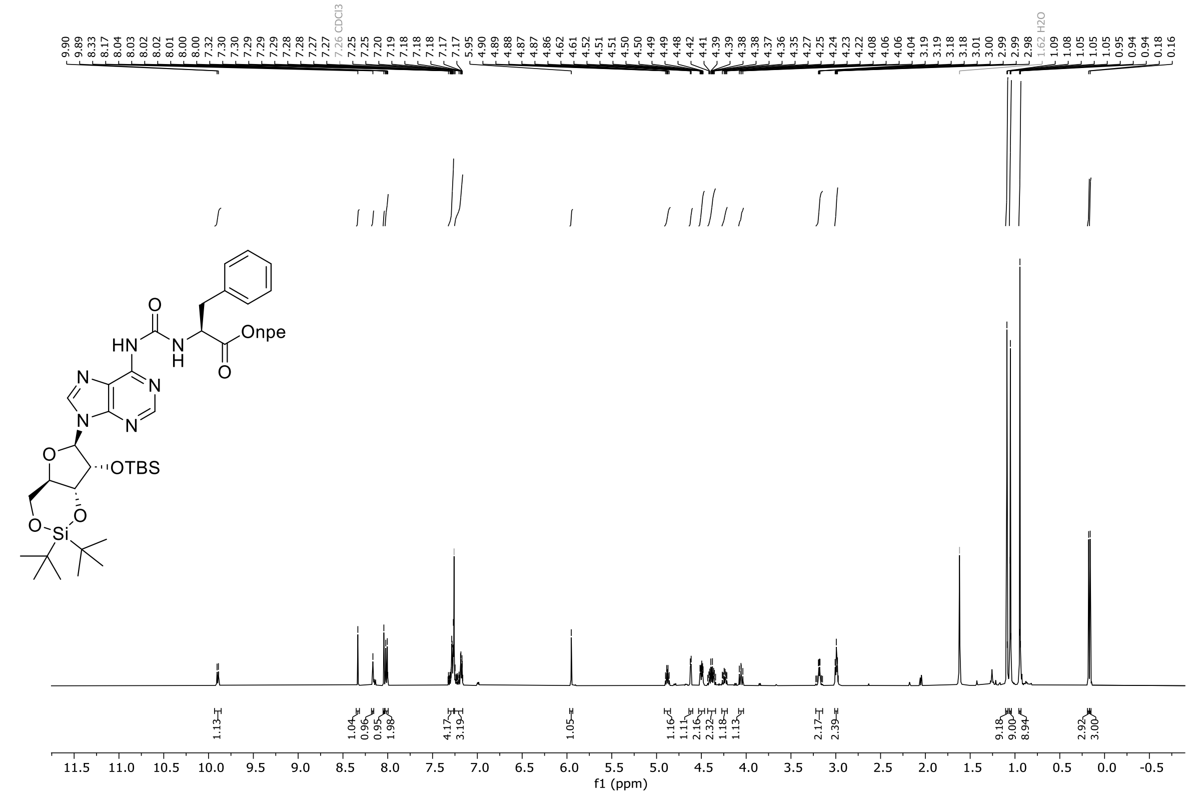

#### $^1$H NMR spectrum of compound 24a

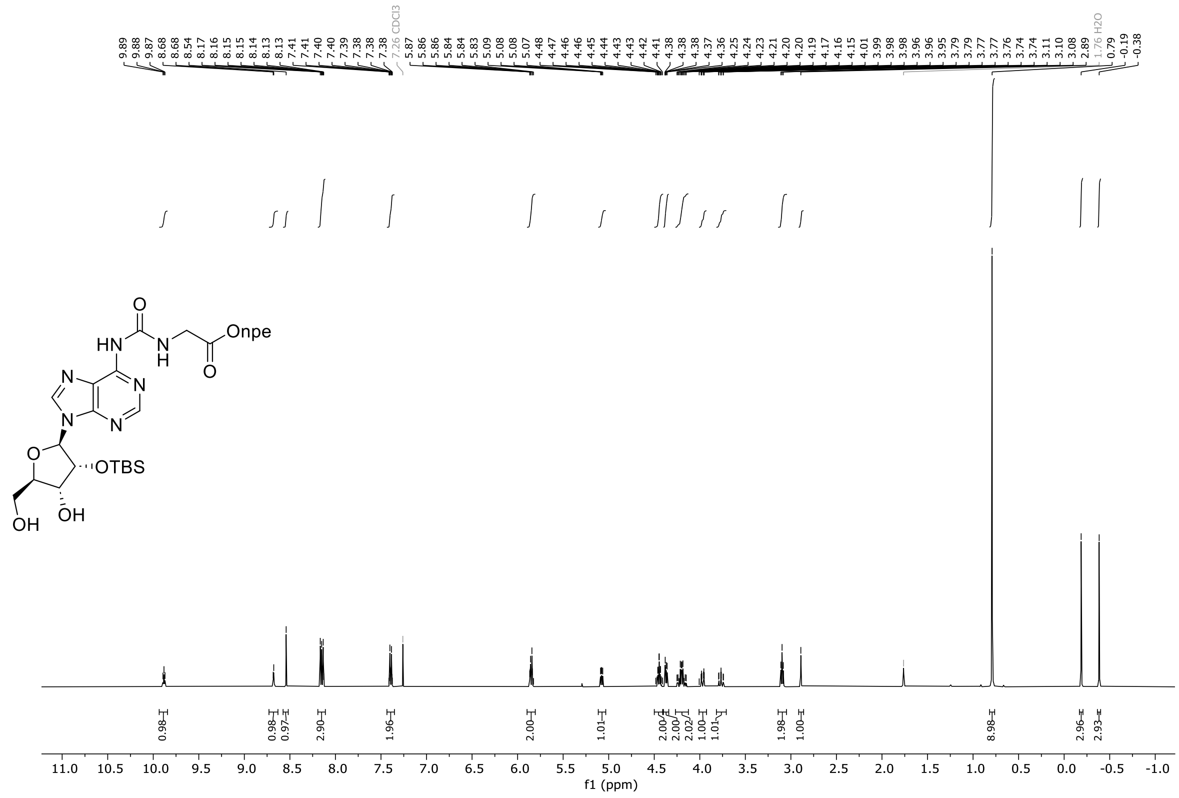

#### $^1$H NMR spectrum of compound 24b

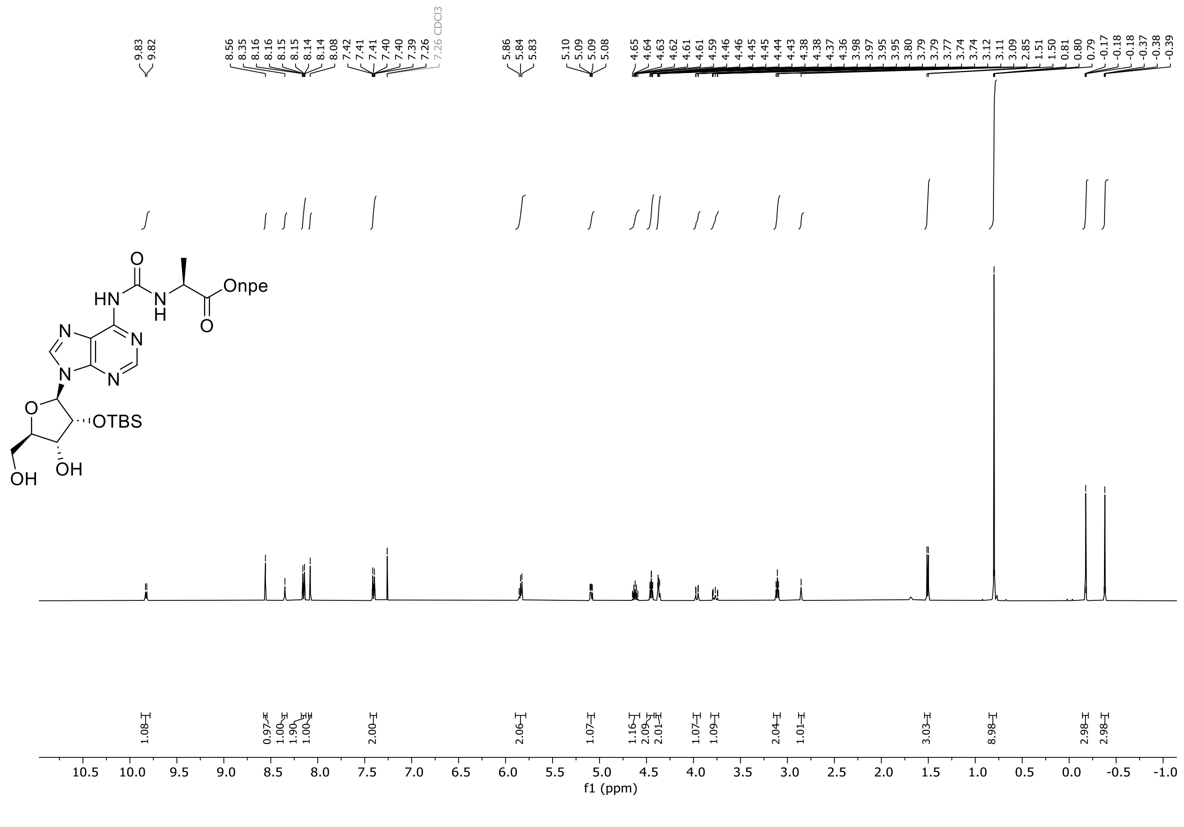

##### $^{13}$C NMR spectrum of compound 24b

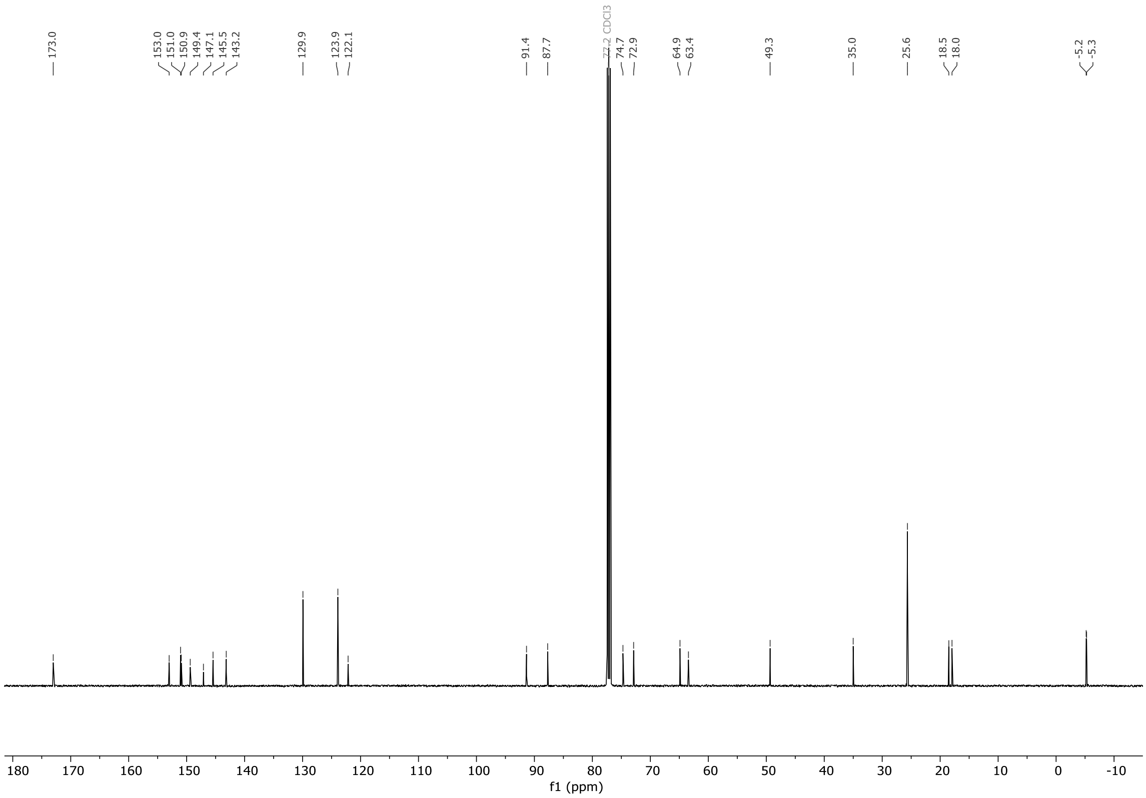

##### $^{1}$H NMR spectrum of compound 24c

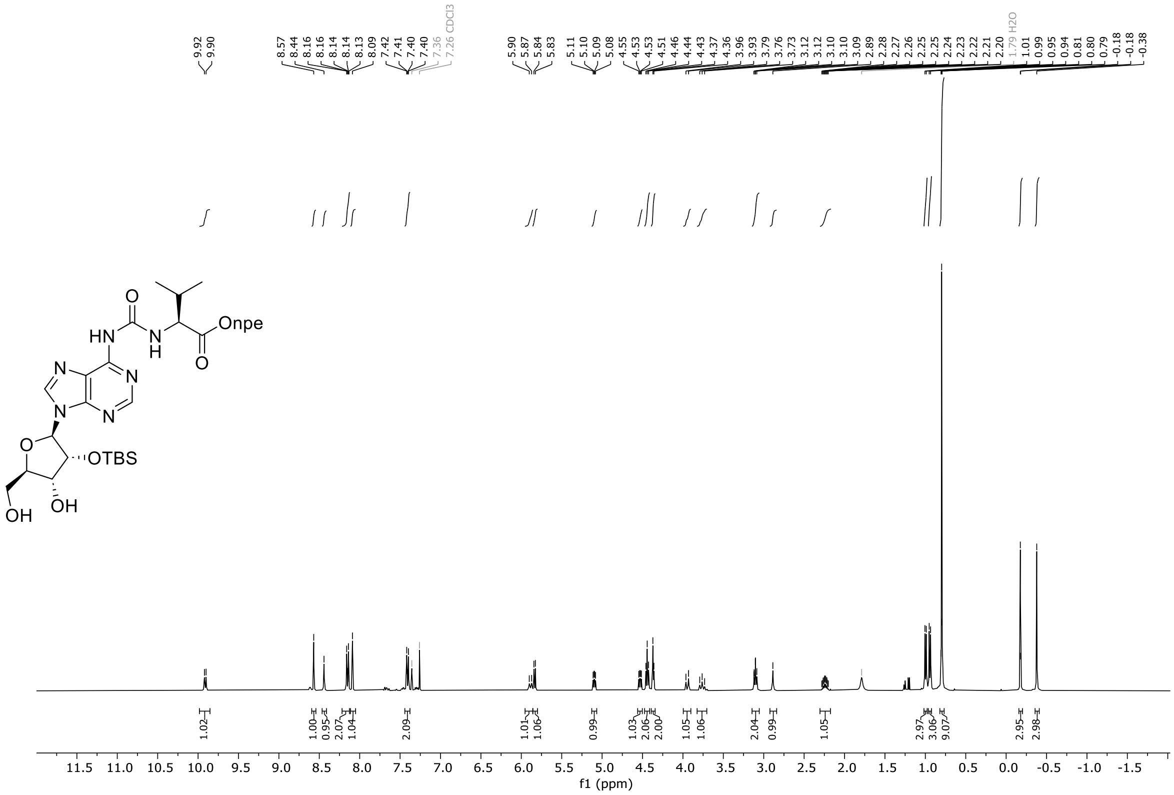


##### $^{1}$H NMR spectrum of compound 24d

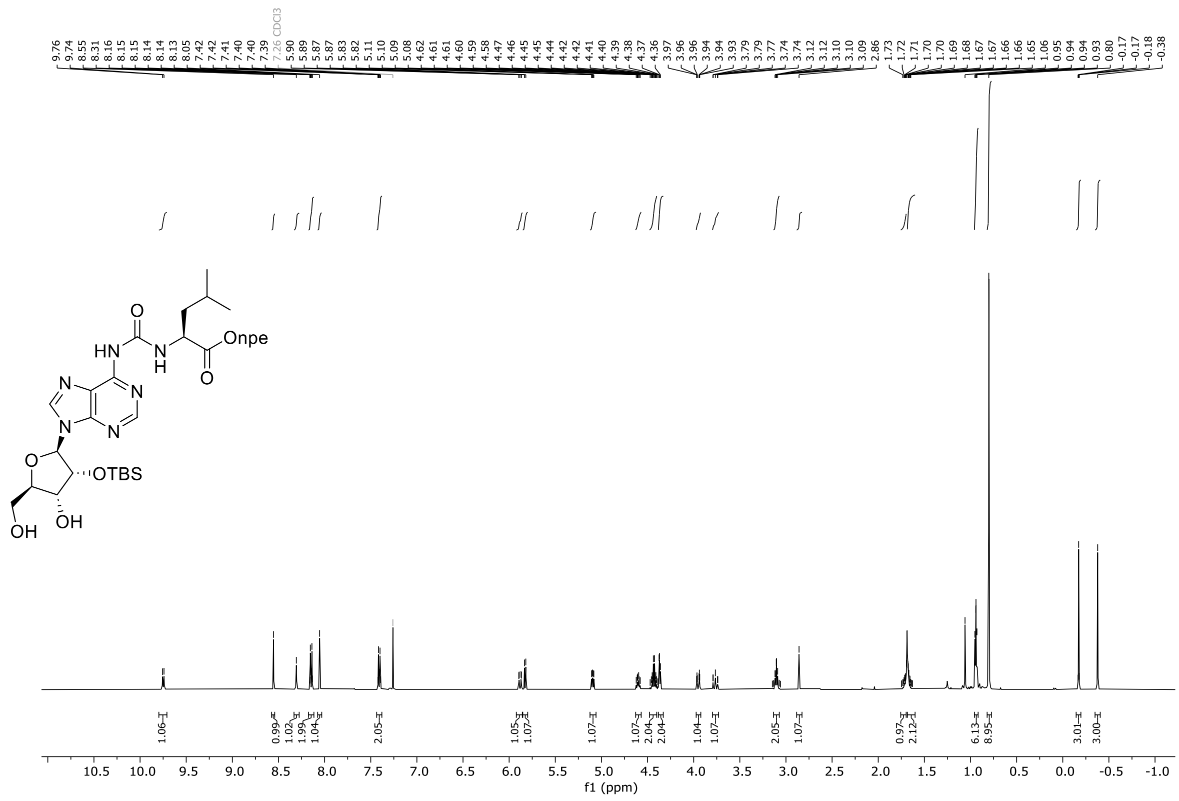


#### $^{13}$C NMR spectrum of compound 24d

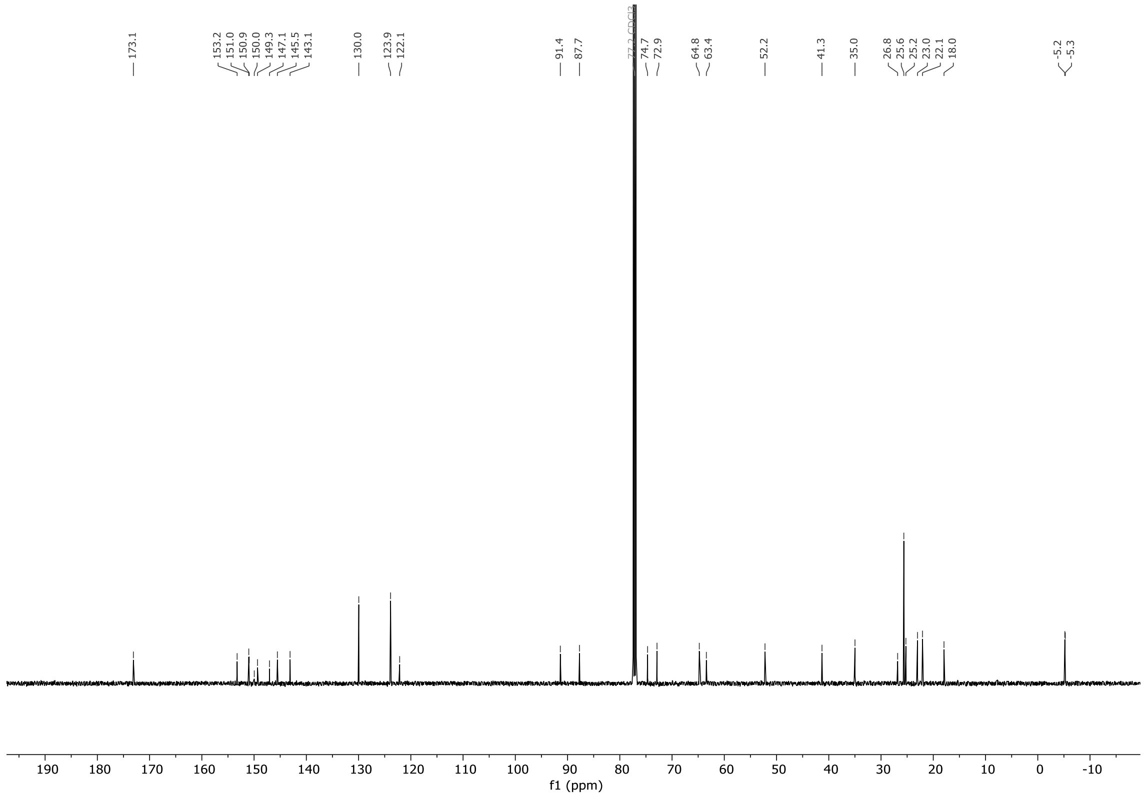


#### $^{1}$H NMR spectrum of compound 24e

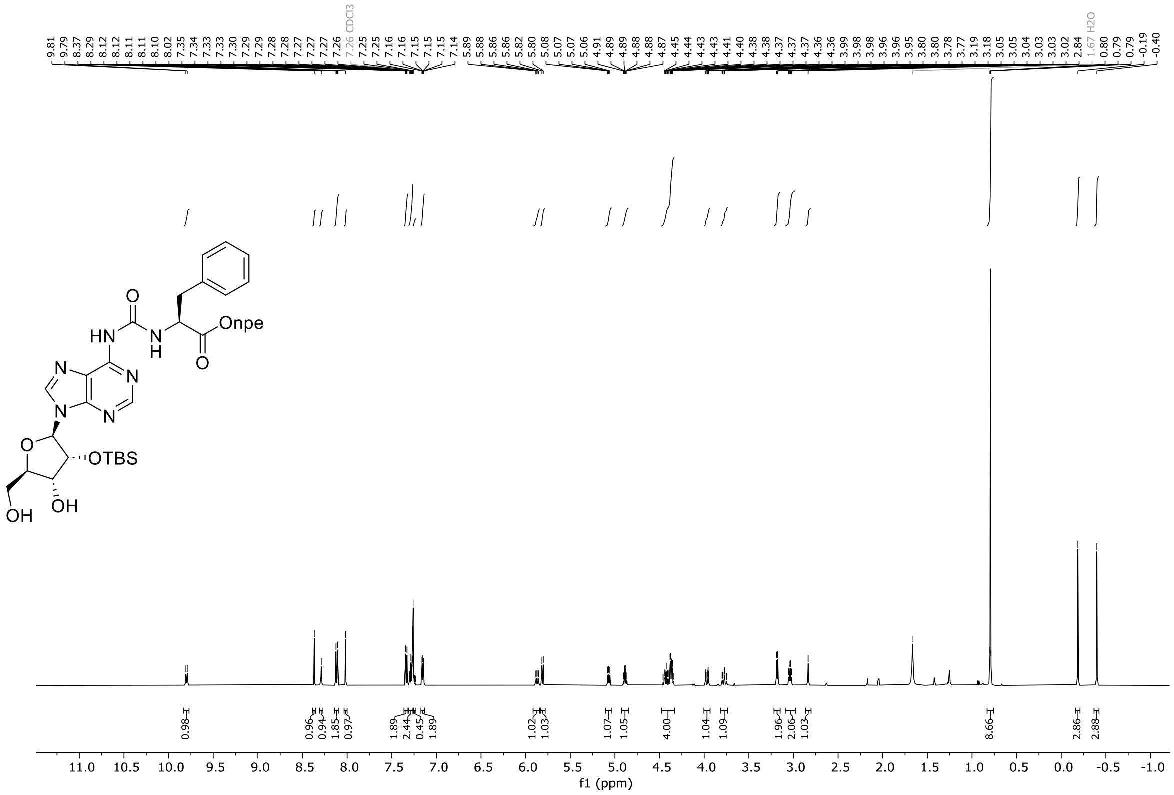


**$^{1}$H NMR spectrum of compound 25a**

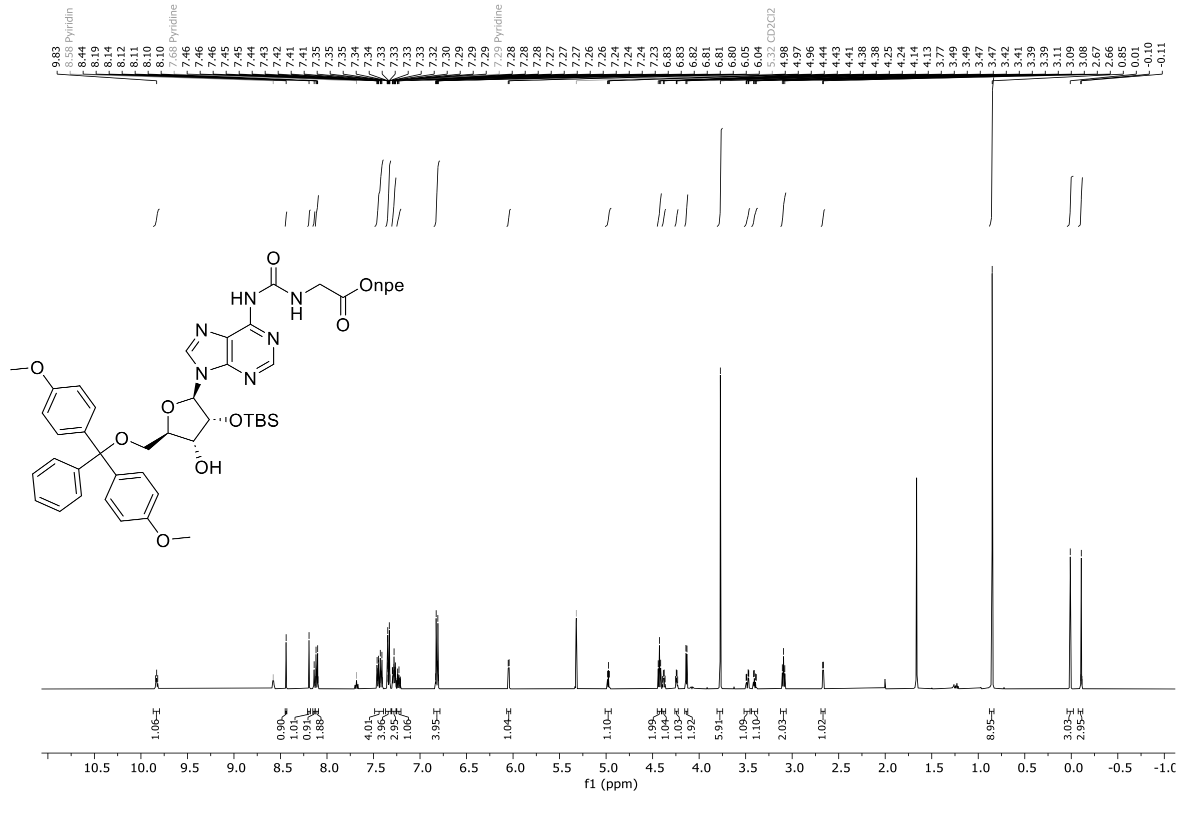


**$^{1}$H NMR spectrum of compound 25b**

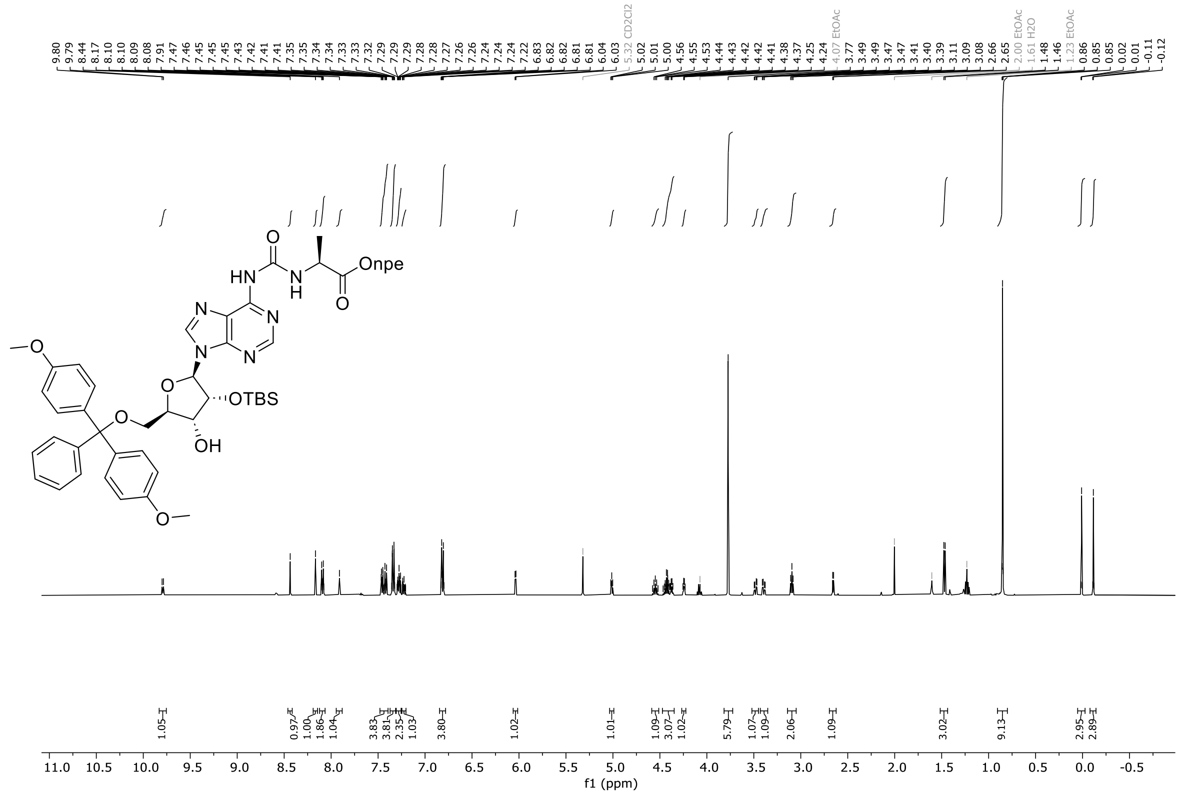


#### $^{13}$C NMR spectrum of compound 25b

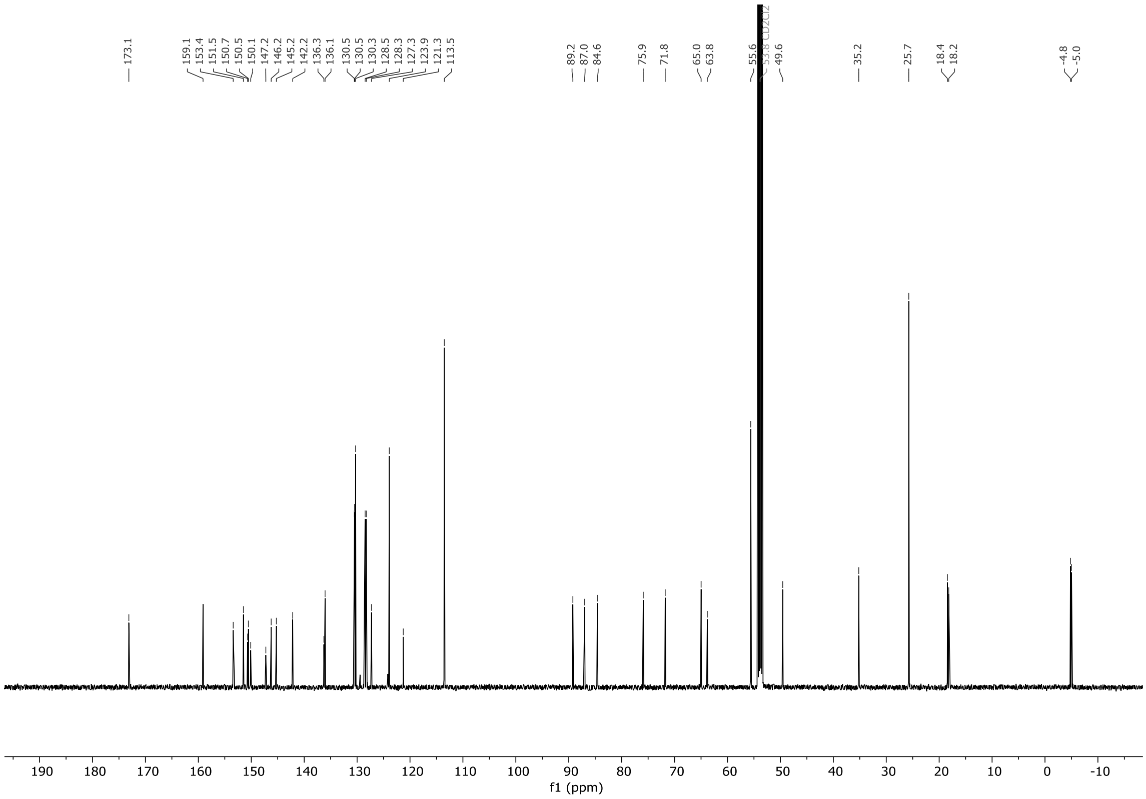


#### $^{1}$H NMR spectrum of compound 25c

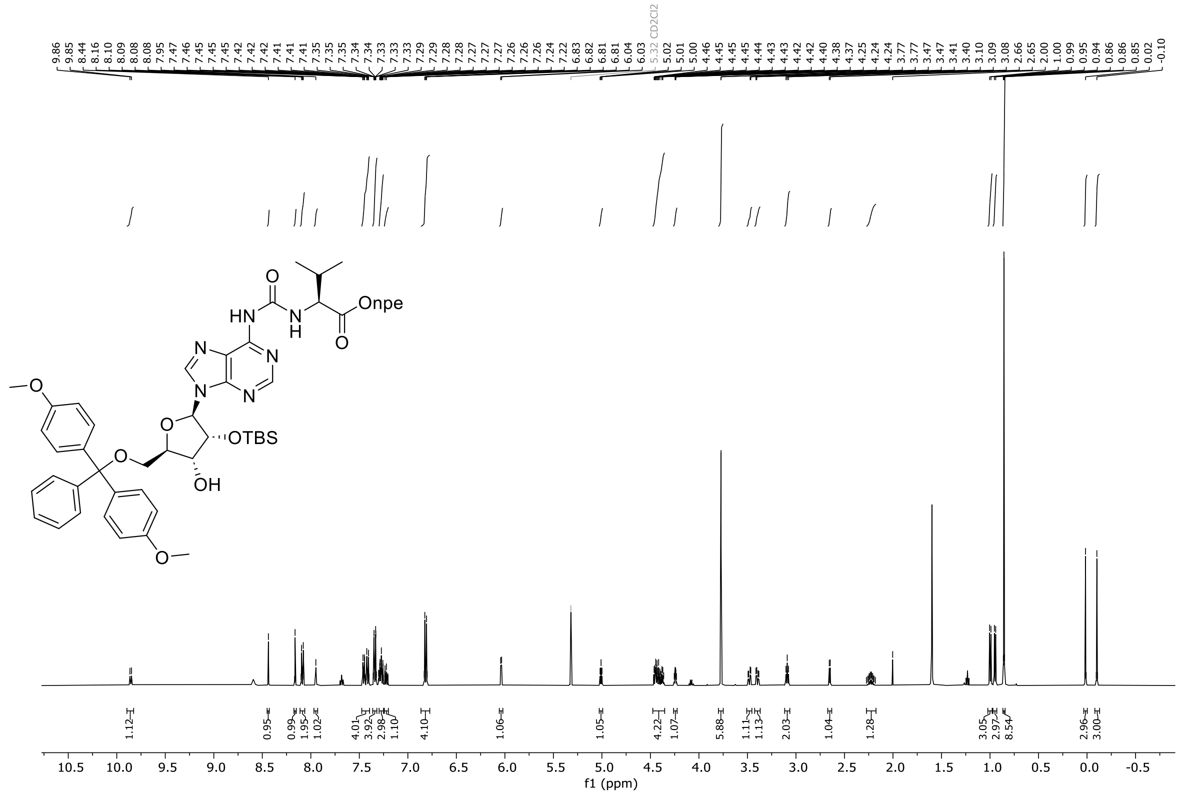


#### $^{1}$H NMR spectrum of compound 25d

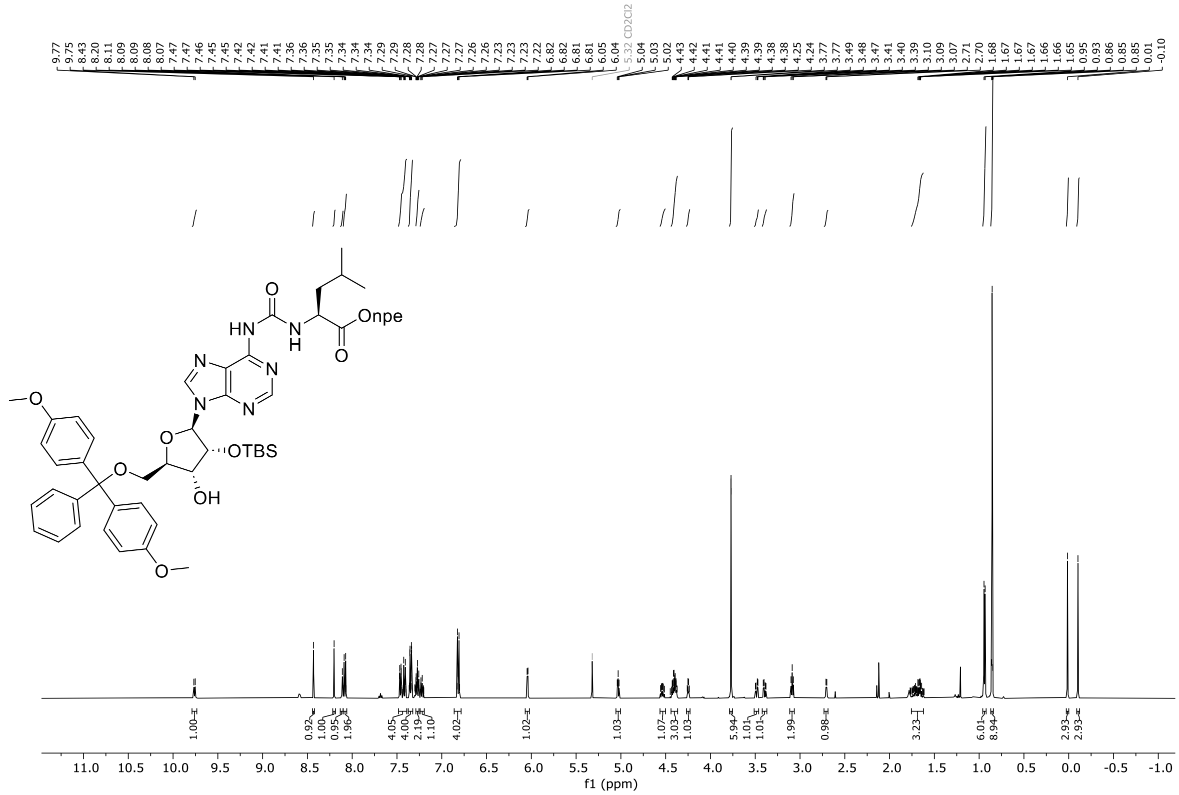


#### $^{13}$C NMR spectrum of compound 25d

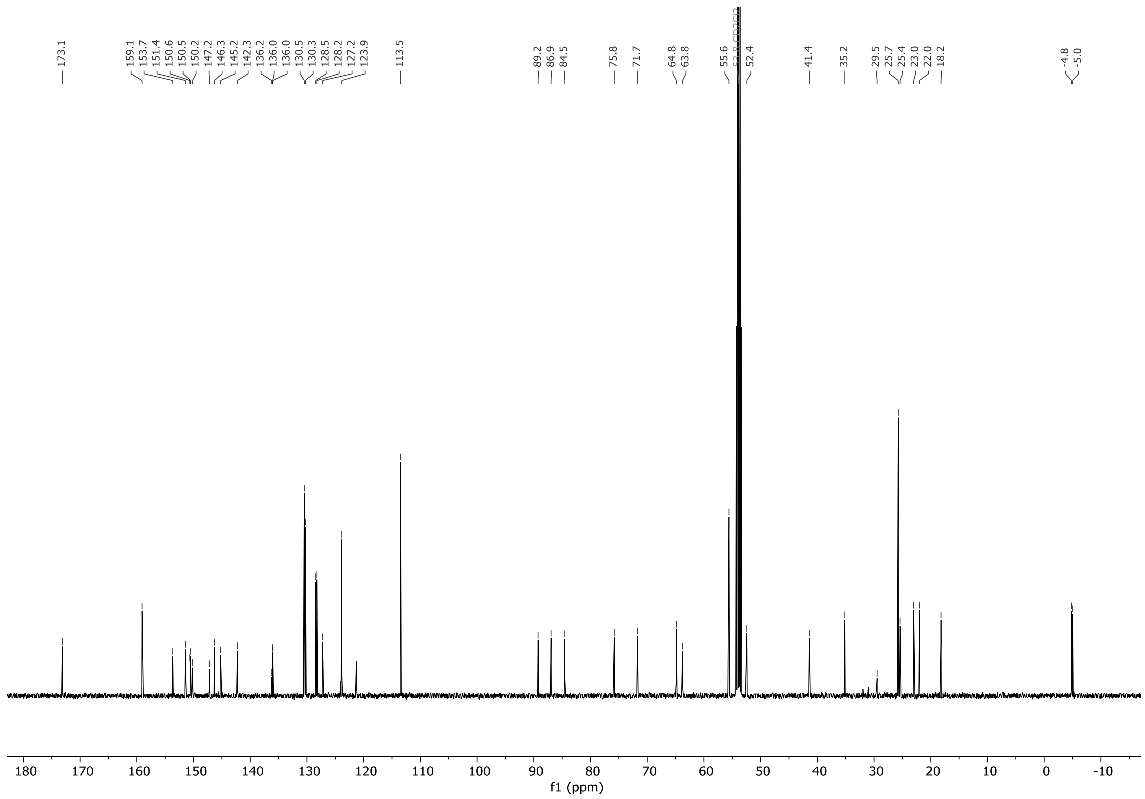


#### $^{1}$H NMR spectrum of compound 25e

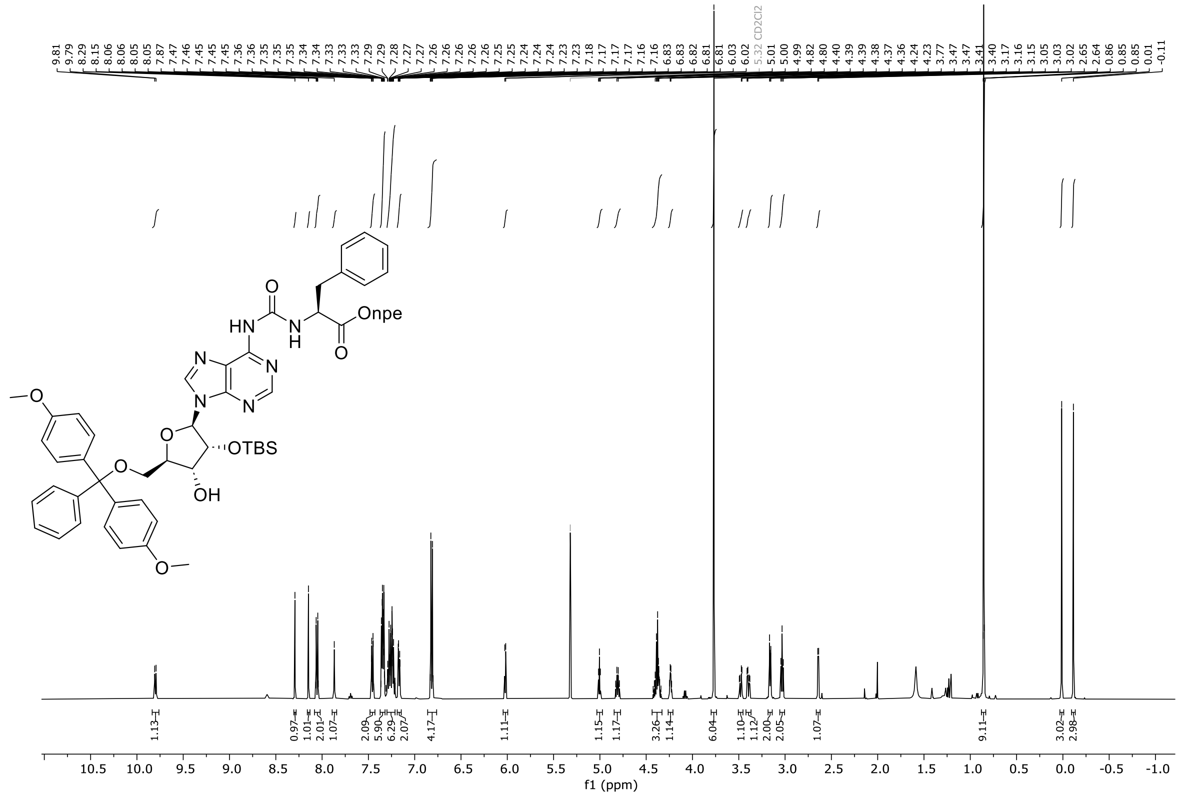


#### $^{31}$P NMR spectrum of compound 26a

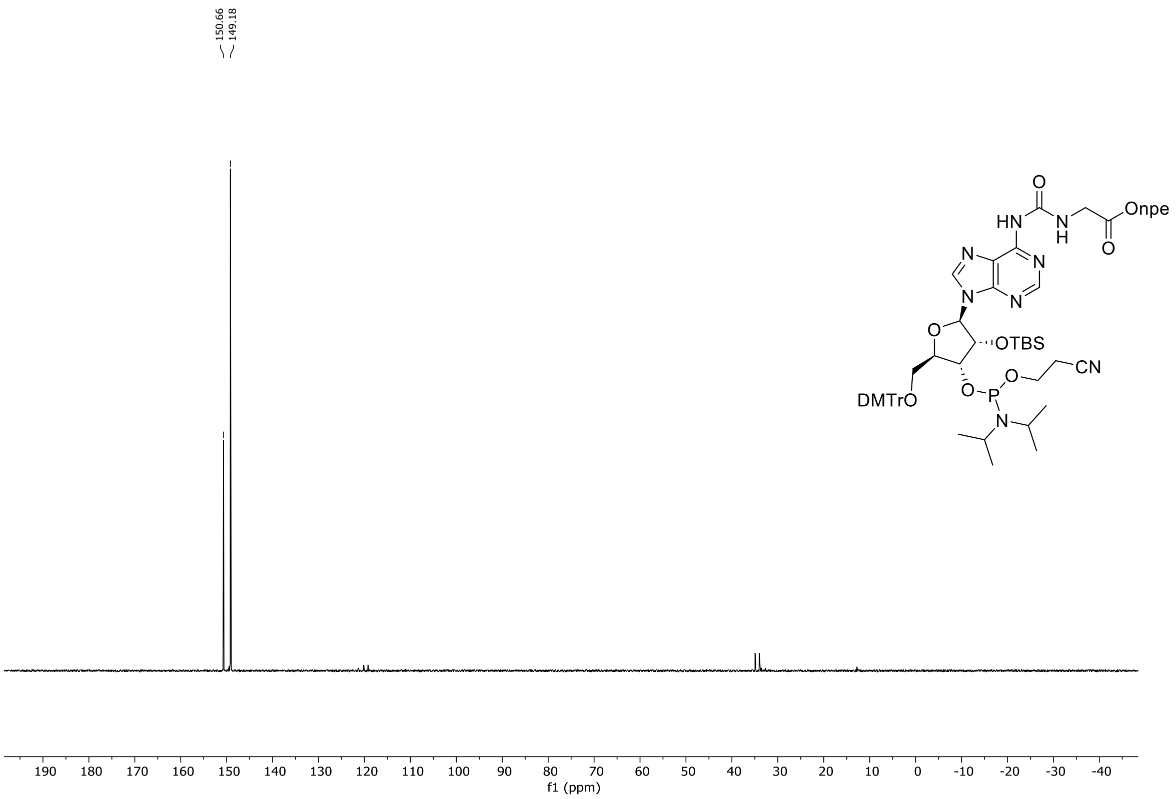


**$^{31}$P NMR spectrum of compound 26b**

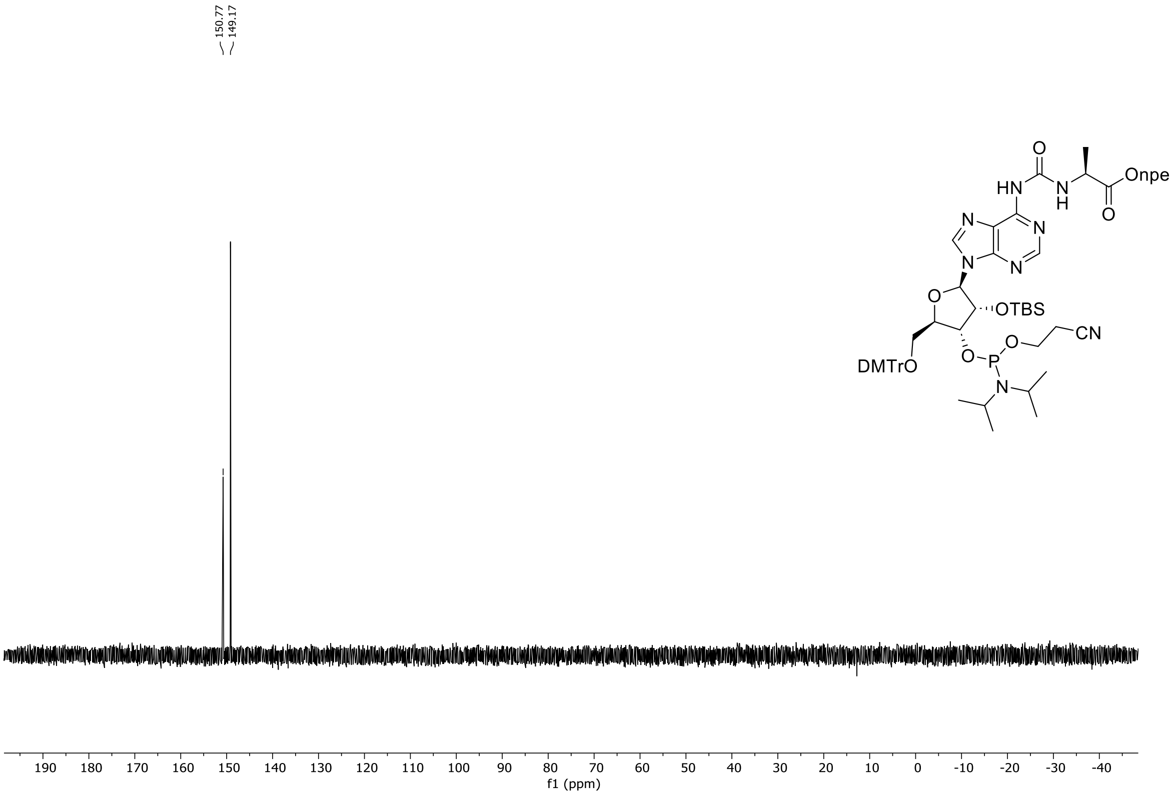


**$^{31}$P NMR spectrum of compound 26c**

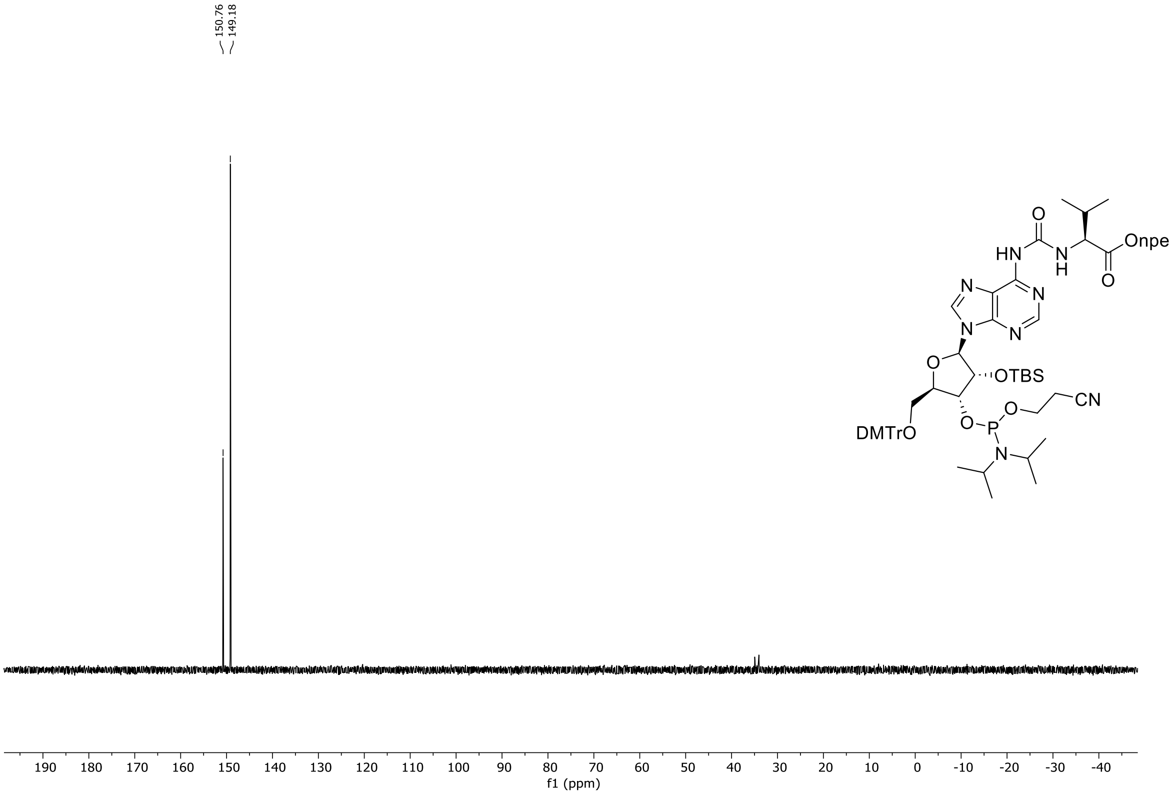


#### $^{31}$P NMR spectrum of compound 26d

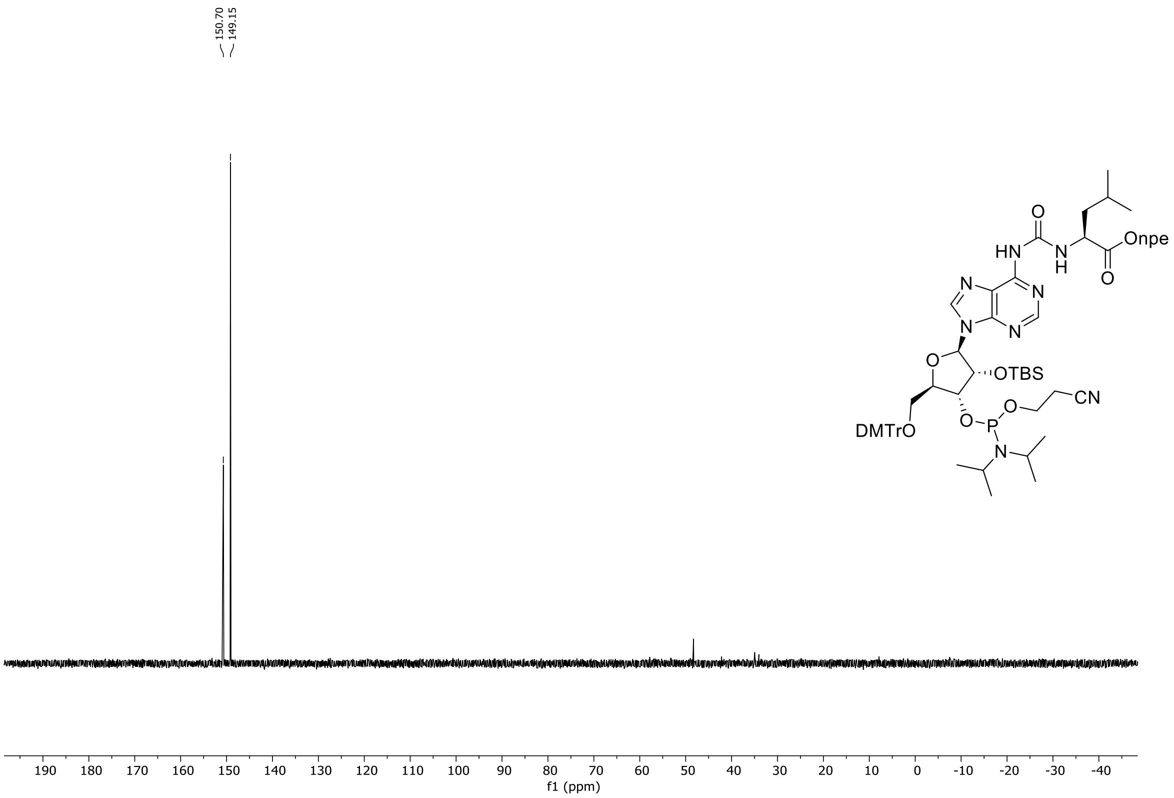


#### $^{31}$P NMR spectrum of compound 26e

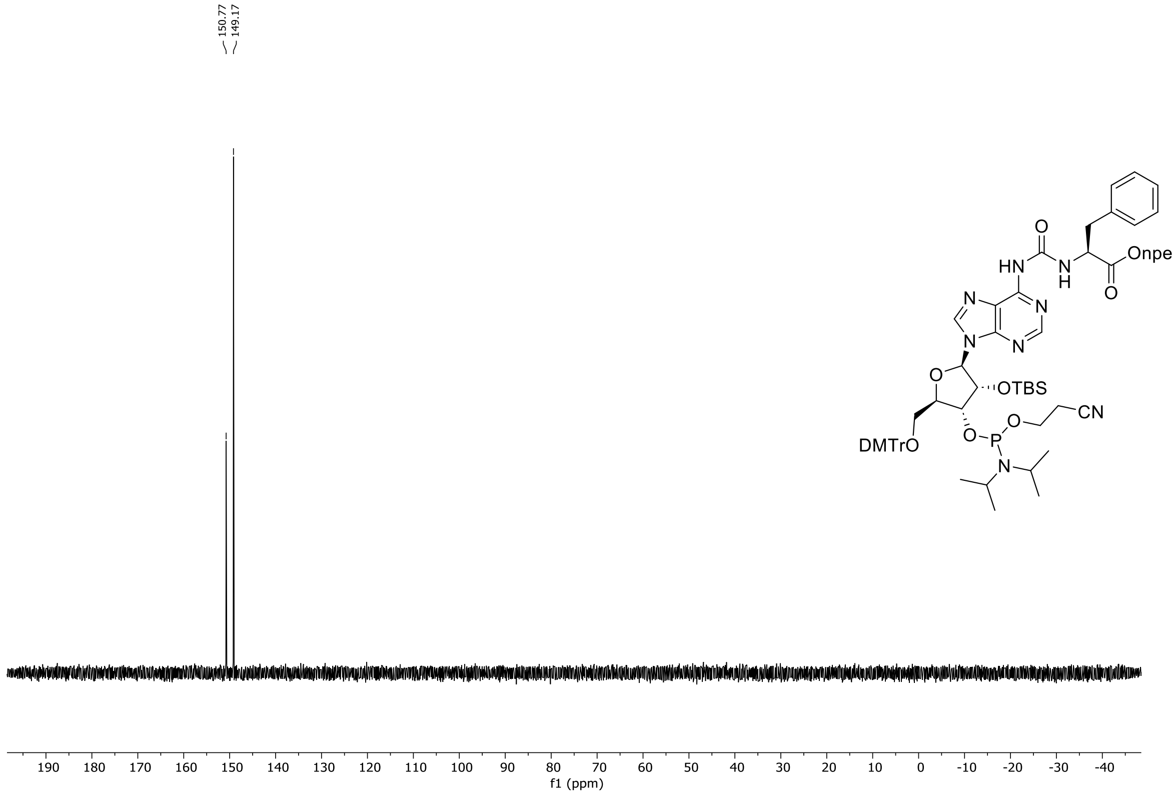

150.77
149.17
190
180
170
160
150
140
130
120
110
100
90
80
70
60
50
40
30
20
10
0
-10
-20
-30
-40
f1 (ppm)
HN
O
N
H
Onpe
O
N
N
N
N
O
OTBS
O
P
O
CN
DMTrO
N
